# Accurate tracking of the mutational landscape of diploid hybrid genomes

**DOI:** 10.1101/507830

**Authors:** Lorenzo Tattini, Nicolò Tellini, Simone Mozzachiodi, Melania Jennifer D’Angiolo, Sophie Loeillet, Alain Nicolas, Gianni Liti

## Abstract

**Background:** Genome evolution promotes diversity within a population via mutations, recombination, and whole-genome duplication. However, quantifying precisely these factors in diploid hybrid genomes is challenging. Here we present an integrated experimental and computational workflow to accurately track the mutational landscape of yeast diploid hybrids (MuLoYDH) in terms of single-nucleotide variants, small insertions/deletions, copy-number variants and loss-of-heterozygosity.

**Results:** Haploid *Saccharomyces* parents were combined into diploid hybrids with fully phased genome and controlled levels of heterozygosity. The resulting hybrids represented the ancestral state and were evolved under different laboratory protocols. Variant simulations enabled to efficiently integrate competitive and standard mapping, depending on local levels of heterozygosity and read length. Experimental validations proved high accuracy and resolution of our computational approach. Finally, applying MuLoYDH to four different diploids revealed striking genetic background effects. Homozygous *S. cerevisiae* showed ~4-fold higher mutation rate compared to *S. paradoxus*. In contrast, interspecies hybrids exhibited mutation rates similar to intraspecies hybrids despite 10-fold higher heterozygosity. MuLoYDH unveiled that a substantial fraction of the genome (~200 bp per generation) was shaped by loss-of-heterozygosity and this process was strongly inhibited by high levels of heterozygosity.

**Conclusions:** We report a comprehensive framework for characterizing the mutational spectrum of yeast diploid hybrids with unprecedented resolution, which can be generalised to other genetic systems. Applying MuLoYDH to laboratory-evolved hybrids provided novel quantitative insights into the evolutionary processes that mould yeast genomes.

## Introduction

High-throughput sequencing (HTS) technologies, both short- and long-read, have had a massive impact on genome research, enabling previously unimaginable and detailed dissection of the genomic landscape with outstanding speed and low costs [1, 2, 3]. The occurrence of variation in sequence, structure, and size of a genome in time is triggered by several factors including DNA mutations, recombination, and whole-genome duplication. These factors contribute to diversity within a population, translate into quantitative phenotypic variation and may eventually result in speciation. Integrated bioinformatic pipelines, along with high-quality reference assemblies, are fundamental to successfully depict the mutational landscape of genomes [4, 5]. However, *de novo* whole-genome assembly and phasing is still highly challenging and results in incomplete sequences [6, 7]. Thus, the mutational landscape of diploid or polyploid organisms has been characterized through resequencing studies which are based on mapping short reads against a single consensus reference, although the latter misses what defines the genetic identity of one individual [8]. For example, the human genome was assembled using the DNA of ~50 individuals with just one of them accounting for ~70% of the sequence, while the yeast reference genome was produced from a single laboratory strain (namely S288C) and its derivatives [9, 10]. Recently, high-quality panels of reference sequences [11, 12, 13] and novel standards for genome assembly [14] have been reported, while graph-based models have been suggested to overcome the limits imposed by reference bias [15, 16, 17]. Nevertheless, using a single reference sequence is a convenient simplification [8] and current technologies are boosting genome quality [18]. Resequencing studies have been proven successful whenever the level of heterozygosity is sufficiently low (e.g. the percentage of polymorphic loci in humans is < 0.16% [19]) or for homozygous genomes. Yet, mapping against a reference genome raises issues such as the impossibility of probing variation in genomic regions missing in the reference and no variant phasing information. The latter is a crucial point since current whole-genome sequencing methods do not provide phase information by default. In fact, current phasing methods rely on computational and experimental techniques that require trio data [20] or population-based statistical phasing and long reads to maximise the performance [21].

Natural diploid genomes harbour varying levels of heterozygosity [22]. Analysing diploid hybrid genomes, characterized by high heterozygosity, against a reference poses the problem of spurious read mapping, which in turn may lead to false positive calls of both single-nucleotide variants and small insertions/deletions (SNV and indels, respectively). High levels of heterozygosity allow for mapping short-read data against hybrid genome assemblies obtained by concatenating the two parental subgenomes (competitive mapping) [23, 24]. This strategy provides direct variant phasing but is risky whenever the number of heterozygous loci is low since it will result in genomic regions characterized by reads with non-unique mapping, preventing the assessment of small variants (namely SNVs and indels).

The study of the role of hybridization in species fitness is an active field of research in evolutionary biology [25]. Unfortunately, notwithstanding the importance of experimental and computational validation of the methods based on HTS data [26, 27, 28], none of the approaches tailored to the analysis of hybrid genomes has been automatized nor tested through simulations [23, 29, 30]. In this context *Saccharomyces cerevisiae*, along with its closely related species, is a leading-edge eukaryotic model system that has long been exploited in genetics, cell biology and systems biology [31, 32, 33]. The *S. cerevisiae* genome was the first fully sequenced eukaryotic genome [34], and more recently it has also played a crucial role in understanding key principles in evolutionary genomics [35, 36, 37]. Species from the *Saccharomyces* genus have been shown to be prone to intra- and interspecies hybridization [35, 38]. Hybridization occurs ubiquitously with natural hybrids associated with multiple fermenting environments [39, 40, 41, 42]. Outbreeding has also played an important role in shaping *S. cerevisiae* population structure with several groups of strains showing mosaic genomes that result from ancient admixtures of extant lineages [43].

The precise laboratory control of the sexual and asexual phases is a major strength of yeast genetics and enables to combine different haploid species and isolates into designed ancestral hybrids. These diploids can be evolved under various laboratory protocols such as return-to-growth (RTG) [29], adaptive evolution (AE) [44, 45], and mutation accumulation lines (MAL) [46, 47, 48, 49, 50]. RTG experiments generate genome-wide recombinant hybrids characterized by loss-of-heterozygosity (LOH) events. LOHs allow the expression of recessive alleles [50, 51, 52] as well as the formation of new combinations of haplotypes and provide an alternative approach for the analysis of complex traits. EE experiments quantify the preferential accumulation of pre-existing and *de novo* genetic variants that are selected in a controlled environment due to their contribution to organismal fitness. On the contrary, in MAL experiments a bottleneck of one or few individuals is imposed on a population, allowing for non-lethal mutations to accumulate with slight or no filtering by natural selection. Forcing population bottlenecks provides a means to evaluate mutational rates and signatures. Compared to fluctuation assay [53], MALs yield unbiased genome-wide estimations of the rates but, so far, they have been mostly restricted to laboratory strains, mutator backgrounds, and haploids or homozygous diploids. Thus, a global picture of the mutational landscape, including genetic background effects and a quantitative measure of the impact of LOH, is still missing. In this study, we present MuLoYDH, a general framework for the comprehensive characterization of the **Mu**tational **L**andscape **o**f **Y**east **D**iploid **H**ybrids in terms of SNVs, indels, copy number variants (CNVs), and LOHs. The genetic cross of haploid parents with fully assembled genomes enables to reconstruct a phased diploid genome that serves as the ancestral state. The latter is otherwise impossible to obtain from direct sequencing of hybrid diploids. After extensive benchmarking against both simulated and experimentally designed diploid *Saccharomyces* hybrids, we use MuLoYDH to accurately characterize intra- and interspecies MALs obtained by crossing domesticated and natural strains. Our strategy reveals striking genetic background effects and quantifies the genome-wide role of LOH in shaping the evolution of hybrid genomes.

## Results and discussion

### Overview of the MuLoYDH strategy

The MuLoYDH workflow begins with experimentally generating ancestral hybrids by combining two haploid founder strains with fully assembled and annotated genomes. This allows the investigation of the fully phased genome of the derived hybrids. *S. cerevisiae* is an ideal genetic system for this approach since haploid strains can be crossed to produce diploids with a broad range of heterozygosity (Figure 1a-b). Designed *Saccharomyces* diploids can range from complete homozygous (0%) when a single strain is used, low heterozygosity (0.1%) in intraspecies hybrids derived from strains of the same subpopulation, moderate (0.5-4%) crossing strains from diverged subpopulations and extremely high (8-35%) in interspecies hybrids [11, 22, 23].

**Figure 1.**
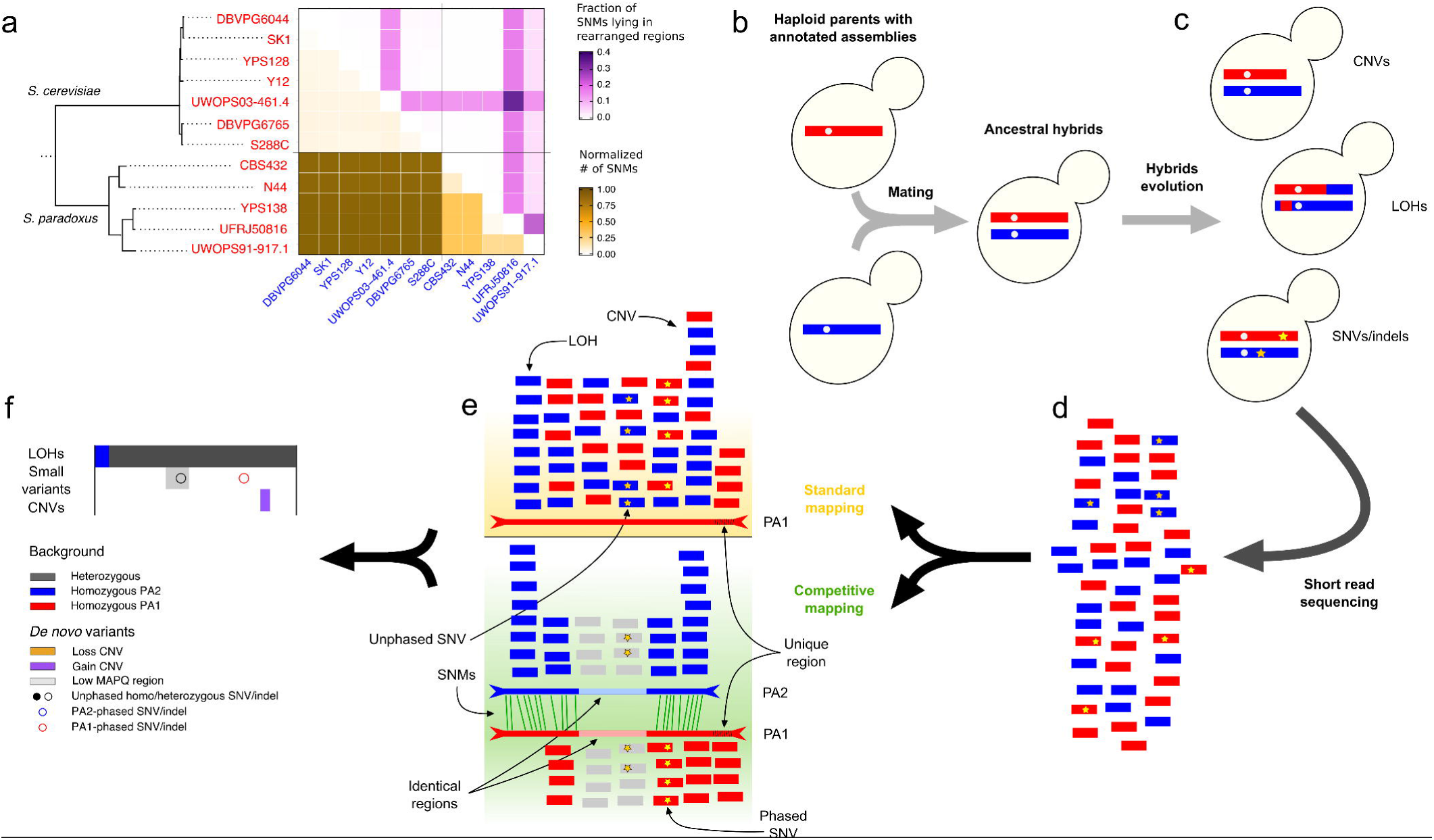
MuLoYDH overview. MuLoYDH comprises: founder parent strains characterization (a), hybrid generation (b) and evolution (c), resequencing of evolved strain (d) and tracking their mutational events (e-f). (a) Complete genome assemblies and annotations guide a rational selection of founder strains with desired genomic distances (bottom left heatmap) and inter-chromosomal rearrangements (top right heatmap). The phylogenetic tree is reproduced from Yue *et al.* [11]. (b) Selected parental strains are combined into ancestral hybrids. The genetic crossing eludes the problems in assembling and phasing diploid genomes. (c) Ancestral hybrids are evolved and accumulate different types of *de novo* variants. (d) Genomes from evolved hybrids are sequenced at high coverage by short-read sequencing. (e) Short reads are mapped against the assemblies of the two parental genomes separately (standard mappings) and against the concatenation of the two assemblies (competitive mapping). For simplicity, the standard mapping against only one parental assembly (PA) is reported. In the competitive mapping, reads from parent 1 (red) are expected to map to the assembly of parent 1 (PA1) on the basis of the presence of SNMs (green lines). Conversely, reads from parent 2 (blue) are expected to map to the assembly of parent 2 (PA2). Regions bearing no SNM due to high sequence identity are characterized by reads with MAPQ = 0 (light grey reads). These regions are probed for small variants from standard mapping, without direct phasing, against one parental assembly (light red region in PA1), whereas the other one is masked (light blue segment in PA2). (f) Small variants obtained from competitive and standard mappings are combined into a single set of calls. CNVs and LOHs are called from standard mappings.

Following mitotic hybrid evolution under different defined laboratory conditions (Figure 1c), the corresponding short-read data can be mapped using both *competitive* and standard approach (Figure 1d-e). The former consists in mapping short-read data against the union of the two parental assemblies, whereas the latter refers to mapping against a single parental assembly. The computational strategy implemented in MuLoYDH for tracking the mutational events relies on the presence of single-nucleotide markers (SNMs) between the two parental subgenomes (Additional file 1: Figure S1-S4). The genomic density and distribution of SNMs are fundamental for our purposes since SNMs are probes for LOH detection and, as detailed in the following section, they allow direct phasing of small variants from competitive mapping. In addition, SNMs genomic positions are determined from the assemblies and can be used to set up a rational quality threshold for LOH detection as well as for filtering *de novo* small variants (see Methods). As expected, the number of SNMs detected aligning *S. paradoxus/S. cerevisiae* assemblies was ~15-fold higher compared to *S. cerevisiae/S. cerevisiae* assemblies (Table 1). SNMs were classified as lying in collinear regions or lying within structural rearrangements, namely inversions or translocations, and the corresponding fractions were calculated (*f*c and *f*r, respectively). Using these values we were able to further differentiate the backgrounds beyond the typical heterozygosity measures simply based on sequence divergence. Hybrids derived from the UWOPS03-461.4 and UFRJ50816 showed the largest fraction of SNMs within structurally rearranged regions (Figure 1a), consistently with massive genomic rearrangements occurring within these lineages [11].

**Table 1.** Statistics of SNMs for the constructed hybrids. *S.c.* and *S.p.* refers to *S. cerevisiae* and *S. paradoxus* respectively. For each cross we report: number of SNMs, the genome-wide percentage of SNMs, the fraction of SNMs lying in collinear regions (*f_c_*), fraction of SNMs lying in rearranged regions (*f_r_*) and low-marker-density-regions (LMDRs) fractions. The latter is the fraction of core genomic regions characterized by less than one marker in 300 bp, calculated from pairwise alignment of different pairs of assemblies.

| Species | Background | Number of SNMs | SNMs % | $f_c$ | $f_r$ | LMDRs fraction | |
| --- | --- | --- | --- | --- | --- | --- | --- |
|  |  |  |  |  |  | Assembly 1 | Assembly 2 |
| *S.c./S.c.* | SK1/S288C | 75547 | 0.62 | 0.98 | 0.02 | 0.32 | 0.32 |
|  | UWOPS03-461.4/YPS128 | 63926 | 0.54 | 0.81 | 0.19 | 0.30 | 0.31 |
|  | YPS128/DBVPG6765 | 78064 | 0.66 | 0.99 | 0.01 | 0.24 | 0.24 |
| *S.p./S.c.* | N17/DBVPG6765 | 1095399 | 9.19 | 0.96 | 0.04 | 0.11 | 0.11 |

The SNMs distribution represents a key feature of the hybrid genome and highly impacts the accuracy of *de novo* variants detection. We calculated the low-marker-density-regions (LMDRs) fraction, i.e. the fraction of genomic regions characterized by less than one marker in 300 bp, namely twice the read length of the sequencing experiments discussed in this study (Table 1). Pairs of genomes characterized by a small number of SNMs showed higher values of the LMDRs fraction. MuLoYDH can be run in two different settings (collinear/rearranged) exploiting *a priori* knowledge of parental genomes reciprocal structure. In the collinear mode SNMs are determined chromosome-by-chromosome, aligning a chromosome of parent 1 against the corresponding homologous from parent 2, whereas with the rearranged option they are calculated through a single whole-genome alignment of parental assemblies. Running MuLoYDH in collinear mode provided a larger number of SNMs with a uniform distribution along the genome compared to the rearranged mode (Figure S5).

The fully phased hybrid genome assembly can be exploited to perform a competitive mapping of the reads obtained from evolved hybrids. This approach, compared to a standard mapping against a single assembly or an unphased reference genome, is expected to provide a larger number of mapped reads in unique regions that belong only to one of the parental assemblies. Indeed, competitive mapping in *S.c*. hybrids reduced the number of unmapped reads of ~8% on average (see Additional file 2 — Table S1). At the same time, it represents a challenge regarding reads mapping to identical regions within the two parental assemblies. In fact, the fraction of reads showing a mapping quality (MAPQ) value equal to zero [54], thus reflecting non-unique mapping, is also expected to increase as the level of heterozygosity decreases. As expected, the number of reads showing MAPQ = 0 increased in the competitive mapping (~43%) with respect to the standard mappings (~15%) in intraspecies hybrids (see Additional file 2 — Table S2). In summary, the crossing phase did provide the unique opportunity to generate diploid hybrids with phased genome to benchmark computational approaches for studying their evolution.

### Benchmarking MuLoYDH against simulated datasets

Highly similar DNA sequences may occur on different genomic scales, from short stretches (such as homopolymers), to complex events (e.g. segmental duplications), up to chromosome level (i.e. homologous chromosomes) [55]. These repetitive sequences are characterized by nearly 100% sequence identity and represent a major challenge of HTS data analysis [56]. The number and the distribution of SNMs affect the performance of small variants calling from competitive mapping. As the number of SNMs and the level of heterozygosity decrease, the mapping algorithm produces a progressively increasing number of reads characterized by MAPQ = 0. In turn, this affects the small variants calling algorithm, since reads characterized by non-unique mapping (i.e. MAPQ = 0) are filtered out. Thus, we investigated the impact of the level of heterozygosity on the performance of MuLoYDH in calling small variants from competitive mapping in simulated genomes (Figure 2a-b). The number of simulated SNMs was chosen to mimic experimental data (Table 1). As expected, the *F*_1_ score sharply decreased with the number of SNMs. Nevertheless, when the percentage of SNMs was ~0.5 (as further discussed in the following paragraph), the score tended to a value close to 1 (see Additional file 1: Figure S6).

**Figure 2.**
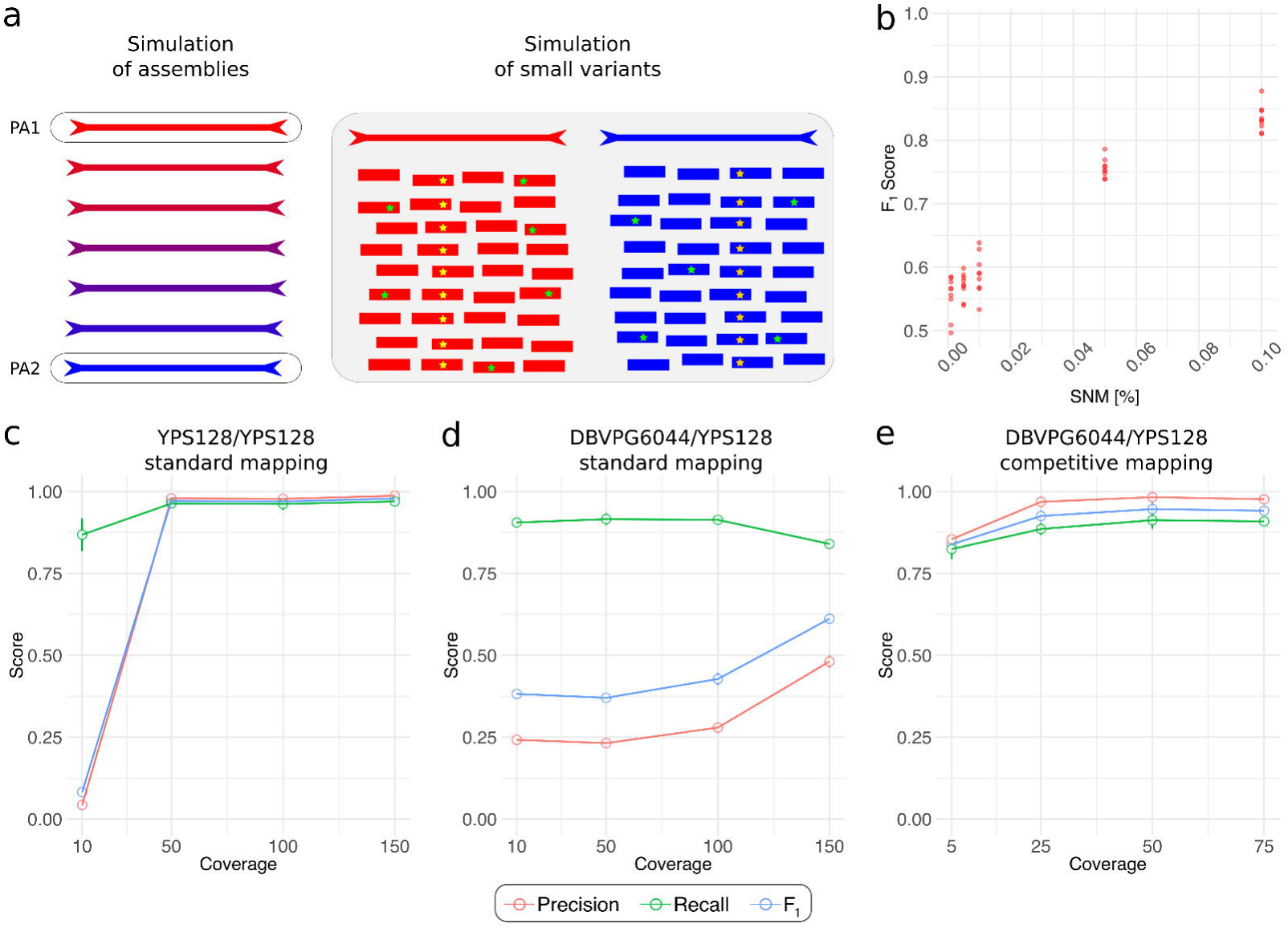
High-accuracy variants detection. (a) Given two assemblies, various levels of heterozygosity are simulated by progressively replacing the polymorphisms in parental assembly 1 (PA1) in the corresponding nucleotides of the assembly of parent 2 (PA2). Small variants (yellow stars) are simulated in known genomic positions to assess the performance of variant calling algorithms from competitive and standard mapping separately with Illumina reads bearing sequencing errors (green stars). (b) The overall performance (*F*_1_ score, see Methods) of small variants detection with competitive mapping decreases with the number of SNMs. For each SNM percentage value, 3 hybrid genomes were simulated from the DBVPG6765/YPS128 hybrid with 3 different replicates of short reads, carrying different variants. The lowest coverage showing *F*_1_ score saturation (25 x, see panel e) was chosen to generate the short-read dataset. DBVPG6044 and YPS128 assemblies are exploited to compare the performance of variant calling from competitive and standard mappings separately. Precision (red), recall (green) and *F*_1_ score (blue) are reported as a function of coverage for: (c) YPS128 complete homozygous diploid data with standard mapping, (d) DBVPG6044/YPS128 heterozygous diploid data with standard mappings against both parental assemblies, and (e) DBVPG6044/YPS128 heterozygous diploid data with competitive mapping. Since we compared the same set of reads in the three approaches, competitive mappings show half of the coverage with respect to standard mappings. Solid lines serve as an eye guide.

Since the competitive approach has never been systematically benchmarked on a dataset of simulated variants and inconsistencies among small variants callers have been reported [57, 58, 59, 60, 61, 62], we compared the performance of calling small variants, with both SAMtools and FreeBayes, from competitive and standard mappings as a function of the coverage (see Methods). As expected, using standard mapping for complete homozygous diploids, the *F*_1_ score increased with coverage showing saturation at 50 x (Figure 2c). On the contrary, calling small variants from heterozygous diploid data mapped with the standard approach provided low *F*_1_ score with few benefits by increasing coverage (Figure 2d). This effect is explained by spurious mapping of reads from parent 2 against the assembly of parent 1 (and vice versa) which leads to false positives (*FP*s). In fact, the low *F*_1_ score can be ascribed to low precision. Instead, the competitive mapping of heterozygous diploid data (Figure 2e) yielded a large number of true positives (*TP*s) and high *F*_1_ score, with a trend similar to the results (Figure 2c) obtained from the standard mapping of the complete homozygous diploid. Therefore, competitive mapping can be exploited to call small variants with direct phasing, although the overall performance is limited by the number of false negatives (*FN*s) (see recall in Figure 2e). Thus, we included in MuLoYDH a module that automatically calculates the boundaries of regions characterized by reads with low mapping quality (i.e. MAPQ < 5). These regions are investigated through standard mapping. Although this prevents direct variant phasing, it allows for testing the presence of small variants in the whole accessible regions of the genome.

Moreover, using DBVPG6765/YPS128 hybrid data, we calculated the *F*_1_ score considering only the variants lying within a 65 kb unique region of the DBVPG6765 strain on chromosome XV, derived from horizontal gene transfer from *Torulaspora microellipsoides* [63]. We obtained *F*_1_ = 0.96 (*TP* = 14, *FN* = 1, *FP* = 0) on the basis of 15 variants (14 SNV and a 1 bp insertion) combining all the simulated short-read data (20 experiments). Hence, MuLoYDH allows for calling small variants in regions which are not reported in the reference genome.

Another aspect of the small variants calling procedure is whether MuLoYDH can correctly genotype variants within LOH regions. In fact, these regions may carry homozygous variants (occurred before LOH) and heterozygous variants (occurred after LOH). Thus, we compared the genotypes of simulated variants with those reported by MuLoYDH, in DBVPG6765/YPS128 hybrids. MuLoYDH correctly called and genotyped 1840 variants in the simulated LOH regions (691 homozygous and 1149 heterozygous variants), producing 62 *FP*s and 207 *FN*s (*F*_1_ = 0.93 *±* 0.01) with, as expected, a larger number of missed events in heterozygous state (121 heterozygous vs 86 homozygous). Details are reported in Additional file 1: Figure S7.

Overall, these results demonstrate that both competitive and standard mapping are required to maximise small variants calling performance. Competitive mapping provides direct variant phasing although it can be used only in regions characterized by a sufficient number of SNMs, while standard mapping is locally necessary if no SNM exists in regions larger than twice the read length.

### Applying the MuLoYDH workflow to a mutator strain

We next applied MuLoYDH to a SK1/BY hybrid with a mutator background (*tsa1*Δ/*tsa1*Δ, see dataset 1 in Methods) evolved for 25 consecutive single-cell bottlenecks [49, 64]. This hybrid evolved drastically from its ancestral state, thus providing a challenging test for our workflow. It accumulated both *de novo* small variants and a series of complex LOH events (Figure 3a-b and Additional file 3: Table S2). MuLoYDH provided a robust genotyping approach of SNMs positions determined by aligning the parental assemblies (Figure 3c). Exploiting high-quality SNMs, genotyped against both parental assemblies (see Methods), 43 LOHs ranging from 97 bp to 591 kb were detected.

**Figure 3.**
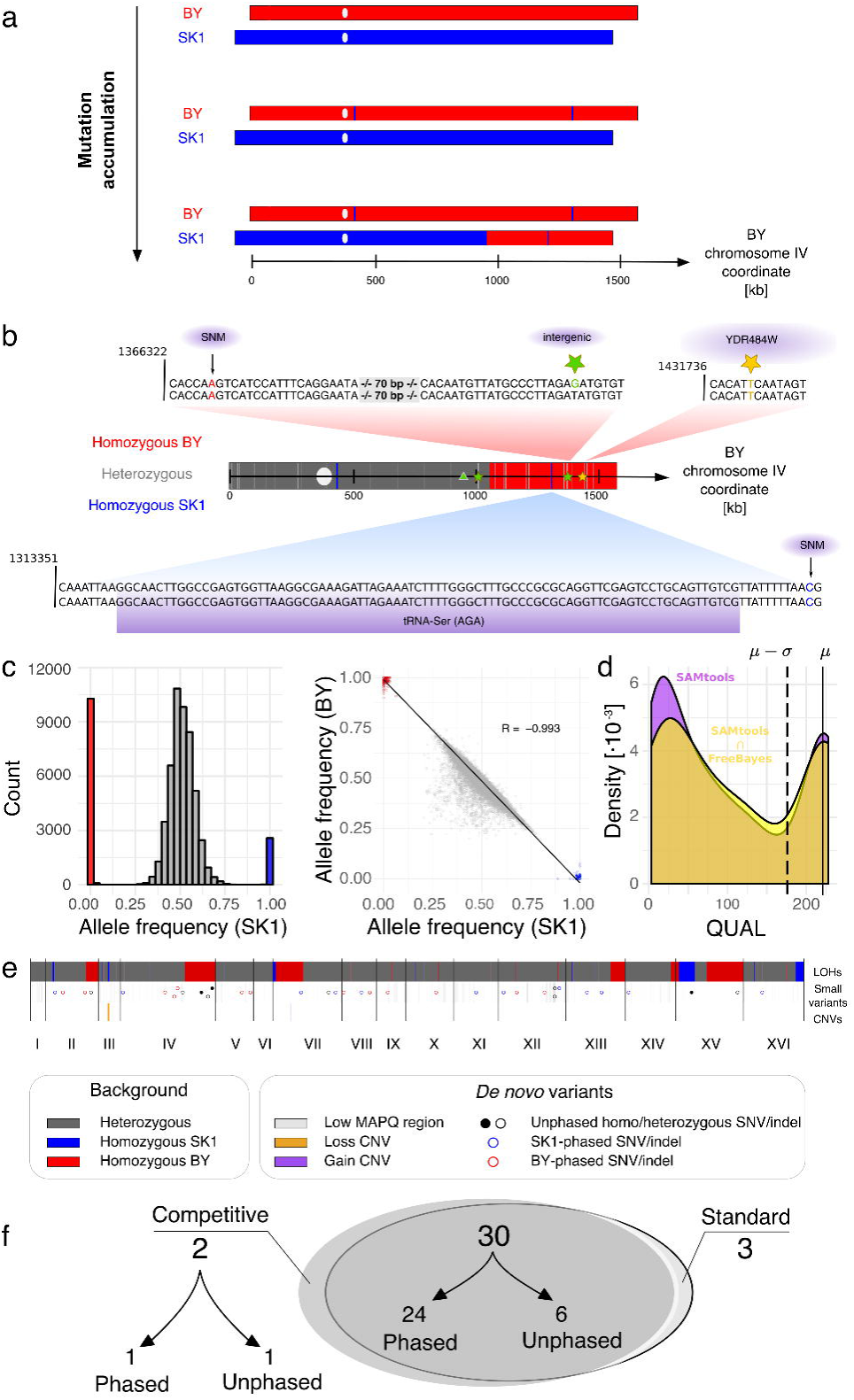
Genome evolution of a mutator hybrid. MuLoYDH provides accurate tracking of the mutational landscape in a SK1/BY *tsa1*Δ*/tsa1*Δ MAL. (a) Hybrid evolution leads to LOH and (b) to small variants. MuLoYDH performs small variants call on the basis of the LOH regions detected. SNVs and indels are automatically annotated. One homozygous (yellow stars) and one heterozygous (green star) SNVs were detected within LOH regions on chromosomes IV (red: BY, blue: SK1, dark grey: heterozygous segments; white oval: centromere). The presence of SNMs (black arrows) allows direct variant phasing through competitive mapping. A 1-bp deletion was detected in a heterozygous segment and phased to the BY chromosome (green triangle). One heterozygous SNV (green star) was detected from standard mapping (light grey segment). (c) The strategy implemented in MuLoYDH for the detection of LOHs allows noise mitigation (see also Additional file 1: Figure S10) as shown by the clear separation of genotypes with different allele frequencies and by the high negative correlation of allele frequencies. R is the Pearson correlation coefficient. Red (blue) dots/columns refer to homozygous BY (SK1) SNMs, while grey dots/columns refer to heterozygous SNMs. SNMs are filtered on the basis of their quality values. (d) The same strategy is applied to *de novo* small variants. Variants with a quality value (QUAL) *< µ −σ* are masked. *µ* is the mean quality and *σ* is the standard deviation calculated from SNMs. Kernel density estimations are calculated from quality values of SAMtools calls (purple) and the intersection of SAMtools and FreeBayes calls (yellow). (e) The genome-wide mutational landscape includes CNVs: one gain event in chromosome VII (3 BY copies) and one loss event of the BY allele in chromosome III. (f) Small variants detected from competitive and standard mapping are reported in the Venn diagram. Variants from competitive mapping are classified as phased and unphased. The latter were all detected within LOH regions.

We further characterized the mutational spectrum which included 34 SNVs, 1 indel and 2 CNVs (Figure 3d-f). 24 out of 34 SNVs were phased (12 to SK1, 12 to BY) as well as the indel that occurred on the BY chromosome IV. The remaining 10 SNVs were called without phasing. 6 of them were detected in BY LOH regions (4 heterozygous, 2 homozygous), 1 in SK1 LOH regions (homozygous), while 3 variants were called from standard mapping. For validation, 11 variants (3 phased and 8 unphased) were tested through PCR and Sanger sequencing. All of them were validated as true positives. 6 out of 8 unphased variants were heterozygous, whereas 2 SNVs, lying in LOH regions, were genotyped as homozygous and thus further supported that it resulted from a mitotic recombination event. We detected a short LOH segment (SK1 allele, 473 bp), supported by 4 SNMs bearing the tRNA-Ser (AGA) gene. The SK1 LOH region lay within a large (*>* 450 kb) BY LOH region.

The latter carried a validated homozygous missense variant (C *→* T, YDR484W, see Figure 3b, yellow star) that likely occurred before the large event. Remarkably, one validated intergenic heterozygous variant (Figure 3b, green star) lay within the aforementioned LOH region (BY chromosome IV). This mutational status suggests that a recombination event led to a short LOH followed by: the occurrence of a SNV (Figure 3b, yellow star), a larger LOH, and finally one heterozygous SNV (Figure 3b, right-most green star). Annotated electropherograms with validated variants and LOH markers are reported in Additional file 1: Figure S8-S9. Altogether, these results demonstrate that the SNMs quality-filtering approach for LOH detection provided accurate results also for events supported by few markers (Additional file 1: Figure S10). In addition, validations of *de novo* small variants showed that the combination of two callers (both applied to competitive and standard mappings) and the filtering strategy proposed yielded reliable tracking of genomic variants (Figure 3d).

### Evolution through complex copy-number variants

Changes in copy-number, from single gene to whole chromosome events, have been observed in both natural and laboratory evolved strains [65, 66, 67, 68]. MuLoYDH produces CNV calls through Control-FREEC [69] normalizing the read count (RC) signal for GC-content and mappability [70]. We tracked genome evolution in the intraspecies UWOPS03-461.4/YPS128 *S. cerevisiae* hybrid evolved via the RTG protocol (see dataset 2 in Methods) [29]. A large fraction of the ancestral hybrid genome is non-collinear, due to a massive genome instability occurred in the Malaysian lineage (UWOPS03-461.4) [11]. In particular, the UWOPS03-461.4 chromosome VIII consists of a 350 kb collinear region that spans the centromere and a 390 kb translocation derived from chromosome VII, while the UWOPS03-461.4 chromosome VII is a complex mosaic harbouring two distinct regions from chromosome VIII (Figure 4a). Recombination between non-collinear homologous chromosome potentially results in complex CNVs.

**Figure 4.**
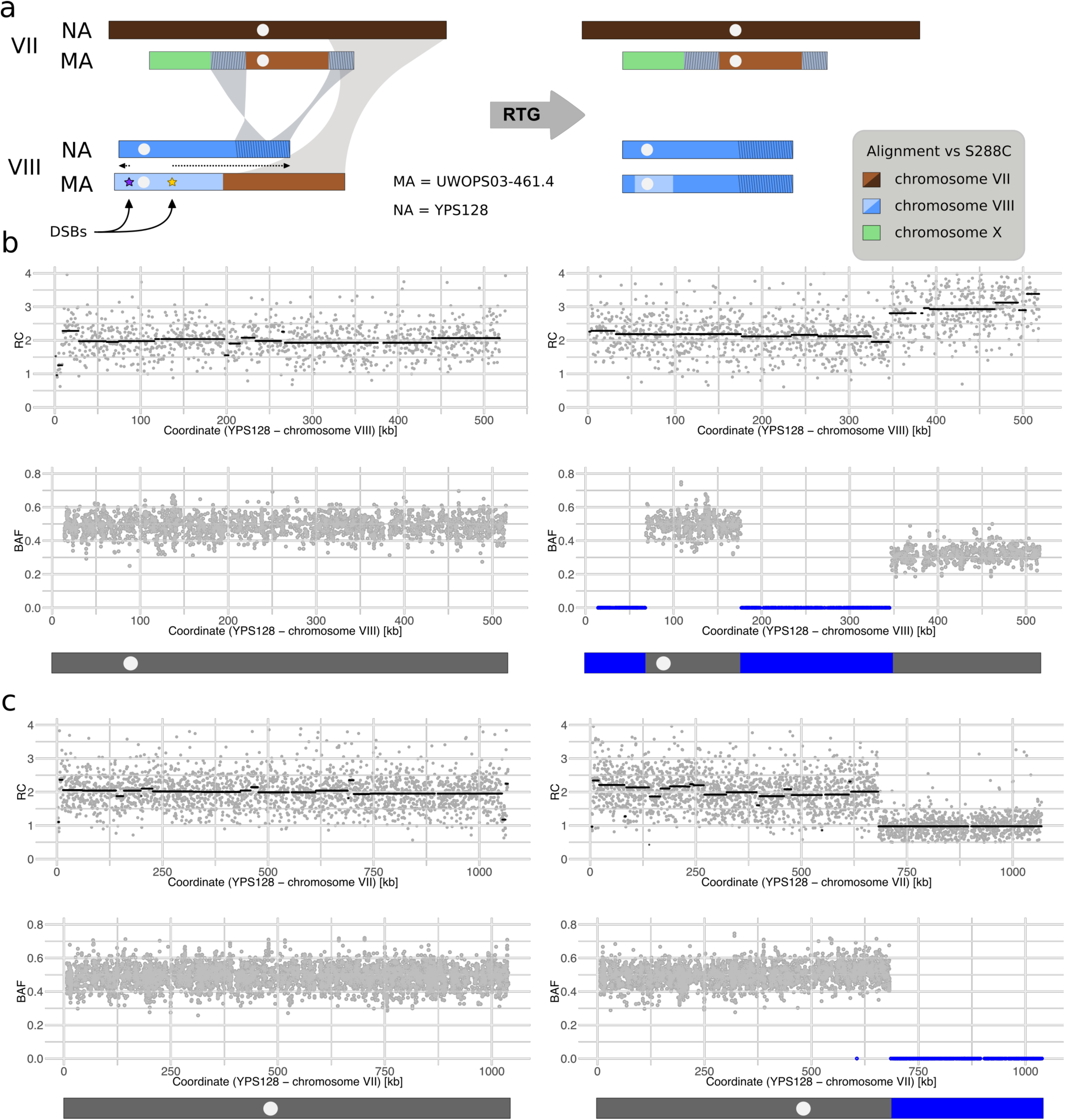
Resolving complex CNVs. (a) Ancestral and RTG-evolved karyotype in a hybrid obtained by crossing UWOPS03-461.4 (a *S. cerevisiae* Malaysian strain, MA) with YPS128 (a *S. cerevisiae* North American strain, NA). Light and dark colors encode for MA and NA alleles respectively. Chromosomal identities were assigned by homologous centromeres. Double-strand breaks in the UWOPS03-461.4 chromosome VIII (purple and yellow stars) were repaired (dotted arrows) using the homologous chromosome from YPS128. These chromosomes bear collinear regions and large inter-chromosomal rearrangements, including a large inversion. Chromosomal identities were determined by centromeres. Repairing a DSB (purple star) in a collinear region with the homologous chromosome leads to a terminal LOH. Repairing a DSB (yellow star) in a chromosome arm bearing collinear and rearranged regions leads to an interstitial LOH and to a complex CNV. (b) Chromosome VIII read count data support the presence of 3 copies of a large segment on the right arm. BAF data from SNMs using mapping against the YPS128 assembly support the presence of a 3-copies region (2 copies of YPS128 and 1 copy of UWOPS03-461.4). The two stretches of BAF values at zero refer to the terminal and interstitial LOH regions (both YPS128 alleles). The former results from DSB repair (purple star, panel a) in chromosome VIII-L, while the latter is the outcome of DSB repair (yellow star, panel a) in chromosome VIII-R. (c) Read count signal and BAF data support the deletion of a large segment of UWOPS03-461.4 chromosome VII-R.

The combination of CNV profiles and B-allele frequencies (BAF) of SNMs, both calculated from standard mapping, shed light on complex events. The UWOPS03-461.4/YPS128 hybrid showed multiple LOHs with two events occurring at both arms of chromosome VIII. Two double-strand breaks (DSBs) likely occurred in the UWOPS03-461.4 chromosome VIII (Figure 4a, purple and yellow stars) and were repaired using the homologous YPS128 chromosome VIII region. Chromosome VIII-L repair occurred within the collinear region and resulted in a simple LOH event without an associated CNV. The same holds for the collinear region in chromosome VIII-R spanning from the DSB to the breakpoint of the chromosome VIII/VII translocation. These regions showed RC ≃ 2 and BAF ≃ 0 (Figure 4b). In contrast, the rearranged chromosome VIII region embedded in the LOH was subjected to CNV, as shown by RC ≃ 3 and BAF ≃ 0.3. The latter supported the presence of 2 copies of YPS128 alleles and 1 copy of UWOPS03-461.4 allele. This complex genomic configuration was further confirmed by the RC signal and the BAF data from chromosome VII, given the loss of the UWOPS03-461.4 chromosome VII translocated regions (Figure 4c and Figure S11). Thus, combining the knowledge of the exact ancestral chromosomal configuration with our computational framework enabled to dissect a series complex genomic events.

### Mutational rates across different genetic backgrounds

Mutational rates and signatures are key parameters for genome evolution but how these vary across natural genetic backgrounds has remained largely unexplored. We applied the MuLoYDH workflow to investigate the effect of genetic background on mutational rates. We constructed four yeast diploids that enabled multiple comparisons (see dataset 3 in Methods). We used a single *S. cerevisiae* background (DBVPG6765) and crossed it to itself, to a different subpopulation of the same species (*S. cerevisiae* YPS128) and to a different species (*S. paradoxus* N17). This resulted in three diploids with approximately 0%, 0.5% and 10% heterozygosity, that enabled the investigation of the effect of heterozygosity in laboratory evolution experiments. We also generated a complete homozygous *S. paradoxus* N17 diploid to compare *S. cerevisiae* to its closely related species. We performed MALs experiments using 8 replicated lines for each of the 4 diploids subjected to 120 consecutive single-cell bottlenecks. The corresponding number of generations was estimated measuring the colony cell population size, observing minimal differences between the four hybrids (see Addition file 1: “Number of generations and mutation rates in MALs”). The mutation rates of SNVs, indels, CNVs and LOHs (the latter only for the heterozygous hybrids) are reported in Figure 5a. Detailed results are reported in Additional file 1 while the lists of *de novo* variants are reported in Additional files 4-7: Tables S3-S6. The SNVs mutation rate in *S. cerevisiae* was consistent with previous reports estimated using diploid laboratory strains, indicating no major differences in our Wine/European background [50]. However, we surprisingly observed a 4-fold lower (*p*-value < 0.0005, Welch’s *t*-test) mutation rate in *S. paradoxus* (95% CI *t*-distribution: [3.39, 11.1] · 10^−11^ bp^−1^) compared to *S. cerevisiae* (95% CI *t* distribution: [1.98, 3.67] ·10^−10^ bp^−1^). The same trend was detected for indels and CNVs. This result suggested that the lower mutation rate of *S. paradoxus* might have contributed to the slower evolution of this species compared to *S. cerevisiae*, as shown by the overall branch length differences observed between the two species since the split from their last common ancestor [11, 71]. The SNVs mutation rate in the two heterozygous hybrids was also slightly different (*p*-value = 0.016, Welch’s *t*-test) with the highly heterozygous interspecies hybrid DBVPG6765/N17 showing higher rate (95% CI *t*-distribution: [1.73, 2.74] · 10^−10^ bp^−1^) compared to the intraspecies hybrid YPS128/DBVPG6765 (95% CI *t*-distribution: [1.09, 1.89] ·10^−10^ bp^−1^). We also compared the mutation rates of different classes of variants for both the intra- and interspecies hybrids. As reported in Figure 5a, the mutation rate calculated for indels and CNVs was one order of magnitude smaller compared to SNVs and LOHs.

**Figure 5.**
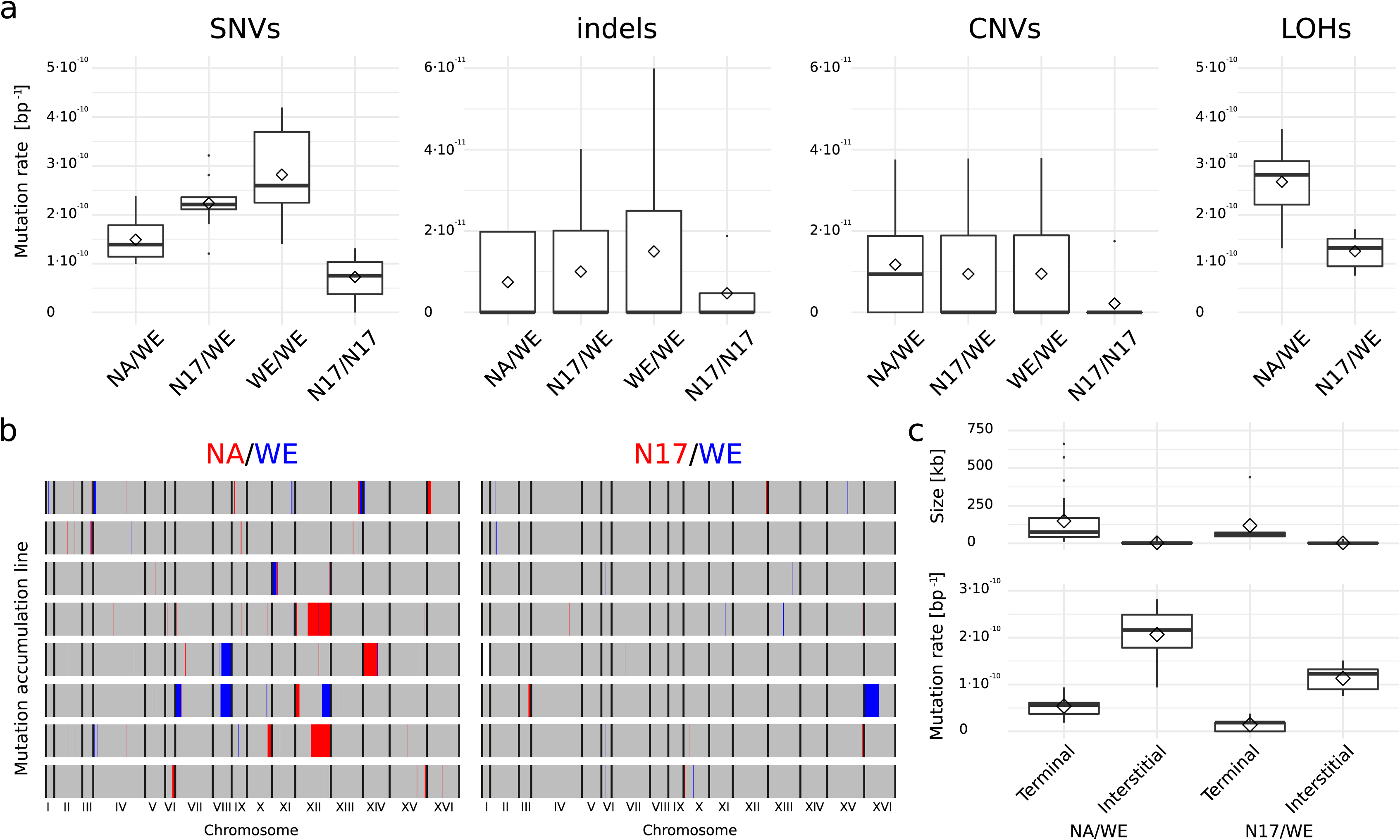
Mutation rates in different backgrounds. (a) Box plots of nuclear mutation rate per generation in different homozygous and heterozygous diploid backgrounds for SNVs, indels, CNVs and LOHs. Rates were calculated from 8 MALs per background propagated for ~2250 generations (120 bottlenecks). Diamonds represent mean values. (b) LOH events in YPS128/DBVPG6765 (NA/WE) and N17/DBVPG6765 (N17/WE) hybrids. LOHs are depicted in red and blue. Heterozygous regions are represented in grey. White regions were masked due to aneuploidies. (c) Box plots of size and mutation rates for terminal and interstitial events. Diamonds represent mean values.

The two heterozygous hybrids showed a substantial difference in term of LOH rates (Figure 5b), while both were characterized by a large fraction of small interstitial events and few large terminal LOHs (Figure 5c). Mitotic recombination events are rare and usually require a selection method to be detected [72]. Nevertheless, given the large number of generations performed in our study, we were able to observe a considerable number of LOH events in hybrids. Intraspecies hybrids showed a larger number of LOHs (111) compared to the interspecies hybrids (53) along with (I) a 5-fold higher fraction of the genome in LOH (0.02 ± 0.01 and 0.004 ± 0.007, respectively), and (II) a larger number of large events (> 25 kb), namely 3 and 0.4 on average per sample (Additional file 4: Table S3). Thus, the mitotic recombination rate was higher in *S. cerevisiae/S. cerevisiae* compared to *S. paradoxus/S. cerevisiae* crosses (*p*-value < 0.001, Welch’s t-test). Heatmaps of the detected events are reported in Additional file 1 (Figure S12) and the corresponding LOH segments are shown in Additional file 8. Overall, these results provided a quantitative genome-wide measure of the importance of LOHs in shaping polymorphisms patterns in diploid hybrid genomes and how this process was strongly inhibited by very high heterozygosity.

## Conclusions

Although HTS has proven revolutionary for genomic sciences, resequencing studies show intrinsic limits, particularly in the context of hybrid genomes. Variation graphs will help overcoming this deficiency although they will require extensive efforts to exhaustively support the shift to the new graph-based paradigm [8, 17]. Long-read sequencing is a valuable approach to provide novel reference genomes by means of *de novo* assembly. The availability of novel reference genomes opens new perspectives on resequencing approaches, allowing for investigations of the genomic mutational landscape with unprecedented resolution via short-read experiments. Still, current methods for assembling and phasing diploid genomes are costly and yield to limited contiguity [7]. Recently, long-read sequencing has been exploited to evaluate the performance of small variants callers through a synthetic-diploid benchmark [73]. Here we extended the benefits of long-read sequencing beyond synthetic diploids to evolved *Saccharomyces* hybrids. MuLoYDH provides a framework for studying genome dynamics by tracking the mutational landscape of designed yeast diploid hybrids and it may be extended to non-clonal populations. Moreover, as the sequencing technologies enhance both read length and accuracy, they will soon allow to produce fully assembled and phased natural hybrid diploid genomes. At this stage, the experimental approach implemented in MuLoYDH will be possibly bypassed while our computational strategy will be readily appropriate for the application to natural genomes.

MuLoYDH was developed to take advantage of the fully phased diploid genome assembly as ancestral state and Illumina short reads from the evolved hybrids. Haploid parents were combined into diploid hybrids with varying levels of heterozygosity which were evolved in different laboratory settings. Remarkably, the hybridization process can be easily automated [74] and as the number of available annotated assemblies increases, the number of potential hybrids grows quadratically. The presence of single-nucleotide markers provided a reliable quality threshold for filtering *de novo* small variants, thus bypassing the need of multiple hard filters. Moreover, it enabled the phasing of *de novo* SNVs, indels and CNVs as well as the precise characterization of LOHs. Our method was designed to yield a quantitative measure of the fraction of the genome which cannot be probed for direct phasing through competitive mapping and to perform variant calling in these regions by means of a standard approach based on a single reference. It was devised to resolve the drawbacks of using a single consensus reference for the analysis of diploid hybrid genomes, namely: (I) spurious read mapping, which may lead to false positive calls of both SNV and indels, (II) the impossibility of probing variation in genomic regions which are not reported in the reference, as well as (III) impracticable direct variant phasing. The latter has been the focus of several computational studies (based on a consensus reference) but solving exactly the problem is demanding since it is a NP-hard problem [7]. Experimental validation of the variants detected by MuLoYDH in a mutator MAL showed that it can be used to trace the time course of mutation occurrence with direct phasing information. Moreover, the analysis of the UWOPS03-461.4/YPS128 hybrid proved that MuLoYDH can be used to dissect complex genomic events.

While several studies focused on the mutation rates in haploid and complete homozygous diploid *S.c.* laboratory strains [46, 48, 47, 50], we used MuLoYDH to track the mutational landscape of 4 MALs with varying levels of heterozygosity, providing the first measurement of mutation rates in *S. paradoxus*. Surprisingly, we observed low mutation rates in the natural *S. paradoxus* N17 homozygous diploids compared to *S. cerevisiae* DBVPG6765. Mutation rates rates in inter- and intraspecies hybrids revealed that SNVs and LOHs in particular, are major sources of genomic variability that play a key role in genome evolution. Our study can be extended to other *Saccharomyces* hybrids, encompassing the whole spectrum of heterozygosity, and to complex genomes bearing structural rearrangements or characterized by ploidy *>* 2 [22, 75].

## Methods

### Simulated data

#### Simulations of hybrid genomes with varying levels of heterozygosity

Diploid genomes with varying levels of heterozygosity were simulated by custom R scripts, modifying the number of SNMs between the two parental subgenomes. Given two input assemblies (DBVPG6765 and YPS128), SNM positions were determined by MUMmer (NUCmer) [76]. Decreasing values of SNMs percentage were obtained by progressively replacing the allele of assembly 1 with the corresponding allele, as determined by NUCmer, of assembly 2 in known positions. The substitution step was repeated in order to provide different levels of SNMs (0.5%, 0.1%, 0.05%, 0.01%, 0.005%, 0.001%). For each SNMs value, 3 replicated assemblies were simulated.

#### Simulations of short reads for heterozygous hybrids

Simulated paired-end short reads were generated using the DWGSIM package [26]. In order to produce simulated short-read data from genome assemblies, two input reference assemblies were concatenated to produce a single multi-FASTA, which was sampled to build simulated paired-end (150 bp, insert size 500 bp) Illumina experiments with different coverage levels (10, 50, 100, 150 x). The mutation rate was set to 10^−5^ with the purpose of balancing a relevant number of small variants (~240 per genome) with the storage and the computational resources required for data processing. All the simulations were performed using the following parameters: 0.01 error rate for both forward and reverse read, and 0.1 indel/SNV ratio (according to estimations from Illumina data). Base quality parameters were set according to the experimental data reported in this study. Each simulation was performed in 5 replicates. The command line is reported in Additional file 1.

#### Simulations of short reads for hybrids bearing LOH regions

Short-read data of DBVPG6765/YPS128 hybrids bearing LOHs (with DBVPG6765 alleles) were obtained using DWGSIM with heterozygous genomes with the exception of chromosome I for which two copies of the FASTA sequence of DBVPG6765 were used as input. In order to have a robust statistic, the mutation rate was set to 10^−3^ for chromosome I and to 10^−5^ for all the other chromosomes. The average coverage was set to 50 x on the basis of the short-read simulations (see Figure 2e). 10 replicates were produced. All the other parameters were set as described above. Overall, we simulated 2304 variants in heterozygous regions of the genome and 2081 in LOH regions (787 homozygous and 1294 heterozygous).

#### Simulations of short reads from simulated hybrid genomes

Short reads from simulated hybrid genomes with different levels of heterozygosity (as described above) were obtained using DWGSIM with the parameters reported above. The average coverage was set at 50 x on the basis of the short-read simulations (see Figure 2e). Each simulation was performed in 3 replicates.

#### Performance of small variants calling

Given a set of relevant elements (i.e. the simulated variants) and a set of selected elements (i.e. the called variants) we classified each element (namely each variant) as true positive (*TP*), false positive (*FP*) or false negative (*FN*). We calculated precision as *P* = *TP/*(*TP* + *FP*) and recall as *R* = *TP/*(*TP* + *FN*). The performance of the small variants calling was quantified in terms of the *F*_1_ score which was calculated as the harmonic mean of (*P*) and (*R*) according to: *F*_1_ = 2 ·(*P ·R*)*/*(*P* + *R*). All the calculations were performed after filtering out polymorphic positions (SNMs and indels) determined by NUCmer as described below. In order to fairly compare competitive and standard mapping, the latter approach was run using a control sample for variant subtraction. This allowed for filtering out polymorphic positions (SNMs and indels) which could not be detected by NUCmer.

### Experimental data

Dataset 1 comprises a mutation accumulation line data from a mutator SK1/BY hybrid (*MATa/MATα; ARG4/arg4-nsp,bgl; his3*Δ*1/HIS3; leu2*Δ*0/leu2; met15*Δ*0/MET15; ura3*Δ*0/ura3; tsa1::KanMX/tsa1::KanMX)* generated as described by Serero *et al.* [49]. This dataset was analyzed using the SK1 and S288C assemblies included in MuLoYDH.

Dataset 2 consists of the UWOPS03-461.4/YPS128 hybrid (low sequence divergence; non-collinear genomes with chromosomal rearrangements) evolved under the RTG protocol (adapted from [29]). The hybrid (*MATa/MATα, ho::HygMX/ho::HygMX, ura3::KanMX/ura3::KanMX, leu2δ0/LEU2, met15δ0/MET15, LYS2/lys2::URA3)* was patched from the −80 °C glycerol stock on YPD solid media (1% yeast extract, 2% peptone, 2% dextrose, 2% agar) and incubated overnight at 30 °C. The following day a streak is made from YPD onto minimal solid media not supplemented with uracil and the plate is incubated at 30 °C for 48 hours. Different single colonies of the hybrid strain were taken and inoculated separatedly in 10 ml of pre-sporulation media YPEG 1% yeast extract, 2% peptone, 3% ethanol, 3% glycerol) for 15 hours at 30 °C with shaking at 220 rpm. Each pre-sporulation culture was washed twice with sterile water and resuspended in 2% potassium acetate (*OD*600 = 0.5) using a 250 ml flask that was incubated at 23 °C with shaking at 220 rpm. 1 ml was from the culture at the beginning of sporulation and another sample of 1 ml after 6 hours of incubation. The two samples were washed twice with 1 ml YPD and incubated in 1 ml YPD for 18 hours at 30 °C without shaking. The following day the YPD liquid cultures were vortexed and 20 μl of each culture were plated on minimal media containing 1 mg/ml 5-fluoroorotic acid (5-FOA) and spread with glass beads. The plate was then incubated at 30 °C for 48 hours.

Dataset 3 is composed by MALs from four distinct diploid backgrounds: N17/DBVPG6765, YPS128/DBVPG6765, N17/N17 and DBVPG6765/DBVPG6765. Each MAL consisted of eight independently propagated lines. *S. cerevisiae* DBVPG6765 homozygous diploids (*MATa/MATα, ho::HygMX/ho::HygMX, ura3::KanMX/ura3::KanMX, LYS2/lys2::URA3)* were derived from the Wine/European subpopulation. *S. paradoxus* N17 homozygous diploids (*MATa/MATα, ho::HygMX/ho::HygMX, ura3::KanMX/ura3::KanMX, LYS2/lys2::URA3)* were derived from the European subpopulation. YPS128/DBVPG6765 *S. cerevisiae* intraspecies hybrids (*MATa/MATα, ho::HygMX/ho::HygMX, ura3::KanMX/ura3::KanMX, LYS2/lys2::URA3)* were obtained by mating of North American (YPS128) and Wine/European (DBVPG6765) haploid strains. N17/DBVPG6765 interspecies hybrids (*MATa/MATα, ho::HygMX/ho::HygMX, ura3::KanMX/ura3::KanMX, LYS2/lys2::URA3)* were obtained by mating a *S. paradoxus* haploid strain from the European subpopulation (N17) and a *S. cerevisiae* haploid strain from the Wine/European subpopulation (DBVPG6765). Eight parallel mutation accumulation lines were propagated from each parental background on YPD solid medium (1% yeast extract, 2% peptone, 2% dextrose, 2% agar) and passed through a single cell bottleneck every ~48 hours (~20 generations) at 30 °C, for a total of 120 bottlenecks (~2400 generations). At each single cell bottleneck, a random colony was streaked to isolate the next single colony. To avoid any involuntary selection, at each streak, the closest colony to the center of the plate was picked, independently of its size. To determine the number of generations passed after 48 hours, three colonies for each parental background were independently resuspended in 100 μl of sterile water and serially diluted. 20 μl of each dilution were plated on solid YPD medium and grown for ~48 hours at 30 °C. The number of colonies was manually counted in the plate with suitable dilution and the number of generations (*G*) was estimated according to: *G* = *log*_2_(*n ·d*), where *n* is the number of cells counted on the plate and *d* is the corresponding dilution factor. The results are reported in Additional file 9: Table S8. After 120 single cell bottlenecks, cells were inoculated in 5 ml liquid YPD cultures and grown overnight at 30 °C in a shaking incubator. DNA was extracted using “Yeast Masterpure kit” (Epicentre, USA) following the manufacturer’s instructions.

### Sequencing

Illumina paired-end libraries (2 × 150 bp) were prepared according to manufacturer’s standard protocols and sequenced with an HiSeq 2500 instrument, at the NGS platform of Institut Curie. Coverage statistics are reported in Additional file 10: Table S9.

### Experimental validation of variants

7 SNMs supporting two LOHs and 9 SNVs variants from the SK1/BY hybrid were validated by Sanger sequencing. SNVs were randomly selected to avoid any bias. A pair of primers (upstream and downstream) was designed for each SNV using Unipro UGENE [77]. PCR products were sequenced by Eurofins Genomics^TM^. The presence and the genotype of the variants were checked by visual inspection of the electropherograms.

### Data analysis

#### Parental assemblies

All the parental assemblies reported in this work were downloaded from the “Population-level Yeast Reference Genomes” website (https://yjx1217.github.io/Yeast_PacBio_2016/welcome/). The assembly of *S. paradoxus* strain N17 was obtained correcting the genome sequence of its close relative CBS432, for which a complete assembly is available [11, 43]. The correction was performed using Pilon [78] with short-read data from Illumina sequencing of a diploid homozygous N17 sample. The command line is reported in Additional file 1.

#### MuLoYDH general description

The MuLoYDH pipeline requires as input: (1) a dataset of short-read sequencing experiments from yeast diploid hybrids and (2) the two parental genomes which were used to produce the hybrids in FASTA format as well as the corresponding annotations in the “general feature format” (GFF) (see Additional file 1: Figure S15). Reads from hybrid data are mapped against the assemblies of the two parental genomes separately (standard mappings) and against the union of the two aforementioned assemblies (namely a multi-FASTA obtained concatenating the two original assemblies) to produce the competitive mappings (Figure 1d-f). In the latter case, reads from parent 1 are expected to map to the assembly of parent 1 on the basis of the presence of single-nucleotide markers. Conversely, reads from parent 2 are expected to map to the assembly of parent 2. Standard mappings are used to determine the presence of CNVs. The latter are also exploited to discriminate LOHs due to recombination from those resulting by deletion of one parental allele. The SNMs between the parental assemblies are determined by the NUCmer algorithm and are exploited to map LOH segments. SNMs are genotyped from standard mappings. *De novo* small variants are determined from both competitive and standard mappings. Competitive mapping allows for direct variant phasing in heterozygous regions. Variant calling from competitive mapping is performed setting ploidy = 1 in heterozygous regions and ploidy = 2 in LOH blocks. Regions characterized by reads with low mapping quality (MAPQ < 5 in the competitive mapping) are assessed from standard mapping using arbitrarily the assembly from parent 1. All the scripts described in the following sections are embedded in MuLoYDH.

#### Quality check, mapping, mapping refinement and coverage calculation

Data quality is assessed by FastQC version 0.11.4. Competitive and standard mappings of Illumina reads are performed with BWA version 0.7.12-r1039 using the MEM algorithm [79]. Assemblies can be downloaded from the “Population-level Yeast Reference Genomes” website (https://yjx1217.github.io/Yeast_PacBio_2016/welcome/). Duplicates are removed by SAMtools 1.3.1 (using HTSlib 1.3.1). Depth of coverage is calculated with SAMtools (depth) and awk scripts. Additional file 11: Table S10 reports the coverage calculated for all the samples analysed in this study.

#### Determination of single-nucleotide marker positions

Single-nucleotide marker positions are determined through the NUCmer algorithm (MUMmer version 3) [with show-snps -ClrT] [76]. In order to obtain reliable SNM positions and take advantage of the “seed and extend” strategy of the algorithm, SNMs are calculated in both direct (assembly 1 vs assembly 2) and reverse (assembly 2 vs assembly 1) ways. The intersection of the two sets is retained for LOH detection and to calculate statistics.

#### Classification of single-nucleotide markers

SNMs are classified as lying in collinear or rearranged regions as determined by MUMmer and custom R scripts. The fraction of SNMs within collinear regions (*f_c_*) is calculated as *f_c_* = 1 − *f_r_*, where *f_r_* is the fraction of SNMs lying within rearranged regions, namely inter- and intra-chromosome inversions and translocations.

#### SNMs genotyping, small variants calling, annotation and filtering

SNMs calling and genotyping is performed using SAMtools (mpileup) [-u -min-MQ5 –skip-indels-E] and BCFtools (call) [-c -Oz] from standard mappings. SNMs are quality filtered removing those with quality < (*µ −σ*), where *µ* is the sample SNM mean quality value and *σ* is the corresponding standard deviation.

The strategy implemented in MuLoYDH for calling small variants relies on a stringent procedure to limit the number of false positives and keep the number of false negatives as low as possible. Thus, in order to balance performance (in terms of *F*_1_ score) and both the required computational resources and running time, two general-purpose small variants callers are implemented in MuLoYDH. SNVs and indels are called with: (i) SAMtools (mpileup) [-u -min-MQ5 -E] and BCFtools (call) [-c -Oz], and (ii) FreeBayes [80, 81]. Only variants called by both are retained. Both callers are exploited using competitive and standard mappings as described above. Regions characterized by reads with MAPQ < 5 in competitive mappings are determined by custom R scripts, bash scripts and BEDTools [82]). Parental and control hybrid variation is subtracted from hybrids data using custom bash scripts, VCFtools [83] and tabix [84]. The resulting variants are quality filtered masking those characterized by quality < (*µ−σ*), where *µ* is the sample SNMs mean quality value and *σ* is the corresponding standard deviation. Variants bearing SNM alleles are filtered out, while those lying within (sub)telomeric regions are masked. Small variants are annotated by means of SnpEff [85]. SnpEff database is built exploiting the annotations from the “Population-level Yeast Reference Genomes” website.

#### Copy-number variants calling and annotation

Copy-number variants are estimated by means of Control-FREEC with no matched normal samples, using standard mappings against both parental genomes [69]. Read-count data are normalized by GC-content and mappability. Mappability is calculated with GEM-mappability [86]. Results are annotated with *p*-values calculated with both Kolmogorov-Smirnov and Wilcoxon Rank-Sum tests.

#### Loss-of-heterozygosity detection and annotation

Loss-of-heterozygosity regions are determined and annotated using custom R scripts. Considering standard mappings of each hybrid against both parental assemblies, SNM positions characterized by non-matching genotype or alternate allele are filtered out, as well as multiallelic sites. Moreover, being our approach based on both parental assemblies, MuLoYDH calls LOHs without filtering out small events using an arbitrary threshold based on the number of supporting markers [29, 30]. This aspect is crucial since we aim at comparing LOH rates in *S. cerevisiae/S. cerevisiae* and *S. paradoxus/S. cerevisiae* crosses. SNMs involved in large deletions, as predicted by Control-FREEC, are masked. Finally, stretches of consecutive SNM positions are grouped in LOH regions. Genomic coordinates of each LOH event are determined using both the “first/last coordinates” and the ‘start/end coordinates”. First/last coordinates are determined using the coordinates of the first and the last SNMs of the event. Start/end coordinates are calculated using the average coordinate of the first (last) SNM and the last (first) SNM of the adjacent event. LOH regions are annotated as terminal/interstitial as well as with genomic features embedded and those potentially involved in breakpoints. Annotation is performed based on the genomic features downloaded from the “Population-level Yeast Reference Genomes” website. Interstitial LOHs are defined as homozygous segments that are flanked on both sides by heterozygous SNMs. Terminal LOHs are defined as homozygous regions extended to the end of the chromosmal arm.

#### Calculation of low-marker-density-regions

Regions characterized by less than one SNM in 300 bp are calculated using custom R scripts which are embedded in the MuLoYDH pipeline.

#### Platform

MuLoYDH was developed, tested and optimized using a Linux environment (OS openSUSE 13.2 x86 64), equipped with 64 Intel® Xeon® CPUs (E7-4820 @ 2.00 GHz).

#### Variants filtering in MALs and calculation of mutation rates

Small variants in DBVPG6044 and N17 homozygous backgrounds were quality-filtered on the basis of the values calculated from the SNMs of DBVPG6044/N17 hybrids as described above. All the small variants called by MuLoYDH were checked by visual inspection using IGV [87]. We also refined the lists of called CNVs by visual inspection in order to (I) avoid FPs due to small events which were not called in the control sample and (II) merge large events (e.g. aneuploidies) which were called as multiple shorter events. For each sample, we calculated the mutation rates dividing the number of variants detected and verified by visual inspection for number of generations calculated and for the length of the corresponding genome. Subtelomeric and telomeric regions were excluded from the calculation of small variants to avoid errors due to repeated regions.

#### Analysis of homozygous diploids

In order to analyze data from homozygous diploids, we set up a dedicated pipeline which is described in the following section. Reads from homozygous diploids were mapped against the proper assembly with BWA version 0.7.12-r1039 (MEM algorithm). Assemblies were downloaded from the “Population-level Yeast Reference Genomes” website. Duplicates were removed by means of SAMtools 1.3.1 (using HTSlib 1.3.1). Depth of coverage was calculated with SAMtools (depth) and awk scripts. Following duplicates removal, small variants were called with SAMtools and FreeBayes. The intersection of their outputs was retained and variants reported in control samples were removed. Small variants were annotated by means of SnpEff. SnpEff database was built exploiting the annotation data downloaded from the “Population-level Yeast Reference Genomes” website. The presence of copy-number variants was assessed by means of Control-FREEC with no matched normal samples. Read-count data were normalized by GC-content and mappability, while the latter was calculated by means of GEM-mappability. Results were annotated with *p*-values calculated with both Kolmogorov-Smirnov and Wilcoxon Rank-Sum tests.

## Supporting Information

### 1 Availability of data and materials

Sequencing data are available at http://ircan.unice.fr/Itattini/Datasets-MuLoYDH/. Samples IDs are described in Additional File 1: Table AF1-3. MuLoYDH is available for download at https://bitbucket.org/lt11/muloydh.git. A Docker image (including the mutator data) is available at http://ircan.unice.fr/ ltattini/OS.MuLoYDH.tar.

### Funding

This work was supported by: (A) Agence Nationale de la Recherche (ANR-11-LABX-0028-01, ANR-13-BSV6-0006-01, ANR-16-CE12-0019 and ANR-10-EQPX-03 Investissement d?Avenir to the IC NGS platform) and Fondation ARC pour la Recherche sur le Cancer (PJA20151203273), and (B) French government, through the UCAJEDI Investments in the Future project managed by the National Research Agency (ANR) with the reference number ANR-15-IDEX-01. Simone Mozzachiodi is funded by the convention CIFRE 2016/0582 between Meiogenix and ANRT. All authors discussed, critically revised and approved the final version of the manuscript.

### Author contributions

LT: designed and implemented computational methods, performed simulations, analyzed data and wrote the manuscript; NT: analyzed data, performed simulations; SM: tested computational methods, performed experimental validations; MDA: performed MAL experiments; SL: conducted experiments; AN: revised the manuscript; GL: coordinated and designed the study and wrote the manuscript.

## Acknowledgements

We thank Matteo De Chiara for discussions, Olivier Croce for technical support, Agnès Llored for experimental help, Gilles Fischer for critical reading of the manuscript, and the NGS platform of the Institut Curie for NGS sequencing.

## Additional Files

Additional file 1 — Supplementary Information

Supplementary figures and text.

Additional file 2 — Table S1. Reads mapping statistics

Statistics of unmapped and MAPQ = 0 reads.

Additional file 3 — Table S2. Variants detected in the A452R14 SK1/BY hybrid and results of the validations

List of the variants detected and validated by means of Sanger sequencing.

Additional file 4 — Table S3. LOHs data in YPS128/DBVPG6765 and N17/DBVPG6765 hybrids

Statistics of the LOHs detected, events detected for all MALs of both hybrids, and legend.

Additional file 5 — Table S4. SNVs data in MALs

Mutation rate statistics and data used for SNVs calculations for all the MALs.

Additional file 6 — Table S5. Indels data in MALs

Mutation rate statistics and data used for indels calculations for all the MALs.

Additional file 7 — Table S6. CNVs data in MALs

Mutation rate statistics and data used for CNVs calculations for all the MALs.

Additional file 8 — LOH segments in all MALs

Images of LOH segments detected in YPS128/DBVPG6765 and N17/DBVPG6765 MALs.

Additional file 9 — Table S8. Number of generations per single-cell bottleneck

The mean number of generations (and the standard deviation) for each MALs used to calculate the mutation rates.

Additional file 10 — Table S9. Mutation rates with mean values and standard deviations

The mean mutation rates and the corresponding standard deviations.

Additional file 11 — Table S10. Coverage statistics

Coverage statistics, after duplicates removal, for all the samples reported in this study.

