## Additional file 1 for "Accurate tracking of the mutational landscape of diploid hybrid genomes"

Supplementary Information

#### Contents

Genomic distribution of single-nucleotide markers
Benchmarking MuLoYDH against simulated datasets
Simulations of small variants in LOH regions
Sanger validation of phased SNVs and LOHs
Noise mitigation in LOH detection
Recombination in rearranged chromosomes from RTG data
Recombination in inter- and intra-species hybrids
Number of generations and mutation rates in MALs
Assessing absence of selection in MALs
MuLoYDH pipeline structure
Testing MuLoYDH
Requirements
Dependencies
Database configuration
Notes
Command lines
Data and software availability
References

### Genomic distribution of single-nucleotide markers

The genomic distribution of single-nucleotide markers (SNMs) is calculated automatically in MuLoYDH (Figures S1-S4). These plots provide insights on regions (e.g. telomers and subtelomers) characterized by few SNMs. Running MuLoYDH in "collinear" mode (whenever this condition holds) provides a more uniform distribution compared to the "rearranged" mode (Figure S5).

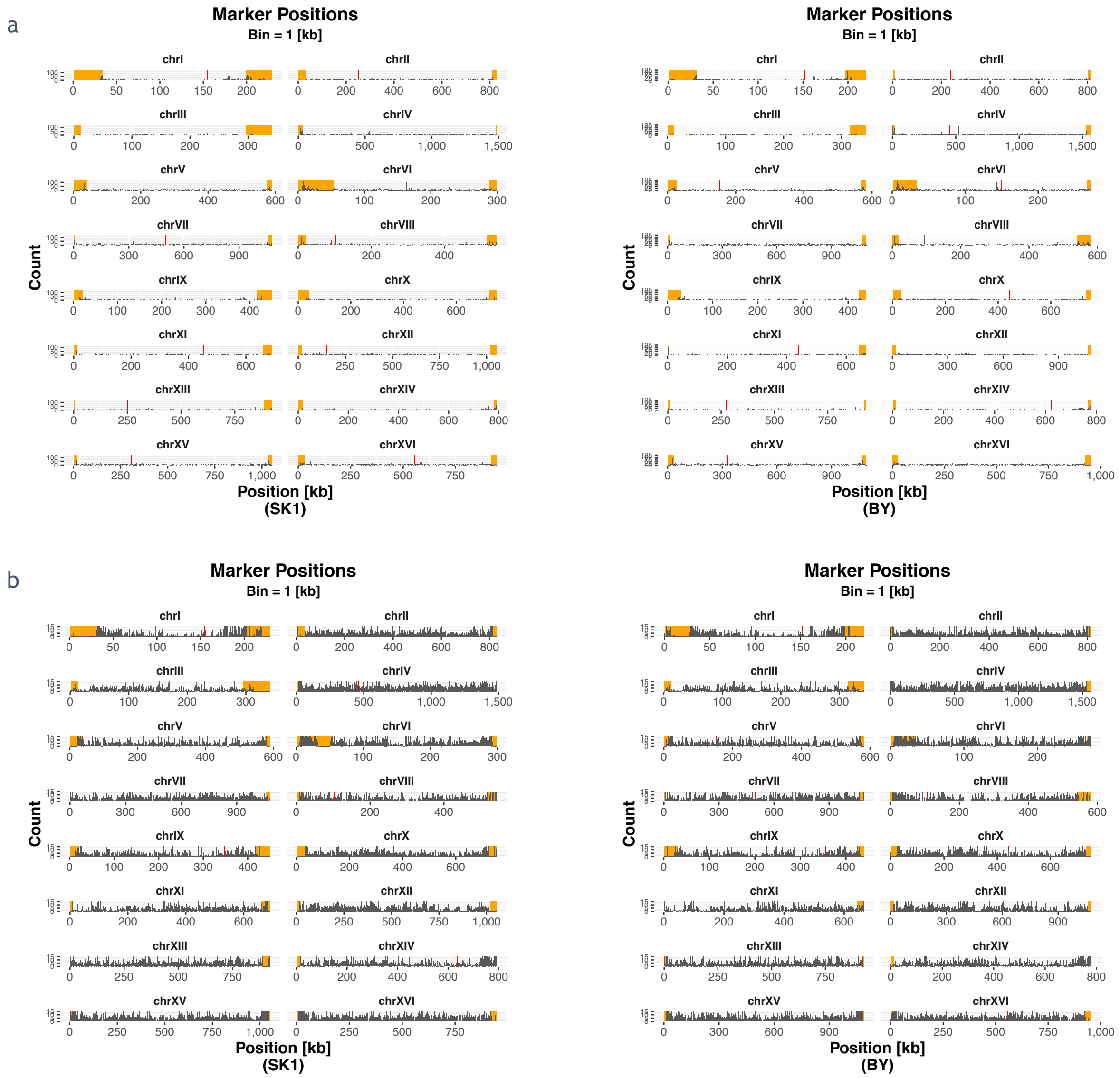


Figure S1 Genomic distribution of SNMs for SK1-BY (SK1-S288C, respectively) assemblies. (a) Histograms of SNM positions obtained from pairwise alignment of different pairs of assemblies. (b) Counts are cut at 15 to highlight regions showing no SNM. Yellow segments refer to (sub)telomeric regions. Centromeres are represented as red lines.

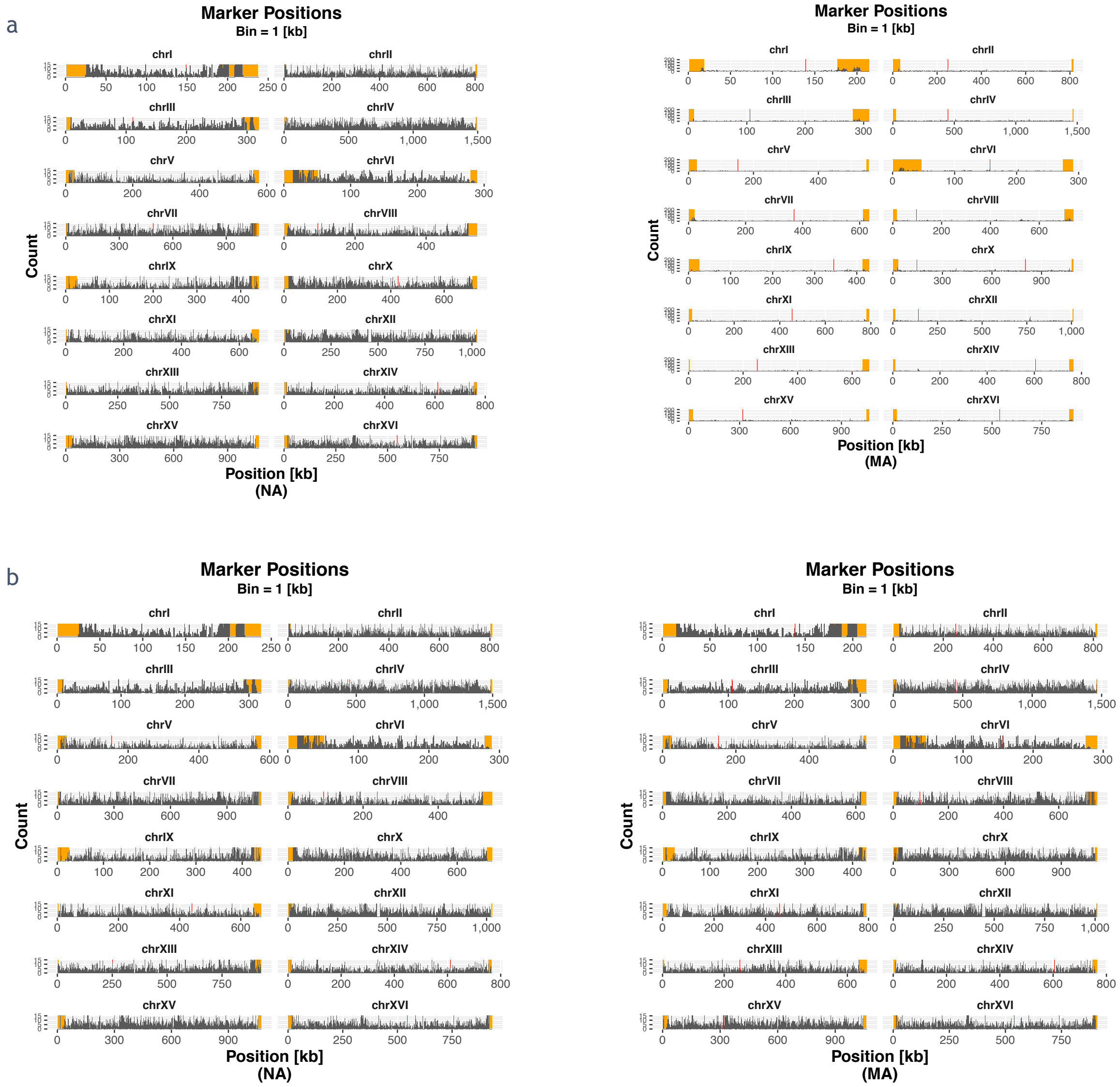


Figure S2 Genomic distribution of SNMs for YPS128-UWOPS03-461.4 (NA-MA, respectively). (a) Histograms of SNM positions obtained from pairwise alignment of different pairs of assemblies. (b) Counts are cut at 15 to highlight regions showing no SNM. Yellow segments refer to (sub)telomeric regions. Centromeres are represented as red lines.

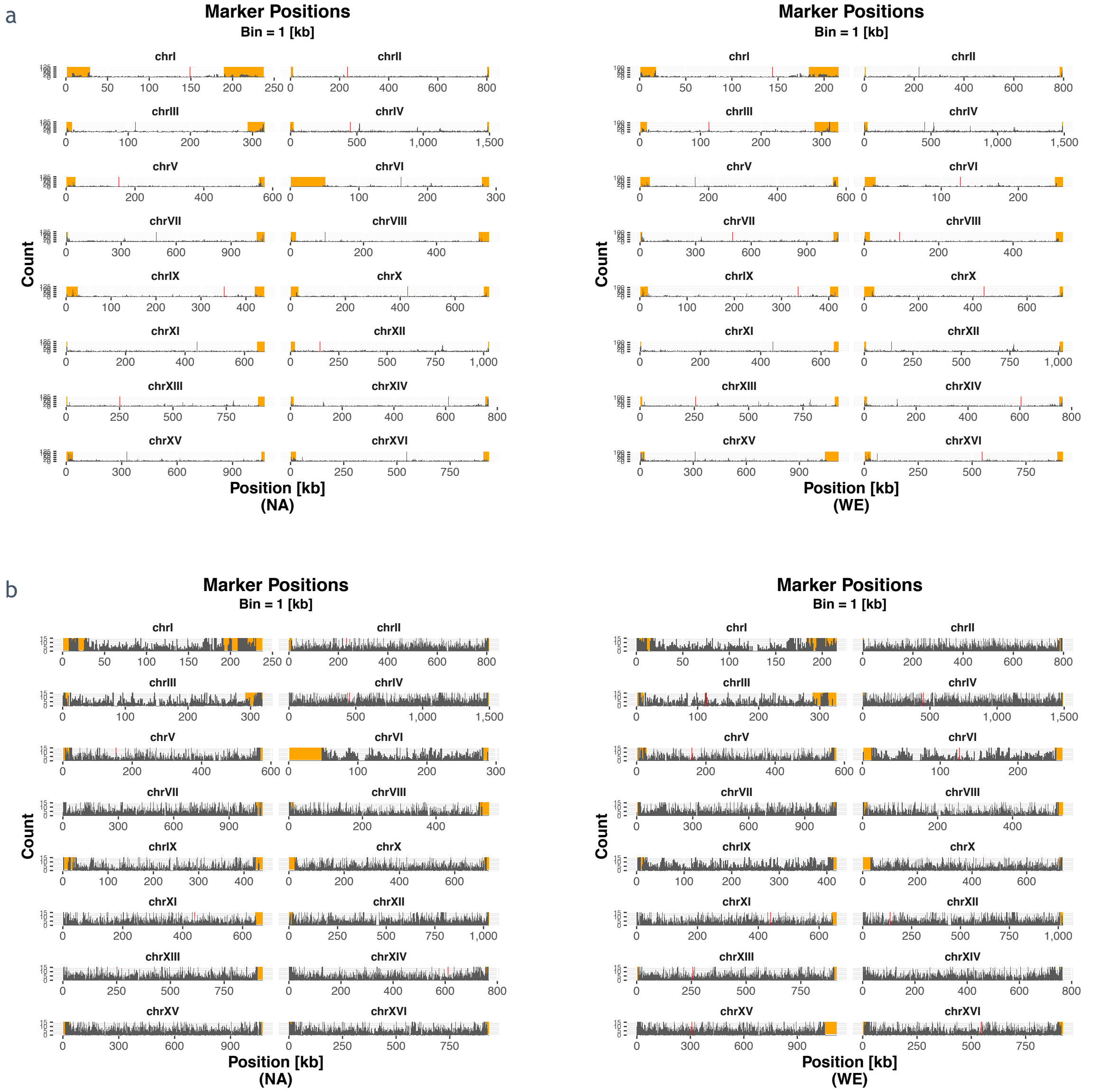


Figure S3 Genomic distribution of SNMs for YPS128-DBVPG6765 (NA-WE, respectively). (a) Histograms of SNM positions obtained from pairwise alignment of different pairs of assemblies. (b) Counts are cut at 15 to highlight regions showing no SNM. Yellow segments refer to (sub)telomeric regions. Centromeres are represented as red lines.

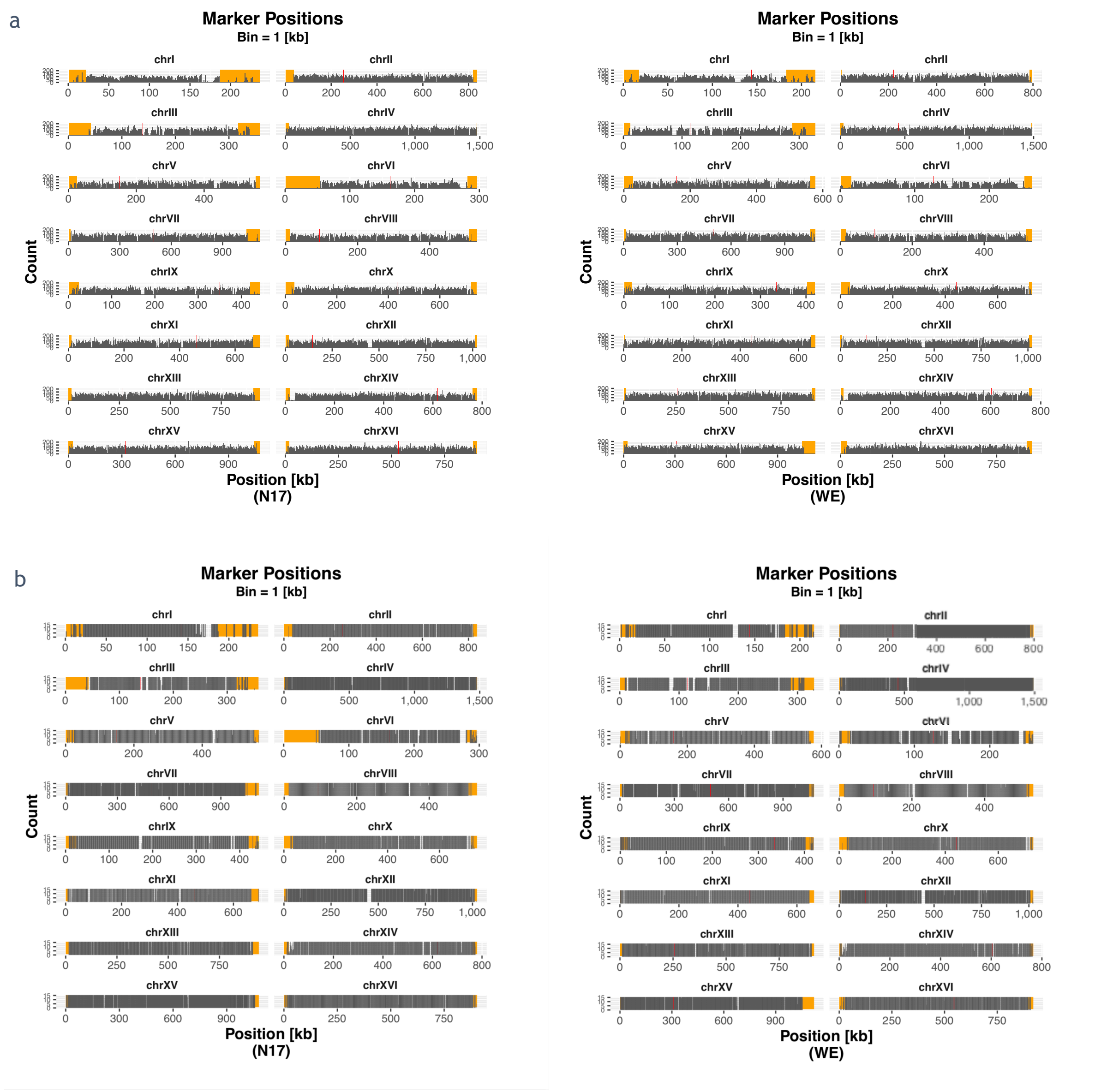


Figure S4 Genomic distribution of SNMs for N17-DBVPG6765 (N17-WE, respectively). (a) Histograms of SNM positions obtained from pairwise alignment of different pairs of assemblies. (b) Counts are cut at 15 to highlight regions showing no SNM. Yellow segments refer to (sub)telomeric regions.

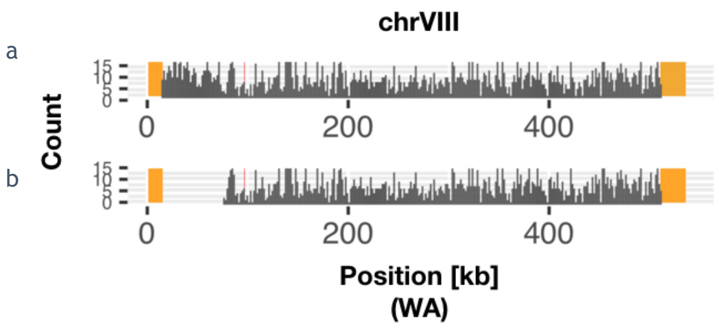


Figure S5 Distribution of SNMs. Histogram of SNM positions (chromosome VIII) obtained from the pairwise alignment of the DBVPG6044 (West African, WA) and YPS128 assemblies. Counts are cut at 15 in order to assess regions with no marker. Centromere is represented as a red line. Running MuLo in collinear mode (a) provide more uniform SNM distribution along chromosome VIII compared to the rearranged mode (b). Yellow segments refer to (sub)telomeric regions.

### Benchmarking MuLoYDH against simulated datasets

Figure S6 shows extended simulated data. The same trends discussed in the main text are observed.

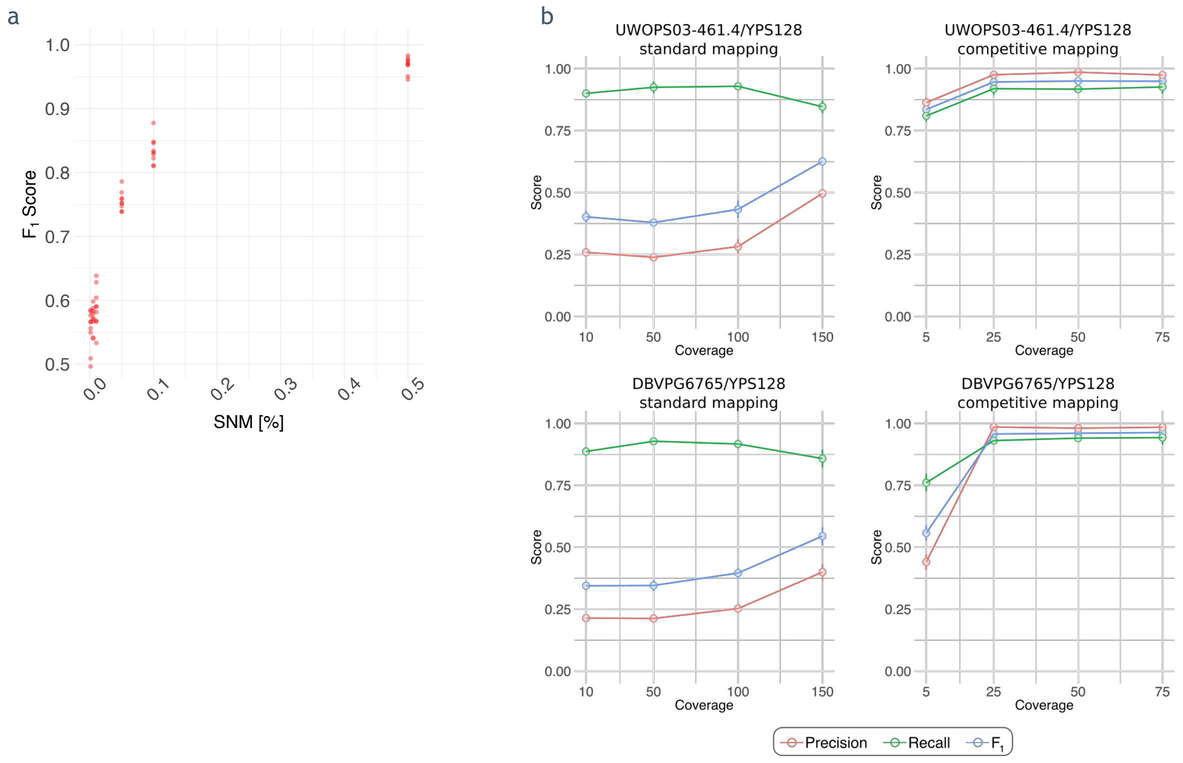


Figure S6 Small variant calling in simulated datasets. (a) The $F_1$ score is reported as a function of the percentage of SNMs. (b) Performance statistics for UWOPS03-461.4/YPS128 and DBVPG6765/YPS128 hybrids from standard and competitive mappings.

#### Simulations of small variants in LOH regions

MuLoYDH correctly called and genotyped 1840 variants in 10 replicates (Figure S7 and Table AF1-1) in the simulated LOH regions (691 homozygous and 1149 heterozygous variants), producing 62 FPs and 207 FNs ($F_1$=0.93 ± 0.01). 61 out of 62 FP variants (2 homozygous and 59 heterozygous) were in variant loci which had been incorrectly genotyped, thus resulting also in FNs. Nevertheless, the loci were correctly called. Among the 61 FPs, we observed 2 SNVs (one homozygous, one heterozygous). We also detected 58 homozygous mis-genotyped small deletions incorrectly called as heterozygous due to mis-mapping of reads which were not supporting the variant. Finally, only one heterozygous small insertion was incorrectly genotyped as homozygous thus resulting in one FP as well as one FN.

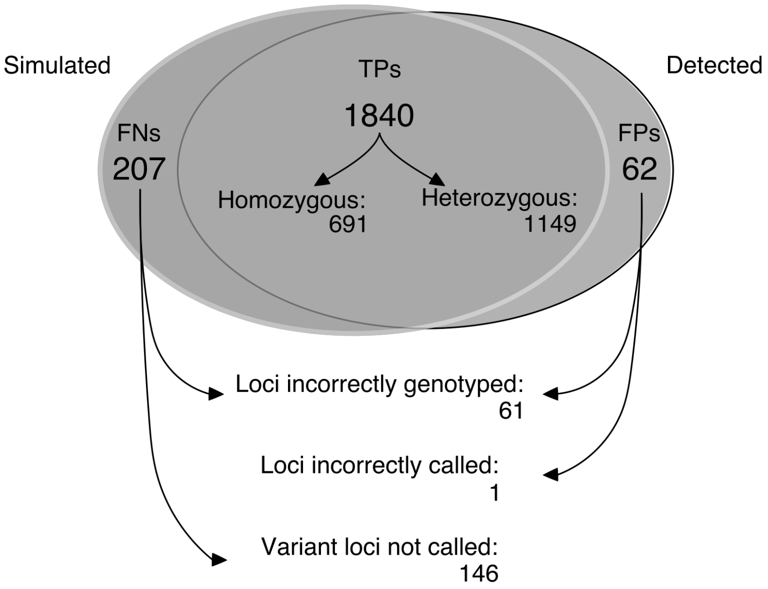


Figure S7 Detection and genotyping of variants in LOH regions.

| replicate | TP | FP | FN | precision | recall | F1 |
|---|---|---|---|---|---|---|
| 1 | 196 | 4 | 24 | 0.980 | 0.891 | 0.93 |
| 2 | 189 | 4 | 15 | 0.979 | 0.926 | 0.95 |
| 3 | 190 | 8 | 22 | 0.960 | 0.896 | 0.93 |
| 4 | 199 | 7 | 21 | 0.966 | 0.905 | 0.93 |
| 5 | 169 | 6 | 23 | 0.966 | 0.880 | 0.92 |
| 6 | 183 | 8 | 24 | 0.958 | 0.884 | 0.92 |
| 7 | 171 | 5 | 18 | 0.972 | 0.905 | 0.94 |
| 8 | 178 | 6 | 18 | 0.967 | 0.908 | 0.94 |
| 9 | 198 | 7 | 20 | 0.966 | 0.908 | 0.94 |
| 10 | 167 | 7 | 22 | 0.960 | 0.884 | 0.92 |
| mean | | | | | | 0.93 |
| SD | | | | | | 0.01 |

Table AF1-1. Performance of variant detection in LOH regions.

### Sanger validation of phased SNVs and LOHs

Electropherograms, annotated with variants and SNMs, are reported in Figures S8-S9. Validated phased variants in SK1 chromosome VIII and XII, and their closest SNMs are shown as well as SNMs in LOH regions (SK1 chrIV:1267782-1268523).

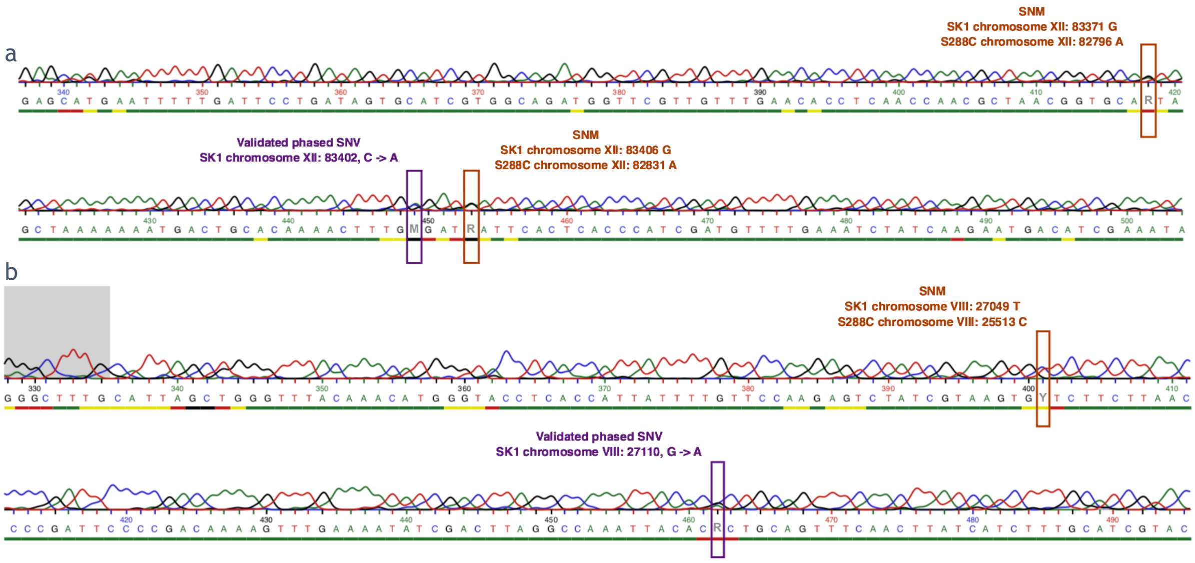


Figure S8 Sanger validation of phased SNVs. (a) Electropherograms showing validated phased variants in SK1 chromosome XII and their closest SNMs. (b) Electropherograms showing validated phased variants in SK1 chromosome VIII and their closest SNMs.

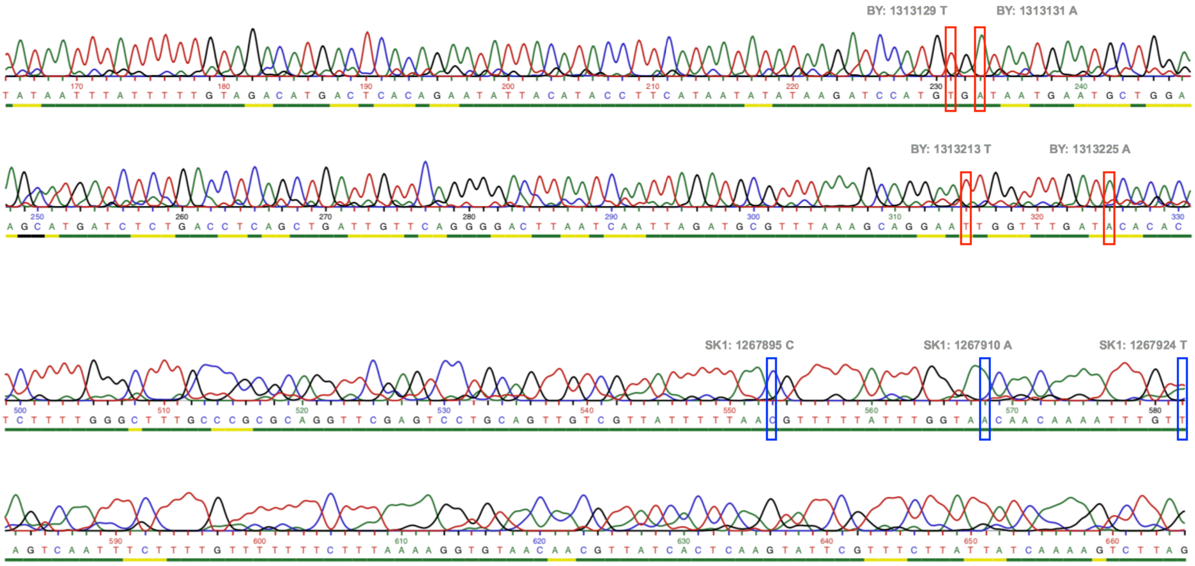


Figure S9 Sanger validation of LOHs. Homozygous SNM supporting two LOHs (BY chrIV: 1043775-1313337 and SK1 chrIV:1267782-1268523).

#### Noise mitigation in LOH detection

In contrast with the methods reported in the literature, the strategy implemented in MuLoYDH allows for noise mitigation and does not rely on any arbitrary threshold for determining SNM genotypes. Figure S10 shows allele frequencies and genotyping criteria for SK1/S288C hybrids that underwent return-to-growth [1]. Short reads were mapped against the SGD reference genome.

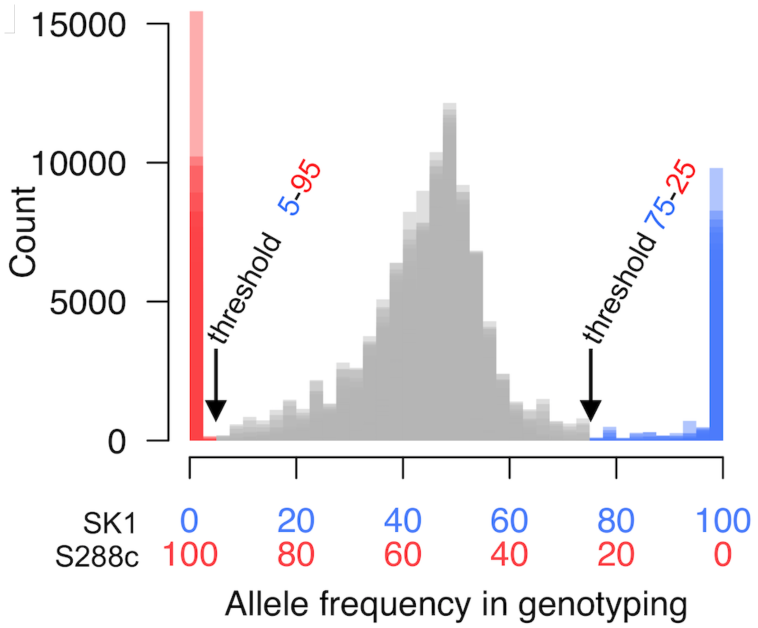


Figure S10 Determination of the genotyping thresholds based on the experimental data. Y-axis: count of sequencing reads. X-axis: percentage of allele specific reads. S288c alleles (red); SK1 alleles (blue); heterozygotes allleles (grey); Blue x-axis: fraction of SK1 frequency, red x-axis: S288c frequency. Note that the distribution of the S228c allele calls is rarely ambiguous because the alignment of the sequencing reads was performed on the SGD reference genome, which is quasi identical to the S288c genome used in the hybrid strain. In contrast, as expected, the call for the SK1 allele is more dispersed due to the lower efficiency of SK1 polymorphic read alignment on the SGD reference genome. Practically, the selected thresholds correspond to the minima experimentally observed in the distribution of the allelic frequencies in the parental and return-to-growth samples. Reproduced with permission from Laureau et al. [1].

#### Recombination in rearranged chromosomes from RTG data

The RC profiles obtained for rearranged chromosomes discussed in the main text can be calculated also for UWOPS03-461.4 chromosomes VII and VIII (Figure S11). As expected they confirm the results (discussed in the main text) calculated from mapping against YPS128.

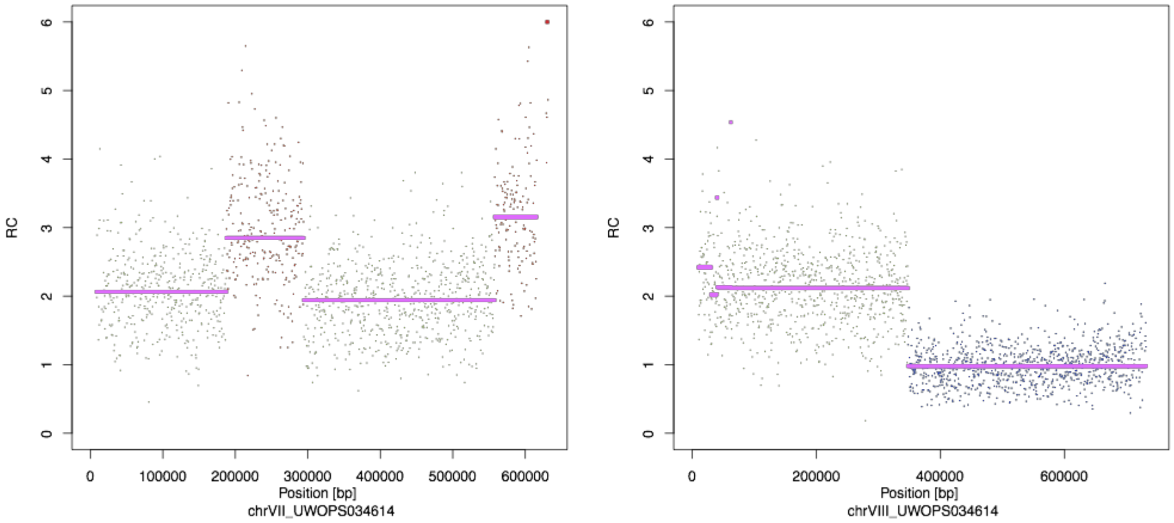


Figure S11 Read count data in rearranged chromosomes. Read count profiles obtained from mapping against UWOPS03-461.4 (corresponding to the events discussed in the main text).

### Recombination in inter- and intra-species hybrids

As reported in Figure S12, MuLoYDH automatically produces heatmaps of LOH events which help determining regions characterized by high recombination rate. For each chromosome, the mean density of LOHs is also plotted automatically (Figure S13). No correlation with chromosome length was observed. For each chromosome, plots of LOH segments are produced automatically as well (Figure S14) using the coordinates of the first and the last SNM supporting an event (upper panel) and start-end coordinates of the event (lower panel). These are calculated e.g. as the average coordinates of the first SNMs and the coordinate of the last SNM supporting the upstream segment. All of the LOH segments used to generate the heatmaps are reported in “Additional file 8 – LOH segments in all MALs”.

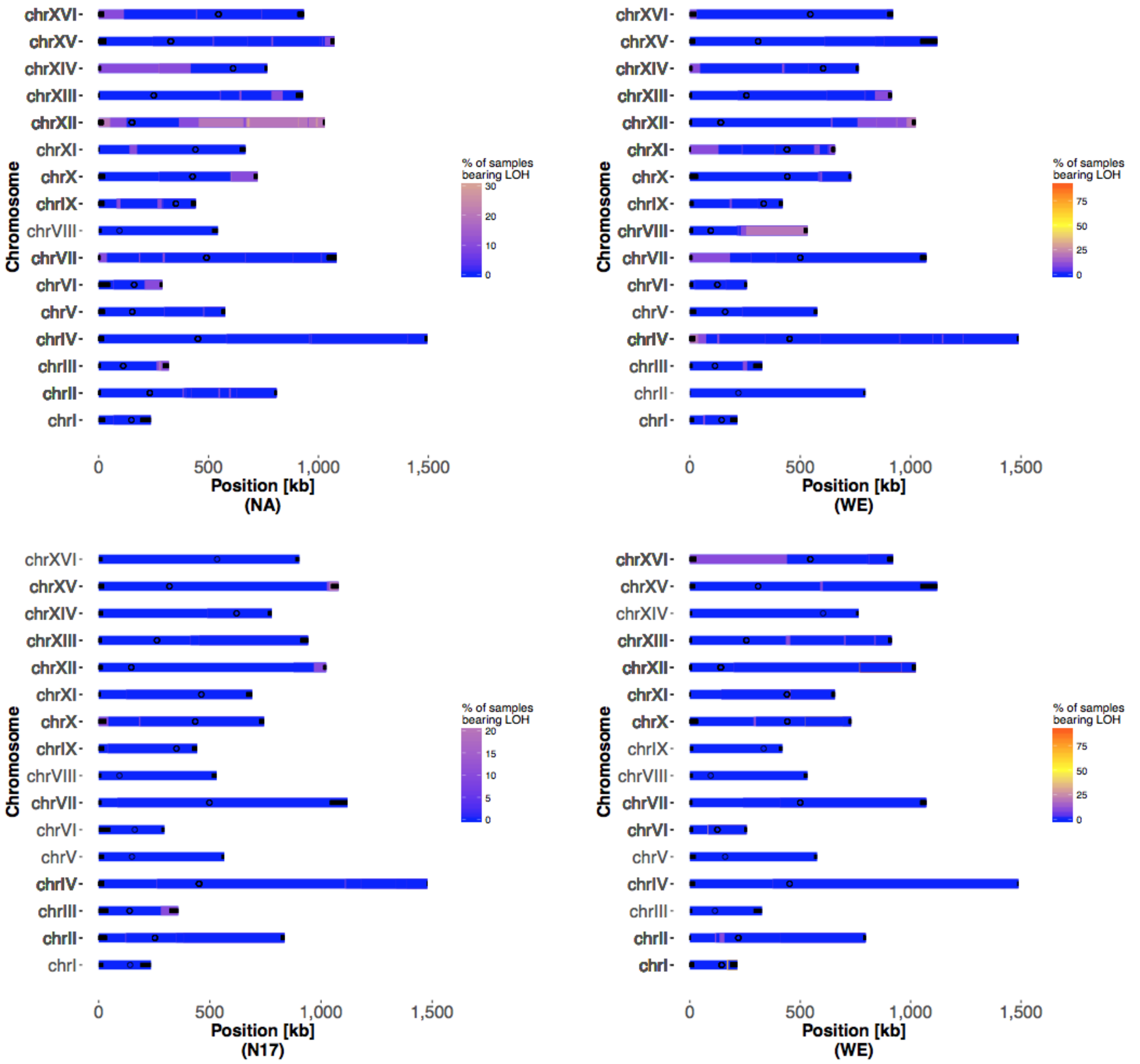


Figure S12 LOH heatmaps. Heatmaps of LOHs occurrence in YPS128/DBVPG6765 hybrids (upper panel) and N17/DBVPG6765 hybrids (lower panel).

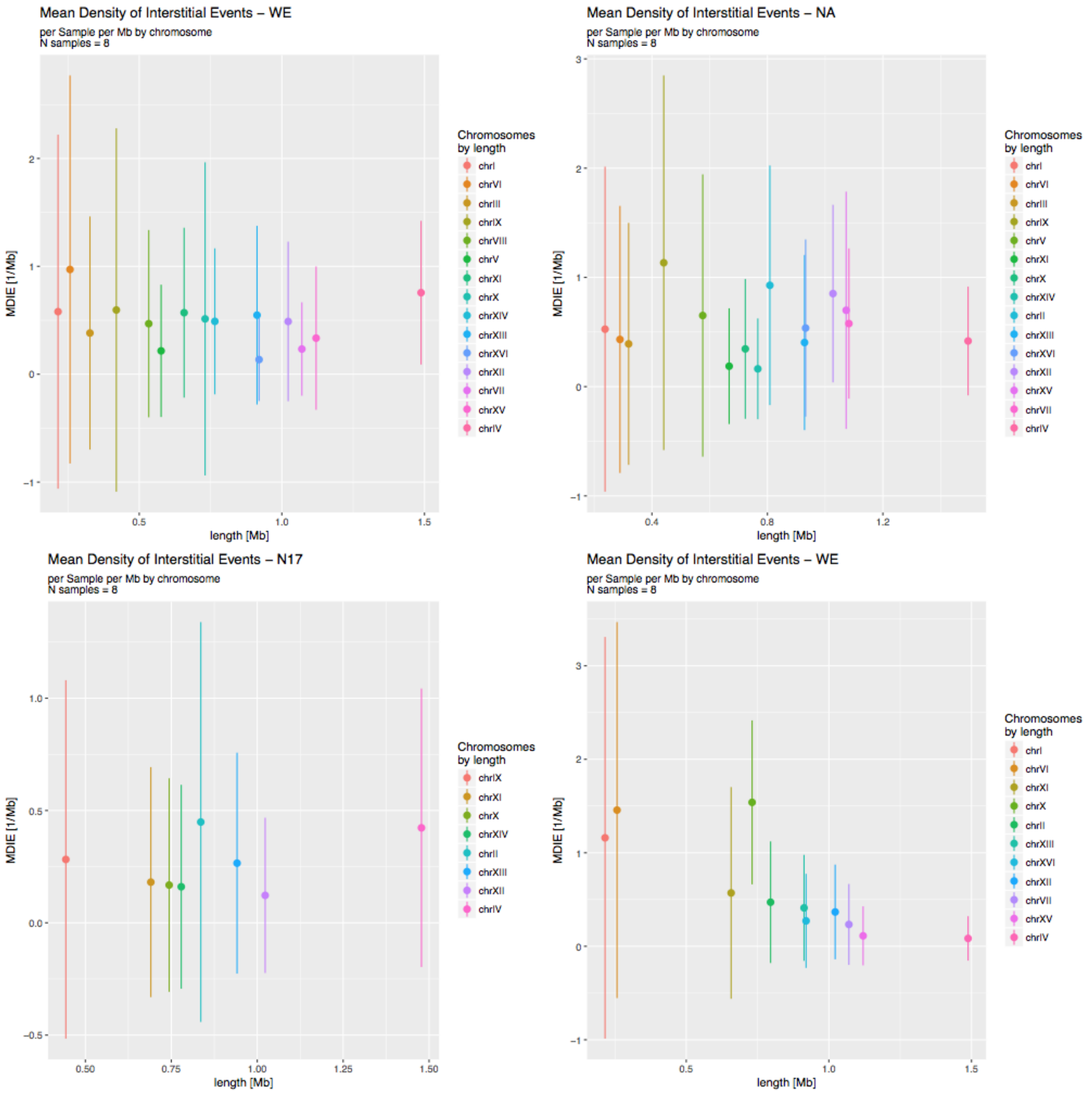


Figure S13 LOH density. Mean density of LOH events in YPS128/DBVPG6765 hybrids (upper panel) and N17/DBVPG6765 hybrids (lower panel). Only chromosomes with at least one event are reported.

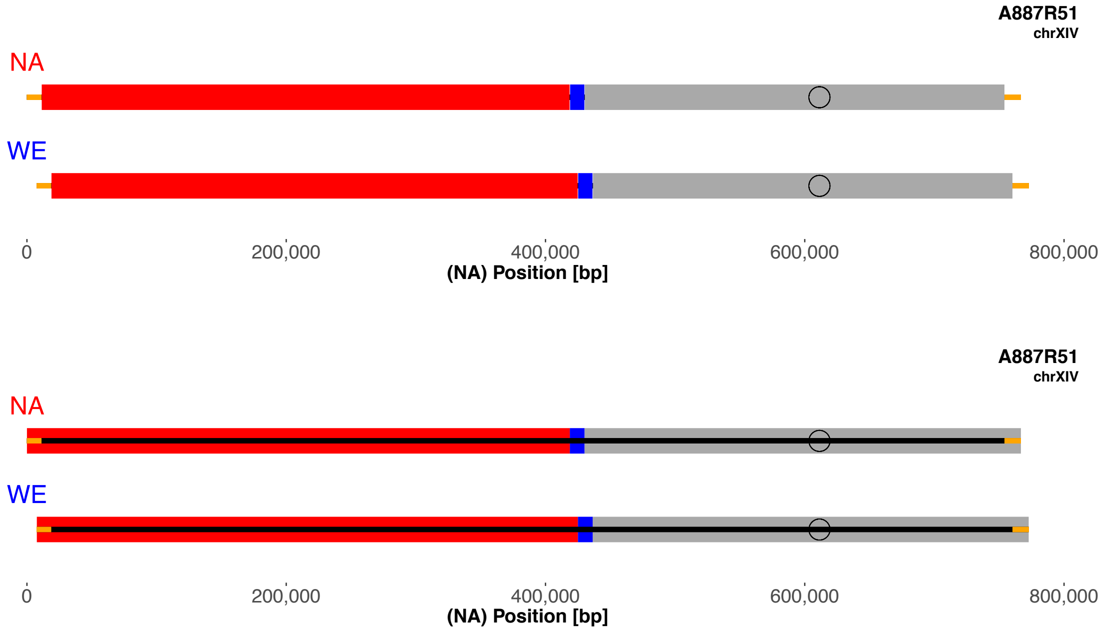


Figure S14. Raw segments plots for chromosome XIV of one YPS128/DBVPG6765 hybrid, (NA/WE, respectively). Yellow segments refer to (sub)telomeric regions.

#### Number of generations and mutation rates in MALs

Table AF1-2 shows the mutation rate per generation in different homozygous and heterozygous backgrounds for SNVs, indels, LOHs and CNVs. Standard deviations are also reported, along with the number of generations.

| Sample | Number of generations per bottleneck | Mutation rate per generation ± standard deviation [1/bp] (Number of variants in 8 lines ) | | | |
|---|---|---|---|---|---|
| | | SNVs | indels | CNVs | LOHs |
| YPS128 x DBVPG6765 | 18.7 ± 1.0 | 1.49E-10 ± 0.47E-10 (60) | 0.74E-11 ± 1.03E-11 (3) | 1.17E-11 ± 1.40E-11 (5) | 2.61E-10 ± 0.73E-10 (111) |
| N17 x DBVPG6765 | 18.5 ± 1.3 | 2.24E-10 ± 0.60E-10 (89) | 1.00E-11 ± 1.52E-11 (4) | 0.95E-11 ± 1.43E-11 (4) | 1.25E-10 ± 0.35E-10 (53) |
| DBVPG6765 x DBVPG6765 | 18.6 ± 0.9 | 2.82E-10 ± 1.02E-10 (113) | 1.50E-11 ± 2.33E-11 (6) | 0.95E-11 ± 1.43E-11 (4) | NA |
| N17 x N17 | 19.8 ± 0.6 | 7.27E-11 ± 4.65E-11 (31) | 4.69E-12 ± 8.69E-12 (2) | 2.19E-12 ± 6.19E-12 (1) | NA |

Table AF1-2. Number of generations, mutation rate and standard deviations of different diploid yeast backgrounds. Nuclear mutation rate per generation in different homozygous and heterozygous backgrounds for SNVs, indels, CNVs and LOHs. The rates were derived from 8 MALs per background propagated for ~2250 generations (120 bottlenecks).

#### Assessing absence of selection in MALs

We calculated the percentage of variants lying within coding regions and compared the results to the expected value (~75%) [2]. We found 20 out of 31 (64.5%) and 64 out of 89 (71.9%) coding variants for N17 diploids and N17/DBVPG6765 hybrids respectively. The percentage for DBVPG6765 diploids (77.9%, 88/113) and YPS128/DBVPG6765 hybrids (76.7%, 46/60) was closer to the expected value. Notably, the less concordant value was calculated from the MAL (N17 diploids) showing the lowest number of variants. We also calculated the percentage of non-synonymous variants occurring in coding regions obtaining 68.4% (13/19), 77.3% (68/88), 73.4% (47/64) and 76.1% (35/46) for N17, DBVPG6765, N17/DBVPG6765 and YPS128/DBVPG6765 respectively. These results suggest that no selection was occurring during the propagation of MALs.

#### MuLoYDH pipeline structure

MuLoYDH integrates several tools. The first part of the pipeline performs quality check of the experiments, the reciprocal alignment of the parental genomes and short-read mappings. Then, CNVs are determined and SNMs are genotyped. These informations are exploited to calculate LOHs. Finally, once the LOH profiles are known, small variants are assessed, filtered and annotated.

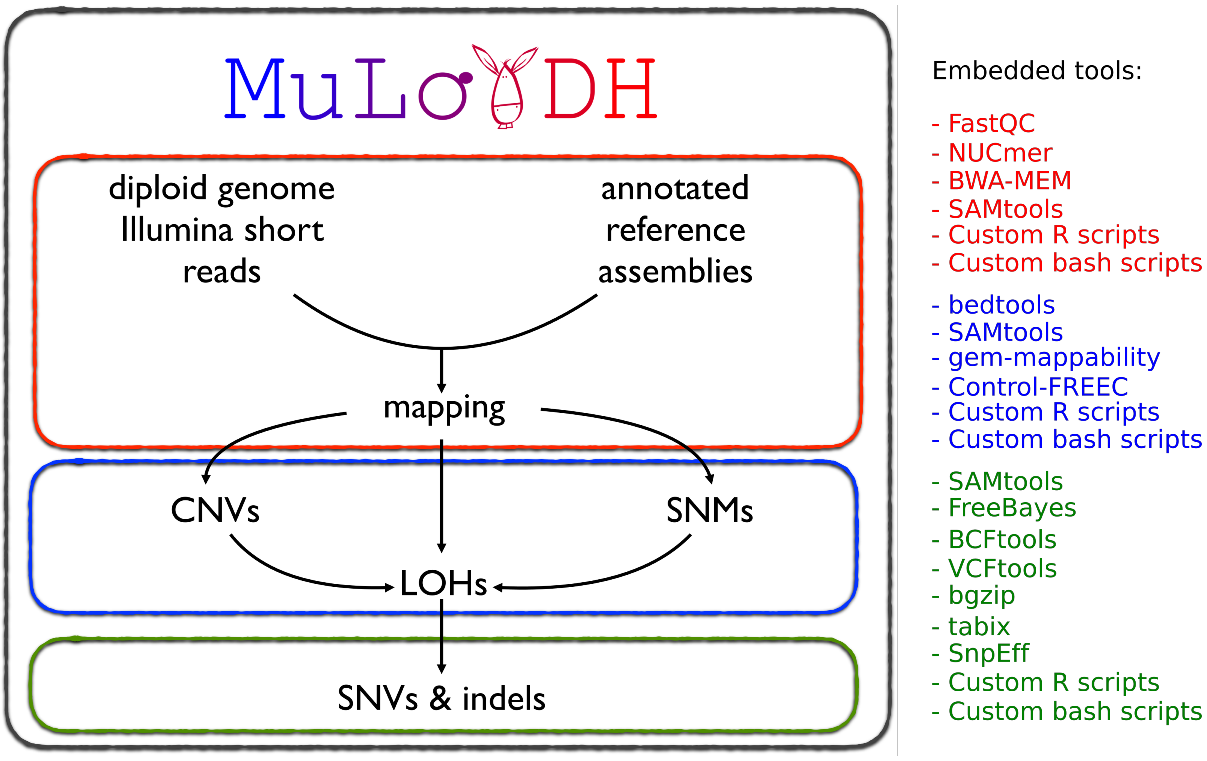


Figure S16 Structure of MuLoYDH pipeline. The MuLoYDH pipeline requires short-read data from evolved hybrids and the corresponding annotated parental assemblies. The tools embedded are reported on the right column.

#### Testing MuLoYDH

To test MuLoYDH you can use its Dockerized version, which already contains all of the softwares and the input data (including fastq data). This only requires the Docker software (https://www.docker.com/) to be installed in your machine. Once you have run the Docker daemon (e.g. in macOS just open the Docker app) and downloaded the tar image of MuLoYDH (http://ircan.unice.fr/~ltattini/OS.MuLoYDH.tar; please note that the tar file is ~12 GB), all you have to do is:

```
cd /path/to/tar/
# load the image
docker load < OS.MuLoYDH.tar
# run the container of the image
docker run -it opensuse/leap:test4MuLoYDH /bin/bash
# run the wrapper
cd /muloydh/Scr
bash MuLoYDH.V4.sh
```

Depending on your system, running the script will produce the results in few hours.

#### Requirements

The following linux (openSUSE, https://www.opensuse.org/) packages are required: vi, wget, unzip, gzip, tar, bzip2, git, R, time, curl, java-1_8_0-openjdk-devel, gcc, gcc-c++, perl, make, python, autoconf, automake, zlib-devel, libbz2-devel, xz-devel, libcurl-devel, libopenssl-devel, ncurses-devel, tcsh, gcc-fortran, libxml2, libxml2-devel.
These can be installed using the following bash lines:

```
# install openSuse basic packages (interactive)
zypper install vi wget unzip gzip tar bzip2 git R time
zypper install curl java-1_8_0-openjdk-devel gcc gcc-c++ perl make python
# more dependencies for samtools (interactive)
zypper install autoconf automake zlib-devel libbz2-devel xz-devel
zypper install libcurl-devel libopenssl-devel ncurses-devel
# more dependencies for mummer (interactive)
zypper install tcsh
# fortran for ape library (interactive)
zypper install gcc-fortran
# packages for rtracklayer (used by control-freec) (interactive)
zypper install libxml2 libxml2-devel
```

#### Dependencies

Several tools are necessary to run MuLoYDH. They must be installed and all the folders containing the corresponding executables must be exported to your PATH. To download the tools in "/download/TOOL_NAME" and install them in a openSUSE linux machine (/usr/local/bin) you can login as root and execute the following bash lines:

```
# set executables' folder
InstallDir="/usr/local/bin"

# download folder for all tools
cd /
mkdir /download

# MuLoYDH dependency 1: download & install samtools
cd /
mkdir /download/samtools
cd /download/samtools
wget https://github.com/samtools/samtools/releases/download/1.9/samtools-1.9.tar.bz2
tar -xjf samtools-1.9.tar.bz2
cd samtools-1.9
./configure
make
make install

# MuLoYDH dependency 2: download & install fastqc
cd /
mkdir /download/fastqc
cd /download/fastqc
wget https://www.bioinformatics.babraham.ac.uk/projects/fastqc/fastqc_v0.11.8.zip
unzip fastqc_v0.11.8.zip
cd FastQC
chmod a+x fastqc
for i in $(find $PWD -maxdepth 1 -type f -perm /a+x); do ln -s $i $InstallDir/$(basename $i);
done
# MuLoYDH dependency 3: download & install bwa
cd /
mkdir /download/bwa
cd /download/bwa
git clone https://github.com/lh3/bwa.git
cd bwa
make
for i in $(find $PWD -maxdepth 1 -type f -perm /a+x); do ln -s $i $InstallDir/$(basename $i);
done

# MuLoYDH dependency 4: download & install bedtools
cd /
mkdir /download/bedtools
cd /download/bedtools
git clone https://github.com/arq5x/bedtools2.git
cd bedtools2
make
cd bin
for i in $(find $PWD -maxdepth 1 -type f -perm /a+x); do ln -s $i $InstallDir/$(basename $i);
done

# MuLoYDH dependency 5: download & install mummer
cd /
mkdir /download/mummer
cd /download/mummer
wget https://sourceforge.net/projects/mummer/files/mummer/3.23/MUMmer3.23.tar.gz
tar -zxvf MUMmer3.23.tar.gz
cd MUMmer3.23
make install
for i in $(find $PWD -maxdepth 1 -type f -perm /a+x); do ln -s $i $InstallDir/$(basename $i);
done

# MuLoYDH dependency 6: download & install bcftools
cd /
mkdir /download/bcftools
cd /download/bcftools
wget https://github.com/samtools/bcftools/releases/download/1.9/bcftools-1.9.tar.bz2
tar -xjf bcftools-1.9.tar.bz2
cd bcftools-1.9
./configure
```

```
make
make install

# MuLoYDH dependency 7: download & install R packages (interactive)
R
source("https://bioconductor.org/biocLite.R")
biocLite("IRanges")
biocLite("rtracklayer")
install.packages("ape")
install.packages("ggplot2")
install.packages("scales")
install.packages("vcfR")

# MuLoYDH dependency 8: download & install gem
cd /
mkdir /download/gem
cd /download/gem
wget http://barnaserver.com/gemtools/releases/GEMTools-static-i3-1.7.1.tar.gz
tar -zvxf GEMTools-static-i3-1.7.1.tar.gz
cd gemtools-1.7.1-i3
for i in $(find $PWD -maxdepth 1 -type f -perm /a+x); do ln -s $i $InstallDir/$(basename $i);
done

# MuLoYDH dependency 9: download & install control-freec
cd /
mkdir /download/control-freec
cd /download/control-freec
git clone https://github.com/BoevaLab/FREEC.git
cd FREEC/src
make
for i in $(find $PWD -maxdepth 1 -type f -perm /a+x); do ln -s $i $InstallDir/$(basename $i);
done

# MuLoYDH dependency 10: download & install htslib (with tabix)
cd /
mkdir /download/htslib
cd /download/htslib
wget https://github.com/samtools/htslib/releases/download/1.9/htslib-1.9.tar.bz2
tar -xjf htslib-1.9.tar.bz2
cd htslib-1.9
./configure
make
make install

# MuLoYDH dependency 11: download & install vcftools
cd /
mkdir /download/vcftools
cd /download/vcftools
git clone https://github.com/vcftools/vcftools.git
cd vcftools
./configure
make
make install

# MuLoYDH dependency 12: download & install freebayes
cd /
mkdir /download/freebayes
cd /download/freebayes
git clone --recursive git://github.com/ekg/freebayes.git
cd freebayes
make
make install
```

Finally, SnpEff is required. It is provided in the libraries embedded in MuLoYDH since it requires manual configuration of the database. Instructions to configure it for novel organisms are reported in BuildSnpEff.txt (Scr folder).

#### Notes

Input fastq file name must be set according to the format: “exp_id.R1.fastq.gz”, “exp_id.R2.fastq.gz”. The prefix “exp_id” may contain only alphanumeric characters. Input fasta files must be named “strain_id.genome.fa” and must be properly sorted, e.g. chrI, chrII, chrIII, chrIV, chrV, chrVI. The prefix “strain_id” may contain only alphanumeric characters.

#### Command lines

| Tool | Command line | Comments |
|---|---|---|
| dwgsim | dwgsim -R 0.1 -e 0.01 -E 0.01 -d Isz -s 125 -1 150 -2 150 -r 0.00001 -q B -Q 2 -C Cov InputFasta Prefix | Isz is the insert size, Cov the coverage value, InputFasta the concatenated parental fasta, Prefix the output prefix string |
| pilon | pilon --genome InputFasta --frags InputBam --changes --fix "bases" | InputBam is the Illumina experiment used to correct InputFasta |

#### Data and software availability

All the sequencing datasets will be uploaded to SRA (https://www.ncbi.nlm.nih.gov/sra). They are currently available at http://ircan.unice.fr/~ltattini/Datasets-MuLoYDH/. Samples IDs are described in Table AF1-3. MuLoYDH is available for download at https://bitbucket.org/lt11/muloydh.git. A Docker image (including the mutator data) is available at http://ircan.unice.fr/~ltattini/OS.MuLoYDH.tar.

#### Samples

| ID | Dataset | Background | Notes | Protocol |
|---|---|---|---|---|
| **A784R51** | 1 | SK1 x S288C | control | MAL |
| **A452R14** | 1 | SK1 x S288C | evolved | MAL |
| **A505R6** | 2 | UWOPS03-461.4 X YPS128 | control | RTG |
| **A347R12** | 2 | UWOPS03-461.4 X YPS128 | evolved | RTG |
| **A887R37** | 3 | DBVPG6765 x DBVPG6765 | control | MAL |
| **A887R38** | 3 | DBVPG6765 x DBVPG6765 | evolved | MAL |
| **A887R39** | 3 | DBVPG6765 x DBVPG6765 | evolved | MAL |
| **A887R40** | 3 | DBVPG6765 x DBVPG6765 | evolved | MAL |
| **A887R41** | 3 | DBVPG6765 x DBVPG6765 | evolved | MAL |
| **A887R42** | 3 | DBVPG6765 x DBVPG6765 | evolved | MAL |
| **A887R43** | 3 | DBVPG6765 x DBVPG6765 | evolved | MAL |
| **A887R44** | 3 | DBVPG6765 x DBVPG6765 | evolved | MAL |
| **A887R45** | 3 | DBVPG6765 x DBVPG6765 | evolved | MAL |
| **A887R19** | 3 | DBVPG6765 x DBVPG6765 | evolved | MAL |
| **A887R1** | 3 | N17 x N17 | control | MAL |
| **A887R2** | 3 | N17 x N17 | evolved | MAL |
| **A887R3** | 3 | N17 x N17 | evolved | MAL |
| **A887R4** | 3 | N17 x N17 | evolved | MAL |
| **A887R5** | 3 | N17 x N17 | evolved | MAL |
| **A887R6** | 3 | N17 x N17 | evolved | MAL |
| **A887R7** | 3 | N17 x N17 | evolved | MAL |
| **A887R8** | 3 | N17 x N17 | evolved | MAL |
| **A887R9** | 3 | N17 x N17 | evolved | MAL |
| **A887R46** | 3 | YPS128/DBVPG6765 | control | MAL |
| **A887R47** | 3 | YPS128/DBVPG6765 | evolved | MAL |
| **A887R48** | 3 | YPS128/DBVPG6765 | evolved | MAL |
| **A887R49** | 3 | YPS128/DBVPG6765 | evolved | MAL |
| **A887R50** | 3 | YPS128/DBVPG6765 | evolved | MAL |
| **A887R51** | 3 | YPS128/DBVPG6765 | evolved | MAL |
| **A887R52** | 3 | YPS128/DBVPG6765 | evolved | MAL |
| **A887R53** | 3 | YPS128/DBVPG6765 | evolved | MAL |
| **A887R54** | 3 | YPS128/DBVPG6765 | evolved | MAL |
| **A887R19** | 3 | N17/DBVPG6765 | control | MAL |
| **A887R20** | 3 | N17/DBVPG6765 | evolved | MAL |
| **A887R21** | 3 | N17/DBVPG6765 | evolved | MAL |
| **A887R22** | 3 | N17/DBVPG6765 | evolved | MAL |
| **A887R23** | 3 | N17/DBVPG6765 | evolved | MAL |
| **A887R24** | 3 | N17/DBVPG6765 | evolved | MAL |
| **A887R25** | 3 | N17/DBVPG6765 | evolved | MAL |
| **A887R26** | 3 | N17/DBVPG6765 | evolved | MAL |
| **A887R27** | 3 | N17/DBVPG6765 | evolved | MAL |

Table AF1-3. List of samples included in this study.
