## Additional file 8 for "Accurate tracking of the mutational landscape of diploid hybrid genomes"

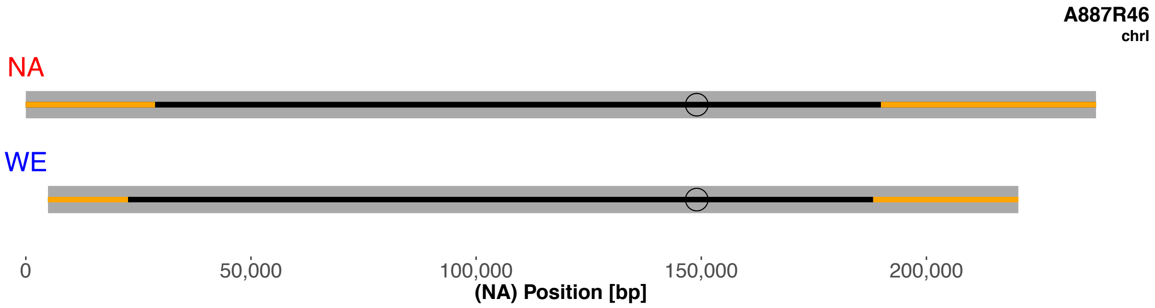
A887R46
chrI
NA
WE
0
50,000
100,000
150,000
200,000
(NA) Position [bp]

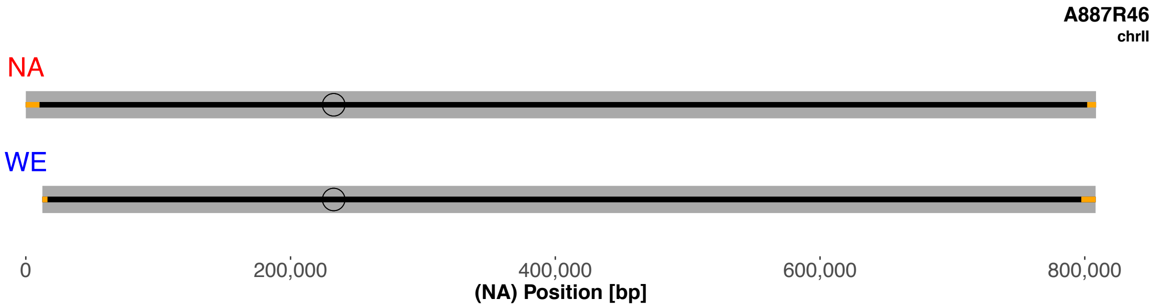
A887R46
chrII
NA
WE
0
200,000
400,000
600,000
800,000
(NA) Position [bp]

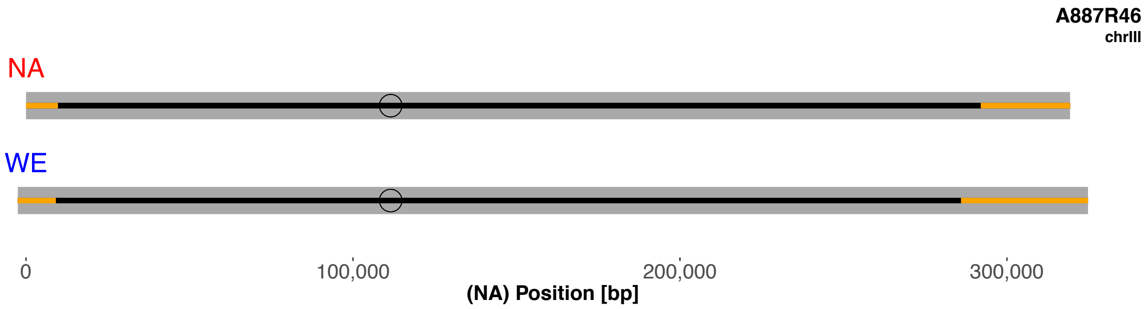
A887R46
chrIII
NA
WE
0
100,000
200,000
300,000
(NA) Position [bp]

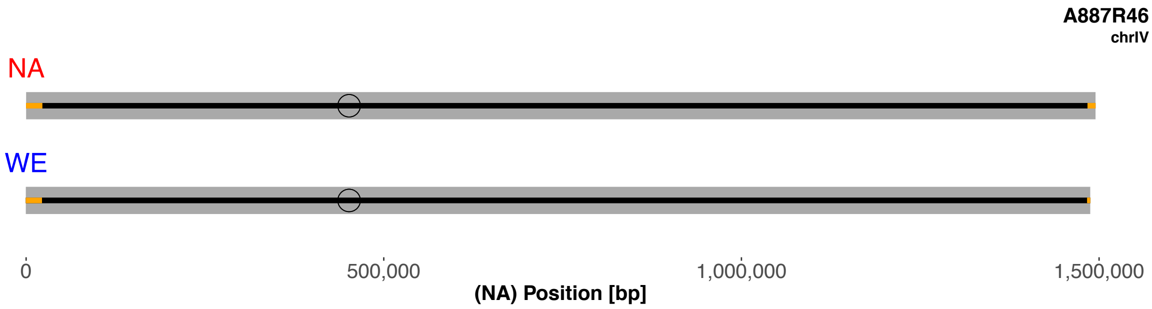
A887R46
chrIV
NA
WE
0
500,000
1,000,000
1,500,000
(NA) Position [bp]

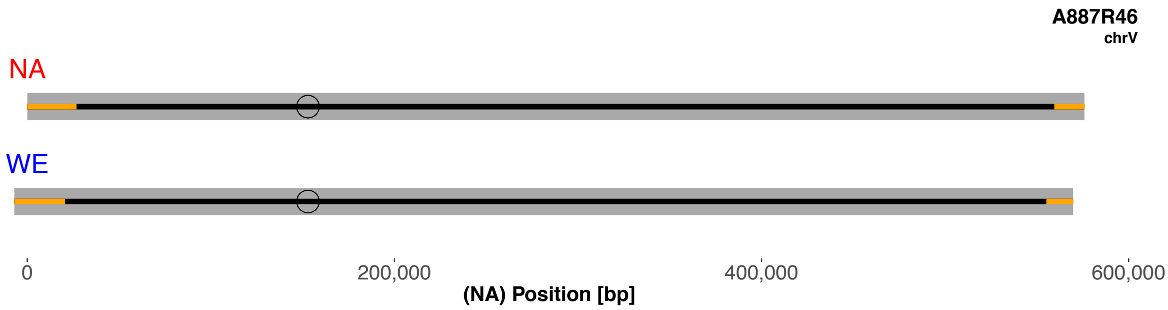
A887R46
chrV
NA
WE
0
200,000
400,000
600,000
(NA) Position [bp]

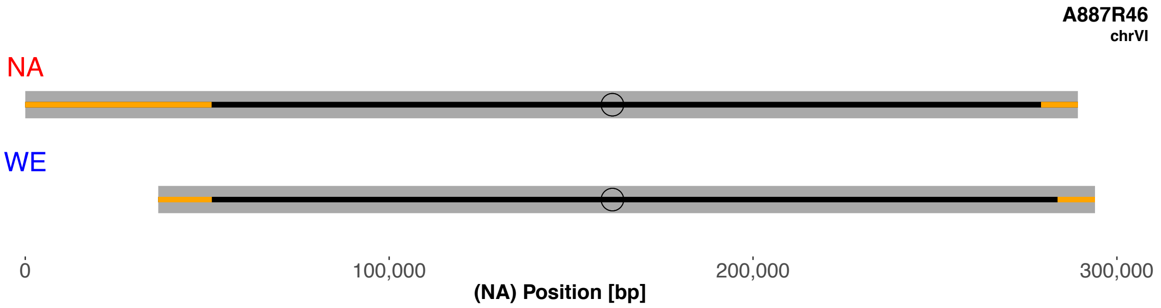
A887R46
chrVI
NA
WE
0
100,000
200,000
300,000
(NA) Position [bp]

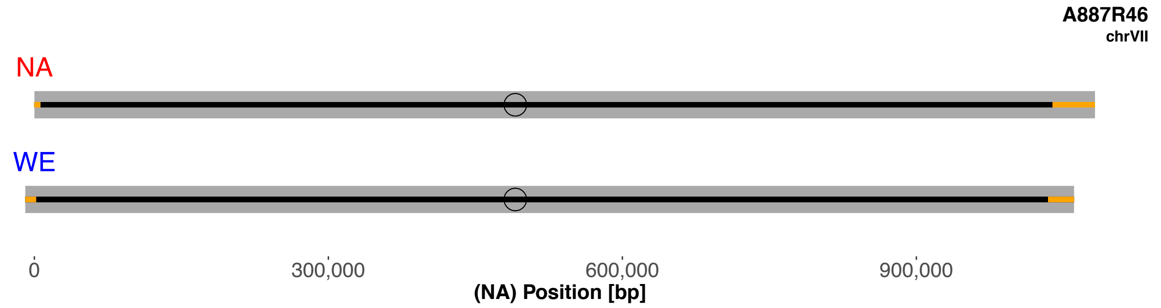
A887R46
chrVII
NA
WE
0
300,000
600,000
900,000
(NA) Position [bp]

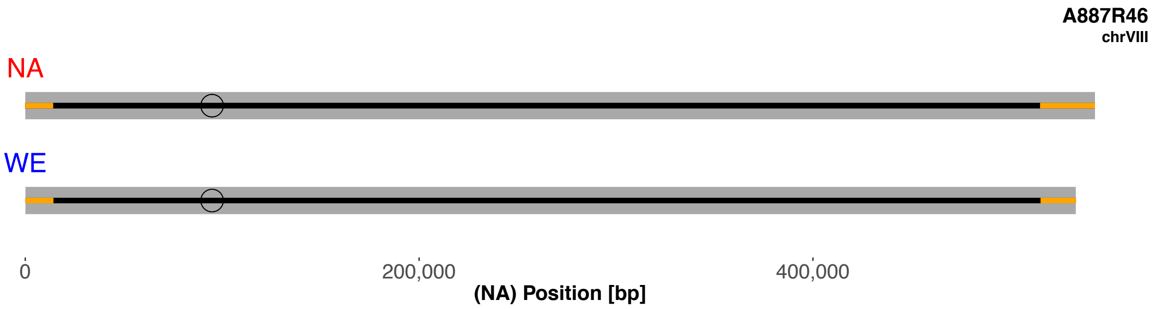
A887R46
chrVIII
NA
WE
0
200,000
400,000
(NA) Position [bp]

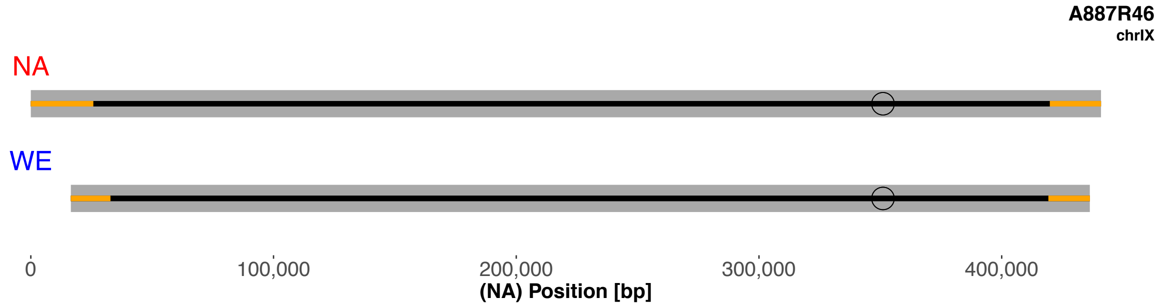
A887R46
chrIX
NA
WE
0
100,000
200,000
300,000
400,000
(NA) Position [bp]

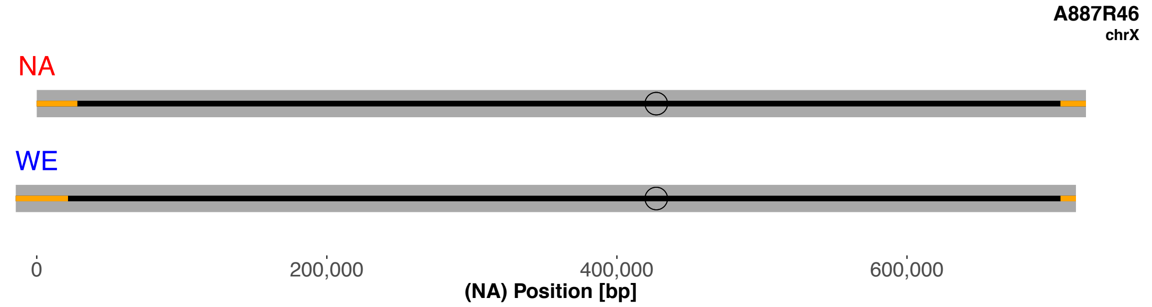
A887R46
chrX
NA
WE
0
200,000
400,000
600,000
(NA) Position [bp]

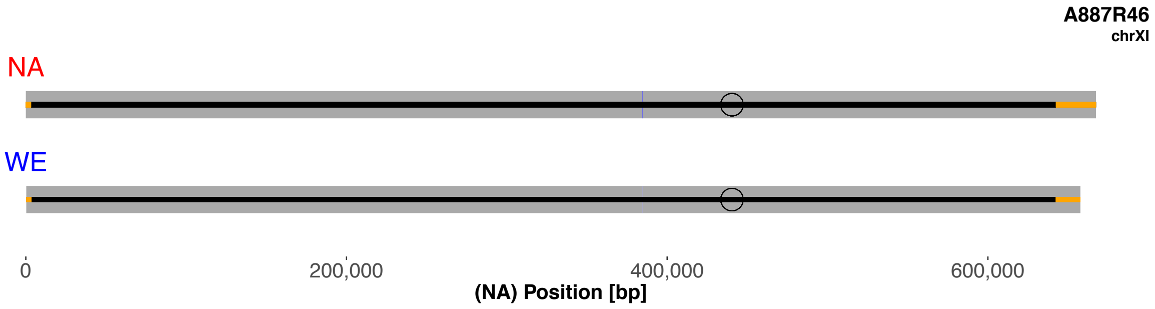
A887R46
chrXI
NA
WE
0
200,000
400,000
600,000
(NA) Position [bp]

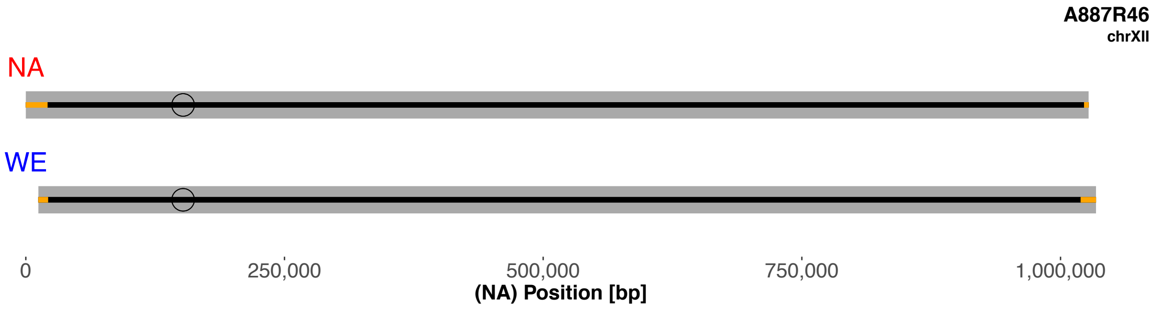
A887R46
chrXII
NA
WE
0
250,000
500,000
750,000
1,000,000
(NA) Position [bp]

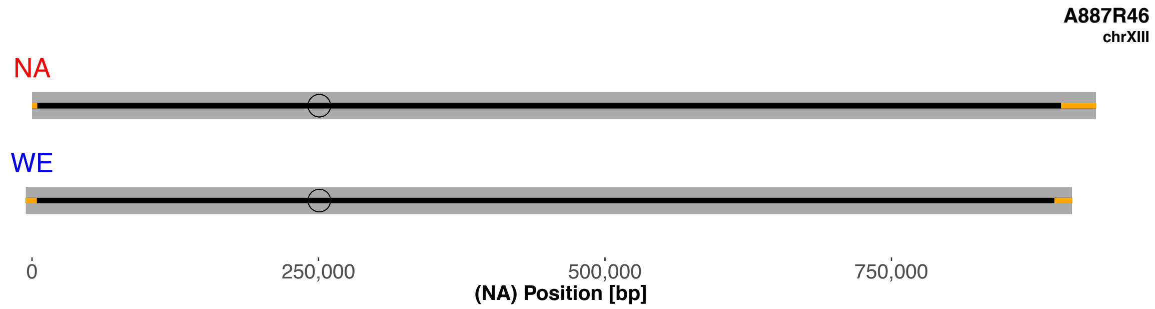
A887R46
chrXIII
NA
WE
0
250,000
500,000
750,000
(NA) Position [bp]

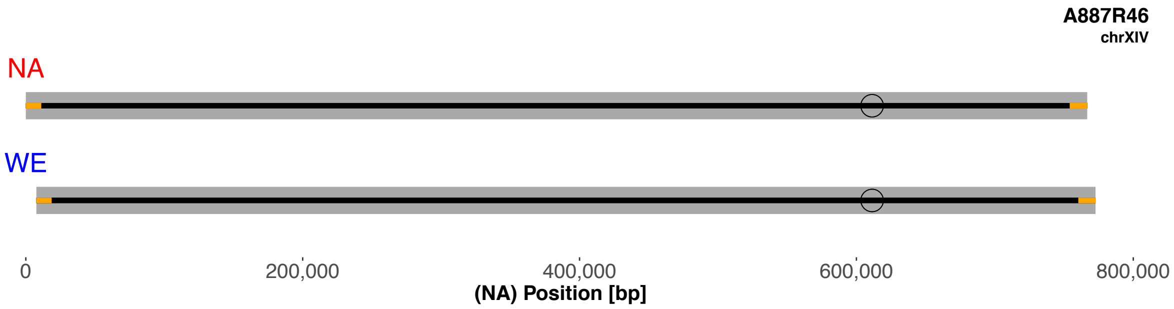
A887R46
chrXIV
NA
WE
0
200,000
400,000
600,000
800,000
(NA) Position [bp]

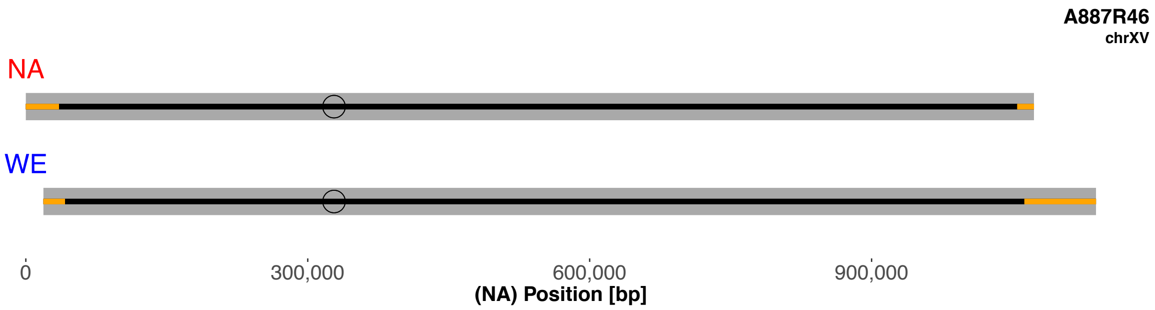
A887R46
chrXV
NA
WE
0
300,000
600,000
900,000
(NA) Position [bp]

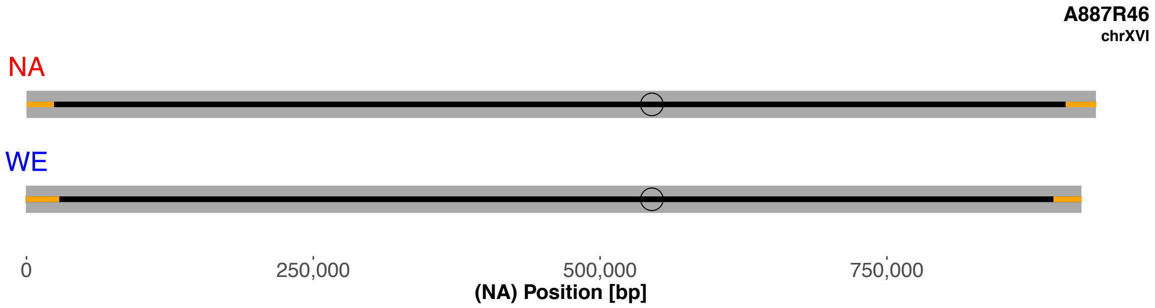
A887R46
chrXVI
NA
WE
0
250,000
500,000
750,000
(NA) Position [bp]

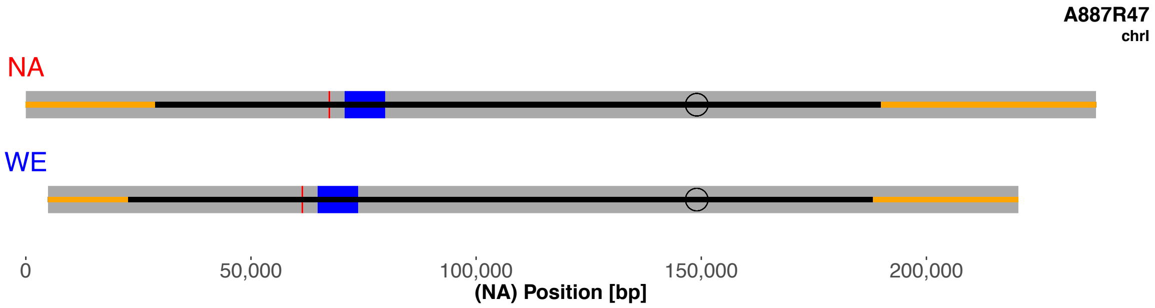
A887R47
chrI
NA
WE
0
50,000
100,000
150,000
200,000
(NA) Position [bp]

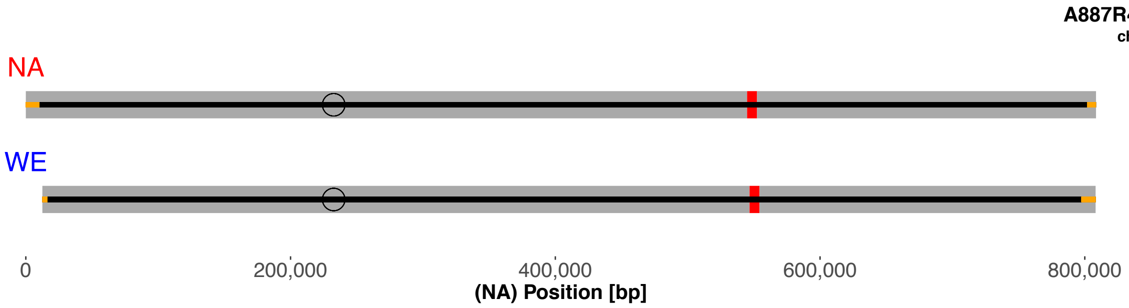
A887R47
chrII
NA
WE
0
200,000
400,000
600,000
800,000
(NA) Position [bp]

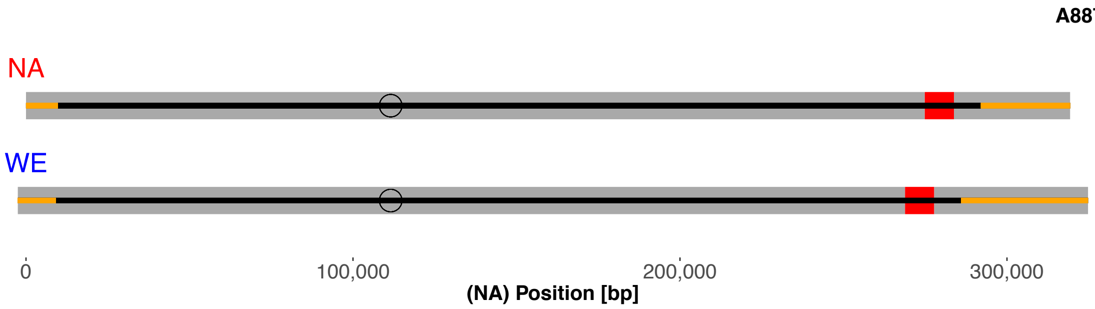
A887R47
chrIII
NA
WE
0
100,000
200,000
300,000
(NA) Position [bp]

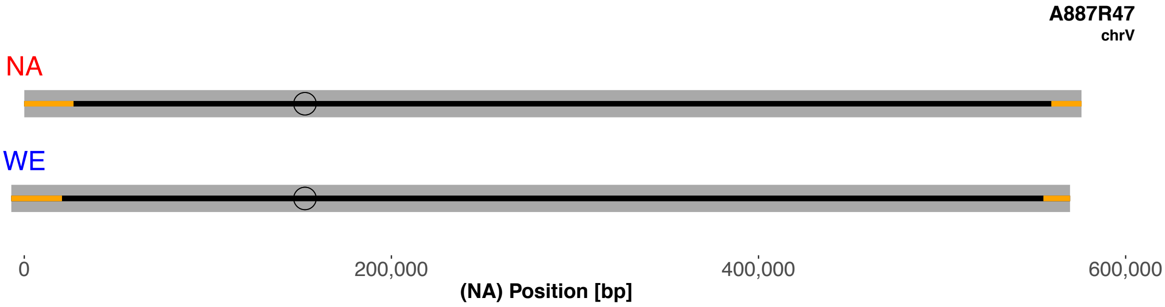
A887R47
chrV
NA
WE
0
200,000
400,000
600,000
(NA) Position [bp]

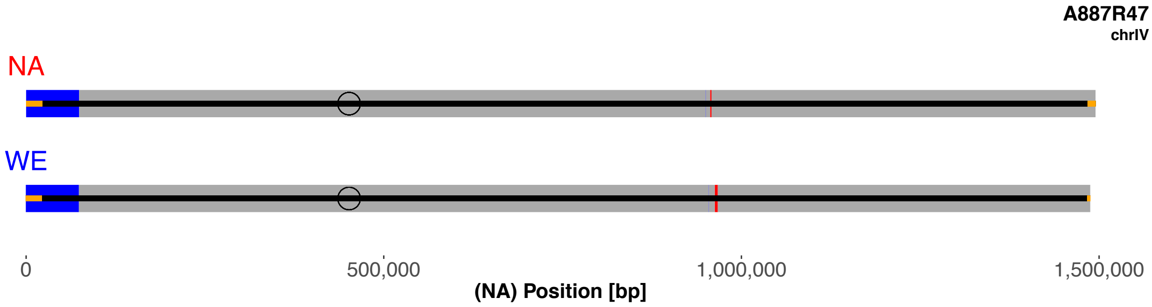
A887R47
chrIV
NA
WE
0
500,000
1,000,000
1,500,000
(NA) Position [bp]

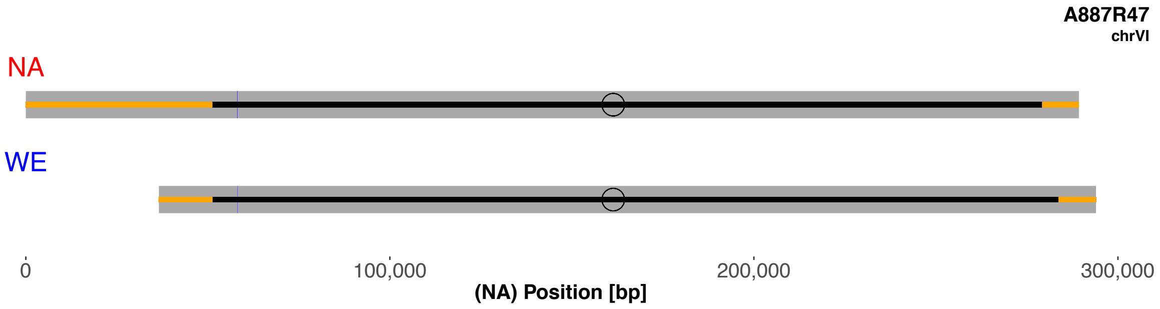
A887R47
chrVI
NA
WE
0
100,000
200,000
300,000
(NA) Position [bp]

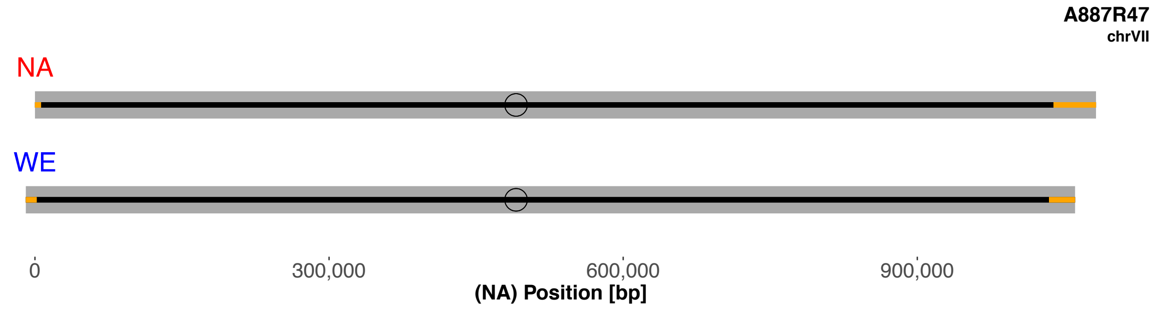
A887R47
chrVII
NA
WE
0
300,000
600,000
900,000
(NA) Position [bp]

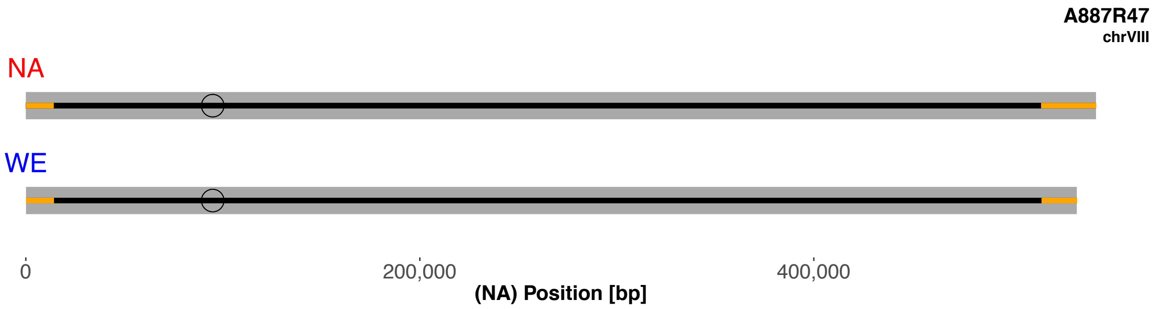
A887R47
chrVIII
NA
WE
0
200,000
400,000
(NA) Position [bp]

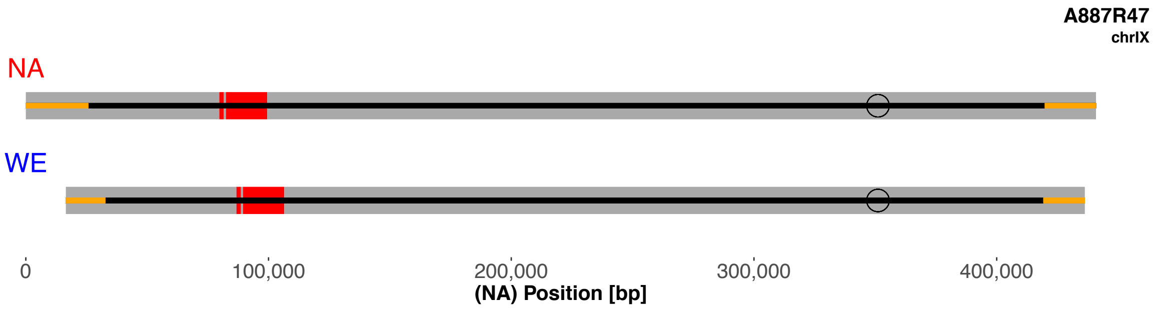
A887R47
chrIX
NA
WE
0
100,000
200,000
300,000
400,000
(NA) Position [bp]

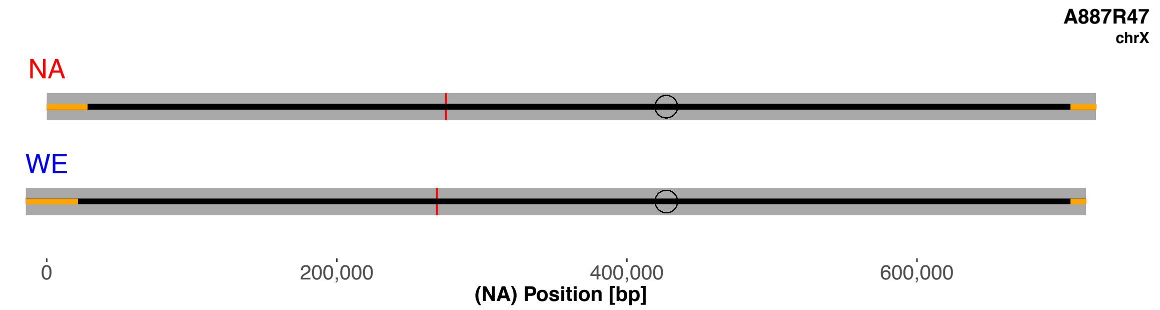
A887R47
chrX
NA
WE
0
200,000
400,000
600,000
(NA) Position [bp]

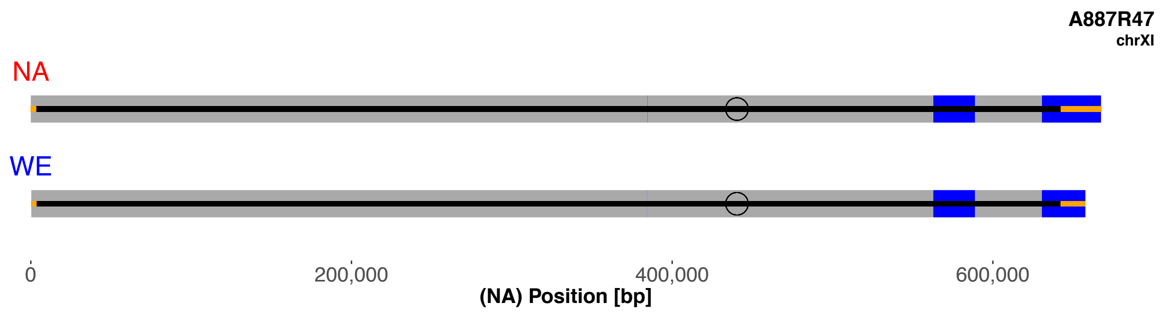
A887R47
chrXI
NA
WE
0
200,000
400,000
600,000
(NA) Position [bp]

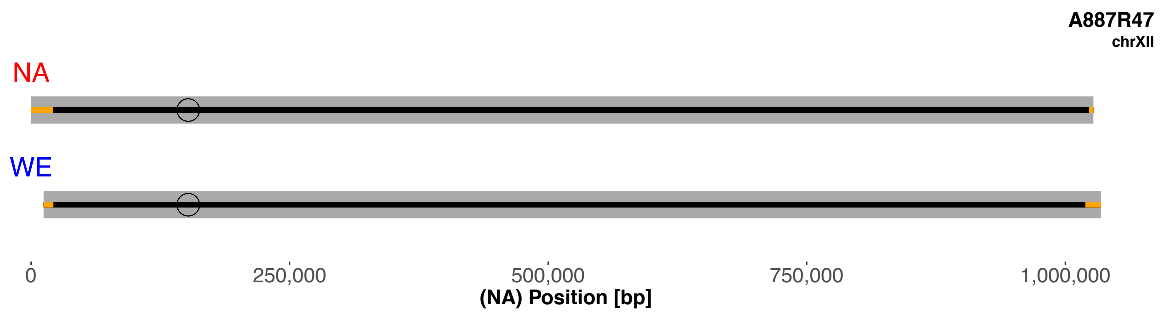
A887R47
chrXII
NA
WE
0
250,000
500,000
750,000
1,000,000
(NA) Position [bp]

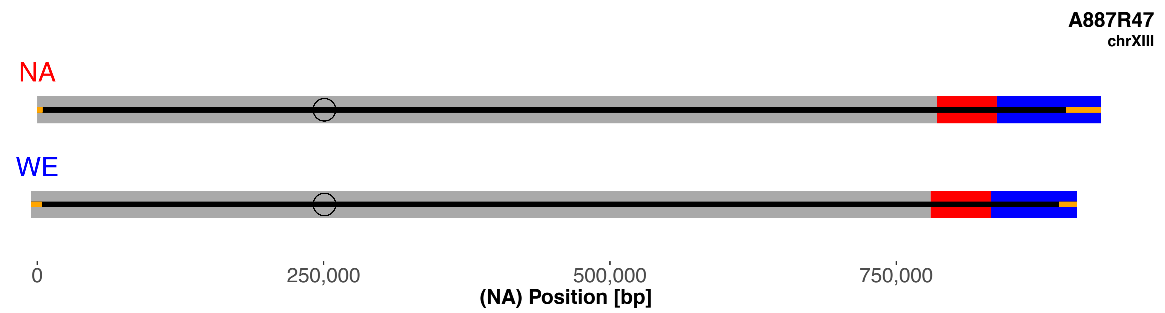
A887R47
chrXIII
NA
WE
0
250,000
500,000
750,000
(NA) Position [bp]

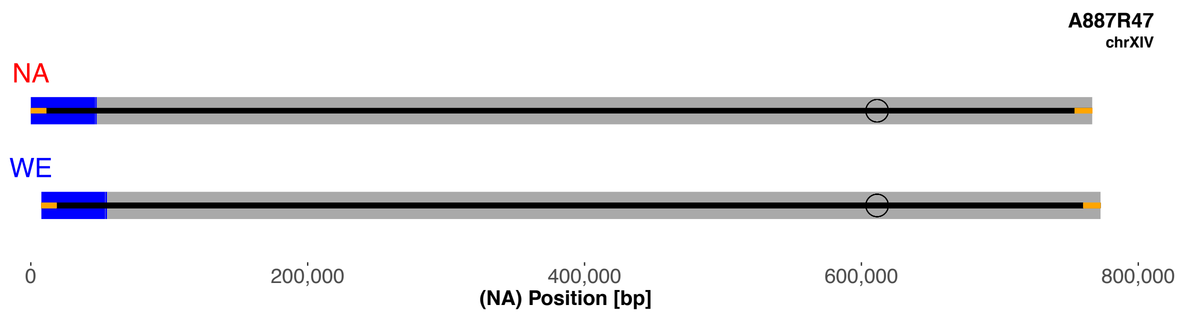
A887R47
chrXIV
NA
WE
0
200,000
400,000
600,000
800,000
(NA) Position [bp]

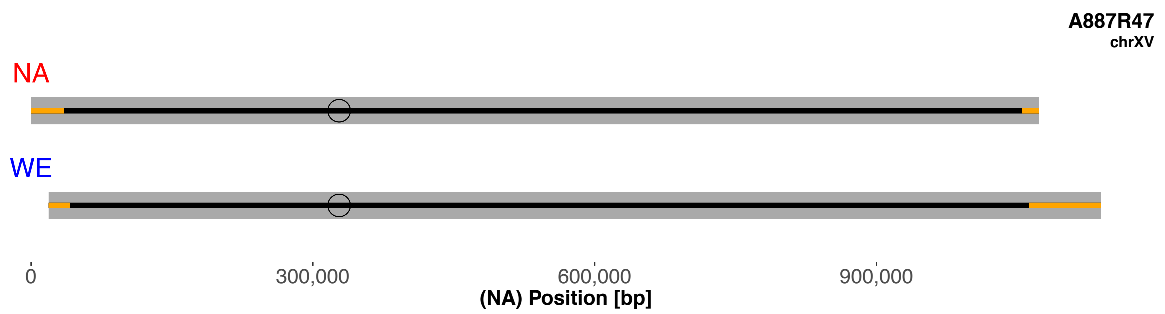
A887R47
chrXV
NA
WE
0
300,000
600,000
900,000
(NA) Position [bp]

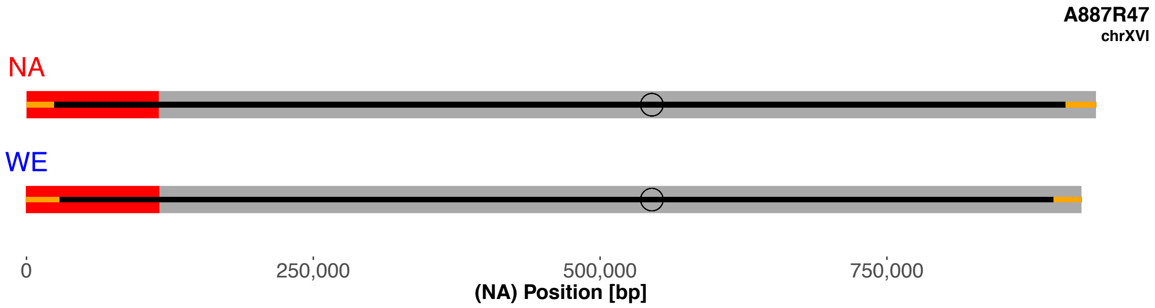
A887R47
chrXVI
NA
WE
0
250,000
500,000
750,000
(NA) Position [bp]

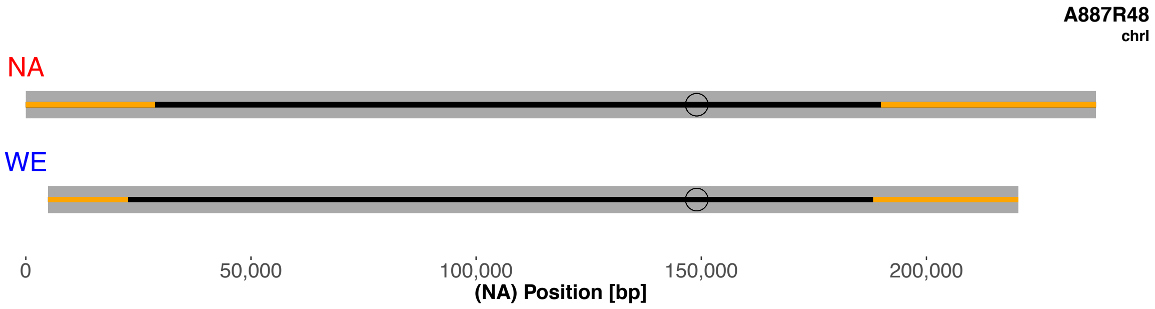
A887R48
chrI
NA
WE
0
50,000
100,000
150,000
200,000
(NA) Position [bp]

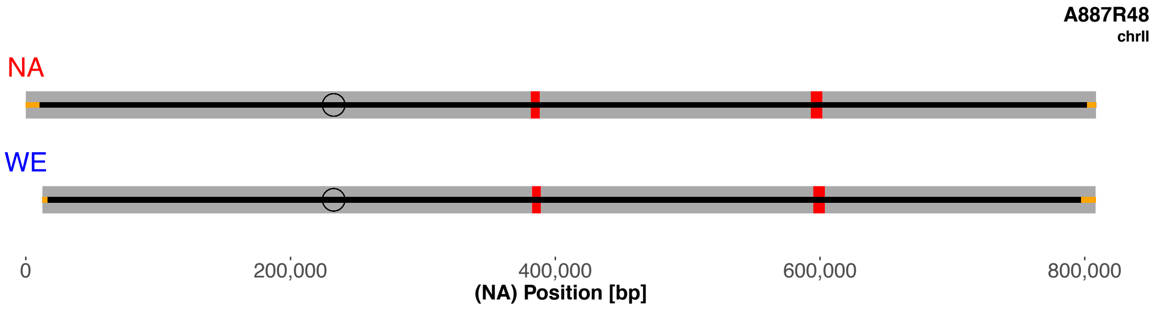
A887R48
chrII
NA
WE
0
200,000
400,000
600,000
800,000
(NA) Position [bp]

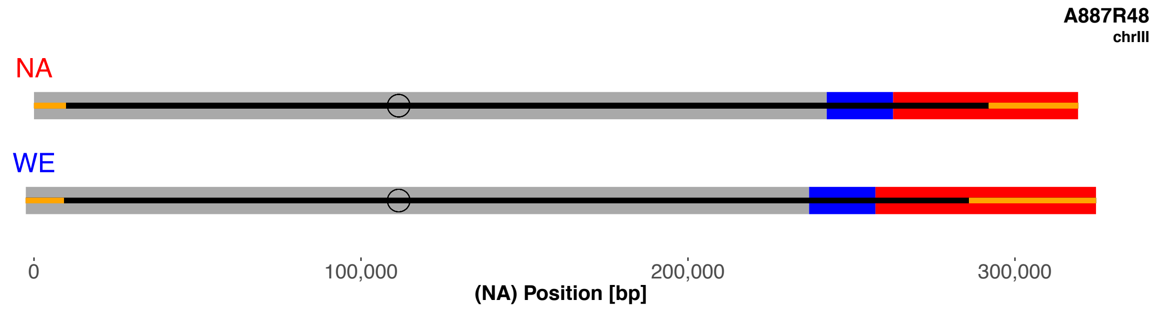
A887R48
chrIII
NA
WE
0
100,000
200,000
300,000
(NA) Position [bp]

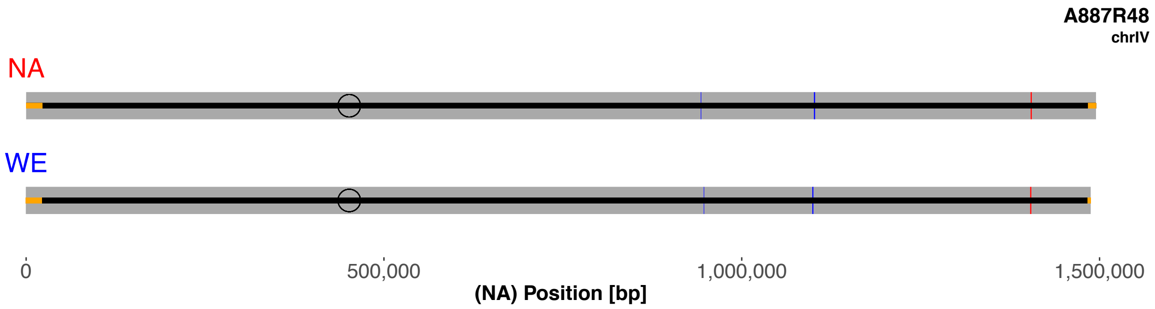
A887R48
chrIV
NA
WE
0
500,000
1,000,000
1,500,000
(NA) Position [bp]

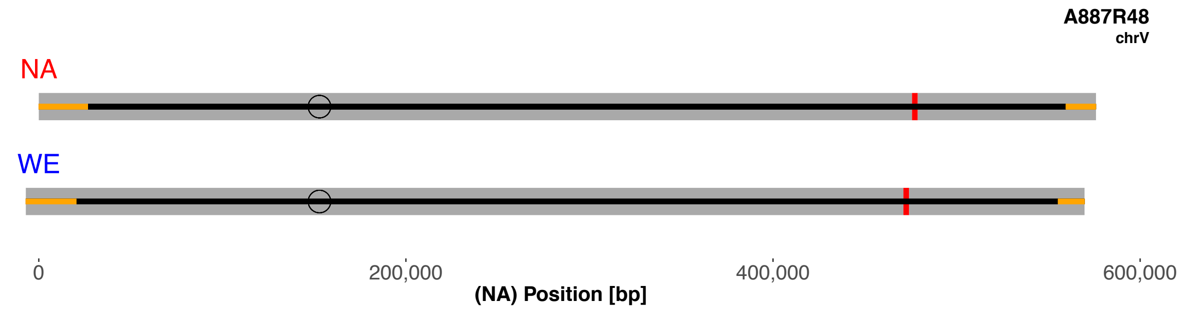
A887R48
chrV
NA
WE
0
200,000
400,000
600,000
(NA) Position [bp]

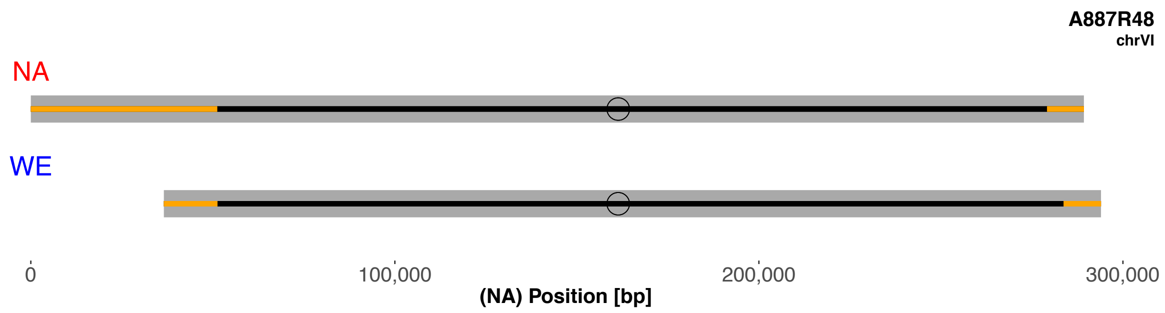
A887R48
chrVI
NA
WE
0
100,000
200,000
300,000
(NA) Position [bp]

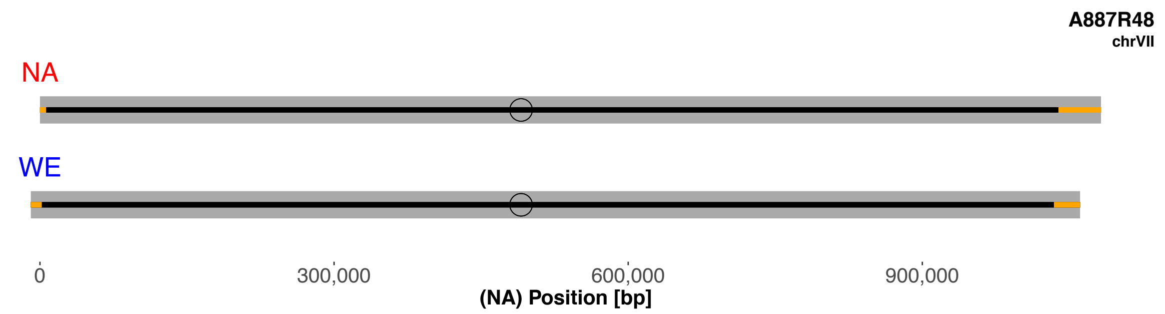
A887R48
chrVII
NA
WE
0
300,000
600,000
900,000
(NA) Position [bp]

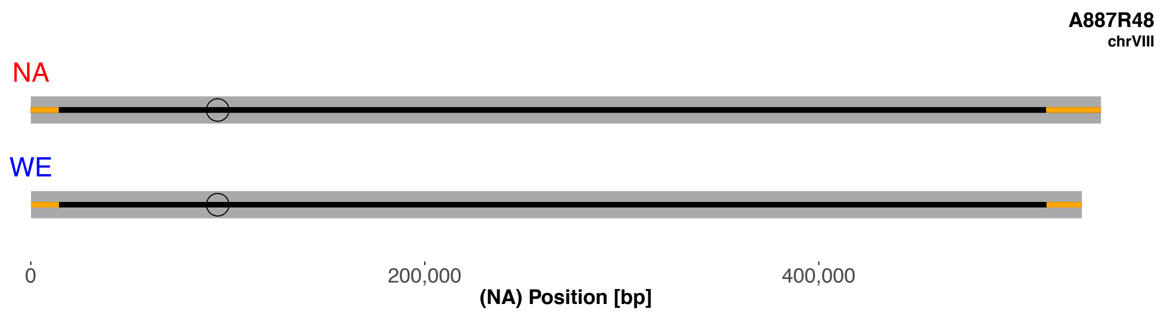
A887R48
chrVIII
NA
WE
0
200,000
400,000
(NA) Position [bp]

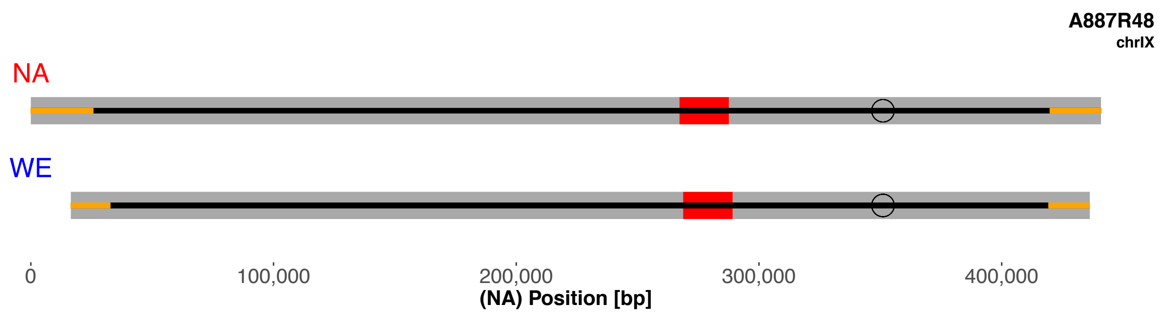
A887R48
chrIX
NA
WE
0
100,000
200,000
300,000
400,000
(NA) Position [bp]

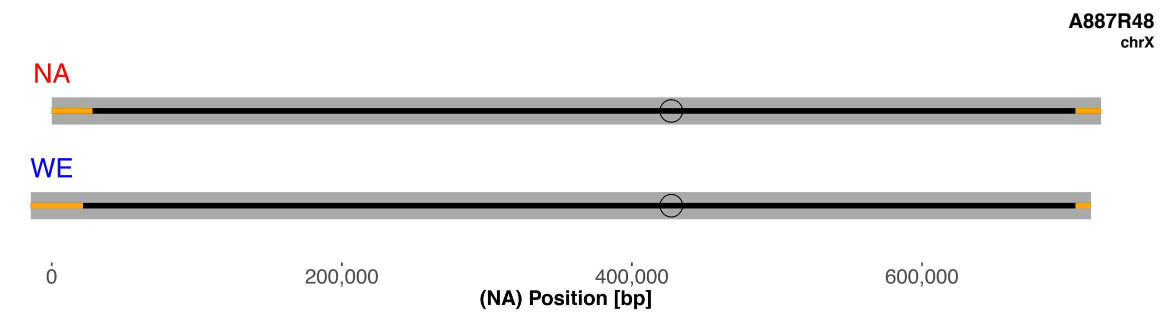
A887R48
chrX
NA
WE
0
200,000
400,000
600,000
(NA) Position [bp]

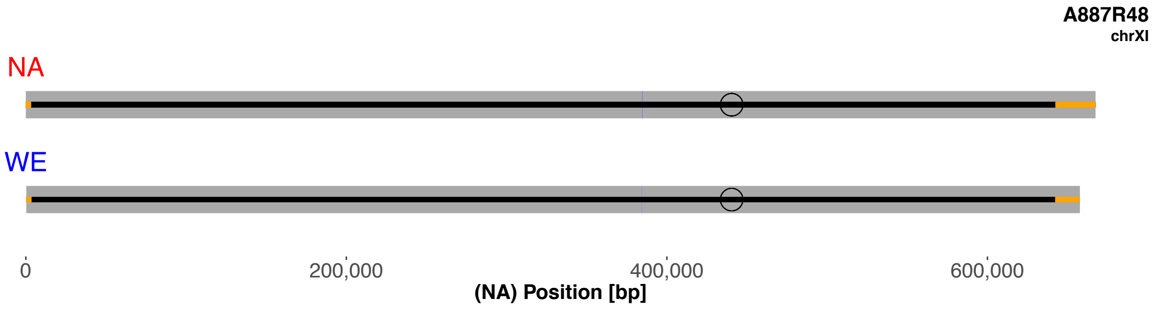
A887R48
chrXI
NA
WE
0
200,000
400,000
600,000
(NA) Position [bp]

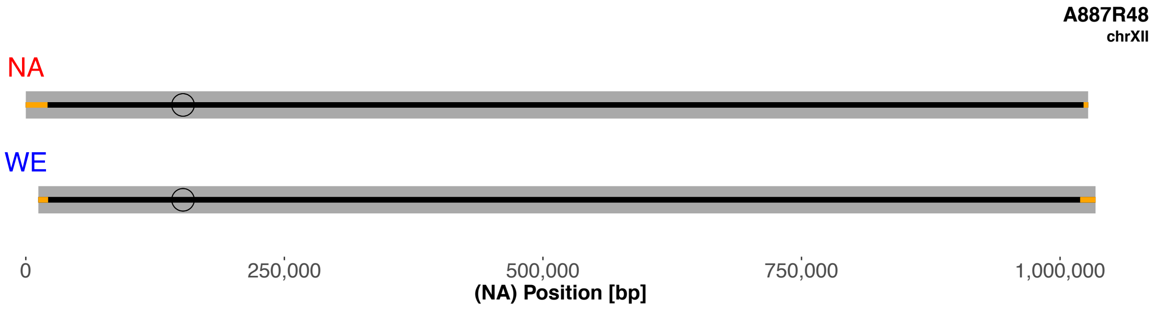
A887R48
chrXII
NA
WE
0
250,000
500,000
750,000
1,000,000
(NA) Position [bp]

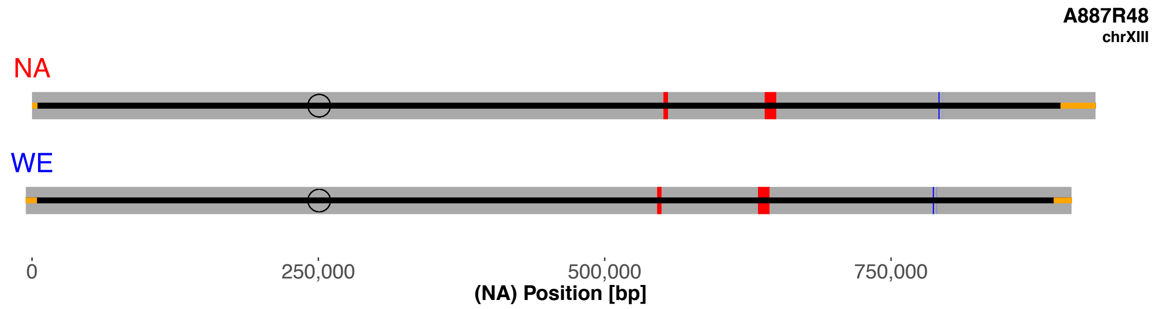
A887R48
chrXIII
NA
WE
0
250,000
500,000
750,000
(NA) Position [bp]

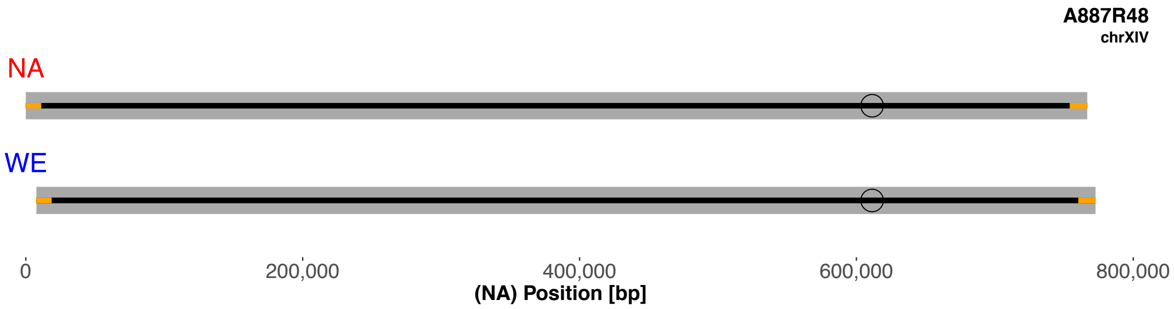
A887R48
chrXIV
NA
WE
0
200,000
400,000
600,000
800,000
(NA) Position [bp]

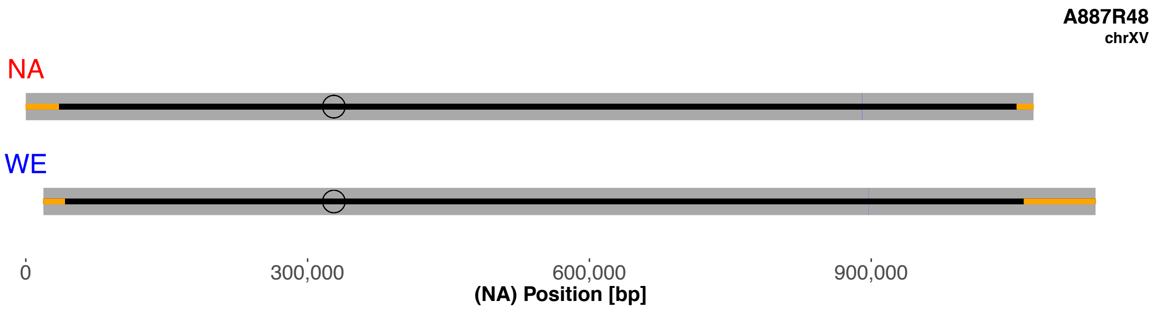
A887R48
chrXV
NA
WE
0
300,000
600,000
900,000
(NA) Position [bp]

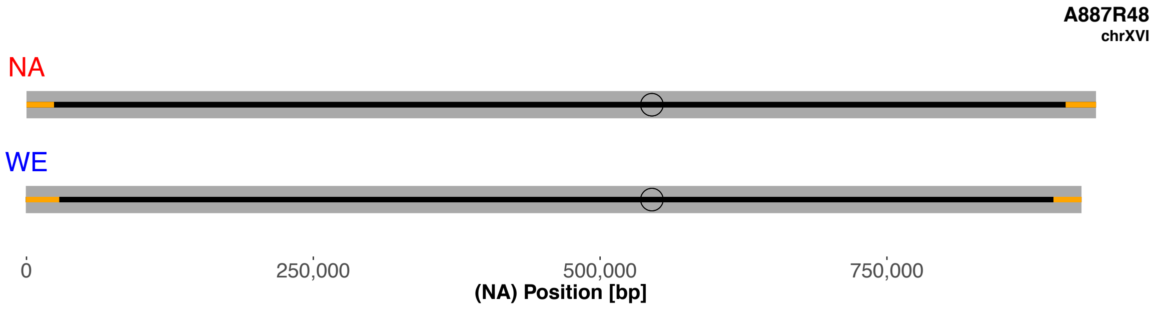
A887R48
chrXVI
NA
WE
0
250,000
500,000
750,000
(NA) Position [bp]

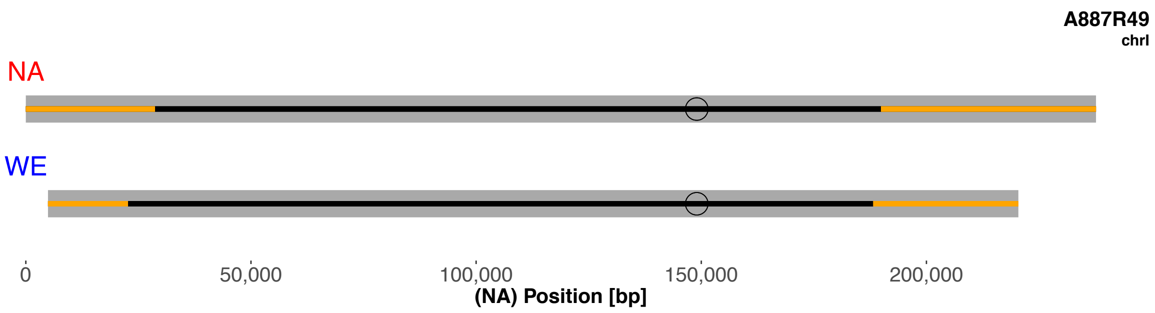
A887R49
chrI
NA
WE
0
50,000
100,000
150,000
200,000
(NA) Position [bp]

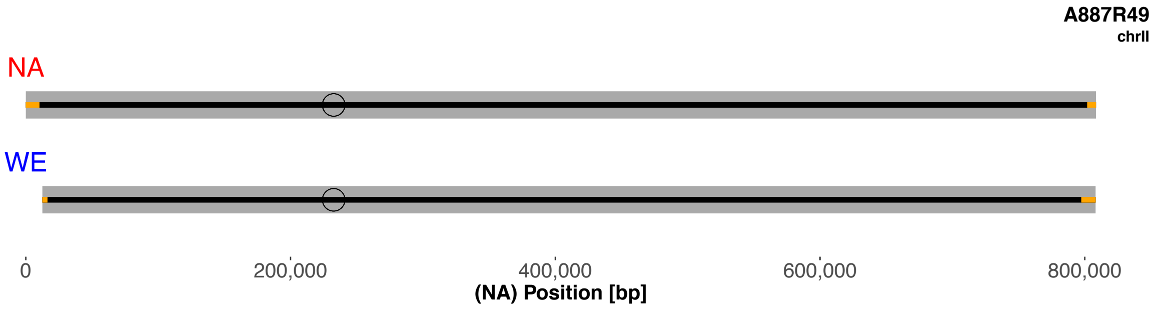
A887R49
chrII
NA
WE
0
200,000
400,000
600,000
800,000
(NA) Position [bp]

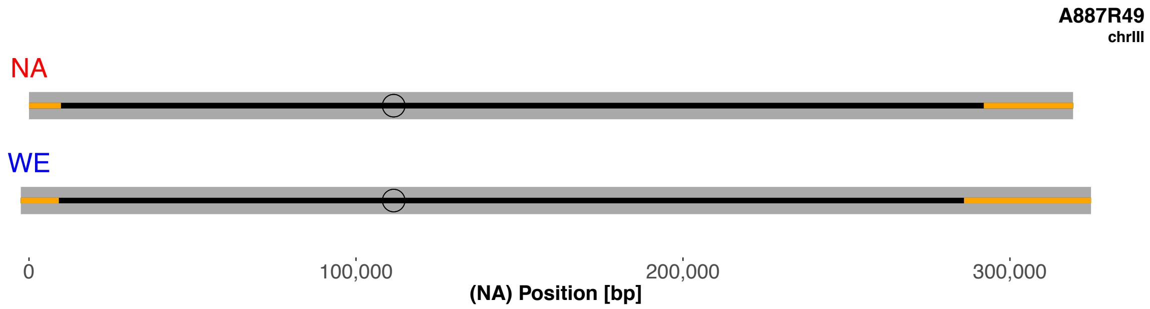
A887R49
chrIII
NA
WE
0
100,000
200,000
300,000
(NA) Position [bp]

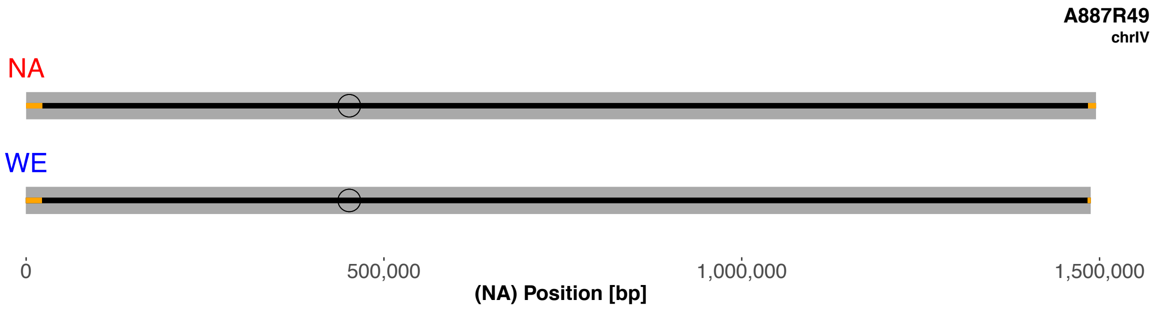
A887R49
chrIV
NA
WE
0
500,000
1,000,000
1,500,000
(NA) Position [bp]

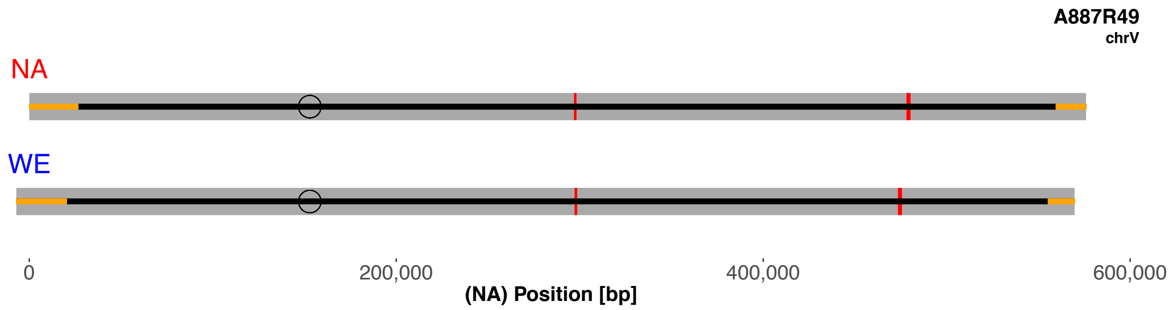
A887R49
chrV
NA
WE
0
200,000
400,000
600,000
(NA) Position [bp]

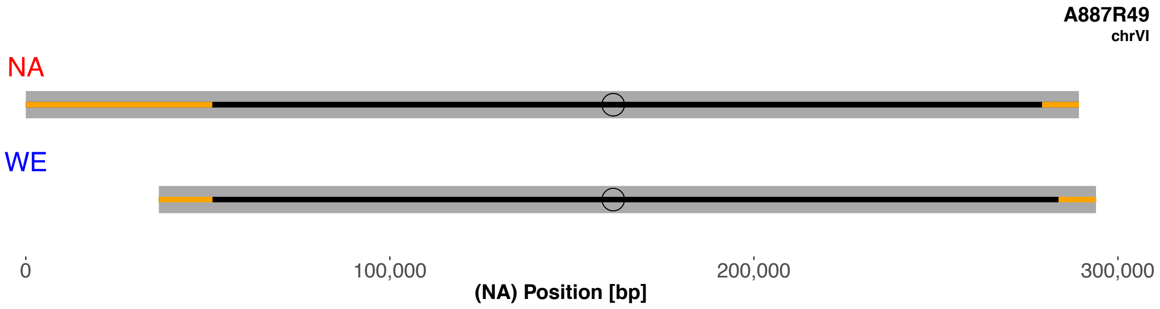
A887R49
chrVI
NA
WE
0
100,000
200,000
300,000
(NA) Position [bp]

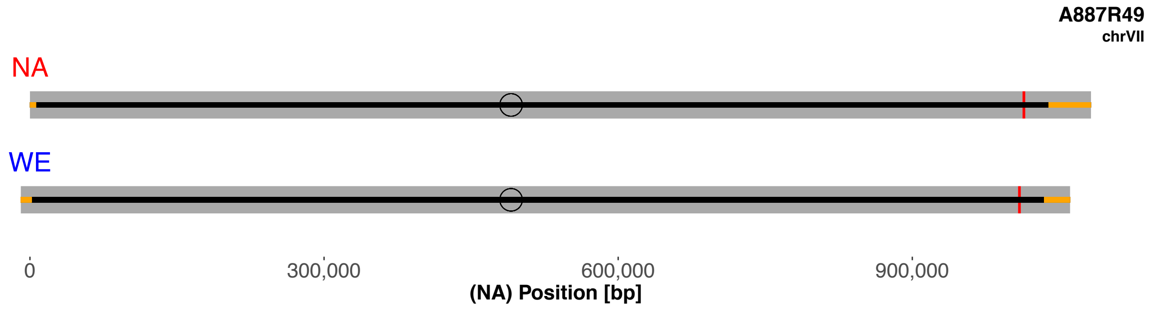
A887R49
chrVII
NA
WE
0
300,000
600,000
900,000
(NA) Position [bp]

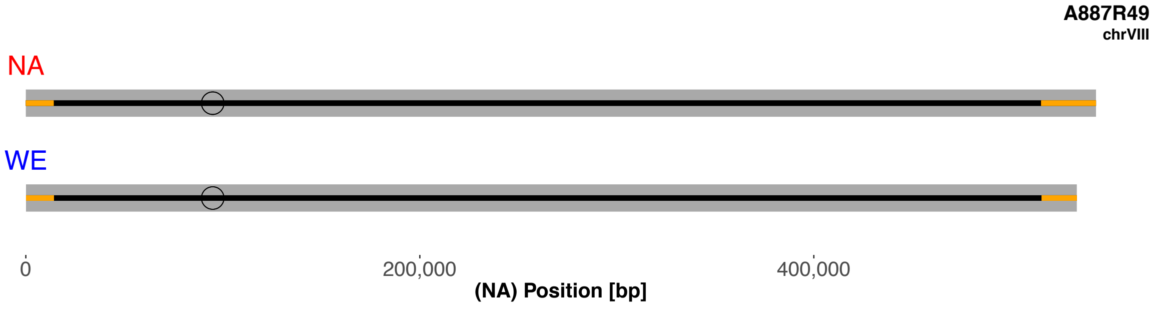
A887R49
chrVIII
NA
WE
0
200,000
400,000
(NA) Position [bp]

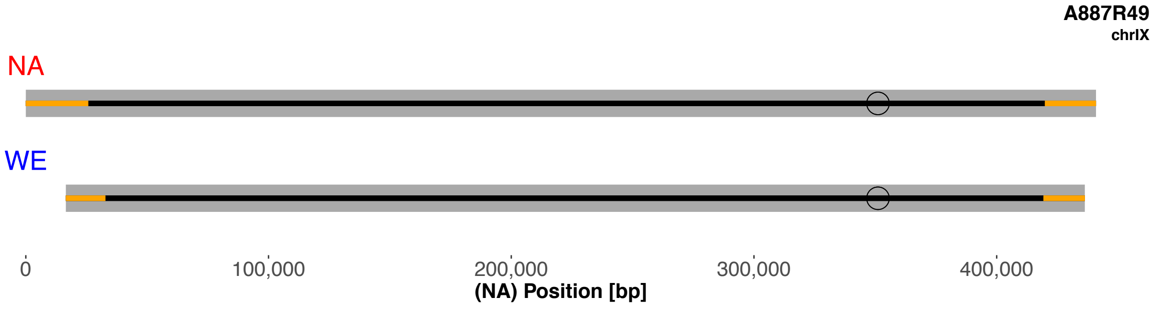
A887R49
chrIX
NA
WE
0
100,000
200,000
300,000
400,000
(NA) Position [bp]

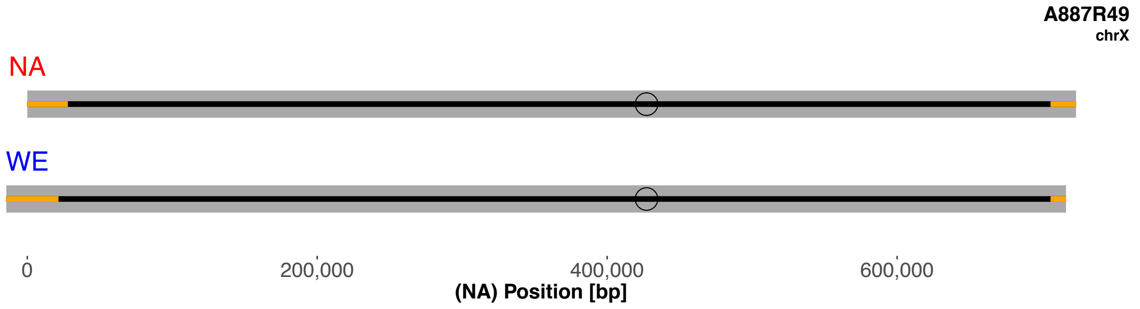
A887R49
chrX
NA
WE
0
200,000
400,000
600,000
(NA) Position [bp]

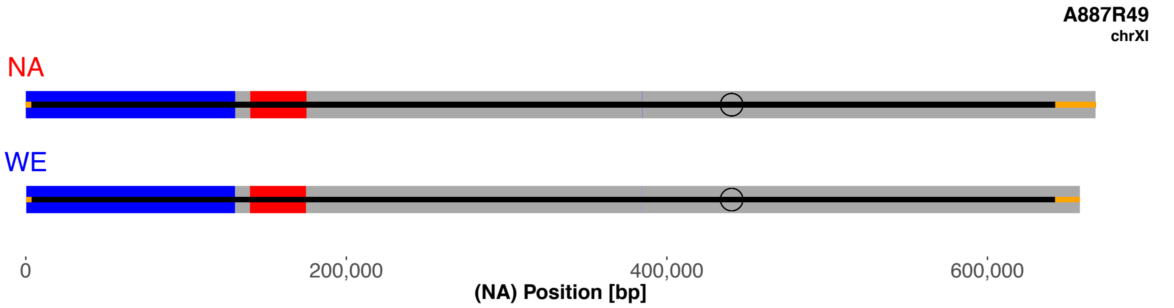
A887R49
chrXI
NA
WE
0
200,000
400,000
600,000
(NA) Position [bp]

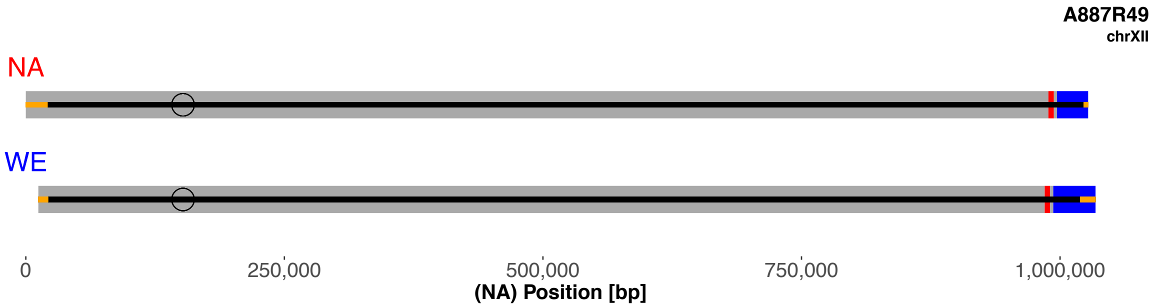
A887R49
chrXII
NA
WE
0
250,000
500,000
750,000
1,000,000
(NA) Position [bp]

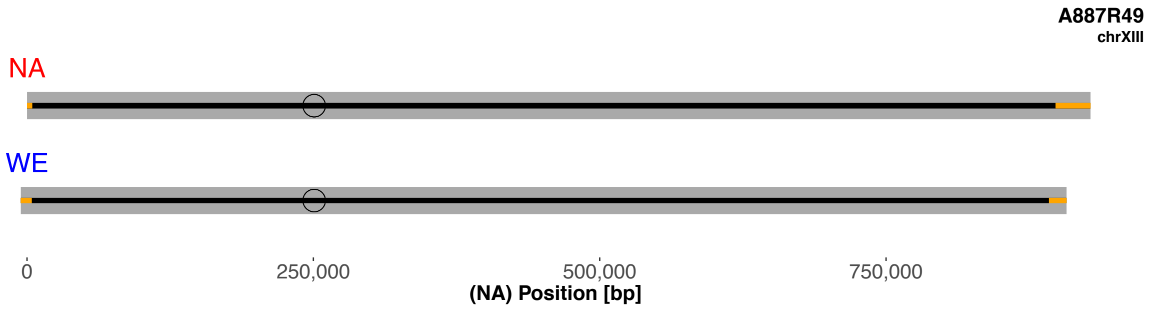
A887R49
chrXIII
NA
WE
0
250,000
500,000
750,000
(NA) Position [bp]

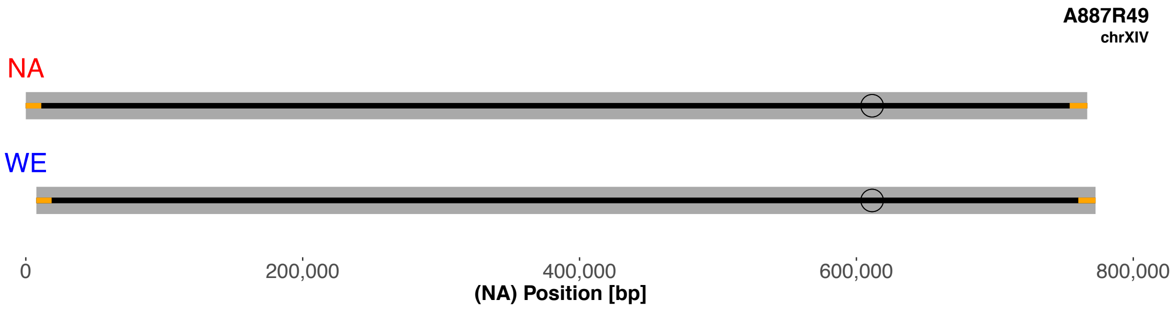
A887R49
chrXIV
NA
WE
0
200,000
400,000
600,000
800,000
(NA) Position [bp]

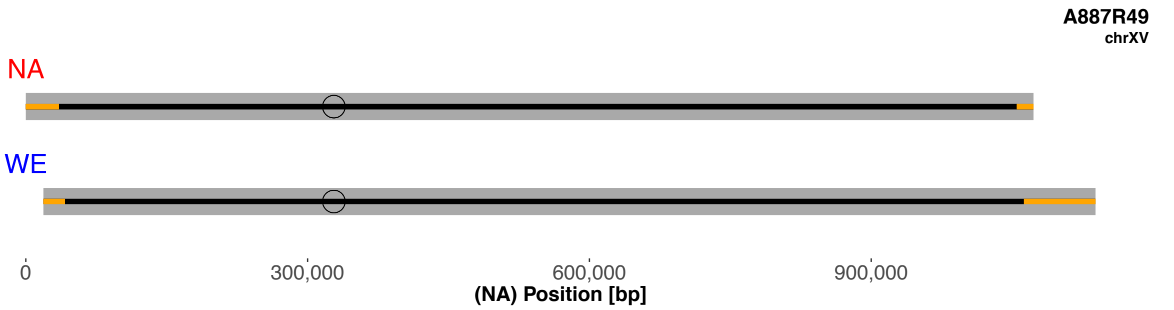
A887R49
chrXV
NA
WE
0
300,000
600,000
900,000
(NA) Position [bp]

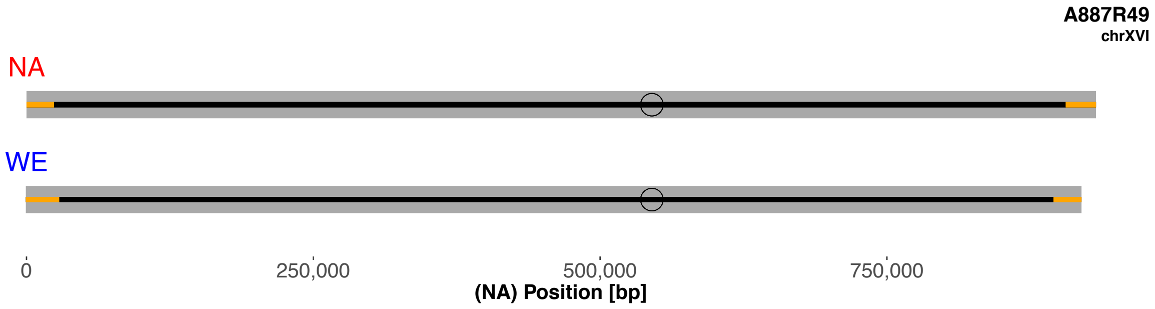
A887R49
chrXVI
NA
WE
0
250,000
500,000
750,000
(NA) Position [bp]

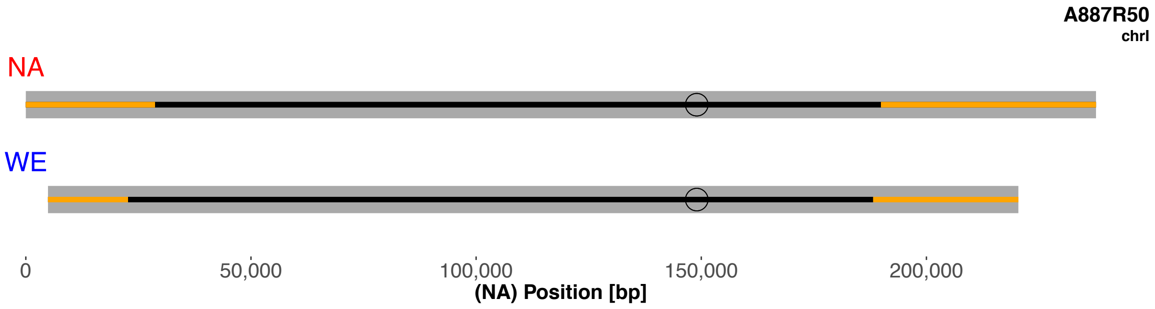
A887R50
chrI
NA
WE
0
50,000
100,000
150,000
200,000
(NA) Position [bp]

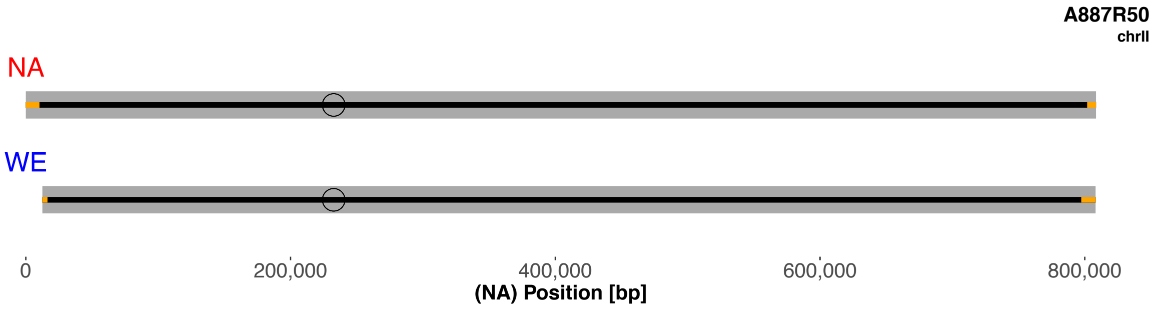
A887R50
chrII
NA
WE
0
200,000
400,000
600,000
800,000
(NA) Position [bp]

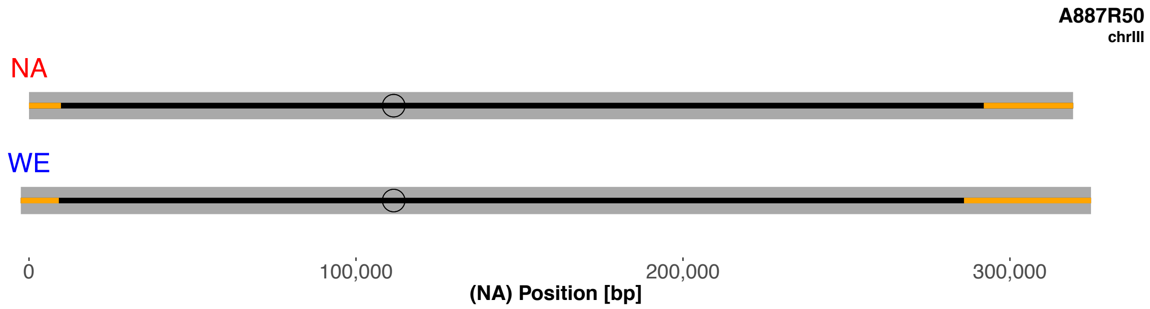
A887R50
chrIII
NA
WE
0
100,000
200,000
300,000
(NA) Position [bp]

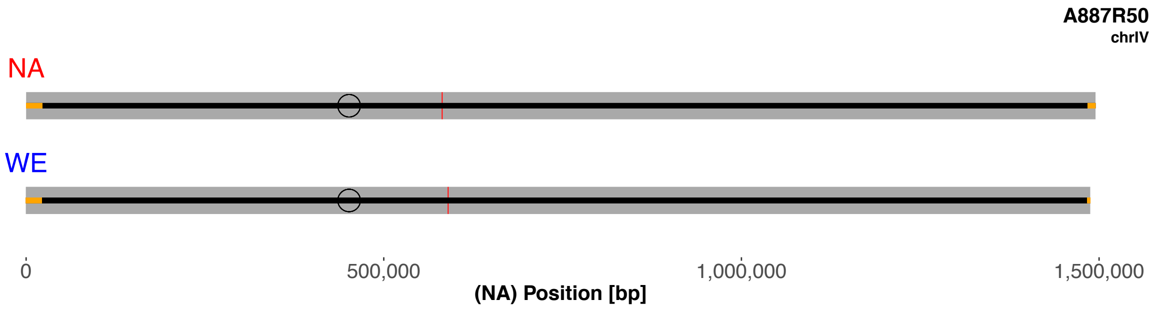
A887R50
chrIV
NA
WE
0
500,000
1,000,000
1,500,000
(NA) Position [bp]

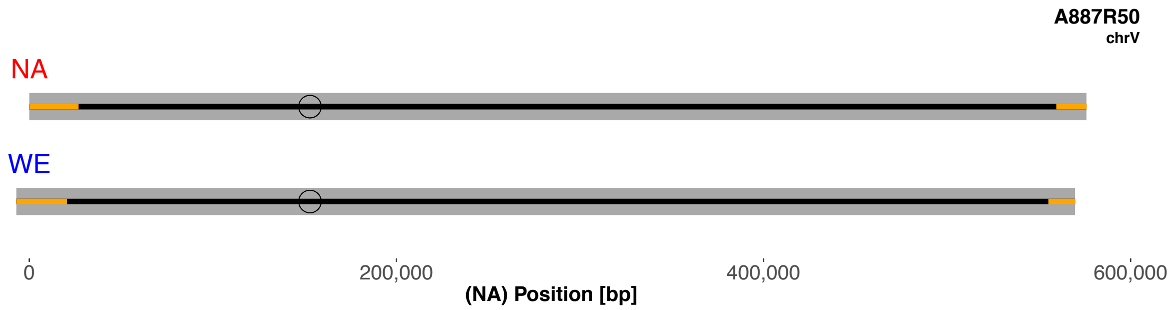
A887R50
chrV
NA
WE
0
200,000
400,000
600,000
(NA) Position [bp]

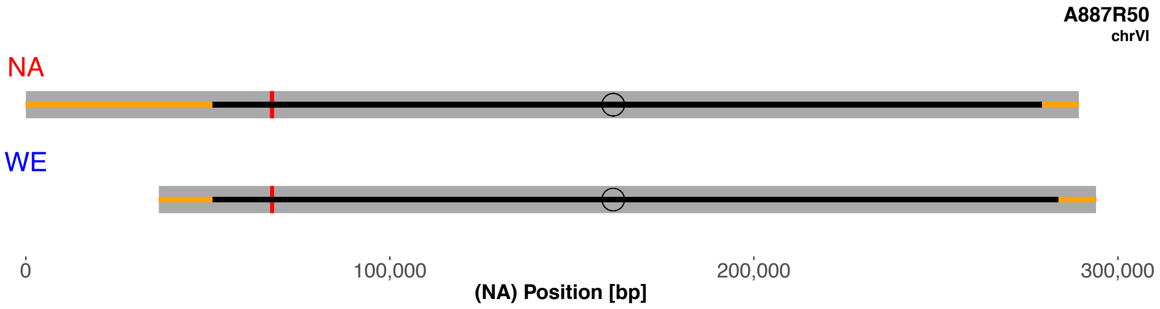
A887R50
chrVI
NA
WE
0
100,000
200,000
300,000
(NA) Position [bp]

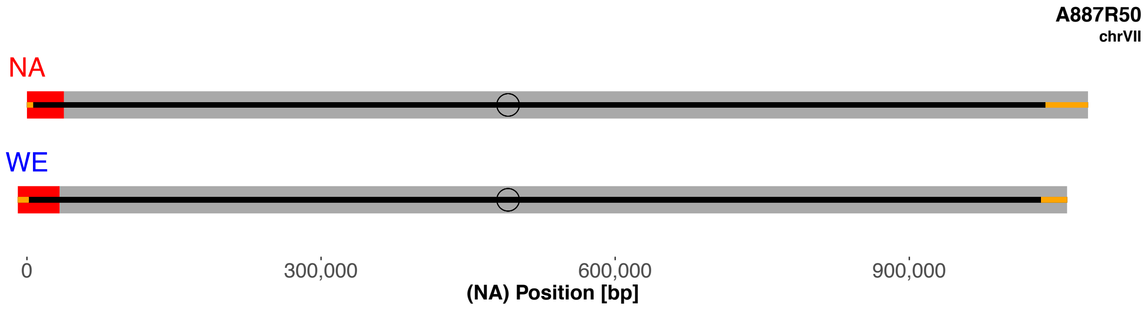
A887R50
chrVII
NA
WE
0
300,000
600,000
900,000
(NA) Position [bp]

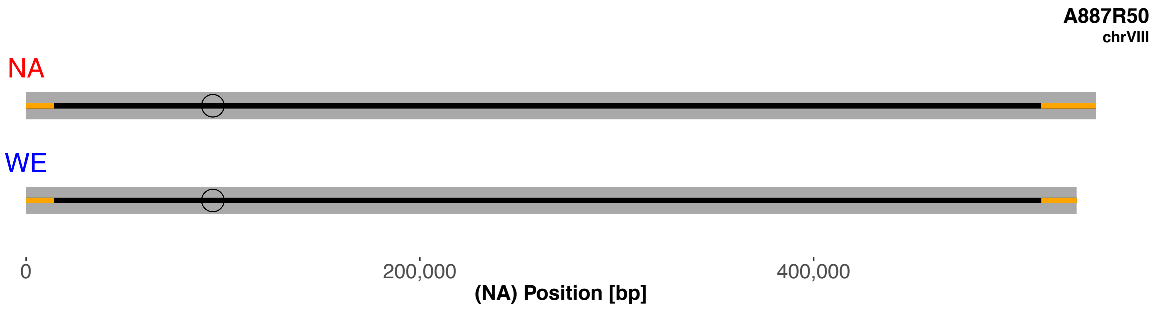
A887R50
chrVIII
NA
WE
0
200,000
400,000
(NA) Position [bp]

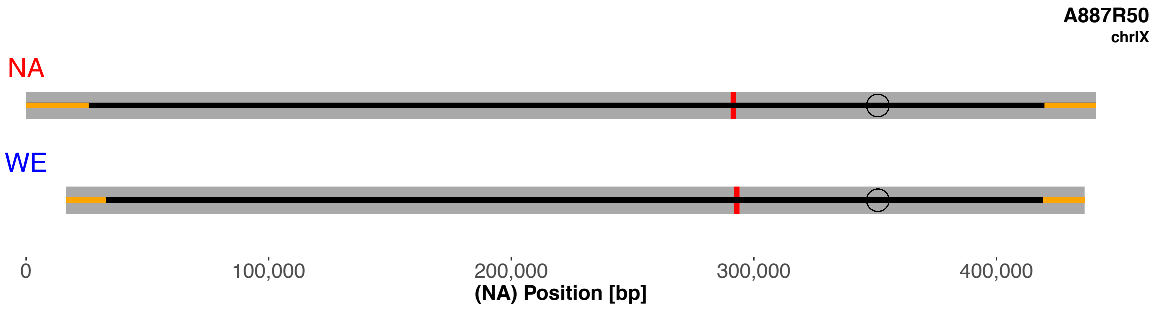
A887R50
chrIX
NA
WE
0
100,000
200,000
300,000
400,000
(NA) Position [bp]

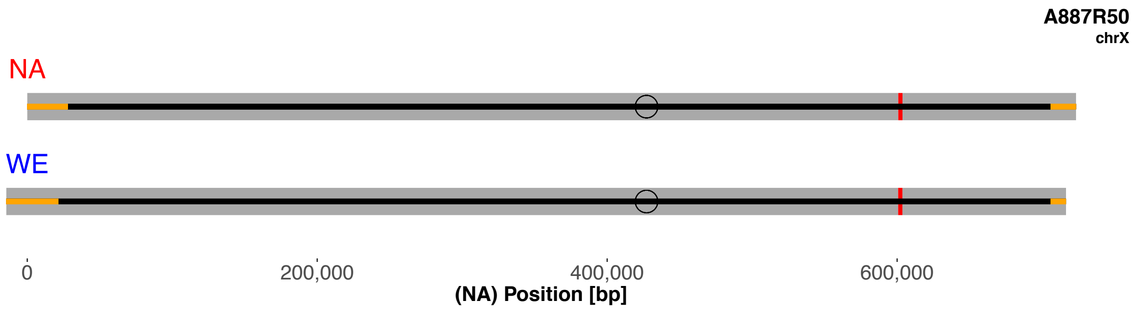
A887R50
chrX
NA
WE
0
200,000
400,000
600,000
(NA) Position [bp]

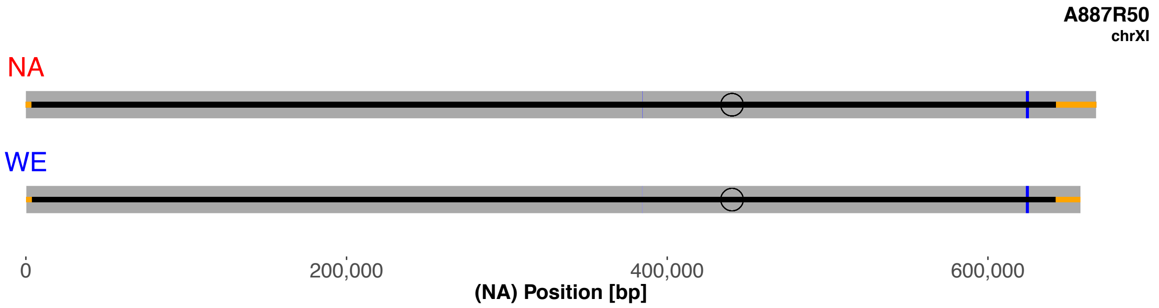
A887R50
chrXI
NA
WE
0
200,000
400,000
600,000
(NA) Position [bp]

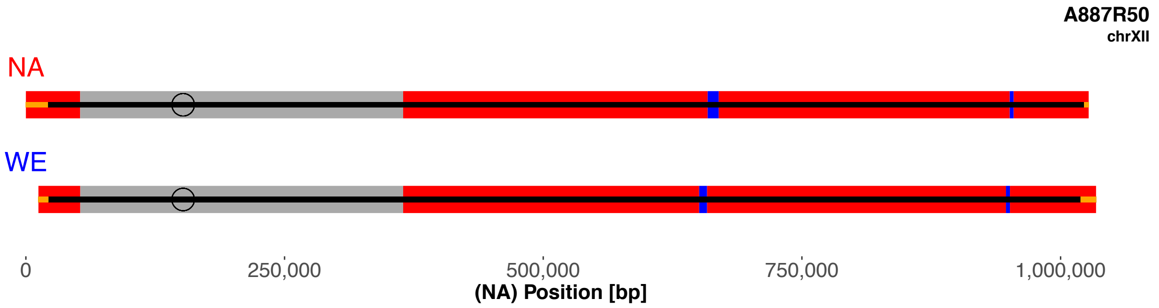
A887R50
chrXII
NA
WE
0
250,000
500,000
750,000
1,000,000
(NA) Position [bp]

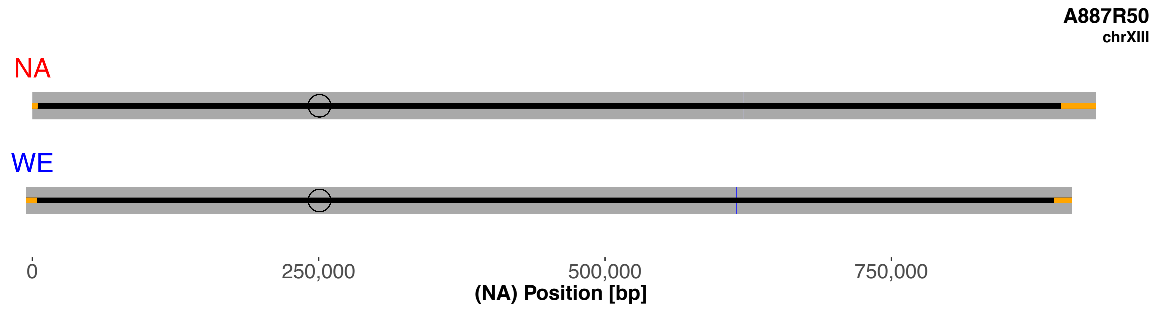
A887R50
chrXIII
NA
WE
0
250,000
500,000
750,000
(NA) Position [bp]

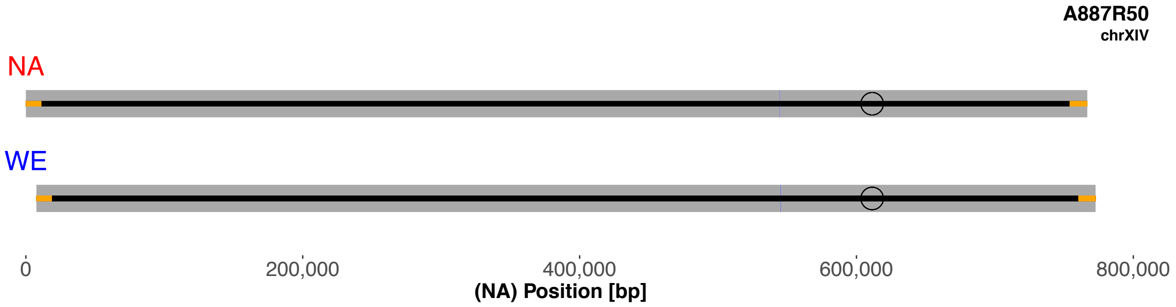
A887R50
chrXIV
NA
WE
0
200,000
400,000
600,000
800,000
(NA) Position [bp]

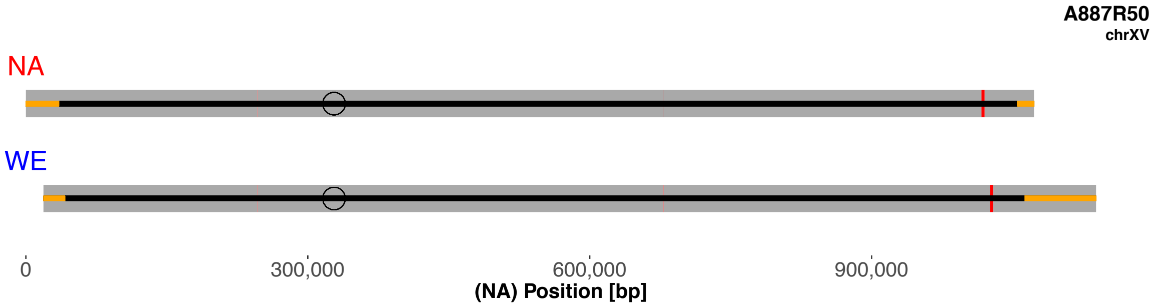
A887R50
chrXV
NA
WE
0
300,000
600,000
900,000
(NA) Position [bp]

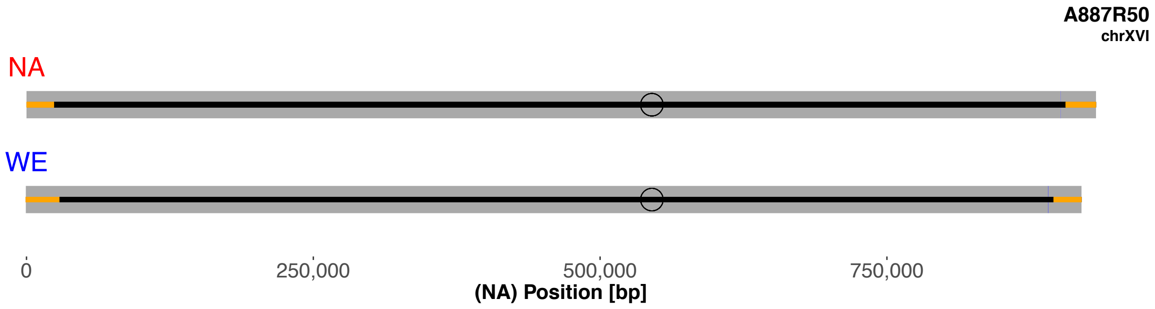
A887R50
chrXVI
NA
WE
0
250,000
500,000
750,000
(NA) Position [bp]

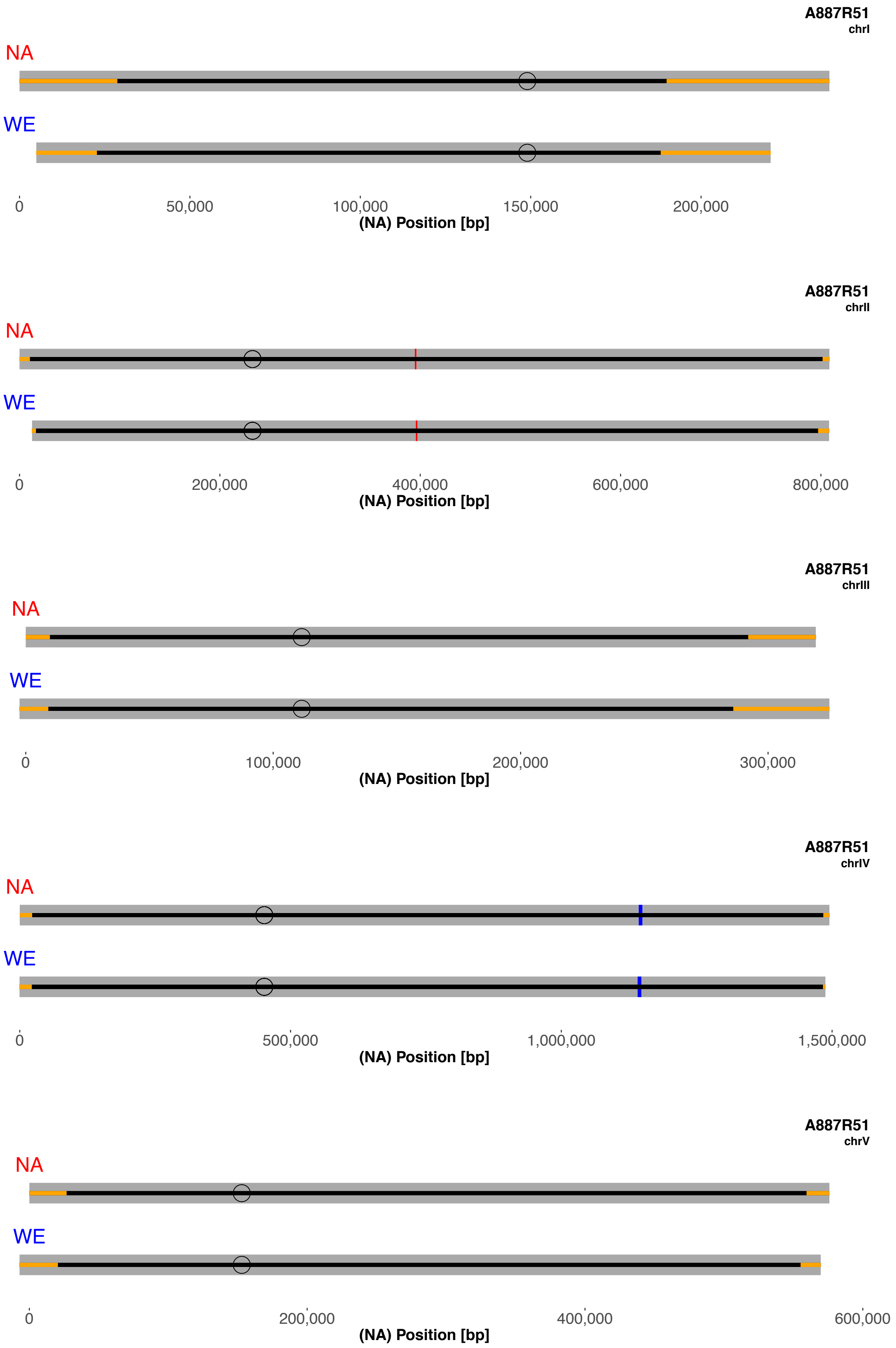
A887R51
chrI
NA
WE
0
50,000
100,000
150,000
200,000
(NA) Position [bp]
A887R51
chrII
NA
WE
0
200,000
400,000
600,000
800,000
(NA) Position [bp]
A887R51
chrIII
NA
WE
0
100,000
200,000
300,000
(NA) Position [bp]
A887R51
chrIV
NA
WE
0
500,000
1,000,000
1,500,000
(NA) Position [bp]
A887R51
chrV
NA
WE
0
200,000
400,000
600,000
(NA) Position [bp]

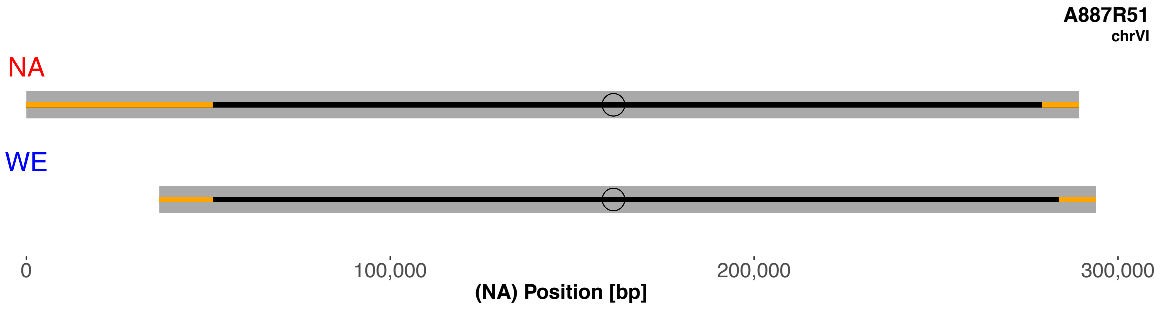
A887R51
chrVI
NA
WE
0
100,000
200,000
300,000
(NA) Position [bp]

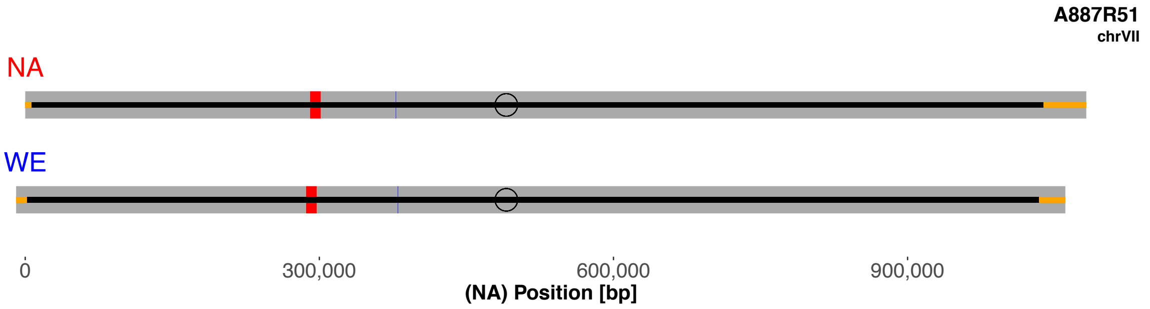
A887R51
chrVII
NA
WE
0
300,000
600,000
900,000
(NA) Position [bp]

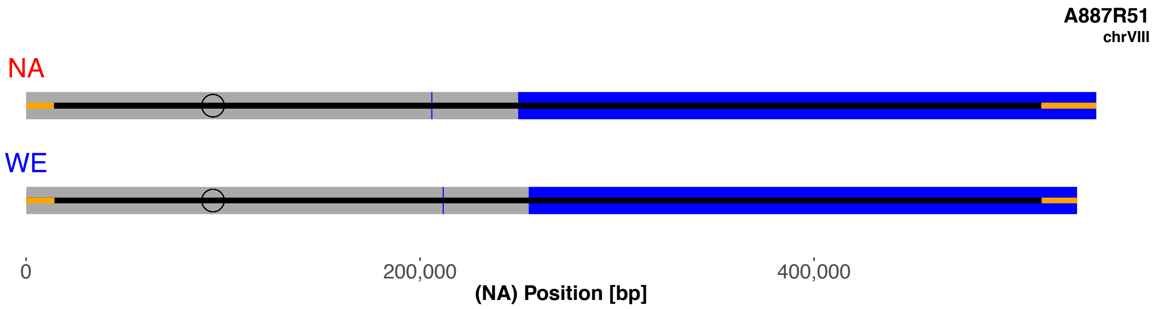
A887R51
chrVIII
NA
WE
0
200,000
400,000
(NA) Position [bp]

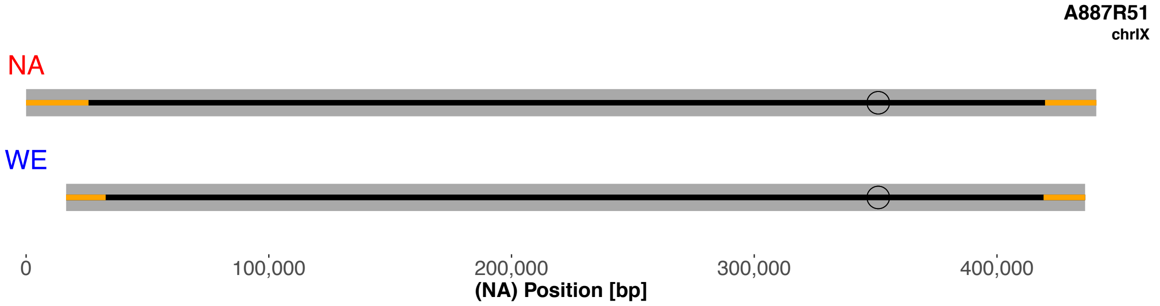
A887R51
chrIX
NA
WE
0
100,000
200,000
300,000
400,000
(NA) Position [bp]

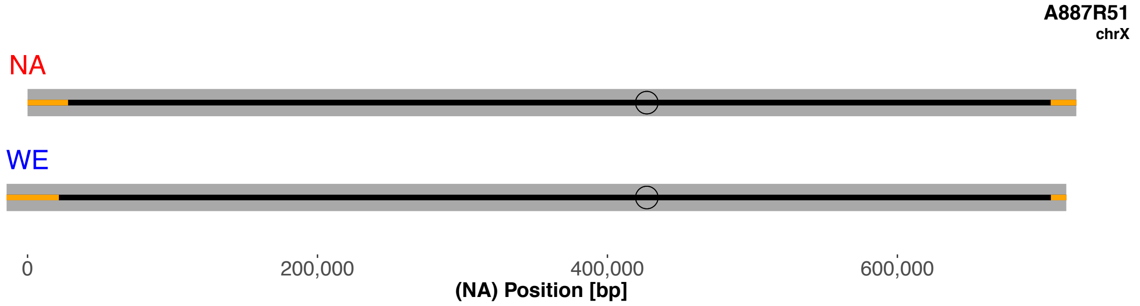
A887R51
chrX
NA
WE
0
200,000
400,000
600,000
(NA) Position [bp]

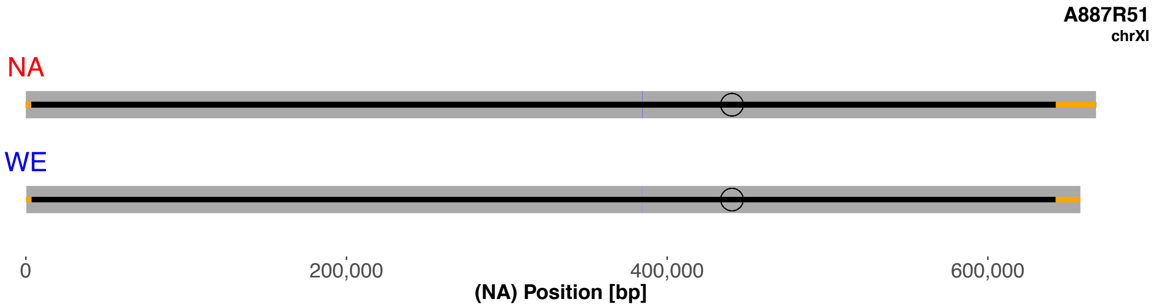
A887R51
chrXI
NA
WE
0
200,000
400,000
600,000
(NA) Position [bp]

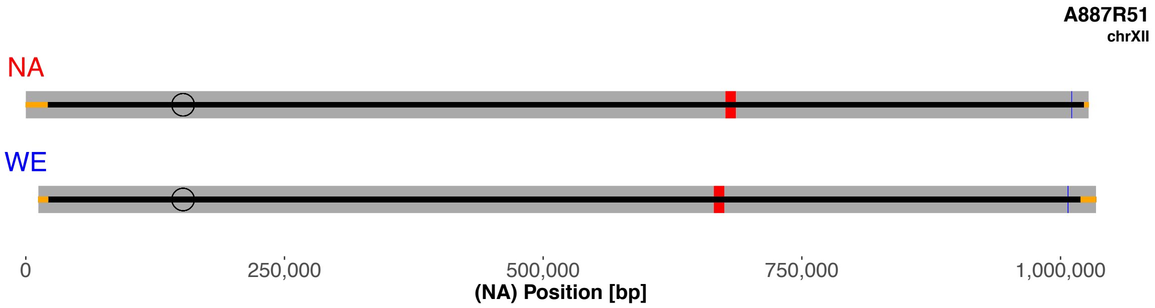
A887R51
chrXII
NA
WE
0
250,000
500,000
750,000
1,000,000
(NA) Position [bp]

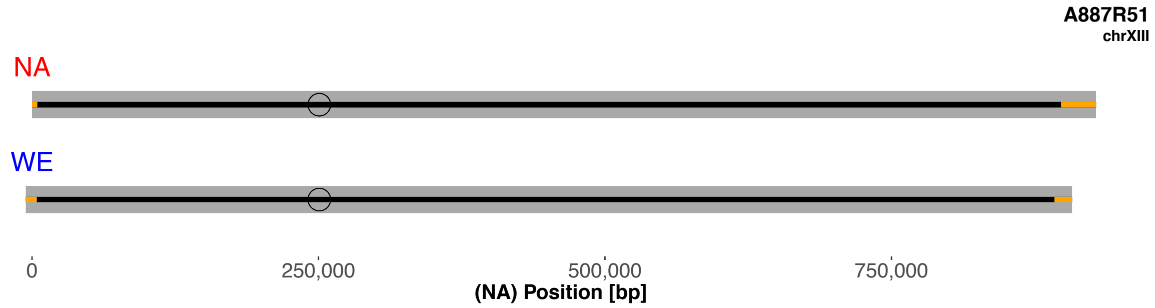
A887R51
chrXIII
NA
WE
0
250,000
500,000
750,000
(NA) Position [bp]

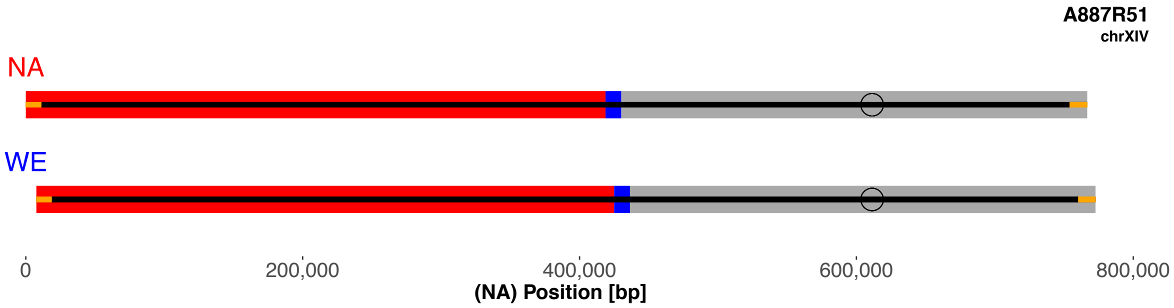
A887R51
chrXIV
NA
WE
0
200,000
400,000
600,000
800,000
(NA) Position [bp]

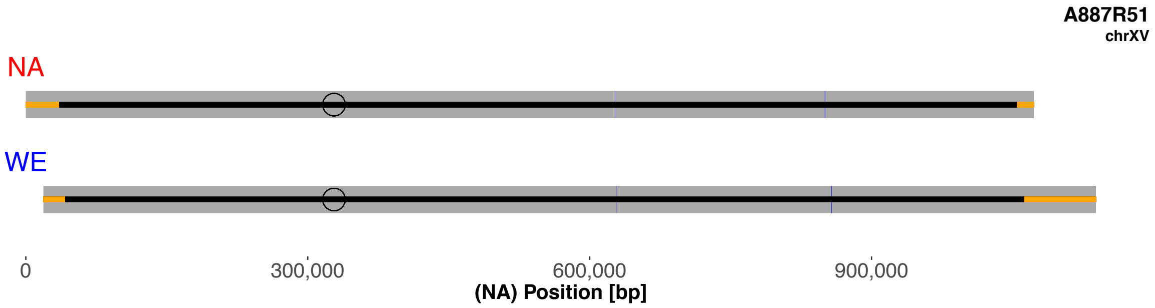
A887R51
chrXV
NA
WE
0
300,000
600,000
900,000
(NA) Position [bp]

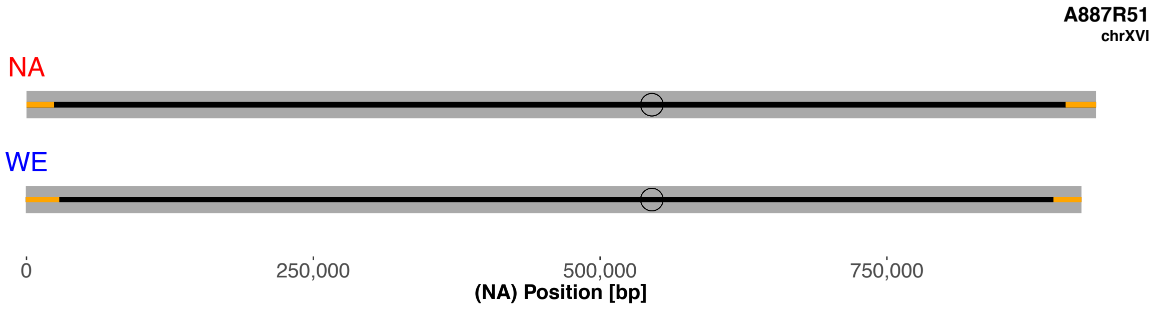
A887R51
chrXVI
NA
WE
0
250,000
500,000
750,000
(NA) Position [bp]

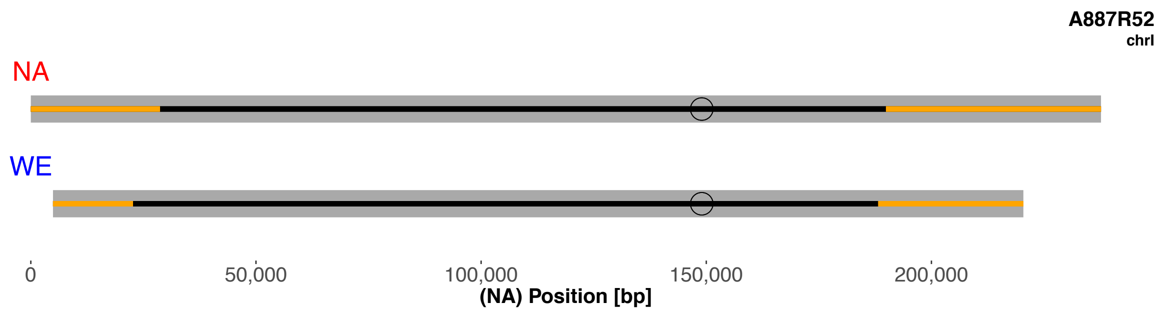
A887R52
chrI
NA
WE
0
50,000
100,000
150,000
200,000
(NA) Position [bp]

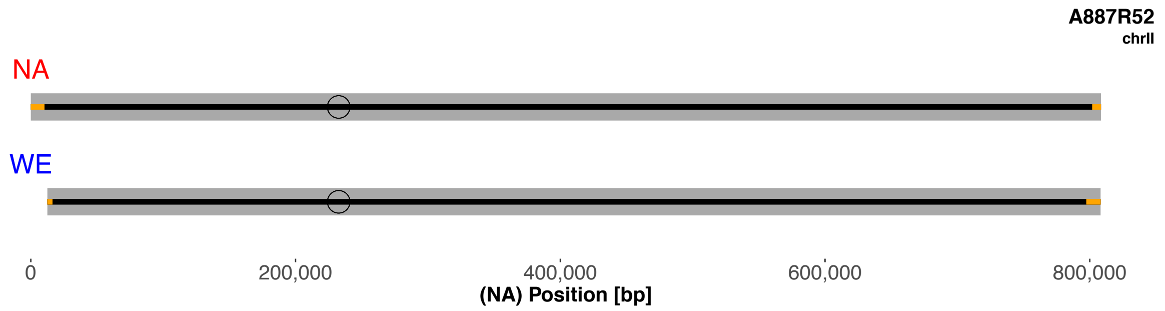
A887R52
chrII
NA
WE
0
200,000
400,000
600,000
800,000
(NA) Position [bp]

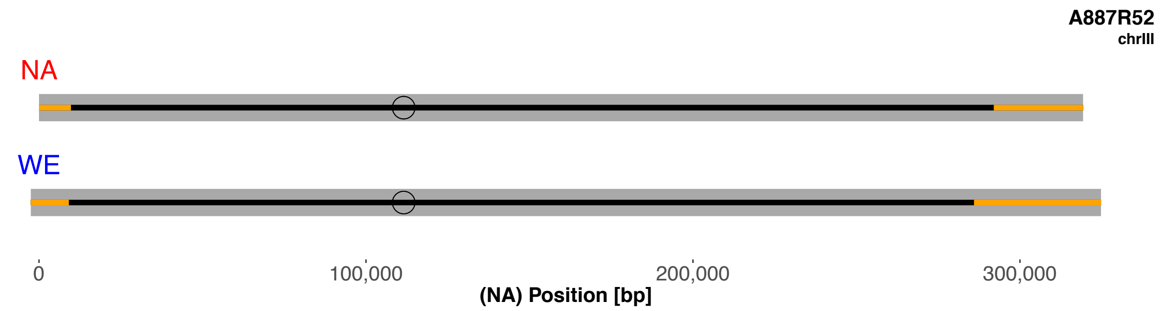
A887R52
chrIII
NA
WE
0
100,000
200,000
300,000
(NA) Position [bp]

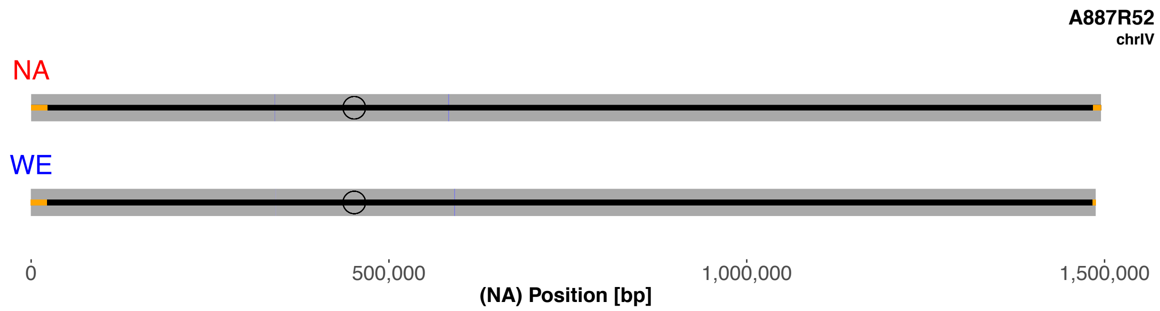
A887R52
chrIV
NA
WE
0
500,000
1,000,000
1,500,000
(NA) Position [bp]

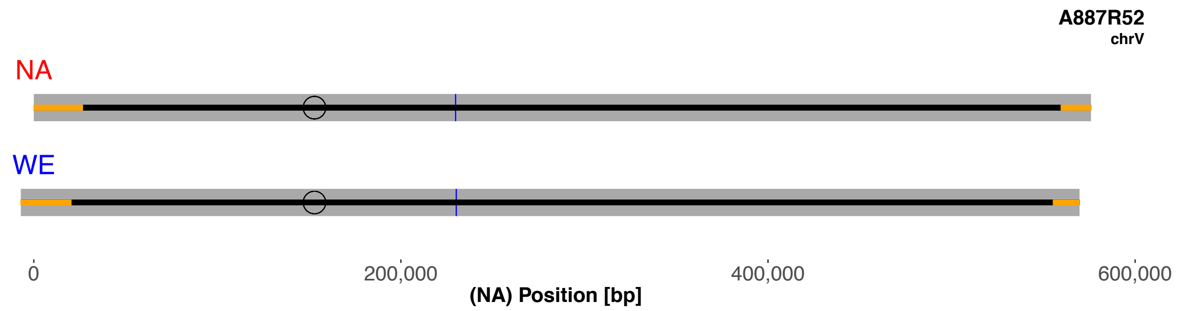
A887R52
chrV
NA
WE
0
200,000
400,000
600,000
(NA) Position [bp]

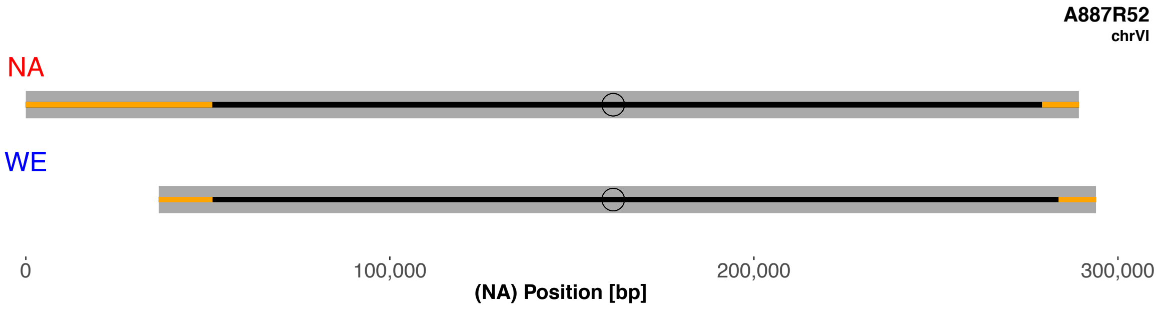
A887R52
chrVI
NA
WE
0
100,000
200,000
300,000
(NA) Position [bp]

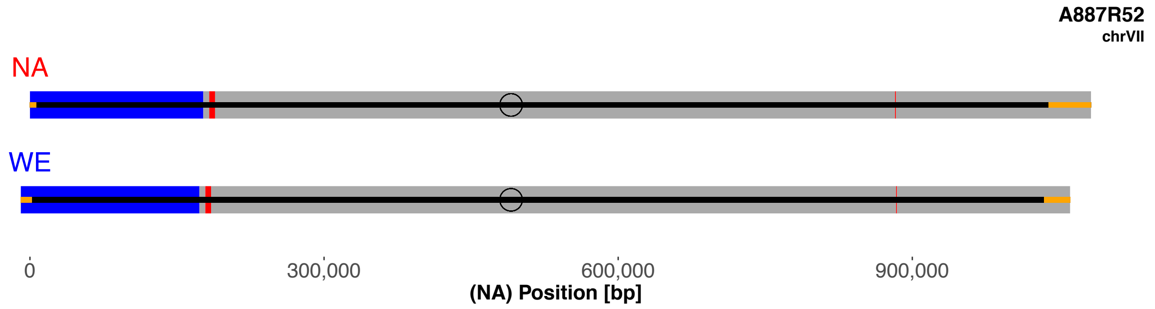
A887R52
chrVII
NA
WE
0
300,000
600,000
900,000
(NA) Position [bp]

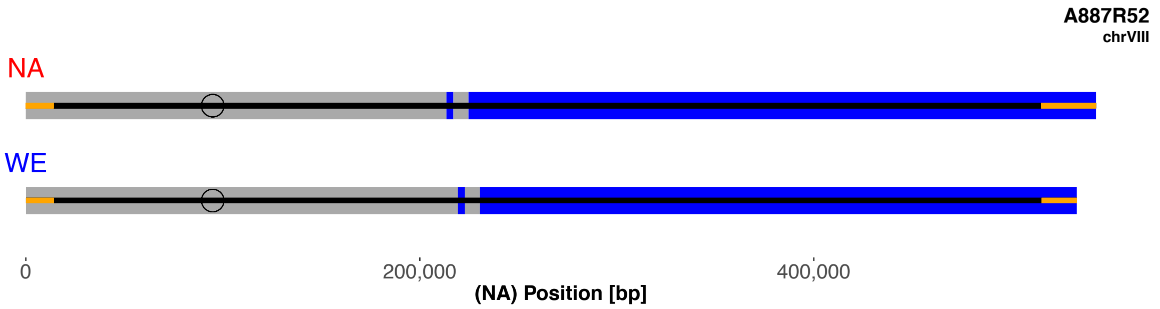
A887R52
chrVIII
NA
WE
0
200,000
400,000
(NA) Position [bp]

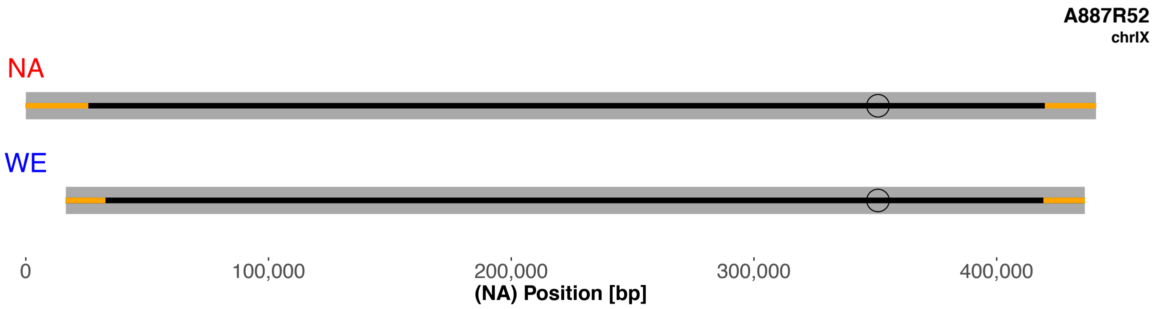
A887R52
chrIX
NA
WE
0
100,000
200,000
300,000
400,000
(NA) Position [bp]

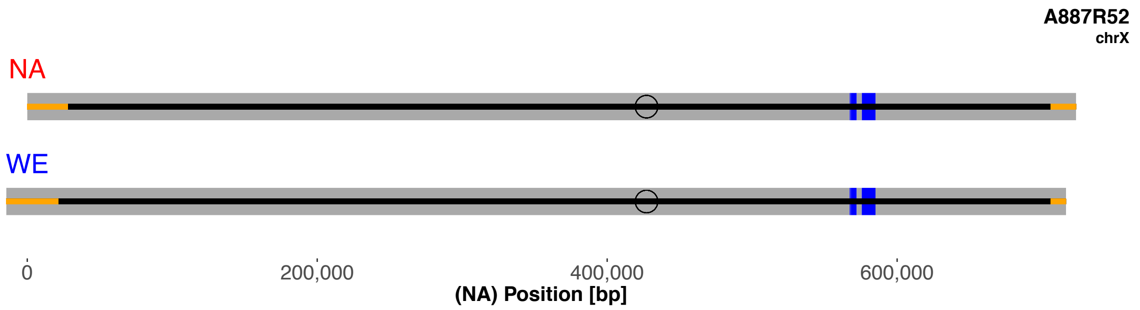
A887R52
chrX
NA
WE
0
200,000
400,000
600,000
(NA) Position [bp]

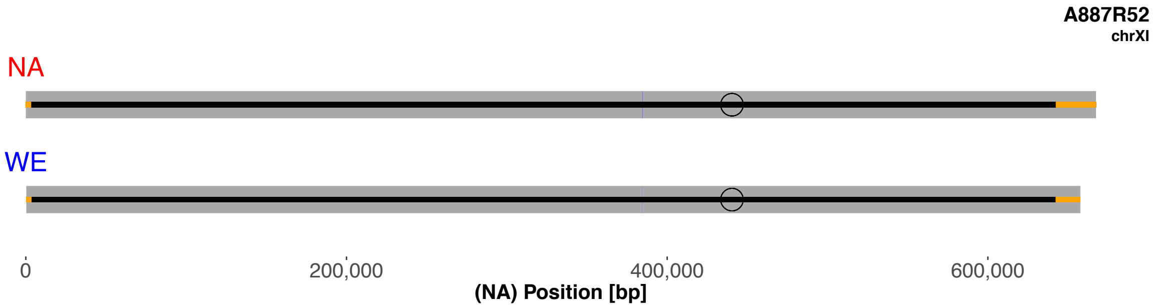
A887R52
chrXI
NA
WE
0
200,000
400,000
600,000
(NA) Position [bp]

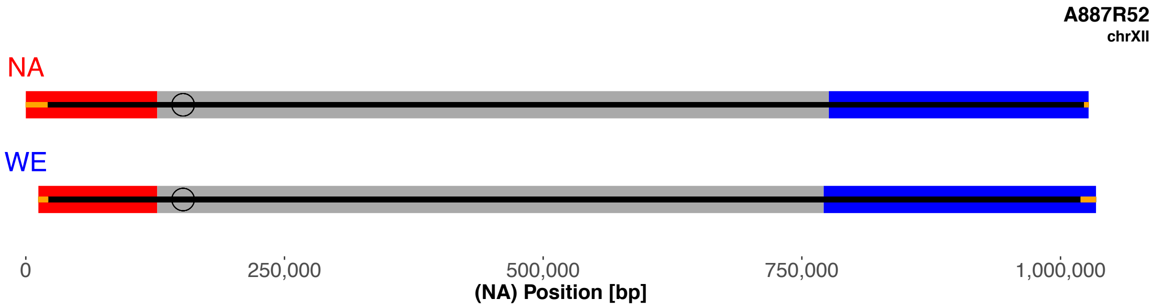
A887R52
chrXII
NA
WE
0
250,000
500,000
750,000
1,000,000
(NA) Position [bp]

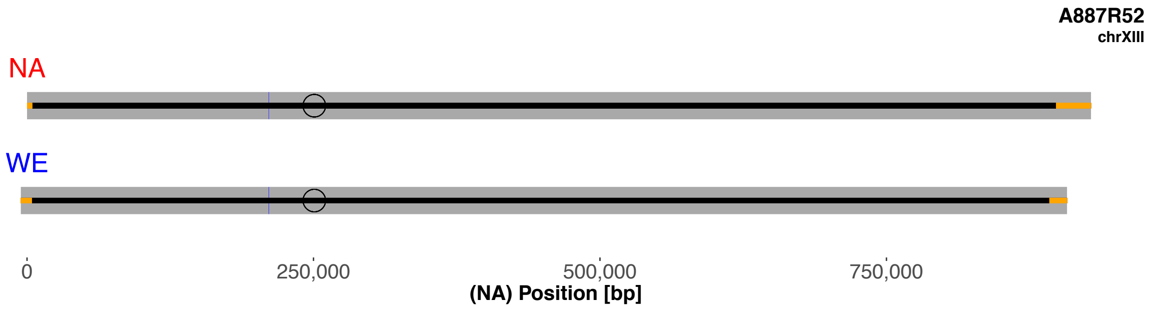
A887R52
chrXIII
NA
WE
0
250,000
500,000
750,000
(NA) Position [bp]

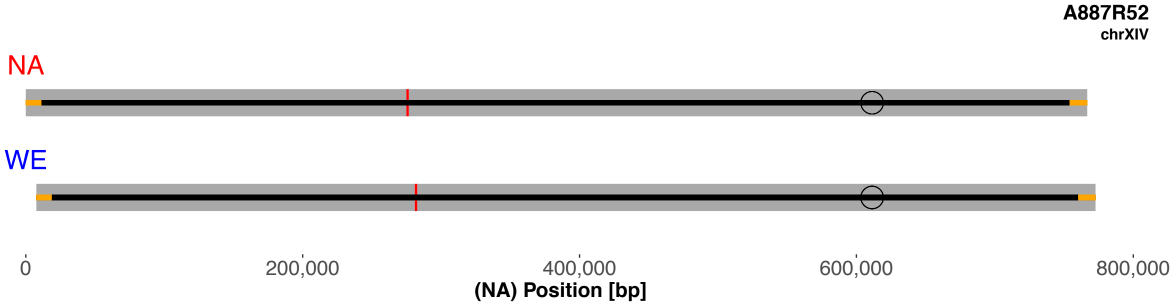
A887R52
chrXIV
NA
WE
0
200,000
400,000
600,000
800,000
(NA) Position [bp]

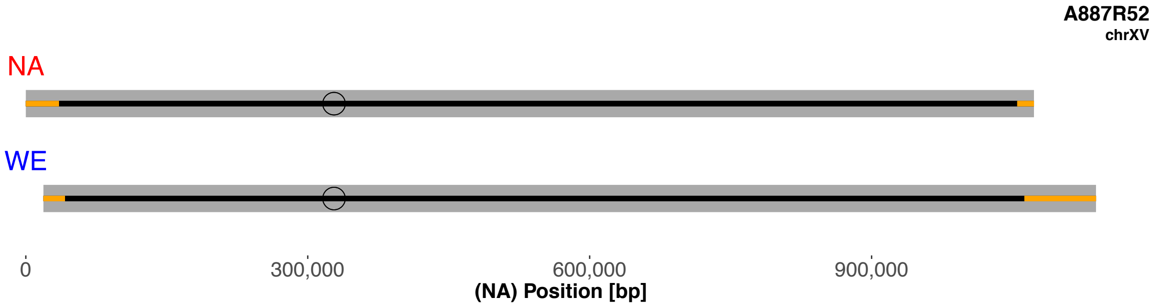
A887R52
chrXV
NA
WE
0
300,000
600,000
900,000
(NA) Position [bp]

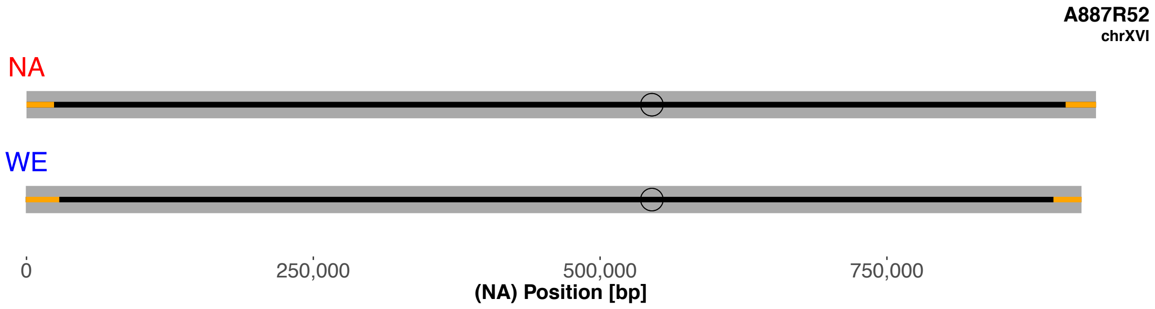
A887R52
chrXVI
NA
WE
0
250,000
500,000
750,000
(NA) Position [bp]

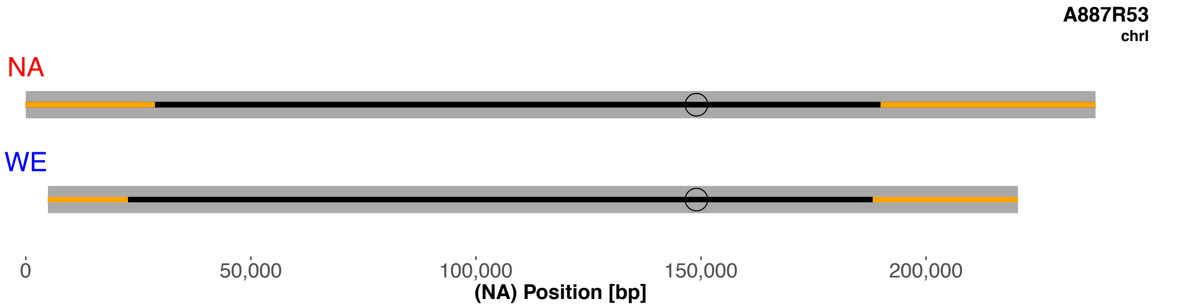
A887R53
chrI
NA
WE
0
50,000
100,000
150,000
200,000
(NA) Position [bp]

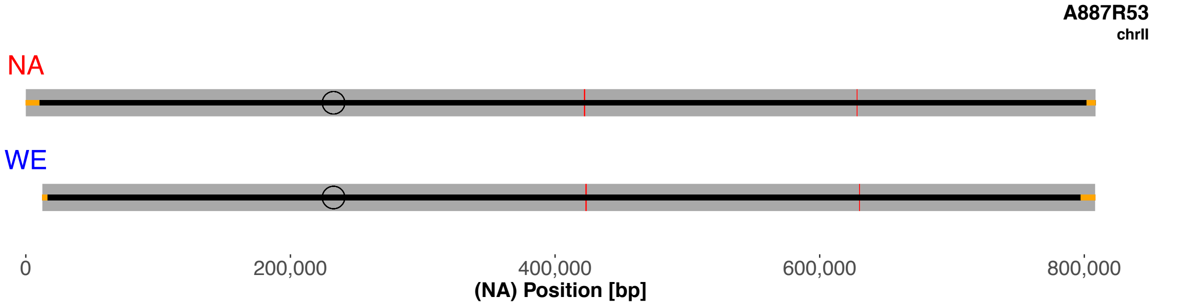
A887R53
chrII
NA
WE
0
200,000
400,000
600,000
800,000
(NA) Position [bp]

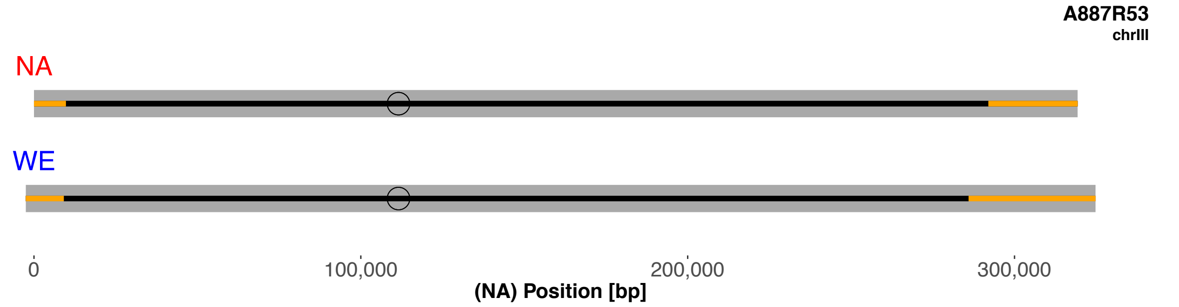
A887R53
chrIII
NA
WE
0
100,000
200,000
300,000
(NA) Position [bp]

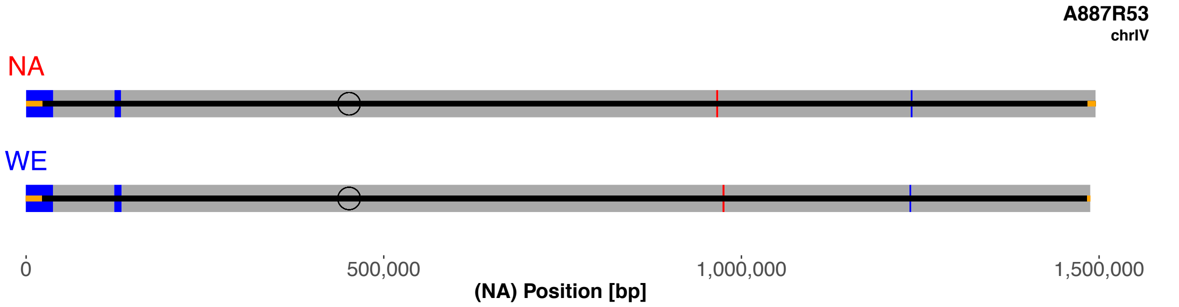
A887R53
chrIV
NA
WE
0
500,000
1,000,000
1,500,000
(NA) Position [bp]

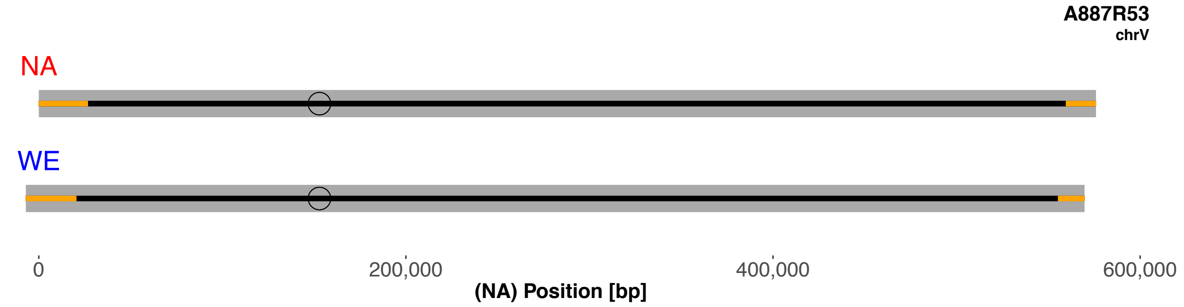
A887R53
chrV
NA
WE
0
200,000
400,000
600,000
(NA) Position [bp]

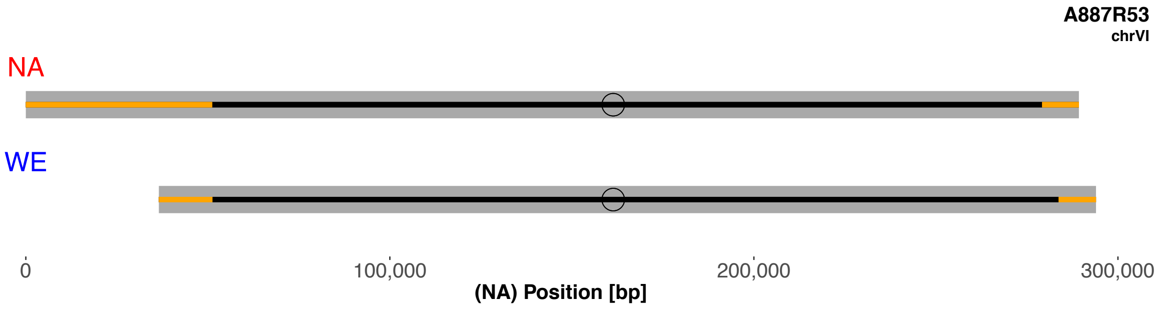
A887R53
chrVI
NA
WE
0
100,000
200,000
300,000
(NA) Position [bp]

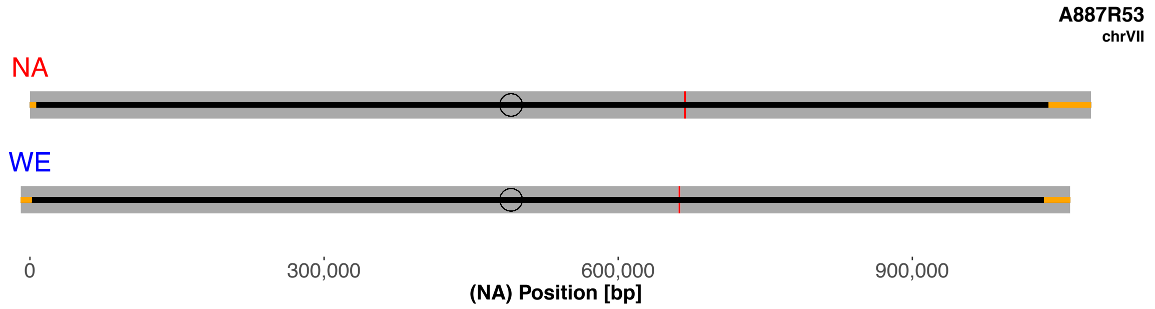
A887R53
chrVII
NA
WE
0
300,000
600,000
900,000
(NA) Position [bp]

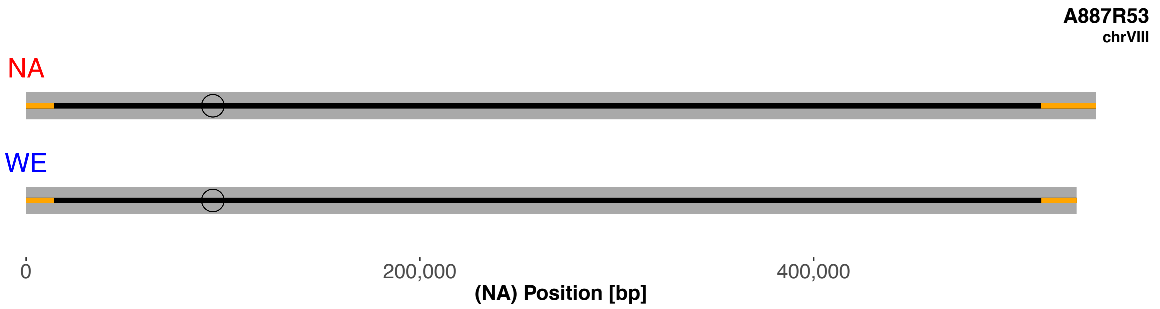
A887R53
chrVIII
NA
WE
0
200,000
400,000
(NA) Position [bp]

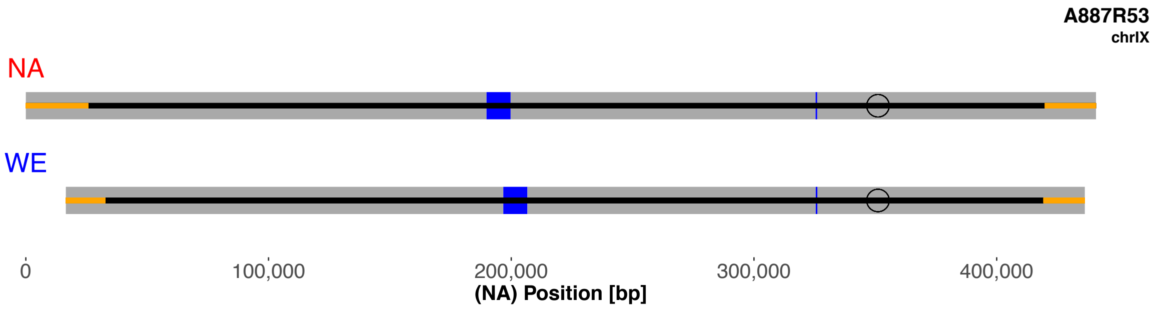
A887R53
chrIX
NA
WE
0
100,000
200,000
300,000
400,000
(NA) Position [bp]

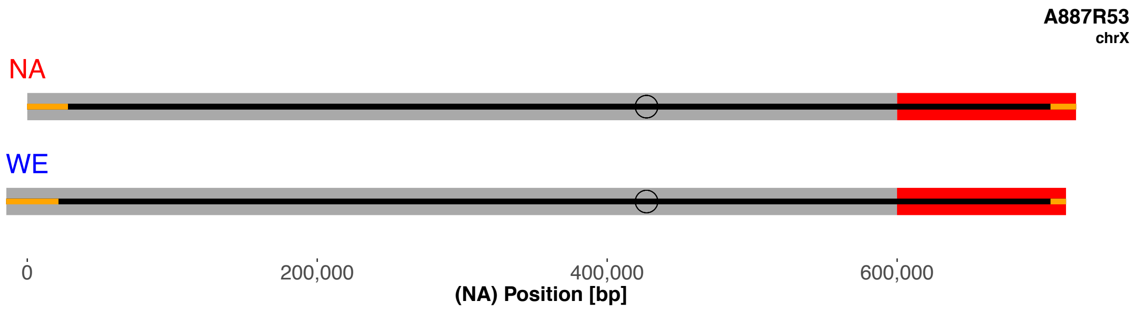
A887R53
chrX
NA
WE
0
200,000
400,000
600,000
(NA) Position [bp]

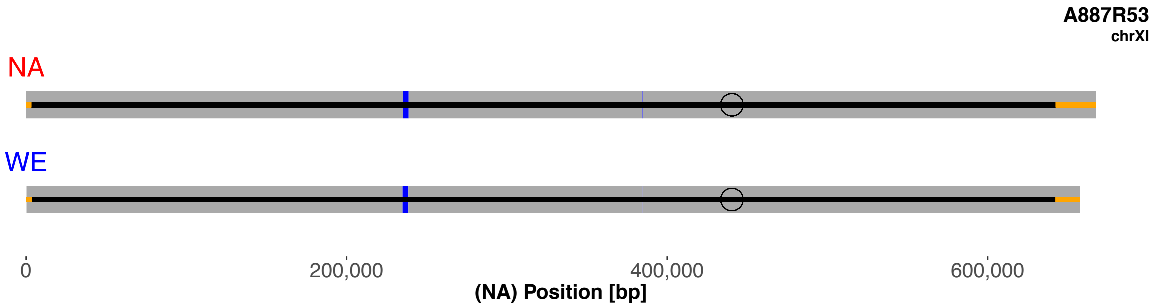
A887R53
chrXI
NA
WE
0
200,000
400,000
600,000
(NA) Position [bp]

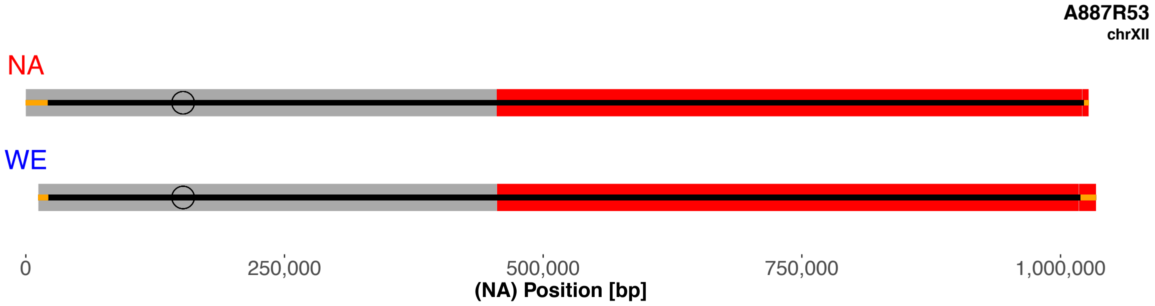
A887R53
chrXII
NA
WE
0
250,000
500,000
750,000
1,000,000
(NA) Position [bp]

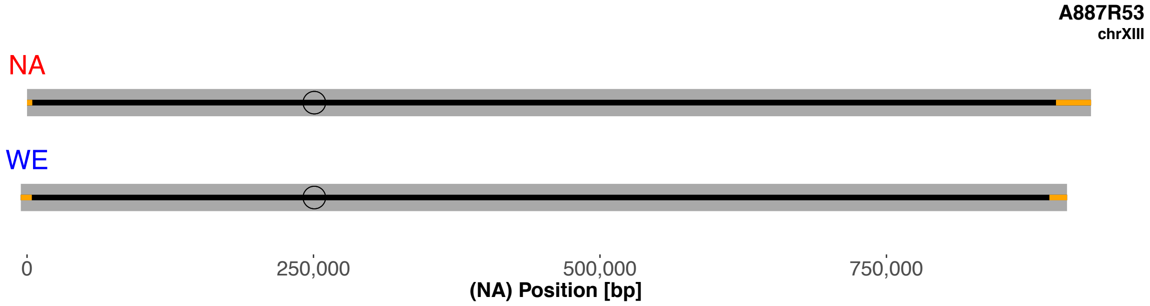
A887R53
chrXIII
NA
WE
0
250,000
500,000
750,000
(NA) Position [bp]

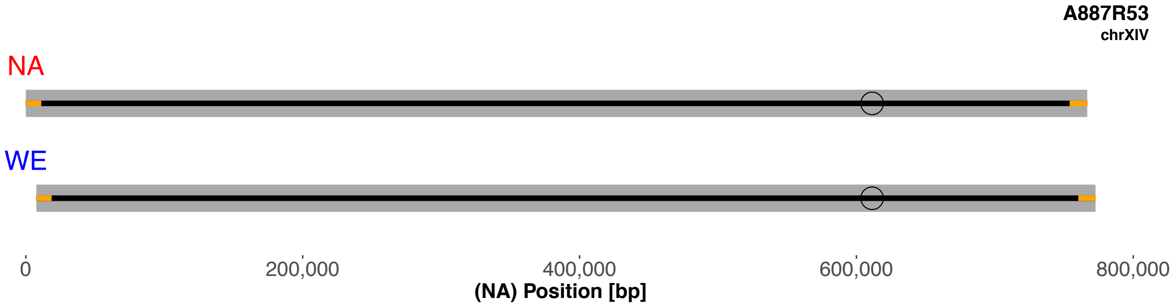
A887R53
chrXIV
NA
WE
0
200,000
400,000
600,000
800,000
(NA) Position [bp]

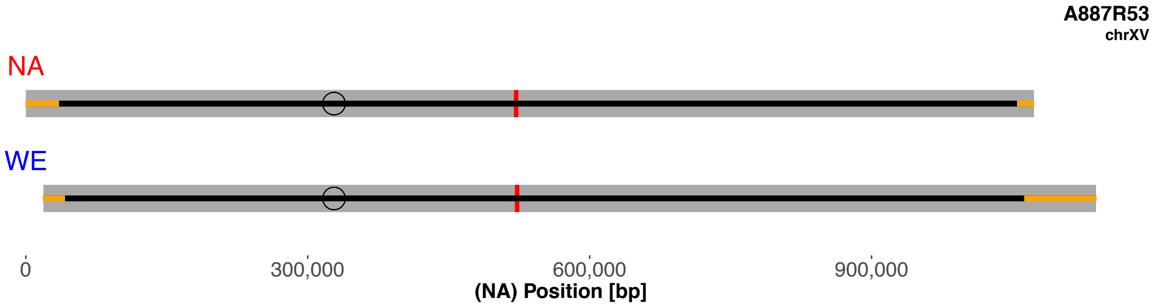
A887R53
chrXV
NA
WE
0
300,000
600,000
900,000
(NA) Position [bp]

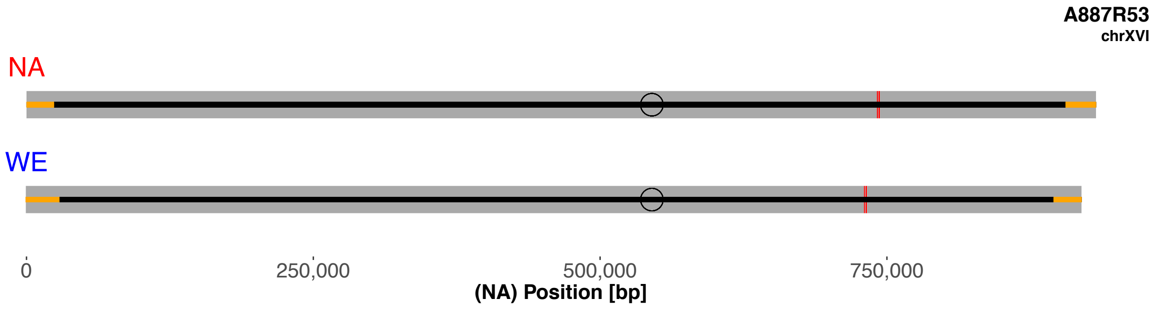

A887R53
chrXVI
NA
WE
0
250,000
500,000
750,000
(NA) Position [bp]

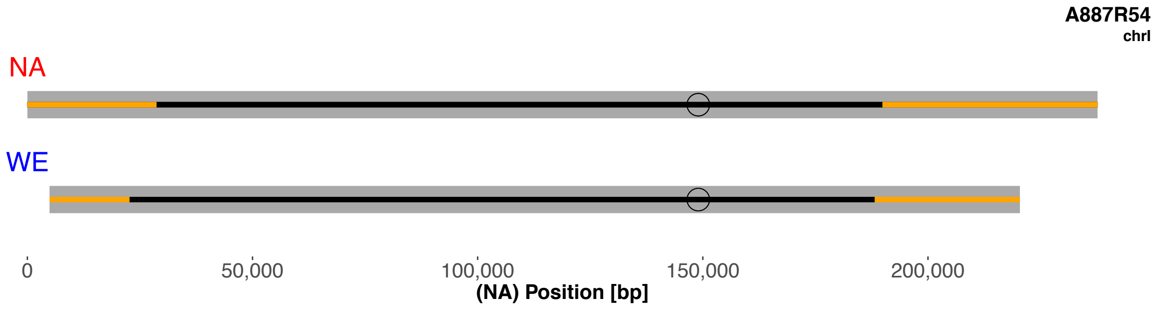
A887R54
chrI
NA
WE
0
50,000
100,000
150,000
200,000
(NA) Position [bp]

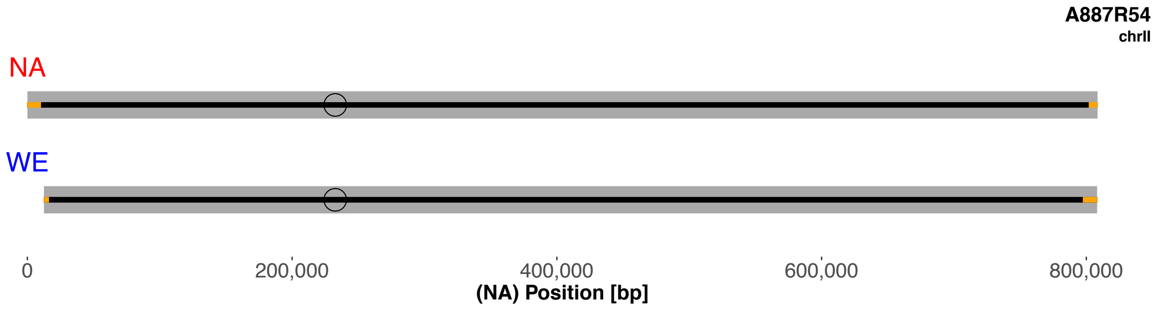
A887R54
chrII
NA
WE
0
200,000
400,000
600,000
800,000
(NA) Position [bp]

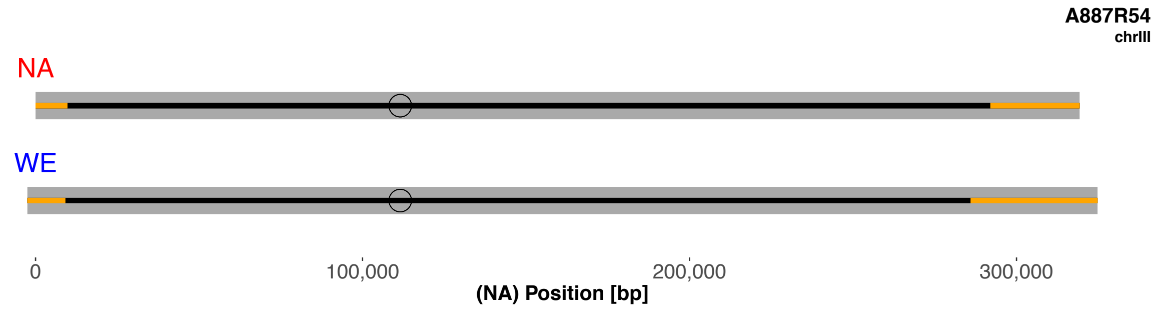
A887R54
chrIII
NA
WE
0
100,000
200,000
300,000
(NA) Position [bp]

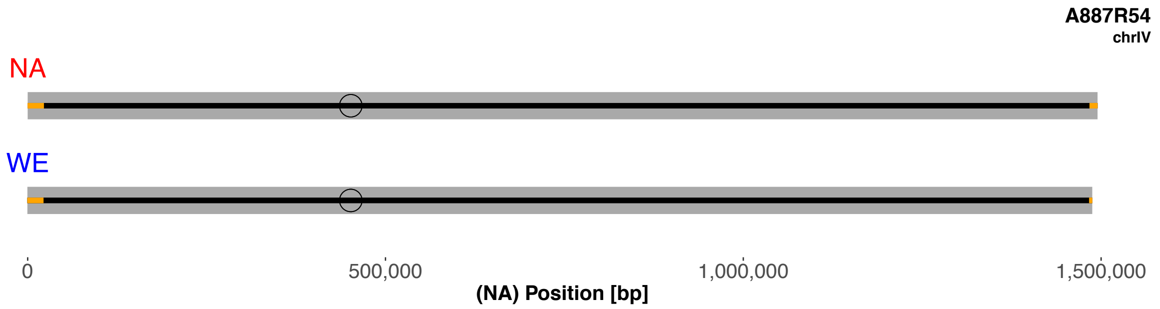
A887R54
chrIV
NA
WE
0
500,000
1,000,000
1,500,000
(NA) Position [bp]

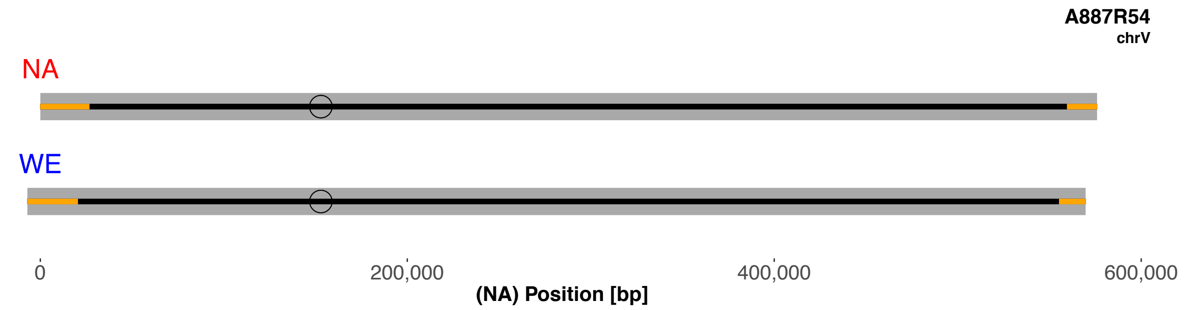
A887R54
chrV
NA
WE
0
200,000
400,000
600,000
(NA) Position [bp]

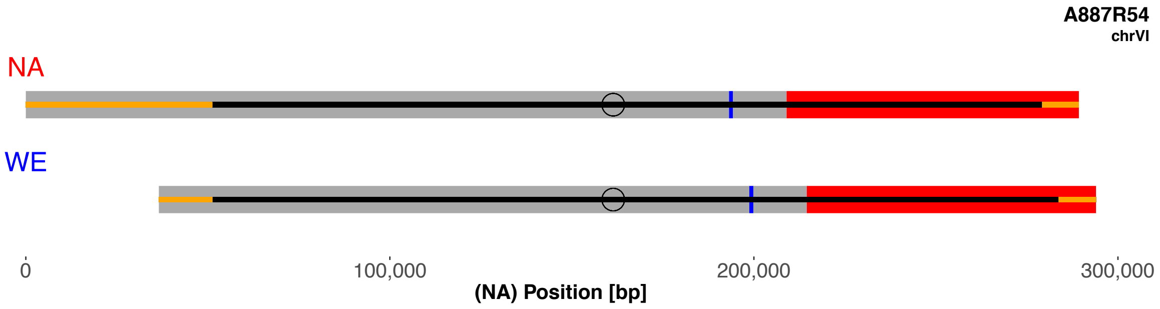
A887R54
chrVI
NA
WE
0
100,000
200,000
300,000
(NA) Position [bp]

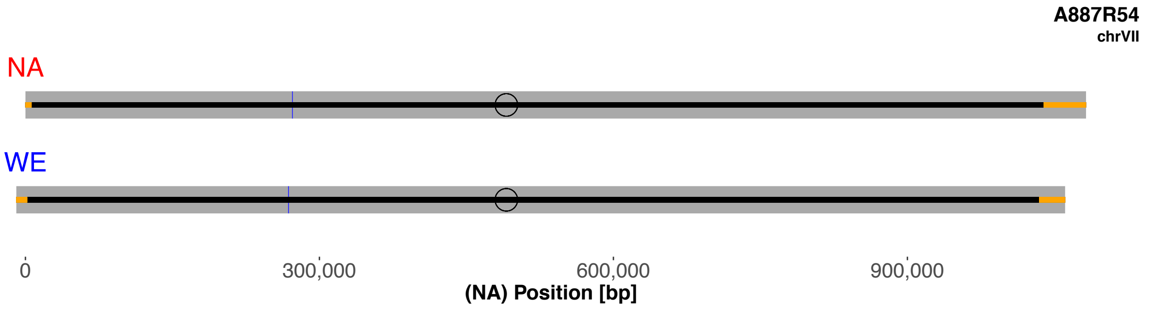
A887R54
chrVII
NA
WE
0
300,000
600,000
900,000
(NA) Position [bp]

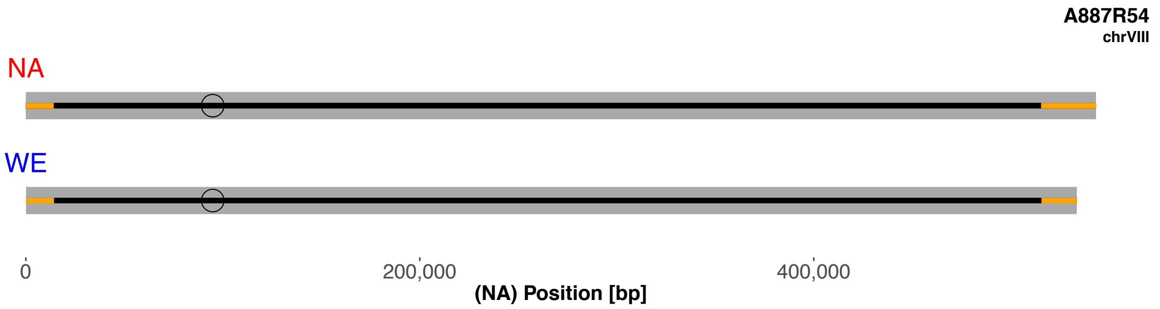
A887R54
chrVIII
NA
WE
0
200,000
400,000
(NA) Position [bp]

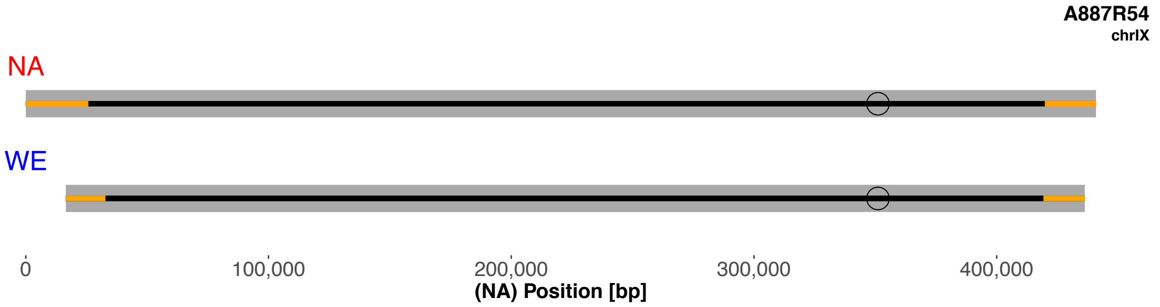
A887R54
chrIX
NA
WE
0
100,000
200,000
300,000
400,000
(NA) Position [bp]

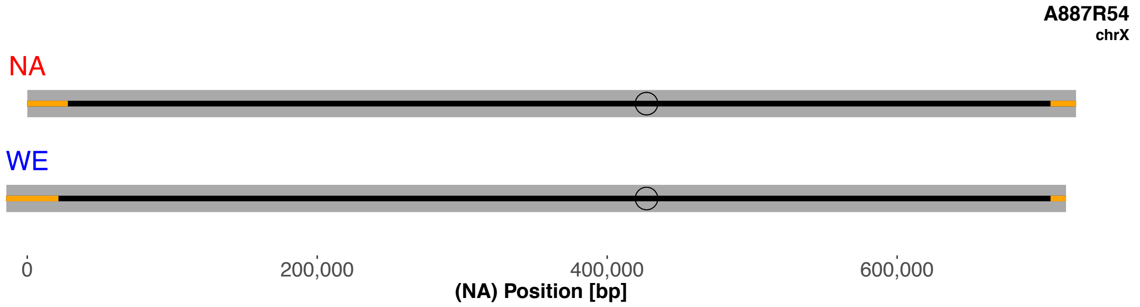
A887R54
chrX
NA
WE
0
200,000
400,000
600,000
(NA) Position [bp]

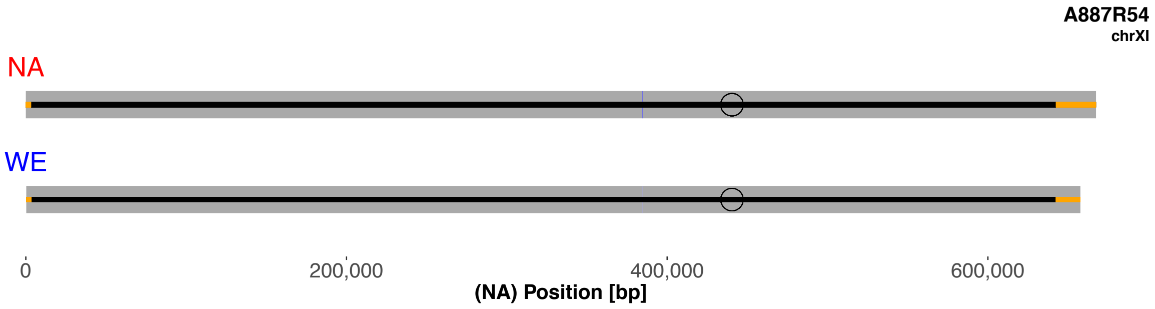
A887R54
chrXI
NA
WE
0
200,000
400,000
600,000
(NA) Position [bp]

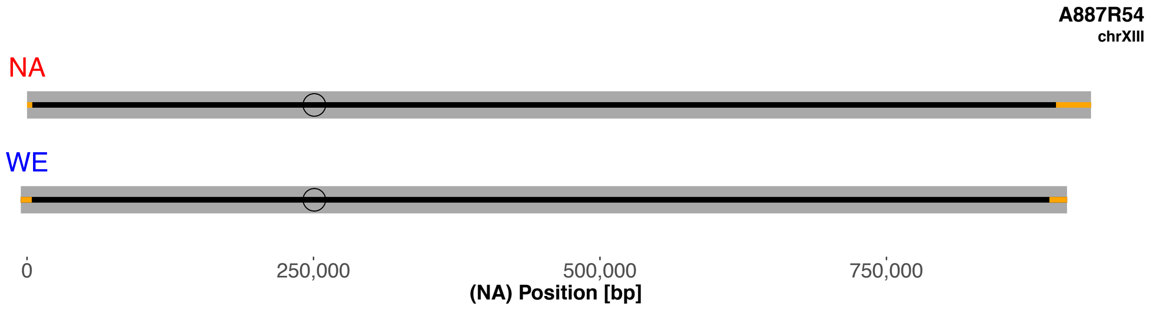
A887R54
chrXIII
NA
WE
0
250,000
500,000
750,000
(NA) Position [bp]

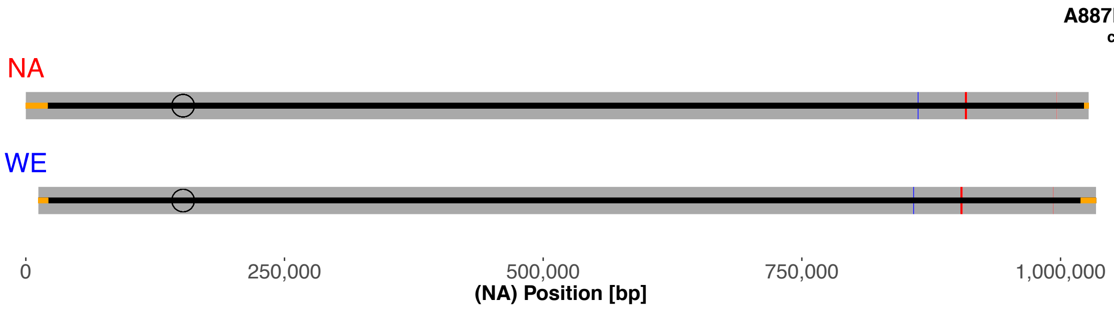
A887R54
chrXII
NA
WE
0
250,000
500,000
750,000
1,000,000
(NA) Position [bp]

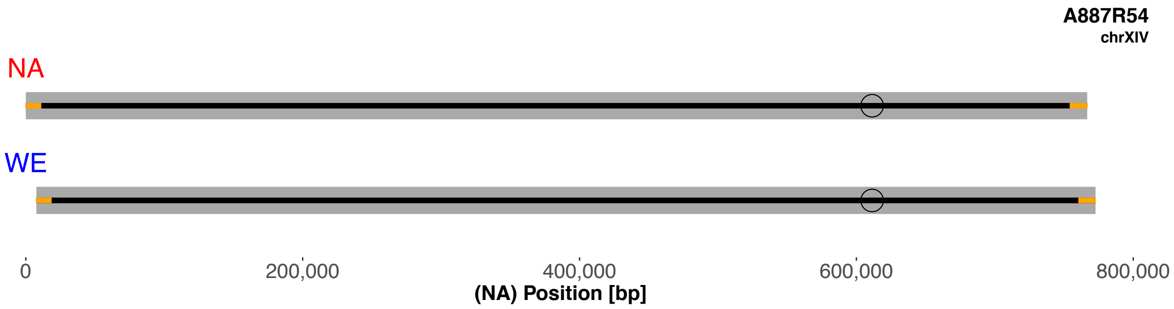
A887R54
chrXIV
NA
WE
0
200,000
400,000
600,000
800,000
(NA) Position [bp]

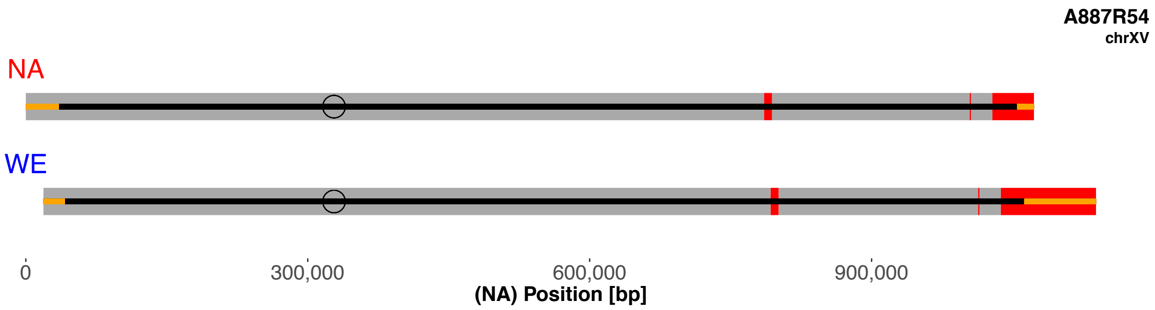
A887R54
chrXV
NA
WE
0
300,000
600,000
900,000
(NA) Position [bp]

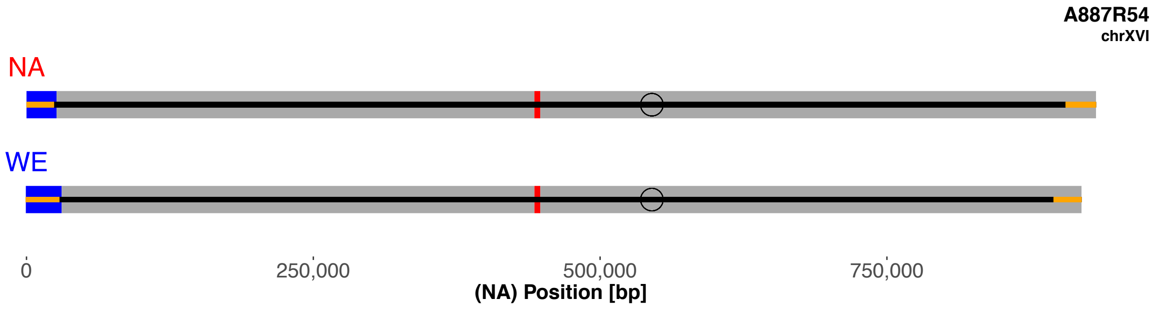

A887R54
chrXVI
NA
WE
0
250,000
500,000
750,000
(NA) Position [bp]

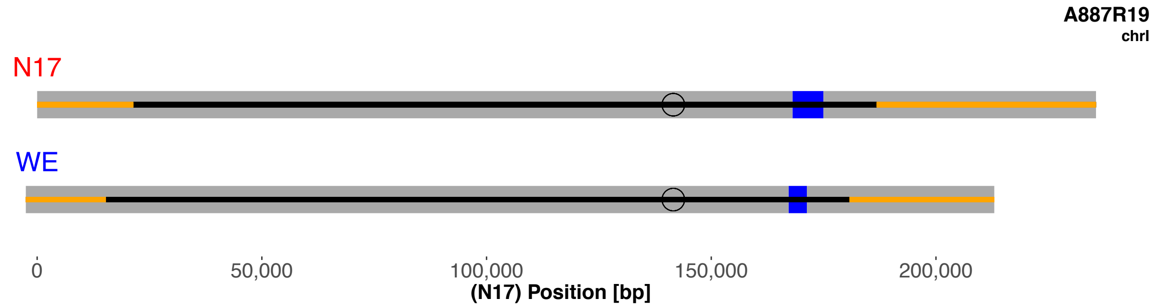
A887R19
chrI
N17
WE
0
50,000
100,000
150,000
200,000
(N17) Position [bp]

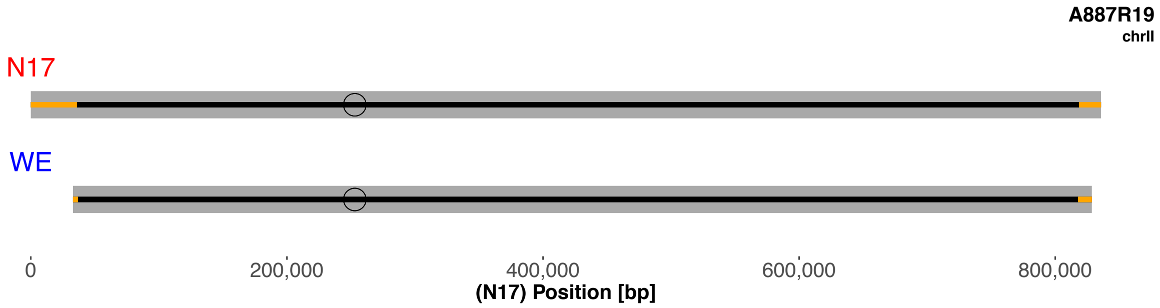
A887R19
chrII
N17
WE
0
200,000
400,000
600,000
800,000
(N17) Position [bp]

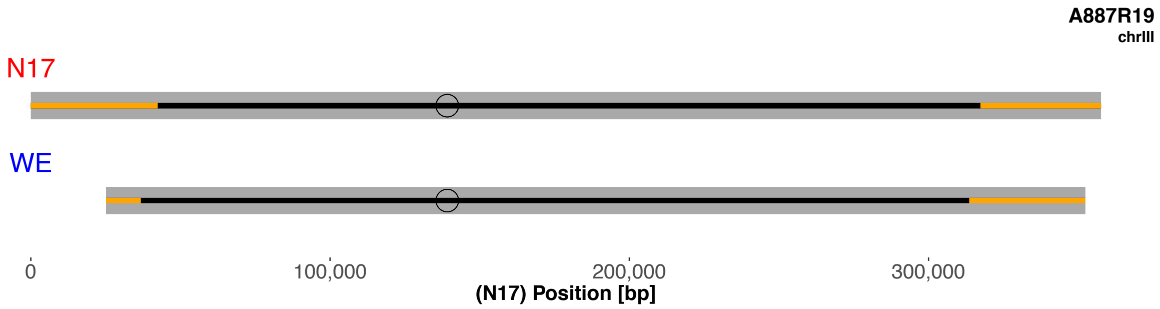
A887R19
chrIII
N17
WE
0
100,000
200,000
300,000
(N17) Position [bp]

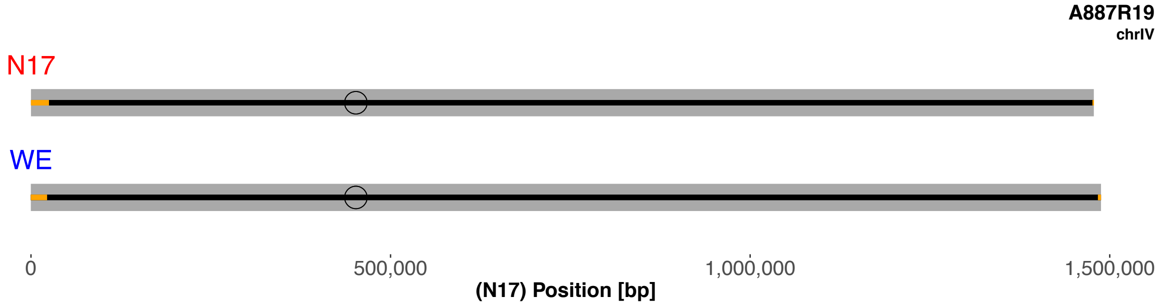
A887R19
chrIV
N17
WE
0
500,000
1,000,000
1,500,000
(N17) Position [bp]

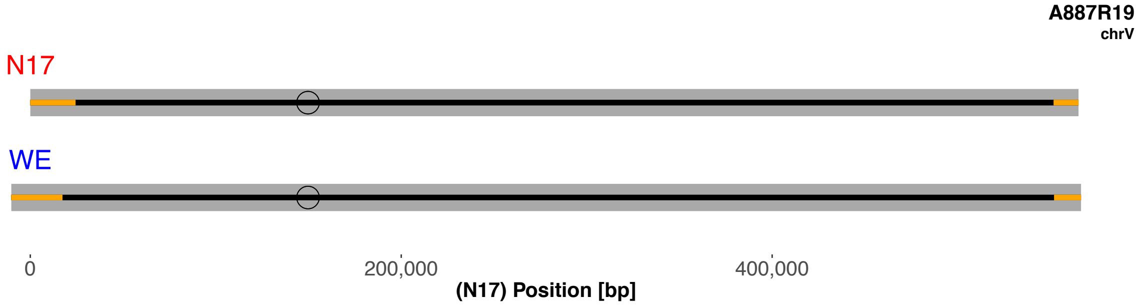
A887R19
chrV
N17
WE
0
200,000
400,000
(N17) Position [bp]

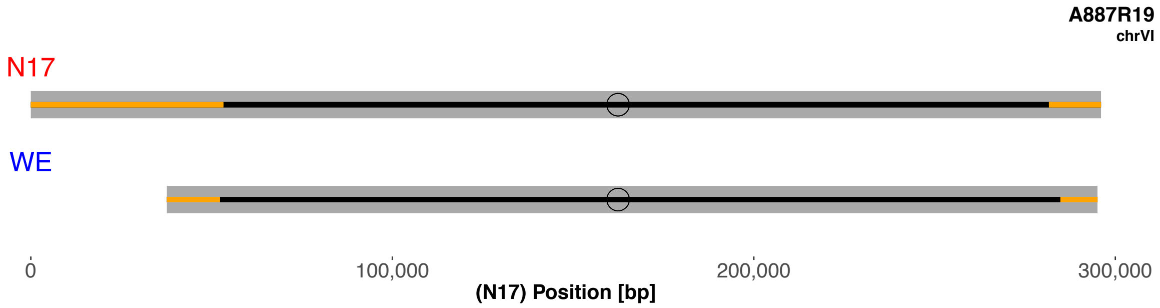
A887R19
chrVI
N17
WE
0
100,000
200,000
300,000
(N17) Position [bp]

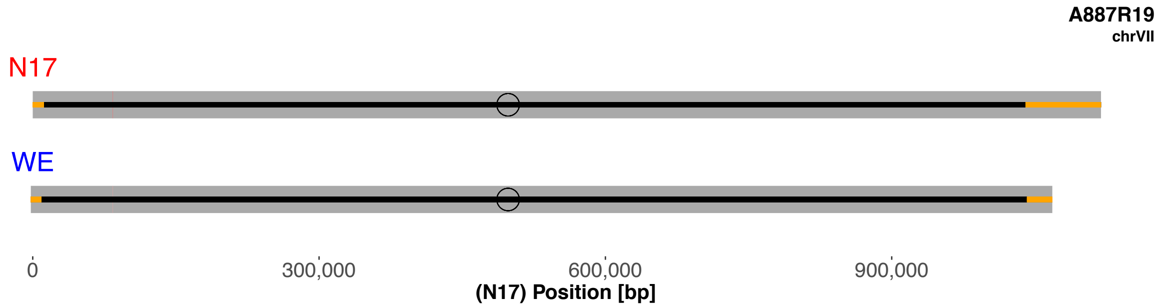
A887R19
chrVII
N17
WE
0
300,000
600,000
900,000
(N17) Position [bp]

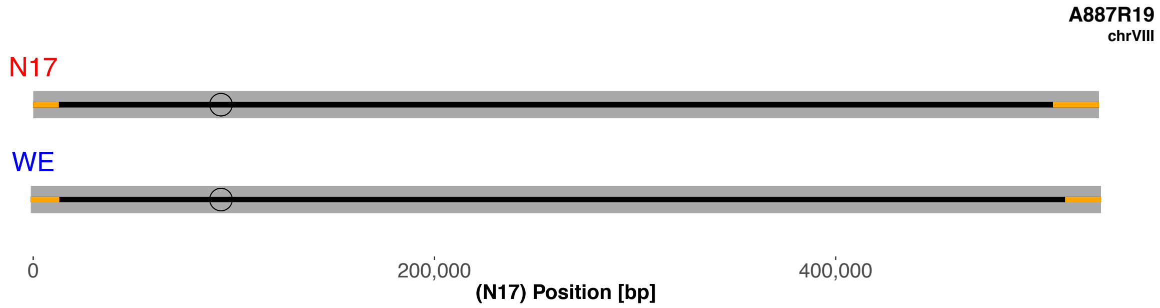
A887R19
chrVIII
N17
WE
0
200,000
400,000
(N17) Position [bp]

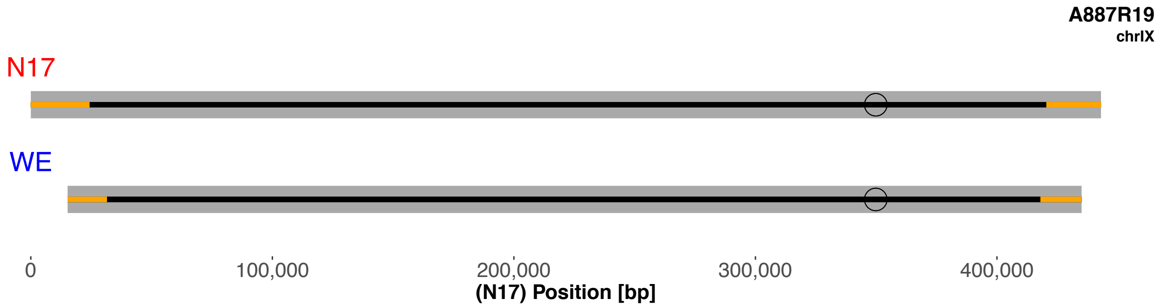
A887R19
chrIX
N17
WE
0
100,000
200,000
300,000
400,000
(N17) Position [bp]

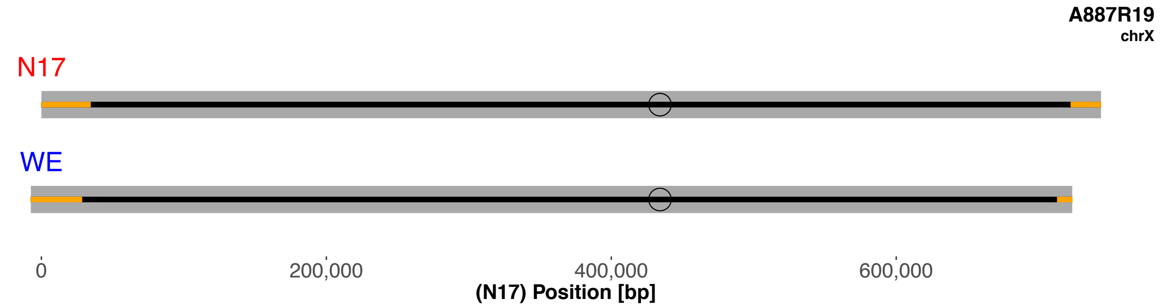
A887R19
chrX
N17
WE
0
200,000
400,000
600,000
(N17) Position [bp]

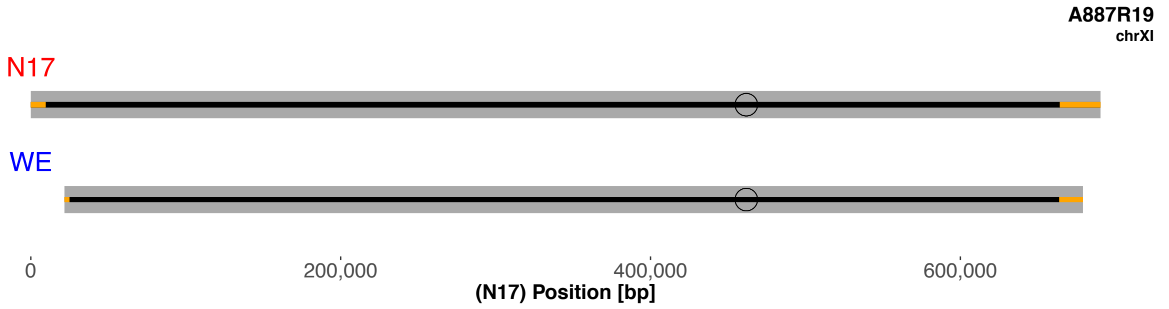
A887R19
chrXI
N17
WE
0
200,000
400,000
600,000
(N17) Position [bp]

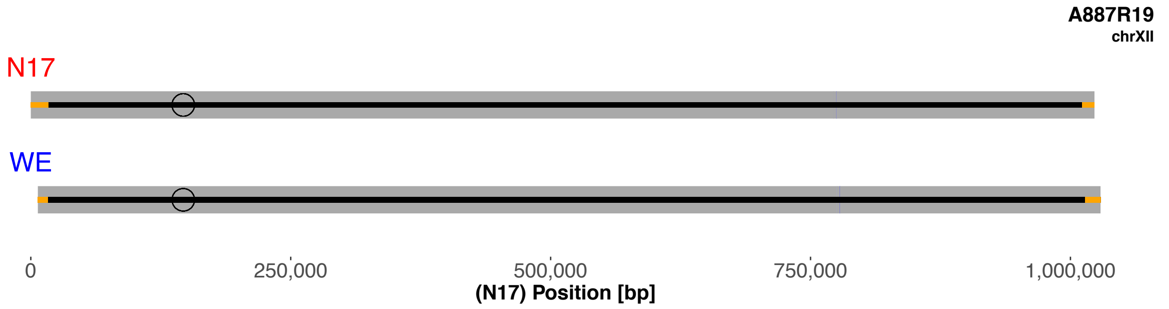
A887R19
chrXII
N17
WE
0
250,000
500,000
750,000
1,000,000
(N17) Position [bp]

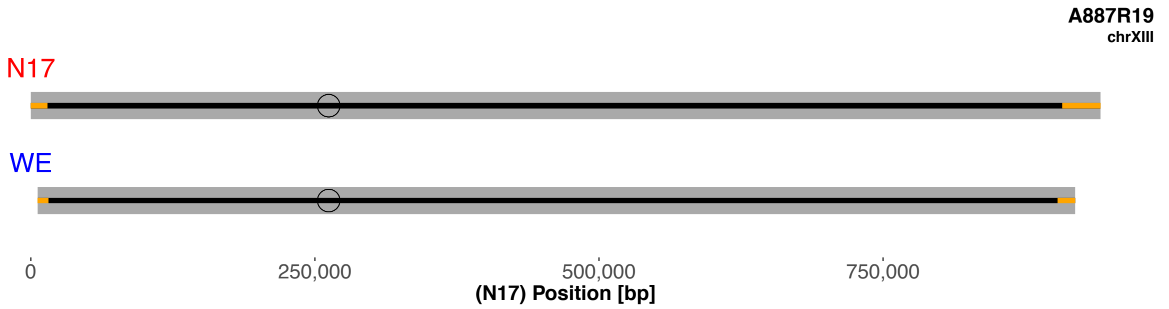
A887R19
chrXIII
N17
WE
0
250,000
500,000
750,000
(N17) Position [bp]

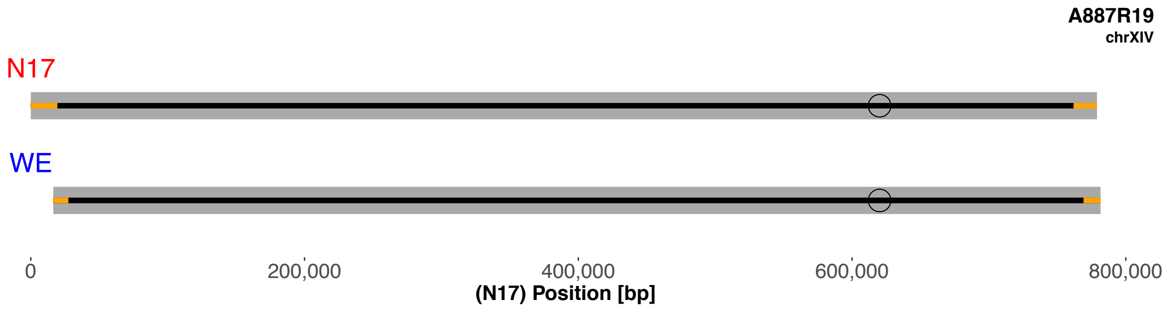
A887R19
chrXIV
N17
WE
0
200,000
400,000
600,000
800,000
(N17) Position [bp]

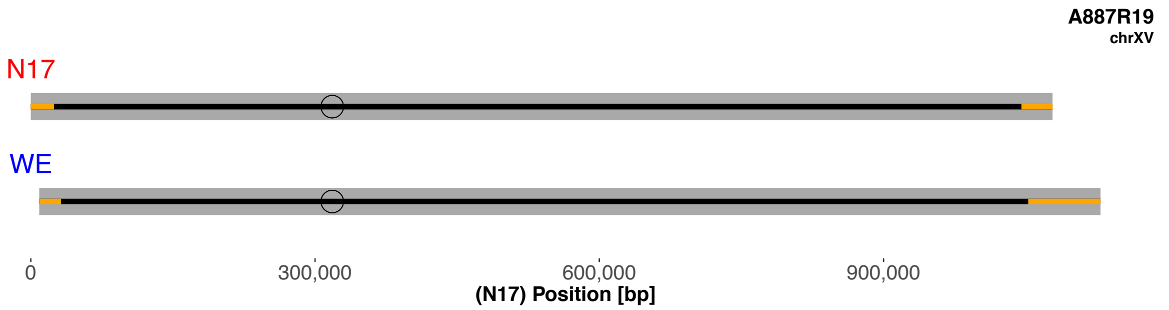
A887R19
chrXV
N17
WE
0
300,000
600,000
900,000
(N17) Position [bp]

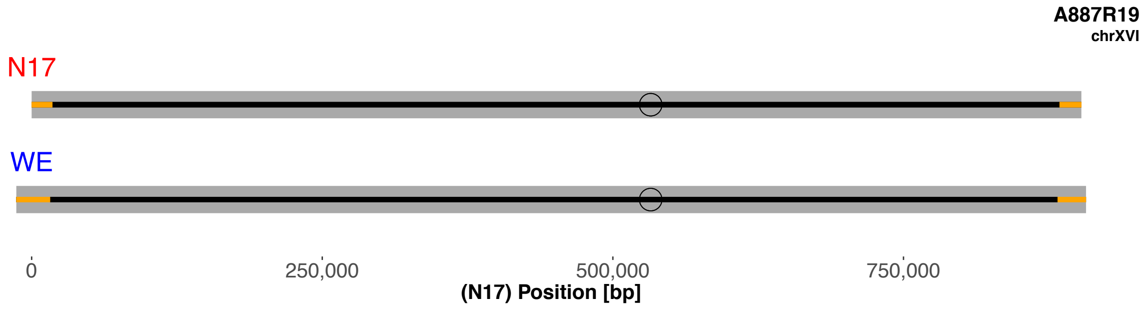
A887R19
chrXVI
N17
WE
0
250,000
500,000
750,000
(N17) Position [bp]

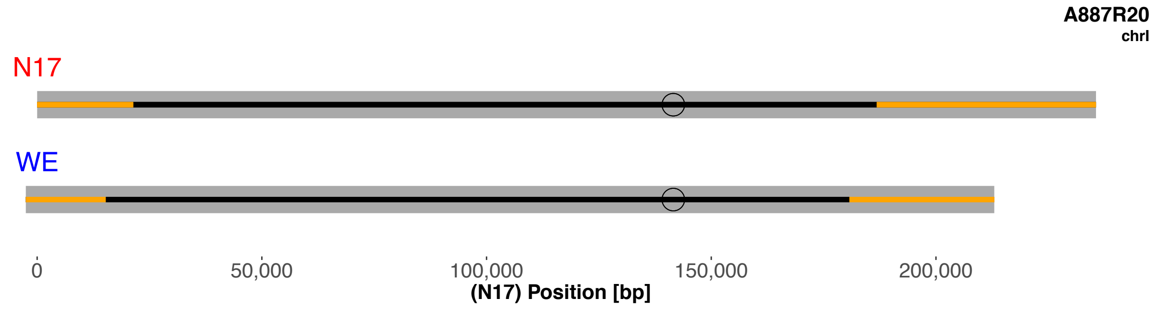

A887R20
chrI
N17
WE
0
50,000
100,000
150,000
200,000
(N17) Position [bp]


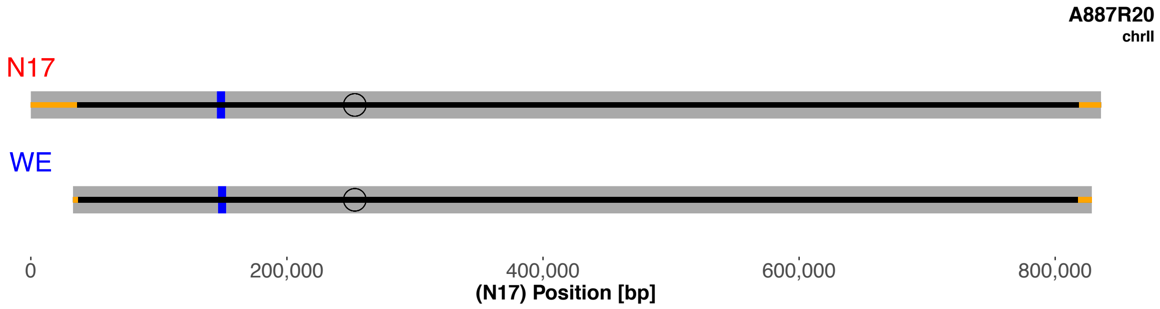

A887R20
chrII
N17
WE
0
200,000
400,000
600,000
800,000
(N17) Position [bp]


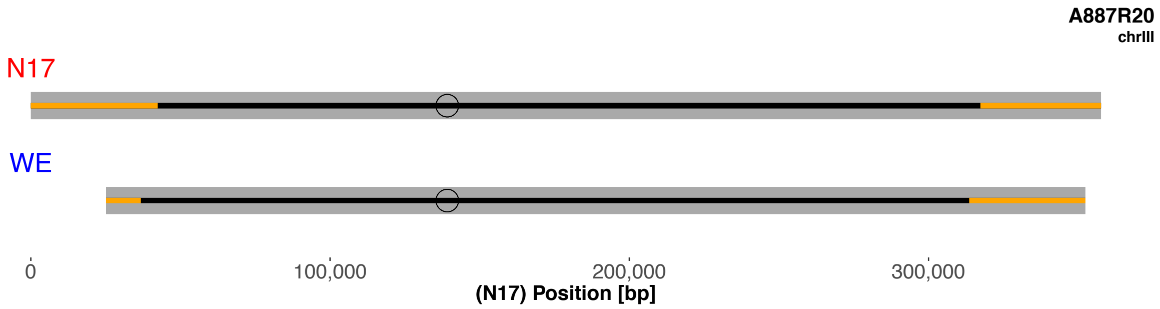

A887R20
chrIII
N17
WE
0
100,000
200,000
300,000
(N17) Position [bp]


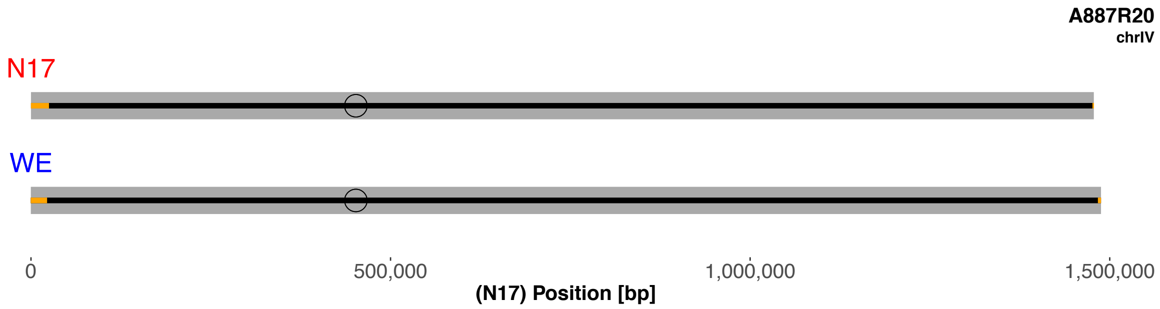

A887R20
chrIV
N17
WE
0
500,000
1,000,000
1,500,000
(N17) Position [bp]


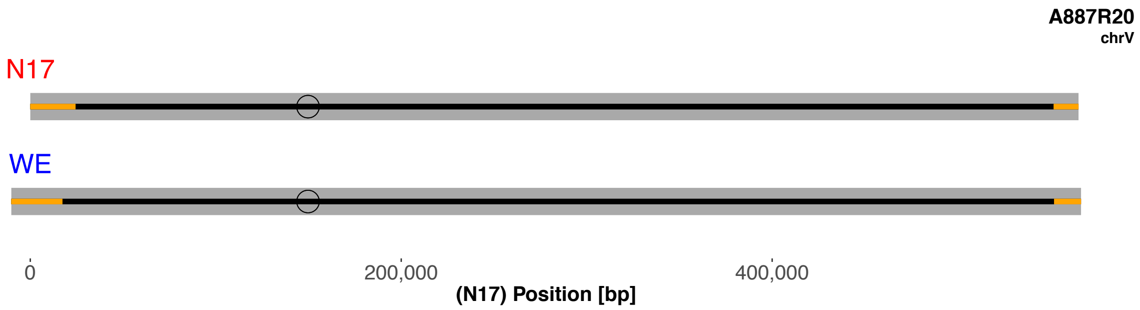

A887R20
chrV
N17
WE
0
200,000
400,000
(N17) Position [bp]

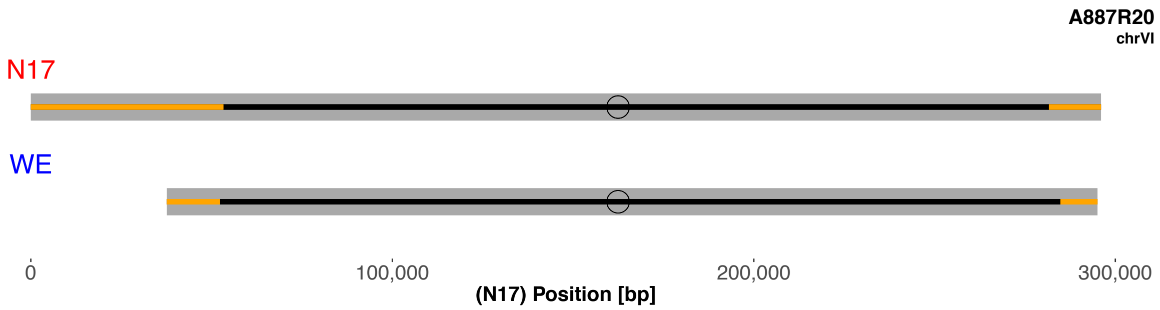
A887R20
chrVI
N17
WE
0
100,000
200,000
300,000
(N17) Position [bp]

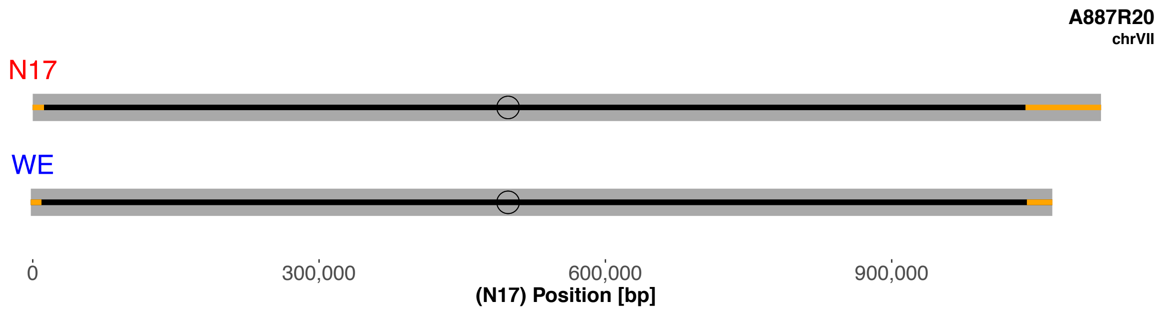
A887R20
chrVII
N17
WE
0
300,000
600,000
900,000
(N17) Position [bp]

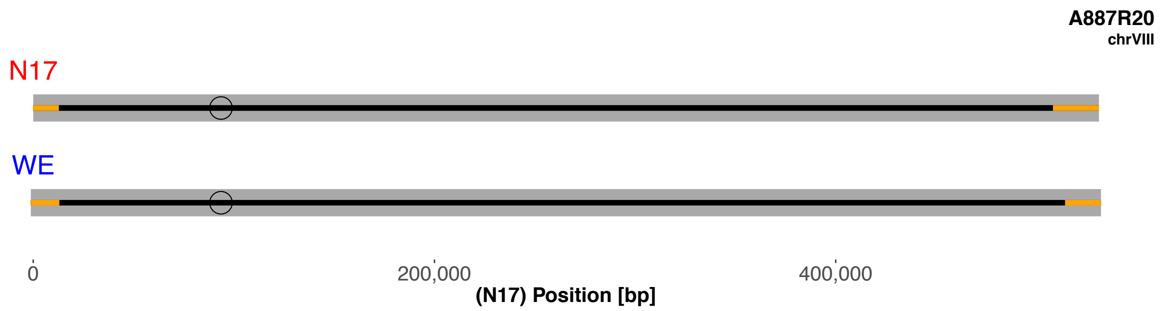
A887R20
chrVIII
N17
WE
0
200,000
400,000
(N17) Position [bp]

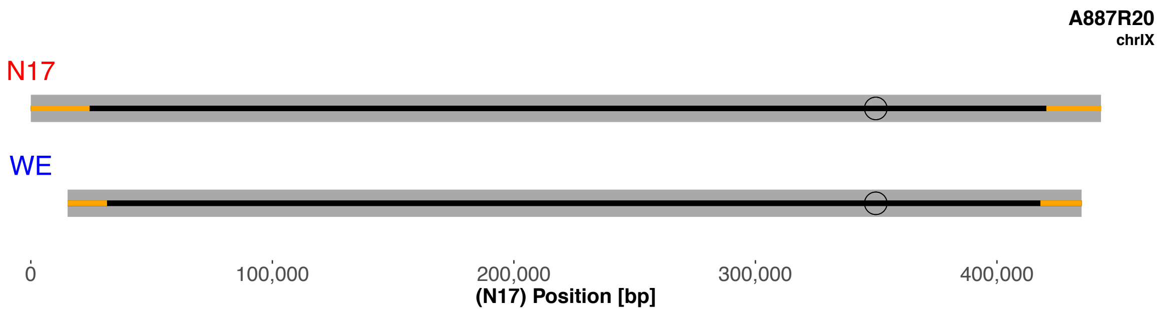
A887R20
chrIX
N17
WE
0
100,000
200,000
300,000
400,000
(N17) Position [bp]

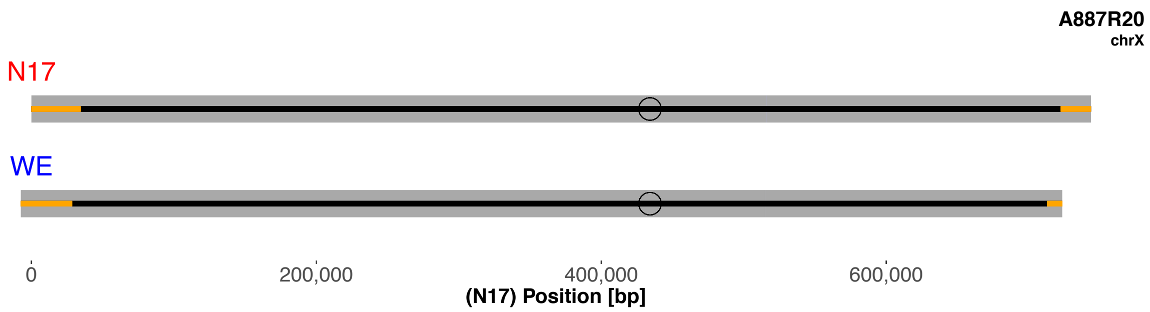
A887R20
chrX
N17
WE
0
200,000
400,000
600,000
(N17) Position [bp]

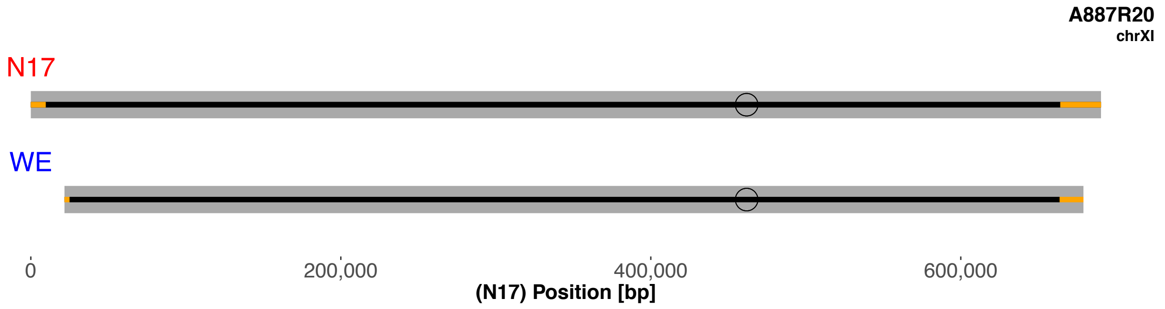
A887R20
chrXI
N17
WE
0
200,000
400,000
600,000
(N17) Position [bp]

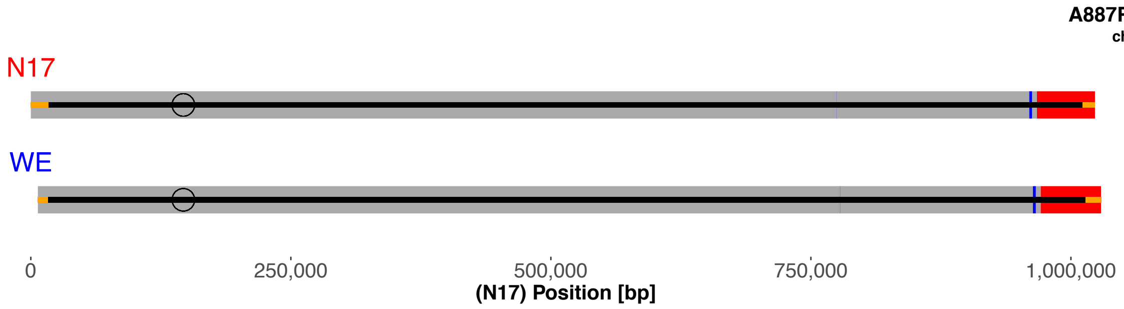
A887R20
chrXII
N17
WE
0
250,000
500,000
750,000
1,000,000
(N17) Position [bp]

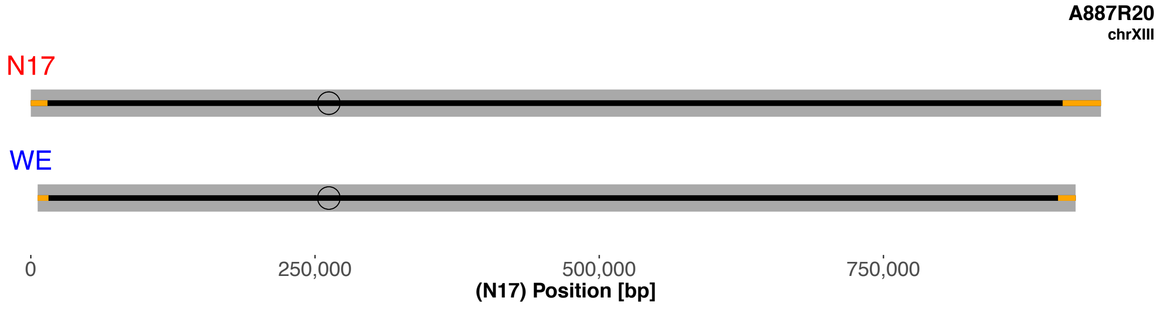
A887R20
chrXIII
N17
WE
0
250,000
500,000
750,000
(N17) Position [bp]

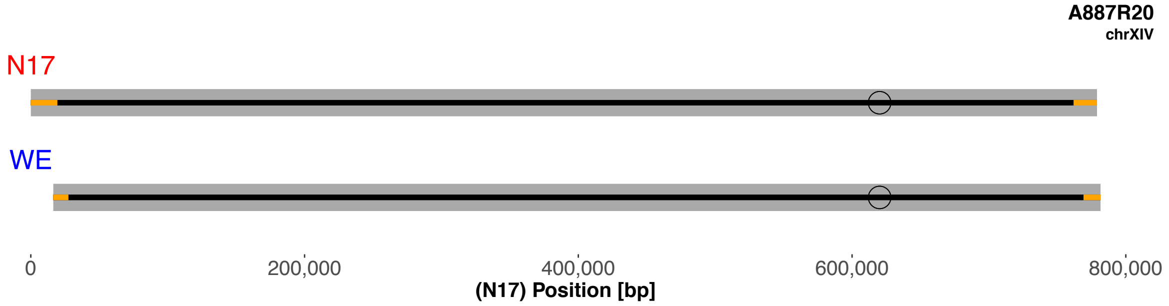
A887R20
chrXIV
N17
WE
0
200,000
400,000
600,000
800,000
(N17) Position [bp]

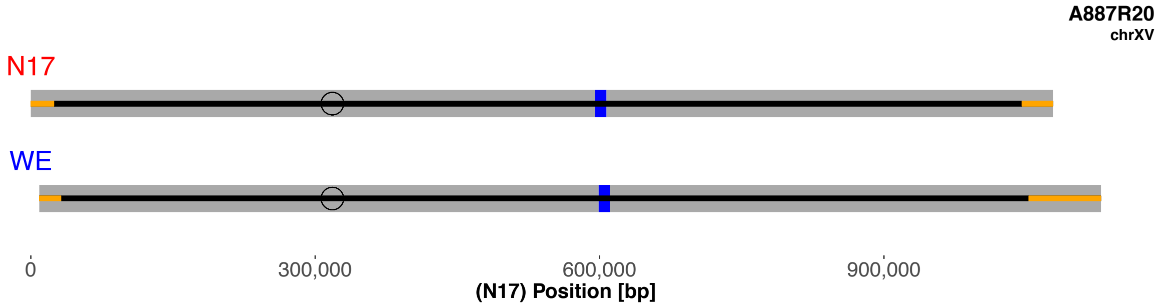
A887R20
chrXV
N17
WE
0
300,000
600,000
900,000
(N17) Position [bp]

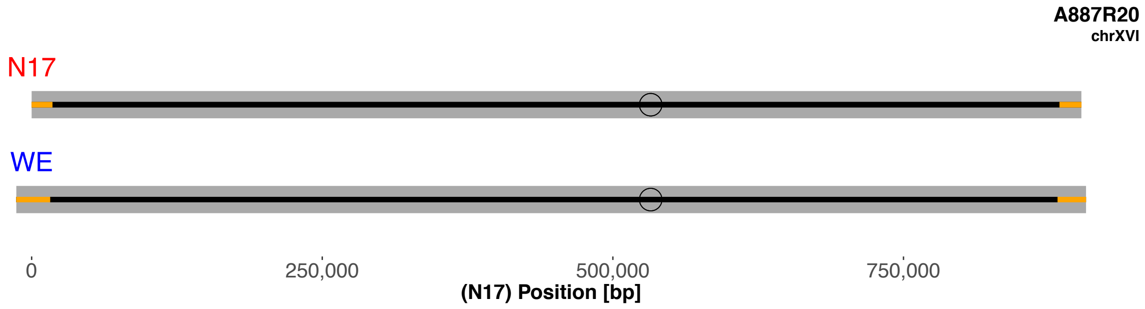
A887R20
chrXVI
N17
WE
0
250,000
500,000
750,000
(N17) Position [bp]

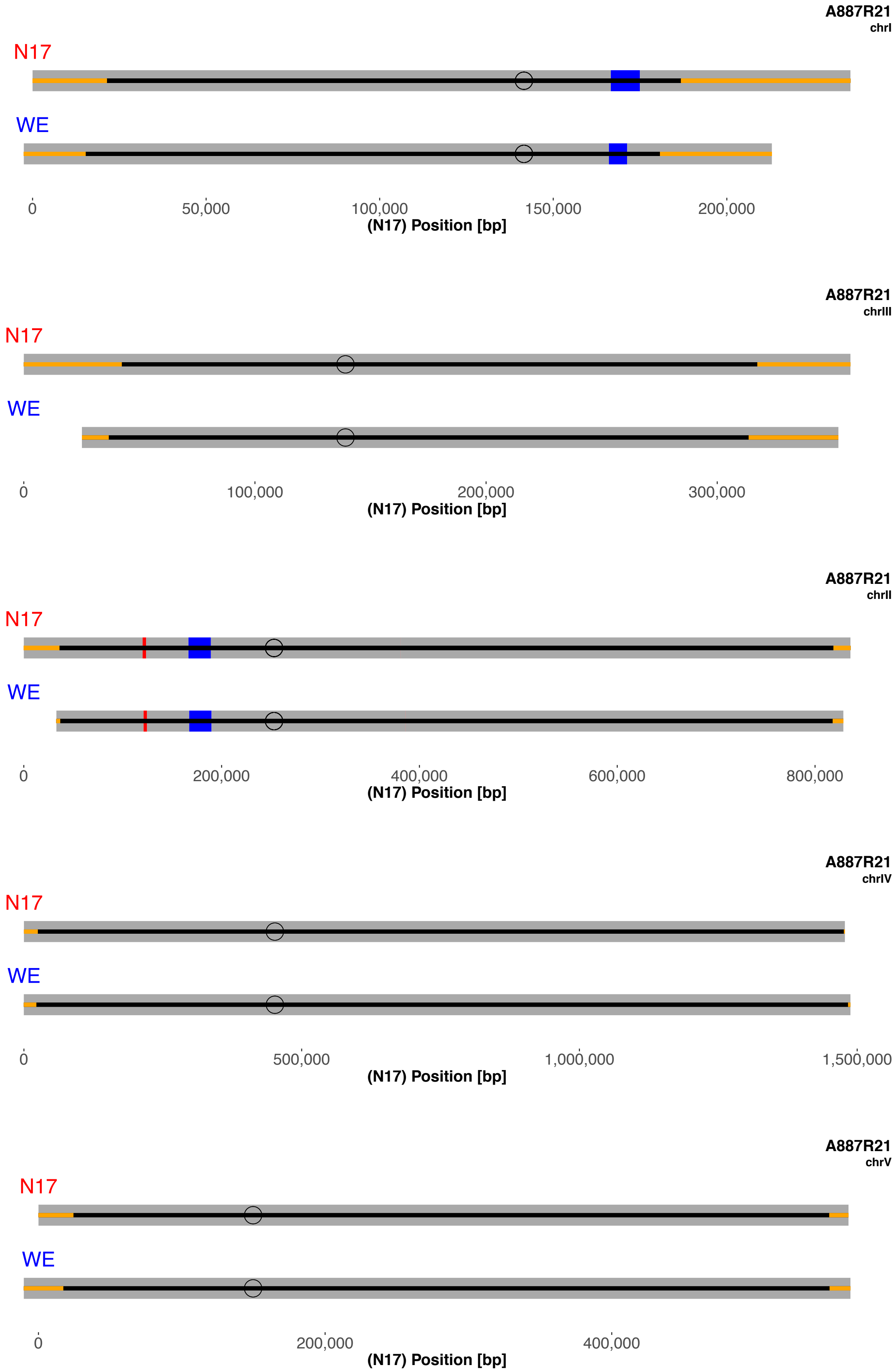
A887R21
chrI
N17
WE
0
50,000
100,000
150,000
200,000
(N17) Position [bp]
A887R21
chrIII
N17
WE
0
100,000
200,000
300,000
(N17) Position [bp]
A887R21
chrII
N17
WE
0
200,000
400,000
600,000
800,000
(N17) Position [bp]
A887R21
chrIV
N17
WE
0
500,000
1,000,000
1,500,000
(N17) Position [bp]
A887R21
chrV
N17
WE
0
200,000
400,000
(N17) Position [bp]

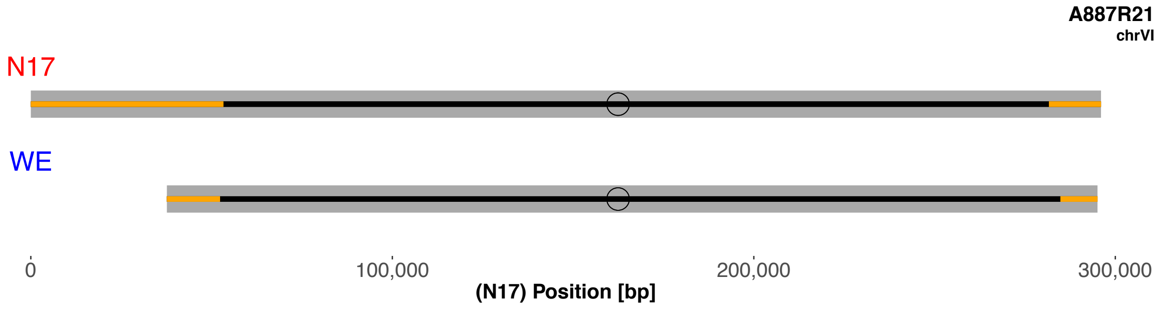
A887R21
chrVI
N17
WE
0
100,000
200,000
300,000
(N17) Position [bp]

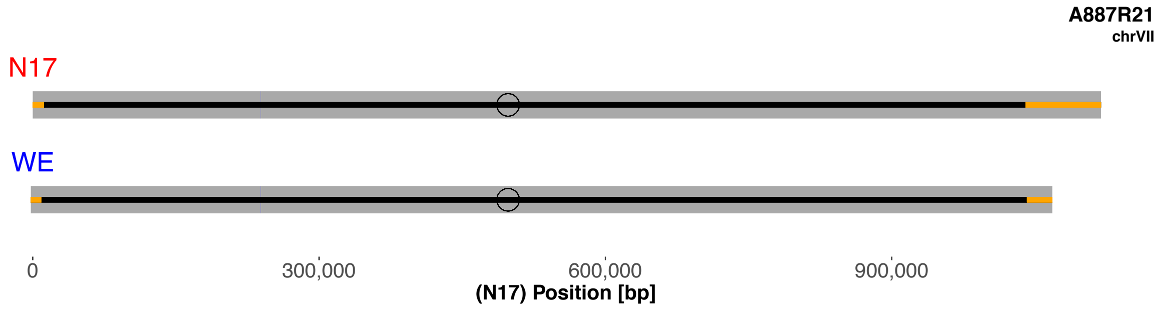
A887R21
chrVII
N17
WE
0
300,000
600,000
900,000
(N17) Position [bp]

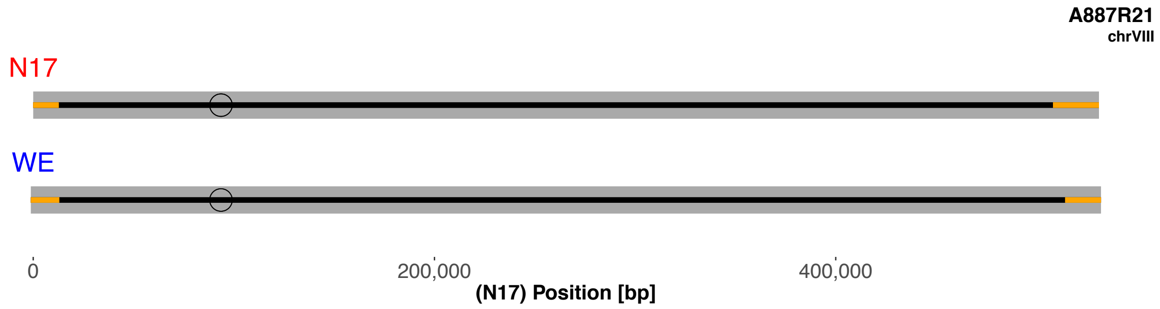
A887R21
chrVIII
N17
WE
0
200,000
400,000
(N17) Position [bp]

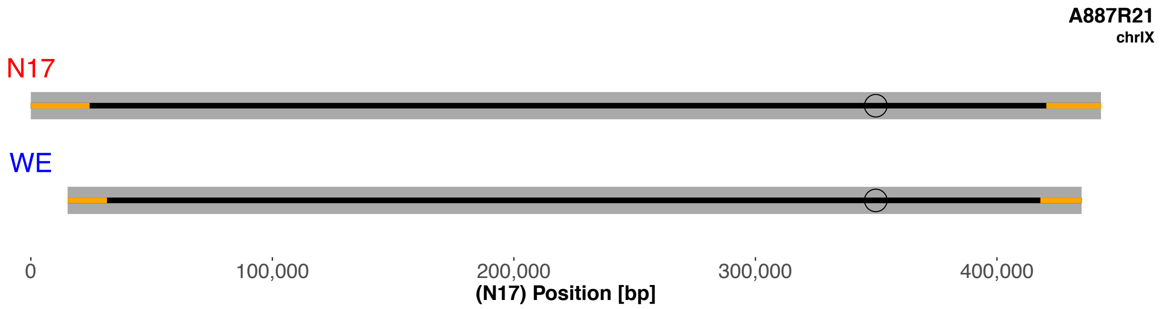
A887R21
chrIX
N17
WE
0
100,000
200,000
300,000
400,000
(N17) Position [bp]

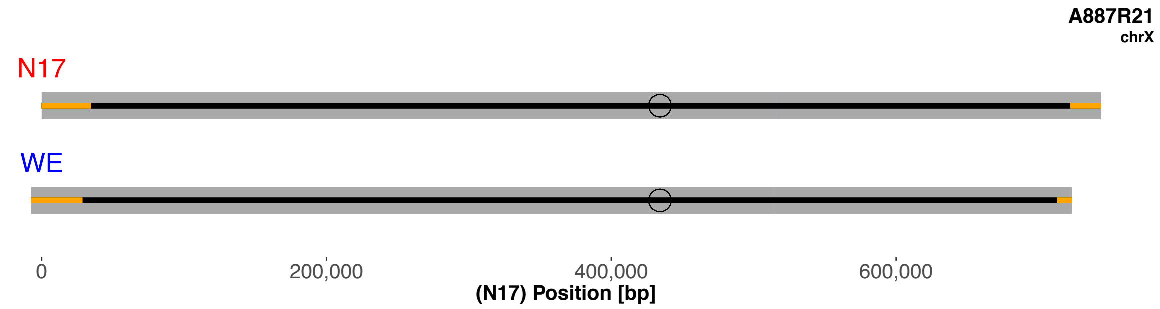
A887R21
chrX
N17
WE
0
200,000
400,000
600,000
(N17) Position [bp]

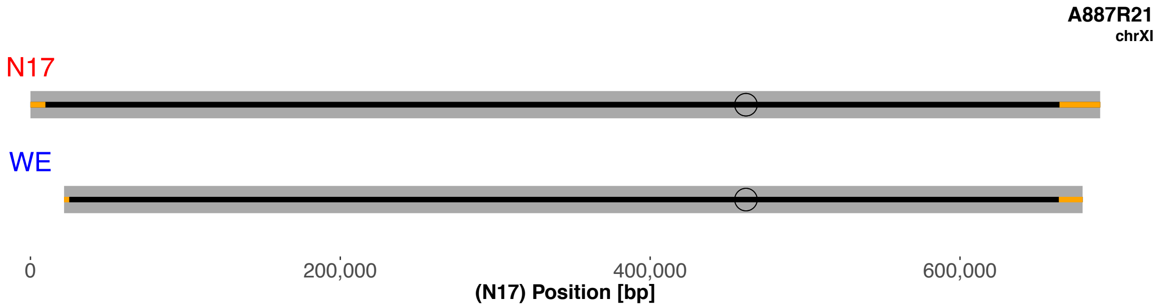
A887R21
chrXI
N17
WE
0
200,000
400,000
600,000
(N17) Position [bp]

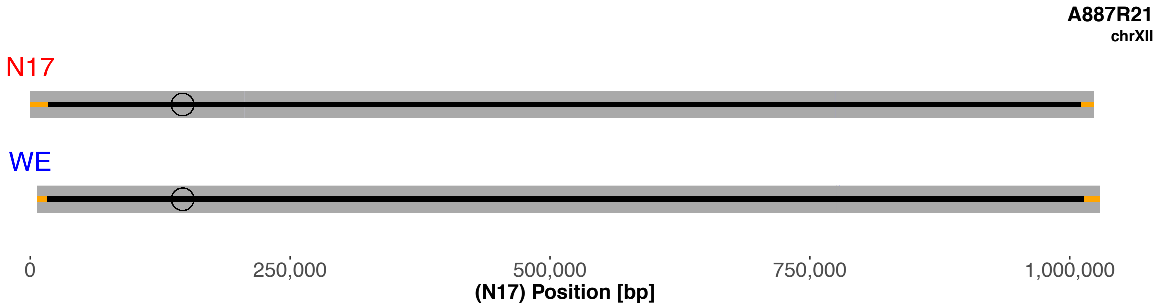
A887R21
chrXII
N17
WE
0
250,000
500,000
750,000
1,000,000
(N17) Position [bp]

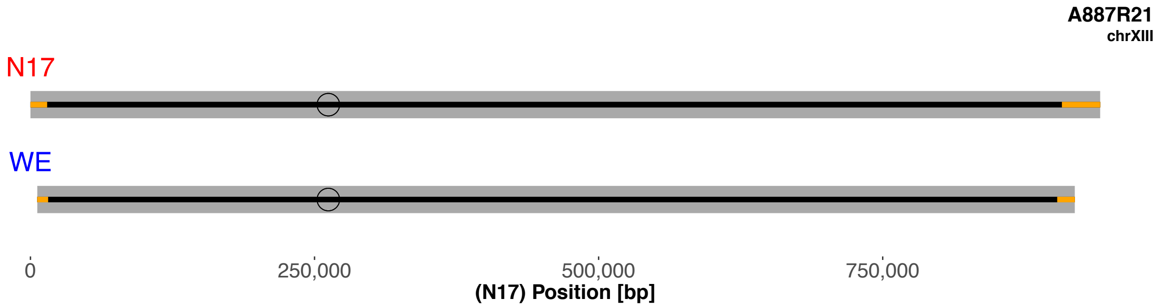
A887R21
chrXIII
N17
WE
0
250,000
500,000
750,000
(N17) Position [bp]

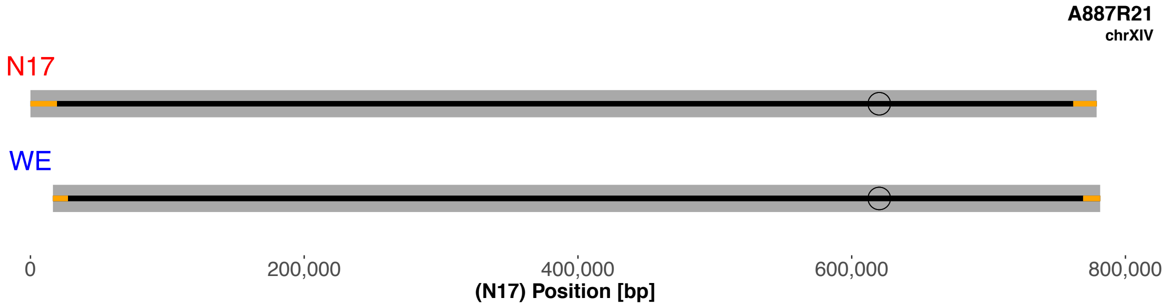
A887R21
chrXIV
N17
WE
0
200,000
400,000
600,000
800,000
(N17) Position [bp]

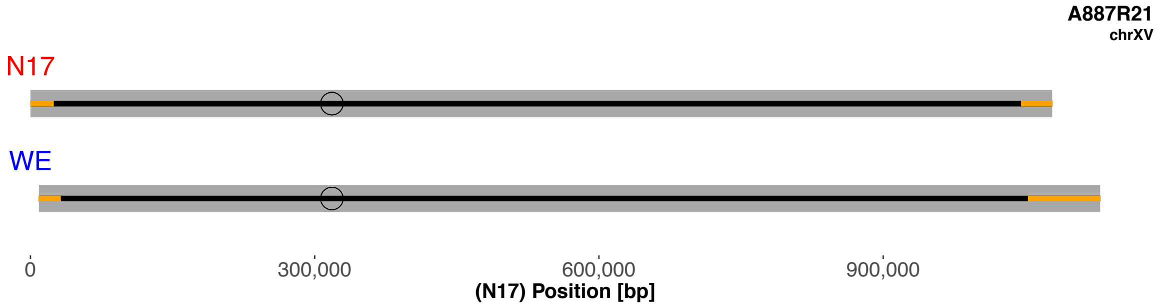
A887R21
chrXV
N17
WE
0
300,000
600,000
900,000
(N17) Position [bp]

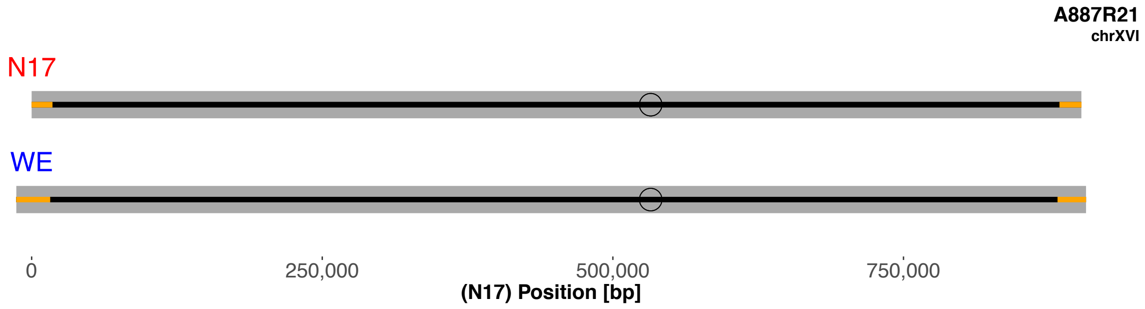
A887R21
chrXVI
N17
WE
0
250,000
500,000
750,000
(N17) Position [bp]

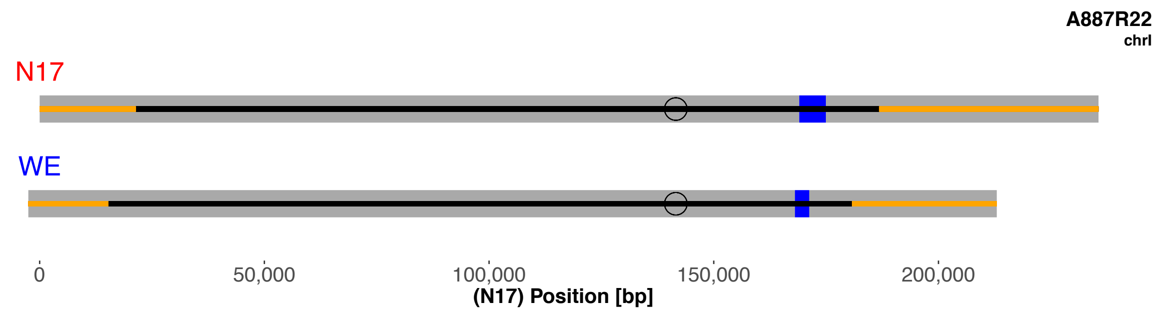
A887R22
chrI
N17
WE
0
50,000
100,000
150,000
200,000
(N17) Position [bp]

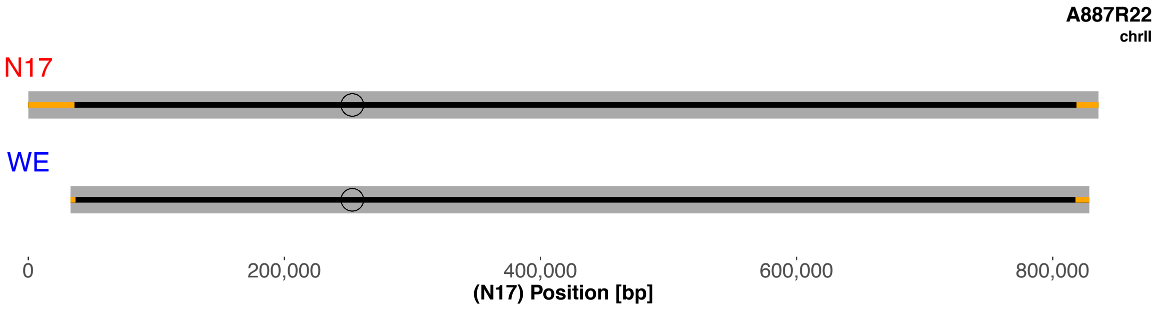
A887R22
chrII
N17
WE
0
200,000
400,000
600,000
800,000
(N17) Position [bp]

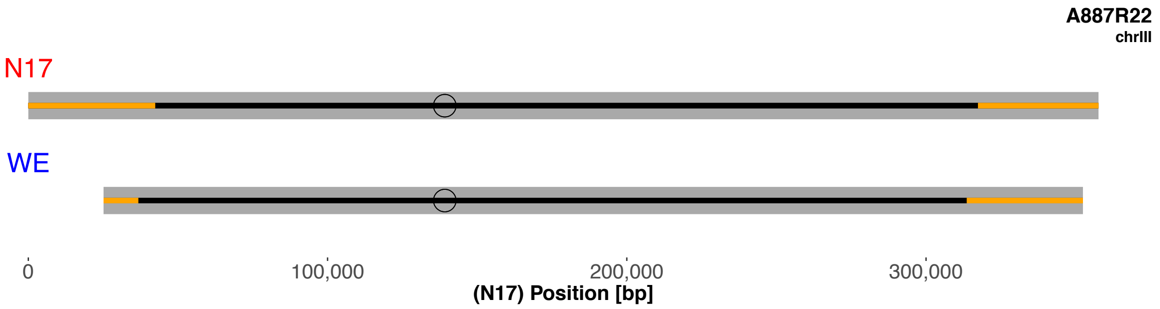
A887R22
chrIII
N17
WE
0
100,000
200,000
300,000
(N17) Position [bp]

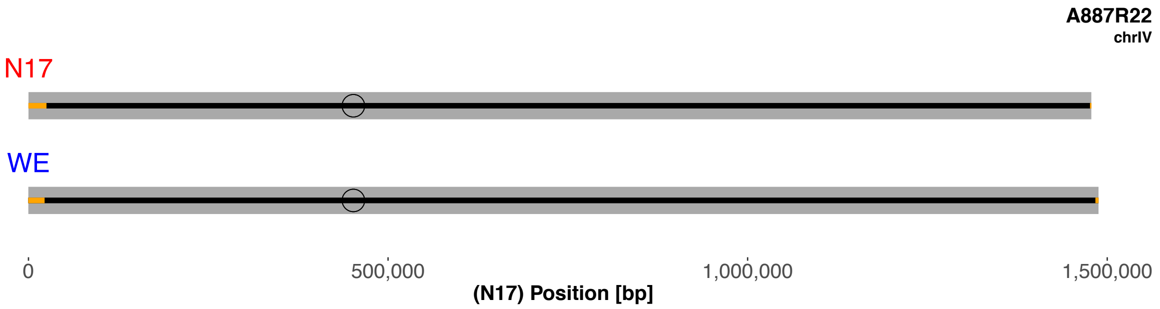
A887R22
chrIV
N17
WE
0
500,000
1,000,000
1,500,000
(N17) Position [bp]

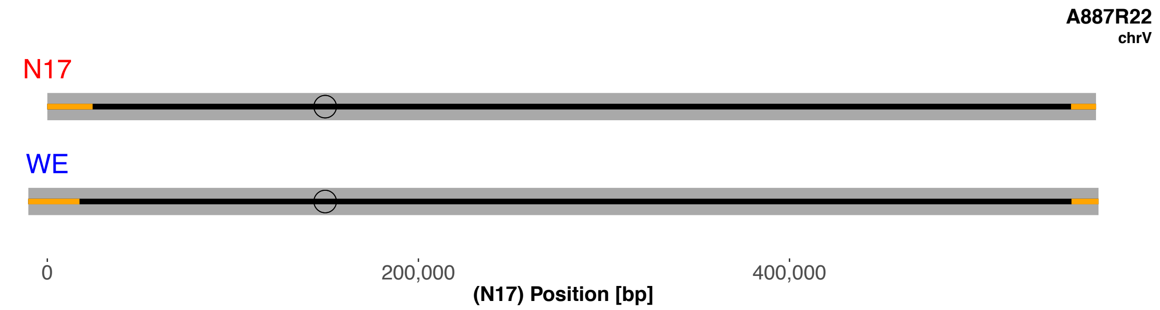
A887R22
chrV
N17
WE
0
200,000
400,000
(N17) Position [bp]

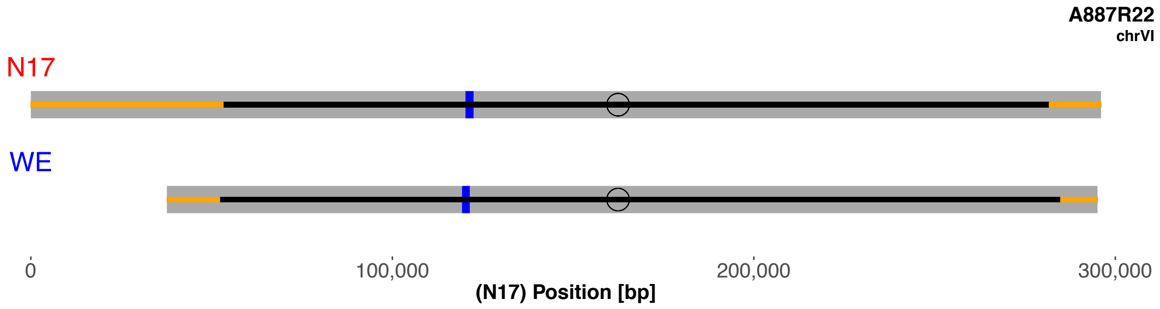
A887R22
chrVI
N17
WE
0
100,000
200,000
300,000
(N17) Position [bp]

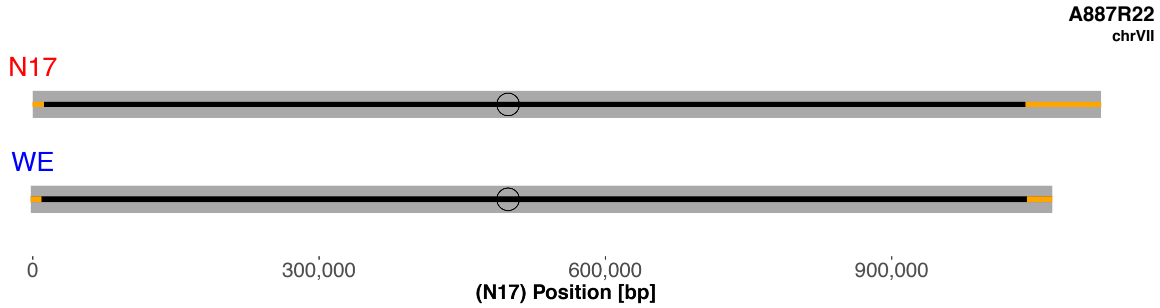
A887R22
chrVII
N17
WE
0
300,000
600,000
900,000
(N17) Position [bp]

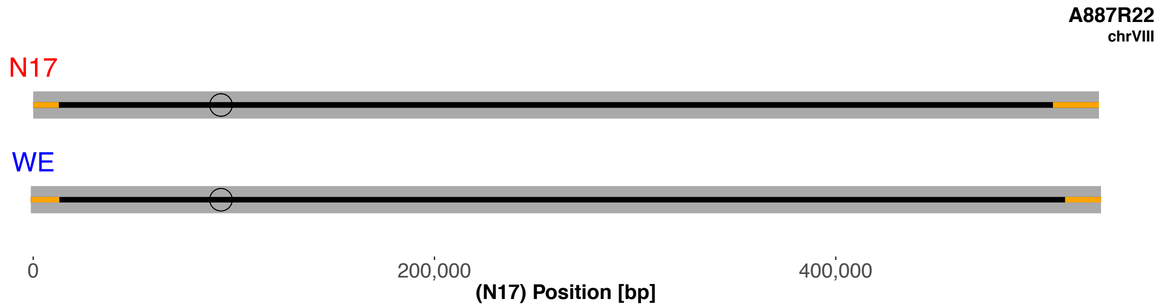
A887R22
chrVIII
N17
WE
0
200,000
400,000
(N17) Position [bp]

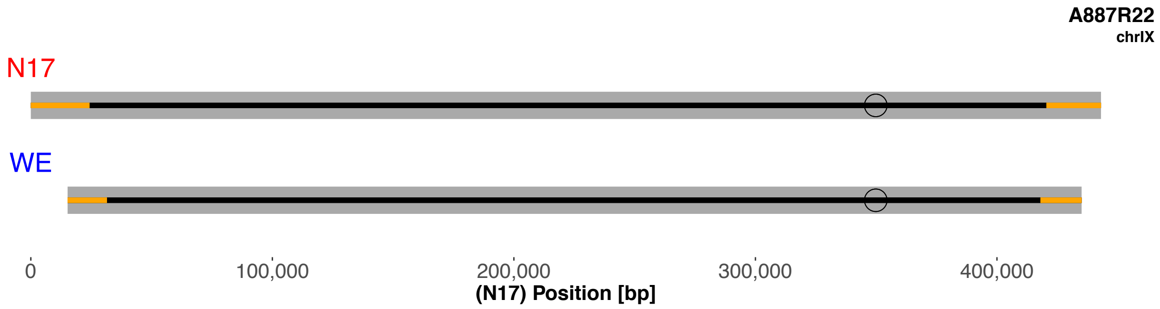
A887R22
chrIX
N17
WE
0
100,000
200,000
300,000
400,000
(N17) Position [bp]

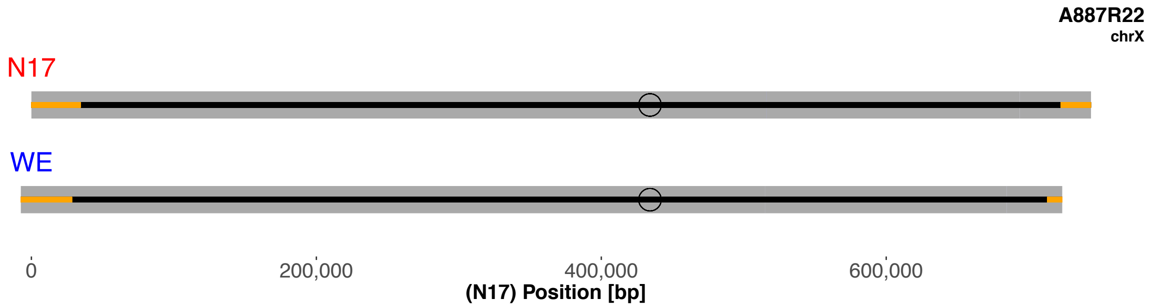
A887R22
chrX
N17
WE
0
200,000
400,000
600,000
(N17) Position [bp]

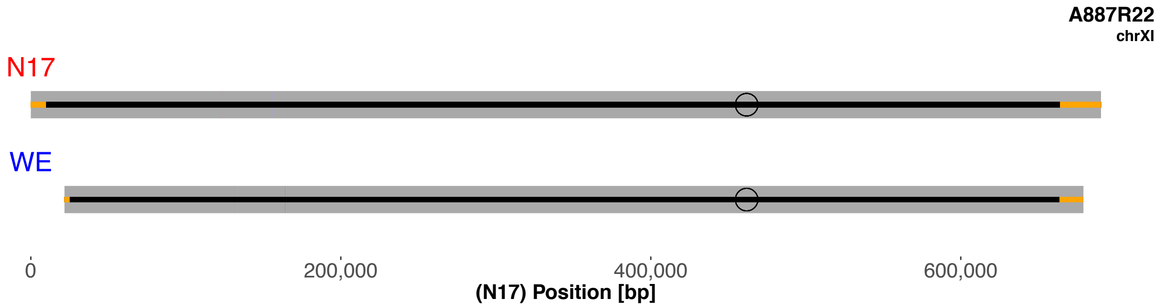
A887R22
chrXI
N17
WE
0
200,000
400,000
600,000
(N17) Position [bp]

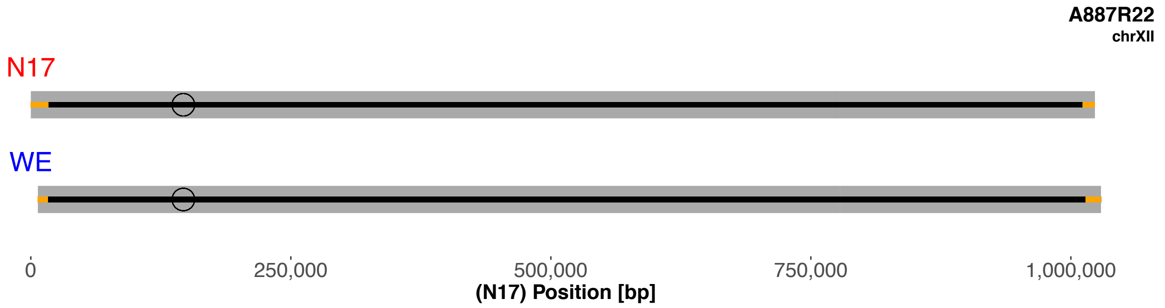
A887R22
chrXII
N17
WE
0
250,000
500,000
750,000
1,000,000
(N17) Position [bp]

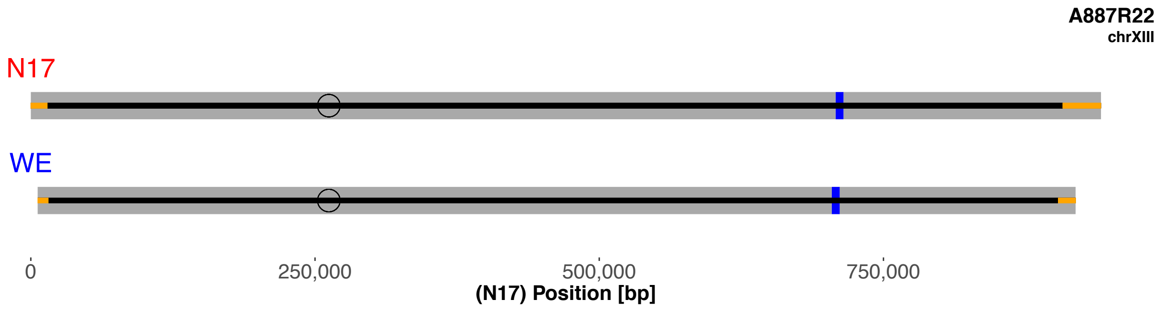
A887R22
chrXIII
N17
WE
0
250,000
500,000
750,000
(N17) Position [bp]

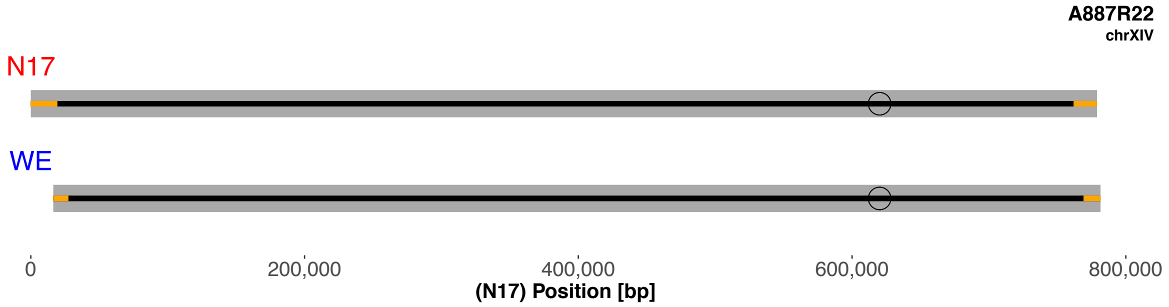
A887R22
chrXIV
N17
WE
0
200,000
400,000
600,000
800,000
(N17) Position [bp]

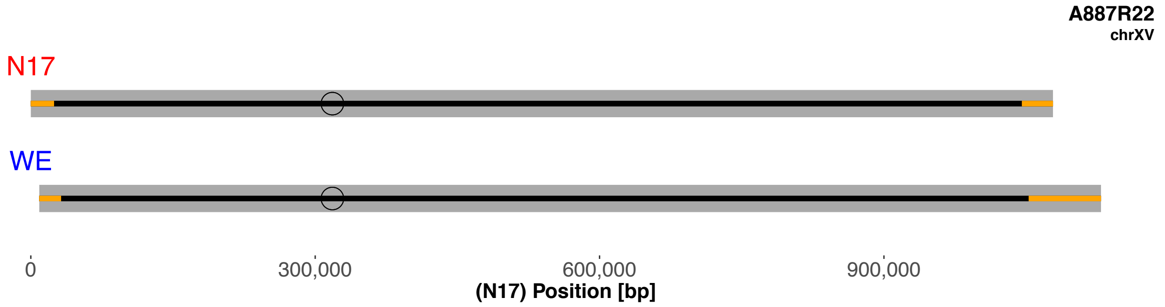
A887R22
chrXV
N17
WE
0
300,000
600,000
900,000
(N17) Position [bp]

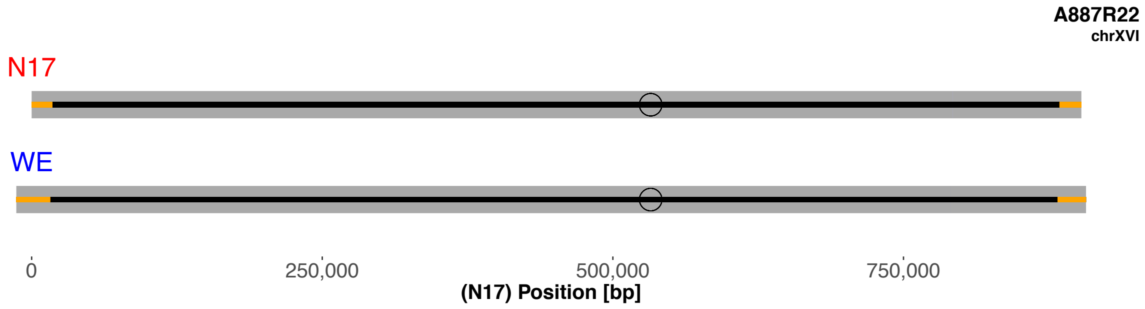
A887R22
chrXVI
N17
WE
0
250,000
500,000
750,000
(N17) Position [bp]

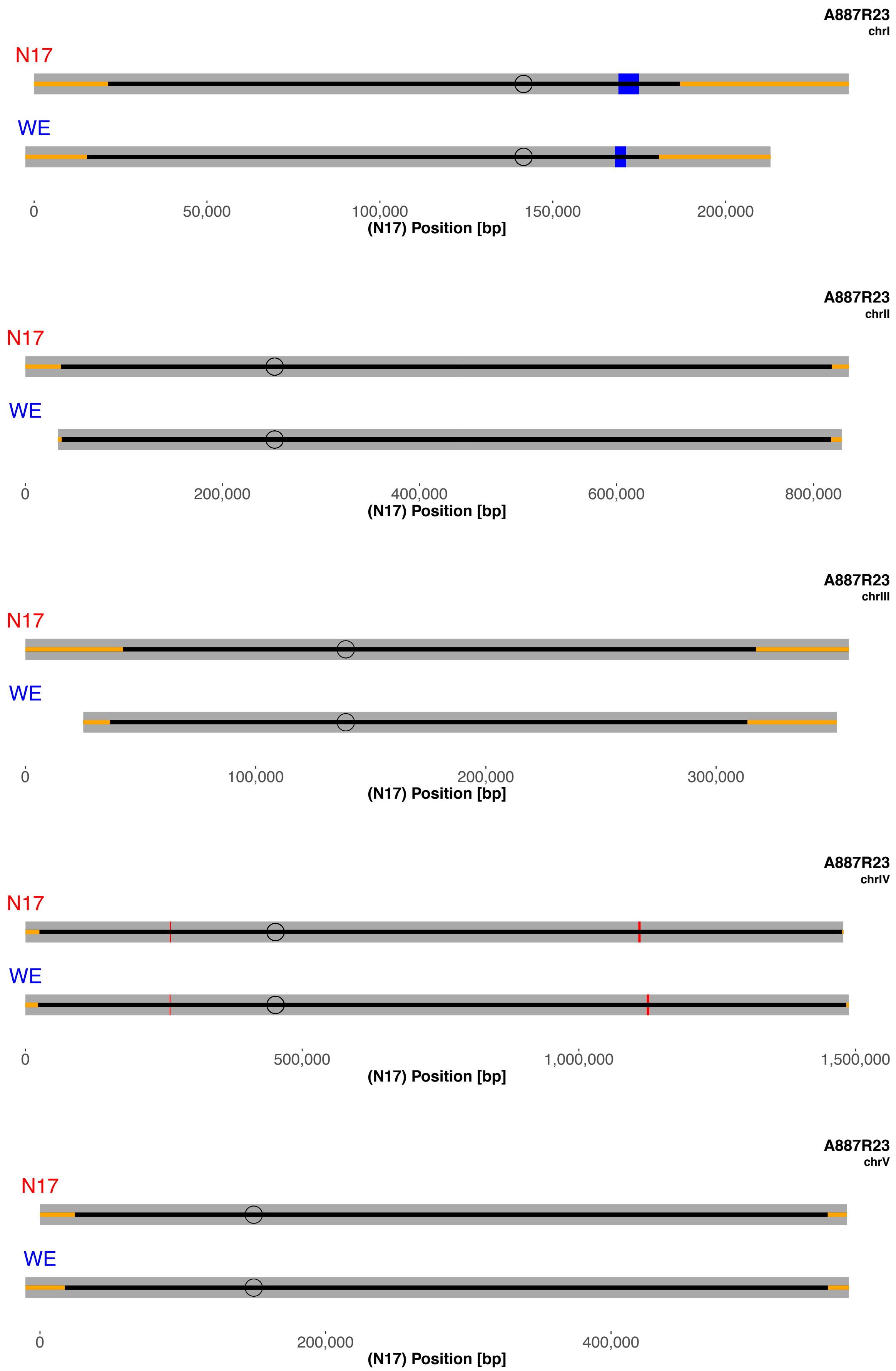

A887R23
chrI
N17
WE
0
50,000
100,000
150,000
200,000
(N17) Position [bp]
A887R23
chrII
N17
WE
0
200,000
400,000
600,000
800,000
(N17) Position [bp]
A887R23
chrIII
N17
WE
0
100,000
200,000
300,000
(N17) Position [bp]
A887R23
chrIV
N17
WE
0
500,000
1,000,000
1,500,000
(N17) Position [bp]
A887R23
chrV
N17
WE
0
200,000
400,000
(N17) Position [bp]

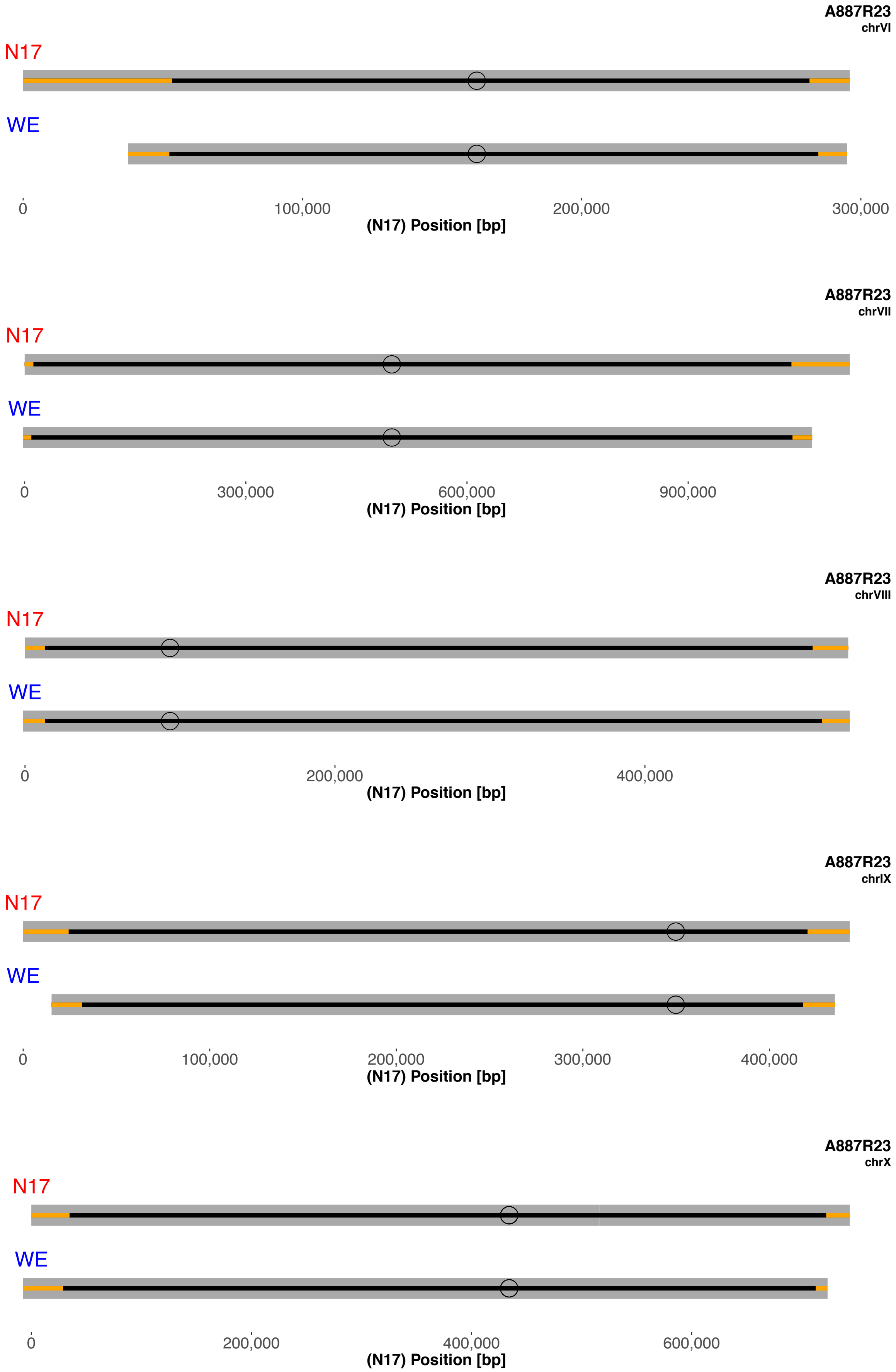
A887R23
chrVI
N17
WE
0
100,000
200,000
300,000
(N17) Position [bp]
A887R23
chrVII
N17
WE
0
300,000
600,000
900,000
(N17) Position [bp]
A887R23
chrVIII
N17
WE
0
200,000
400,000
(N17) Position [bp]
A887R23
chrIX
N17
WE
0
100,000
200,000
300,000
400,000
(N17) Position [bp]
A887R23
chrX
N17
WE
0
200,000
400,000
600,000
(N17) Position [bp]

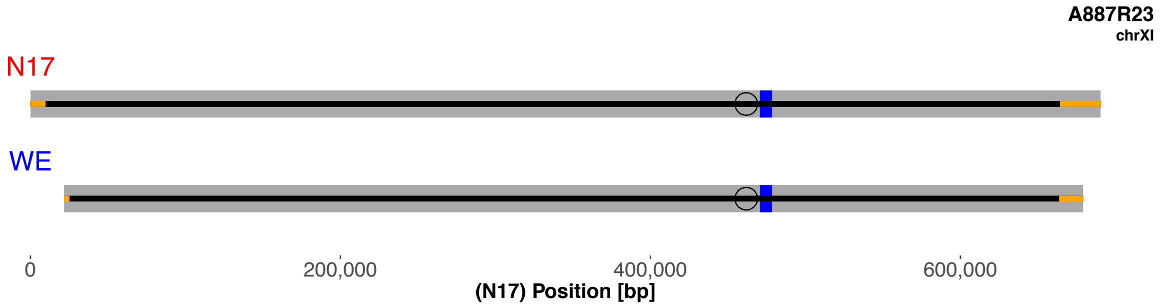
A887R23
chrXI
N17
WE
0
200,000
400,000
600,000
(N17) Position [bp]

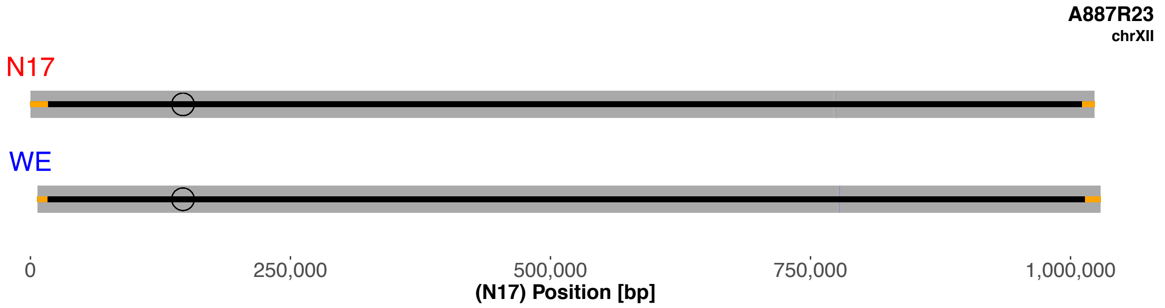
A887R23
chrXII
N17
WE
0
250,000
500,000
750,000
1,000,000
(N17) Position [bp]

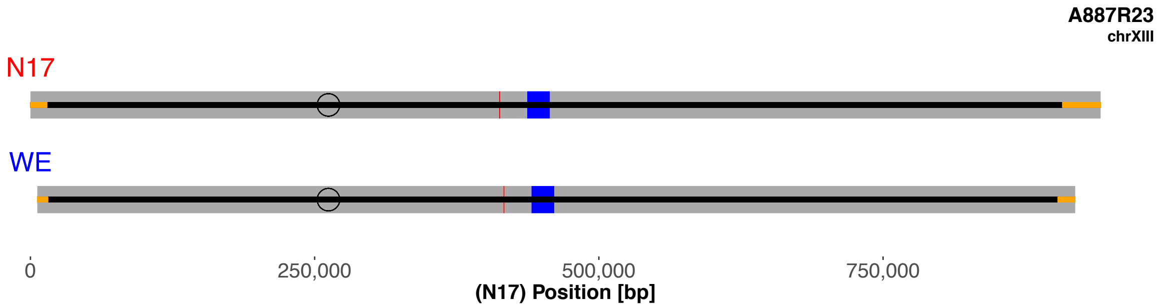
A887R23
chrXIII
N17
WE
0
250,000
500,000
750,000
(N17) Position [bp]

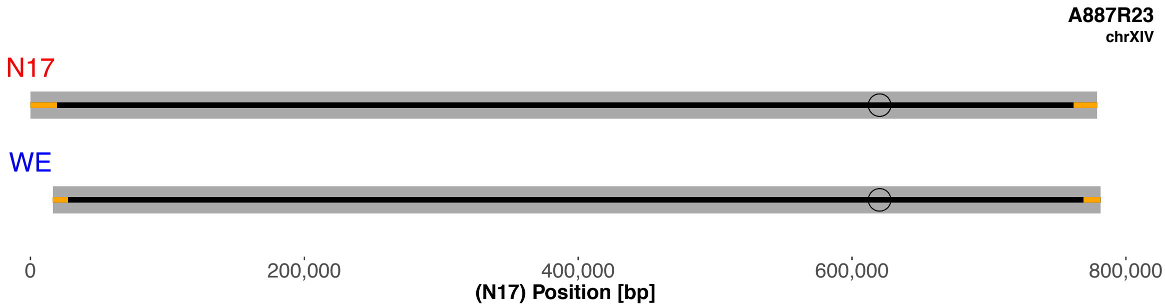
A887R23
chrXIV
N17
WE
0
200,000
400,000
600,000
800,000
(N17) Position [bp]

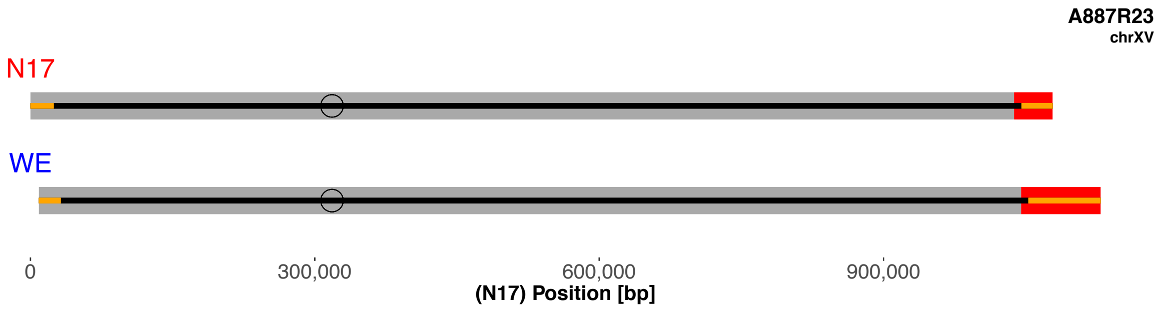
A887R23
chrXV
N17
WE
0
300,000
600,000
900,000
(N17) Position [bp]

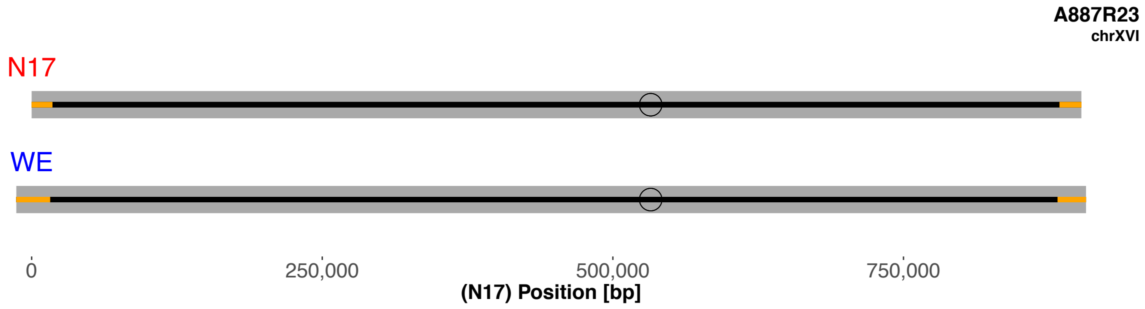
A887R23
chrXVI
N17
WE
0
250,000
500,000
750,000
(N17) Position [bp]

Aneuploidy: chromosome I N17 deleted.

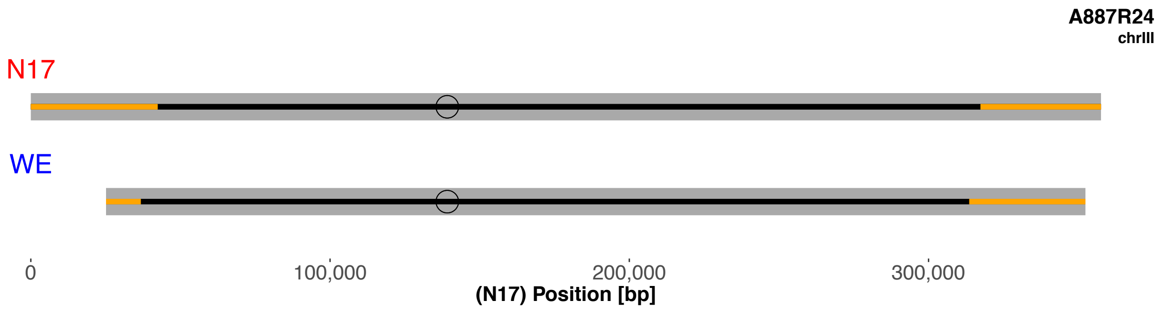


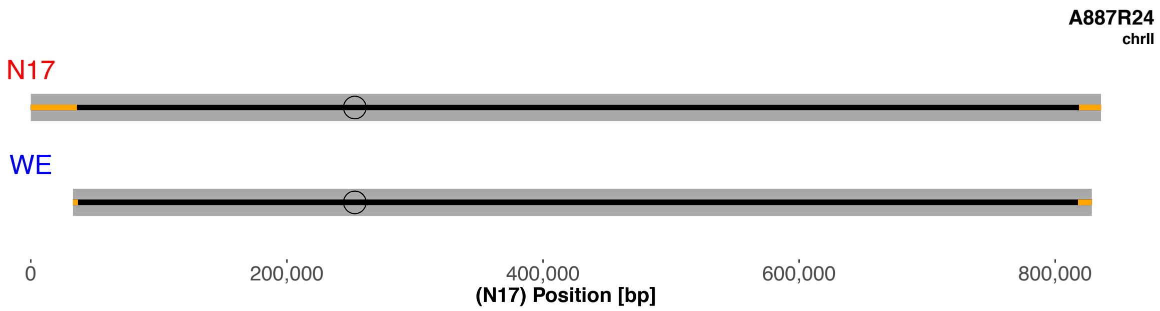


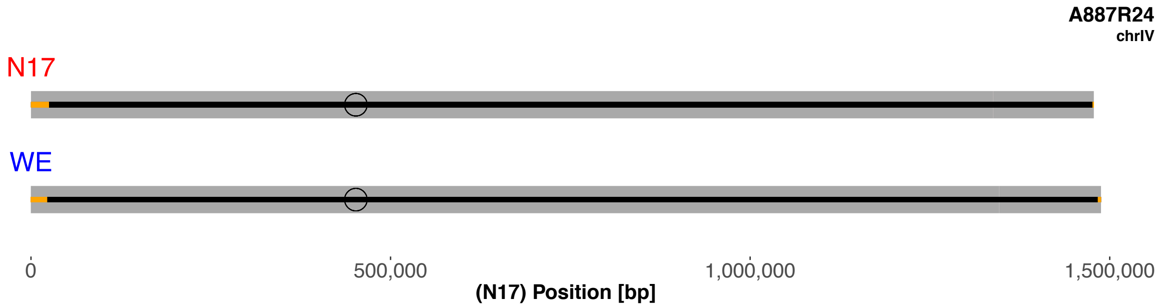


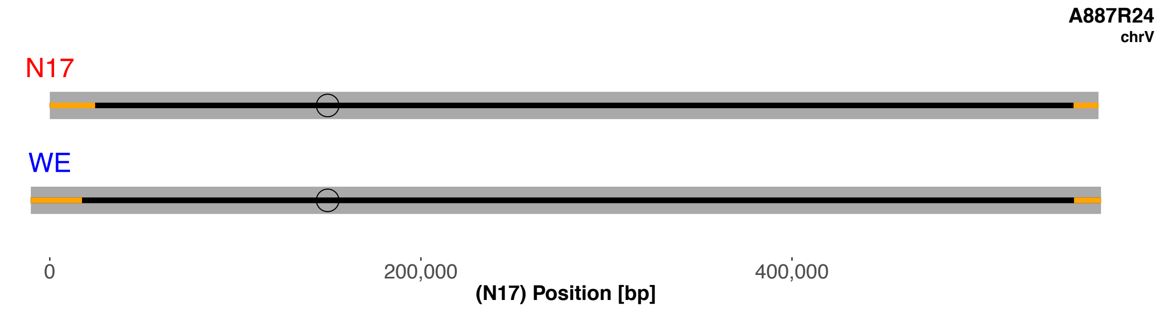

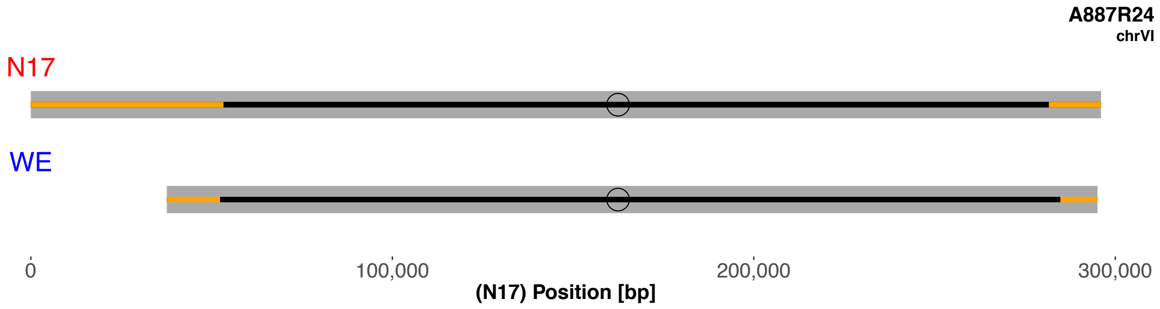
A887R24
chrVI
N17
WE
0
100,000
200,000
300,000
(N17) Position [bp]

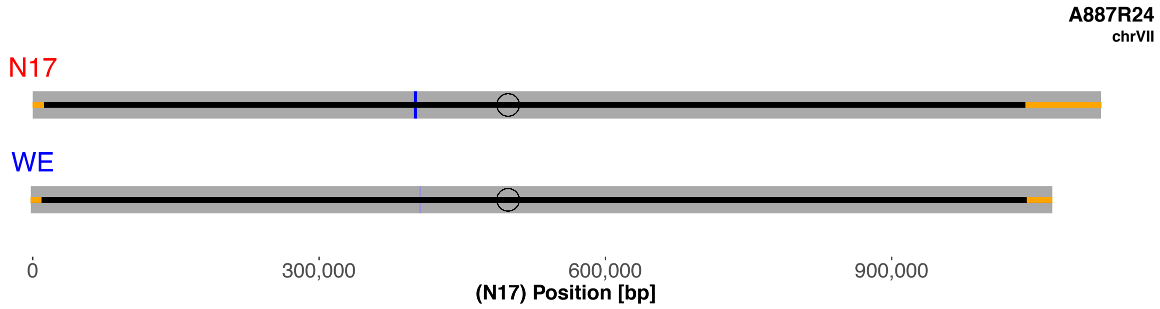
A887R24
chrVII
N17
WE
0
300,000
600,000
900,000
(N17) Position [bp]

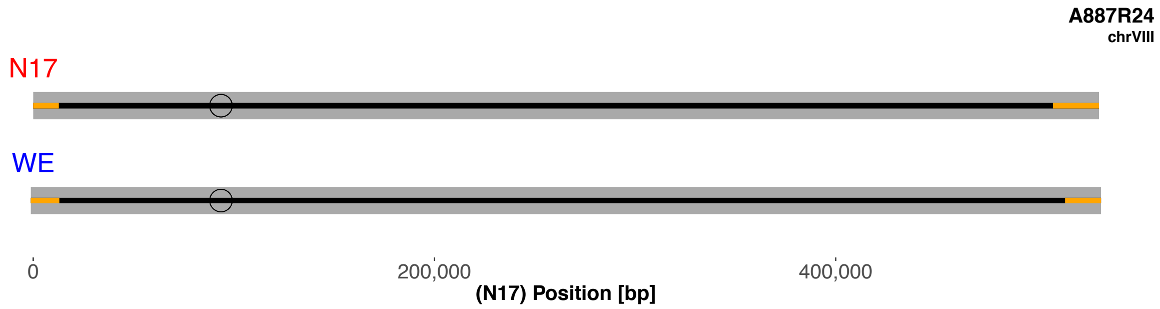
A887R24
chrVIII
N17
WE
0
200,000
400,000
(N17) Position [bp]

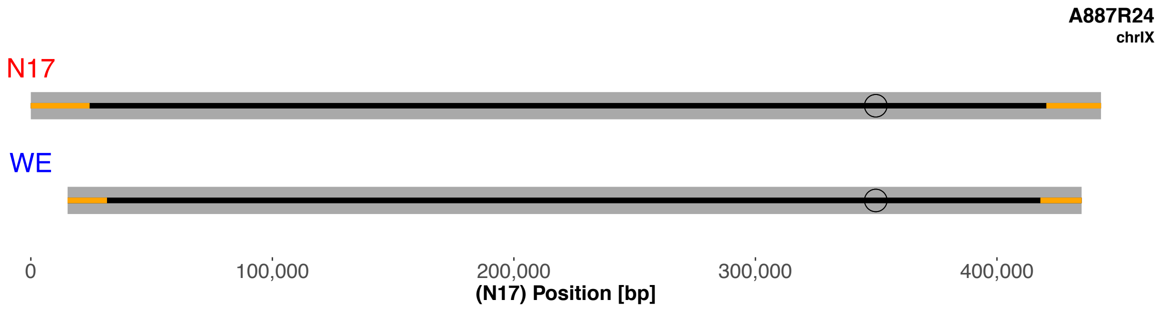
A887R24
chrIX
N17
WE
0
100,000
200,000
300,000
400,000
(N17) Position [bp]

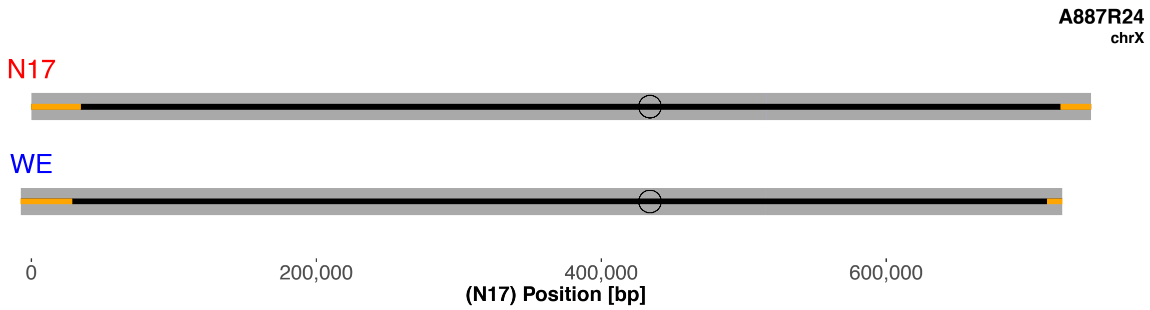
A887R24
chrX
N17
WE
0
200,000
400,000
600,000
(N17) Position [bp]

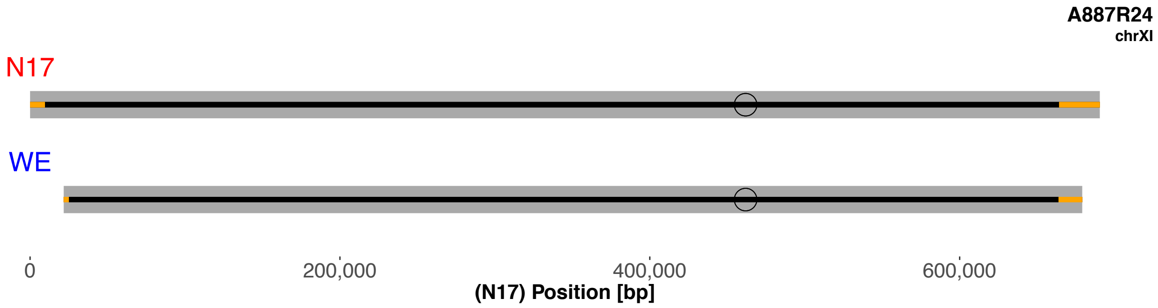
A887R24
chrXI
N17
WE
0
200,000
400,000
600,000
(N17) Position [bp]

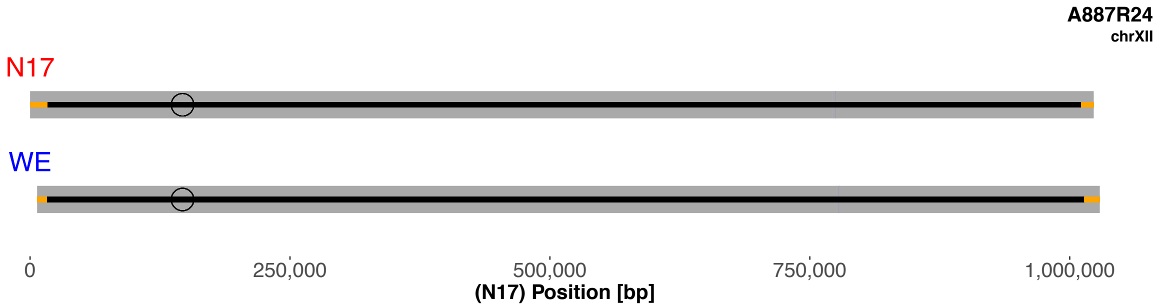
A887R24
chrXII
N17
WE
0
250,000
500,000
750,000
1,000,000
(N17) Position [bp]

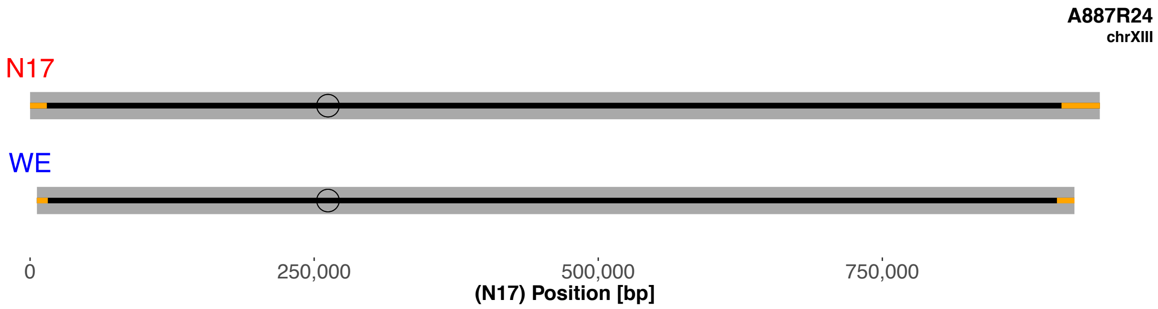
A887R24
chrXIII
N17
WE
0
250,000
500,000
750,000
(N17) Position [bp]

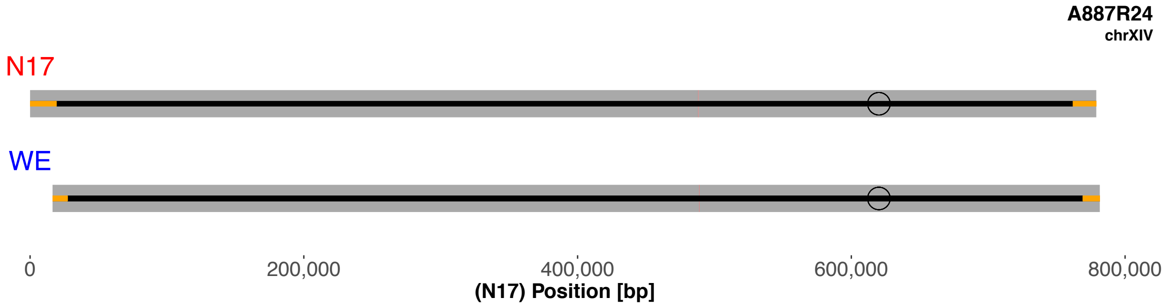
A887R24
chrXIV
N17
WE
0
200,000
400,000
600,000
800,000
(N17) Position [bp]

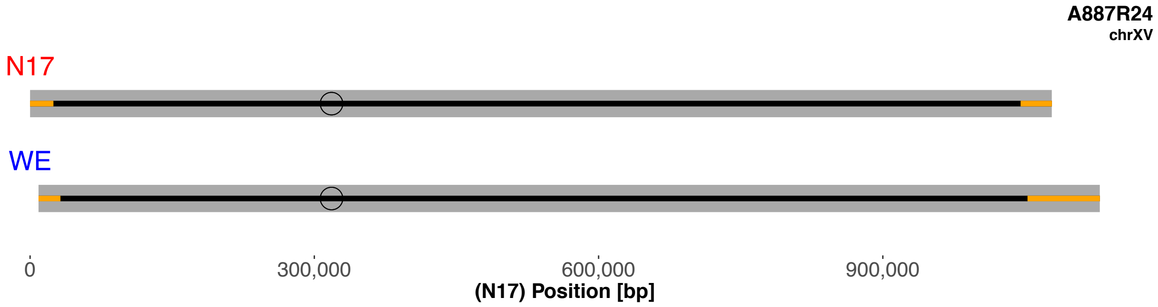
A887R24
chrXV
N17
WE
0
300,000
600,000
900,000
(N17) Position [bp]

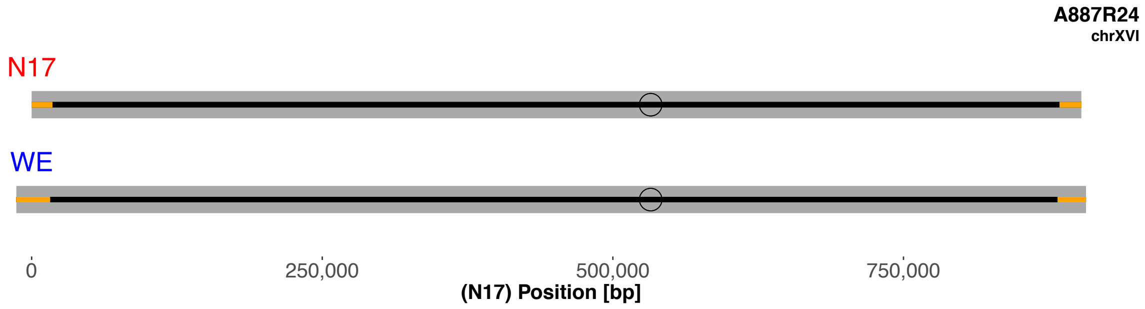
A887R24
chrXVI
N17
WE
0
250,000
500,000
750,000
(N17) Position [bp]

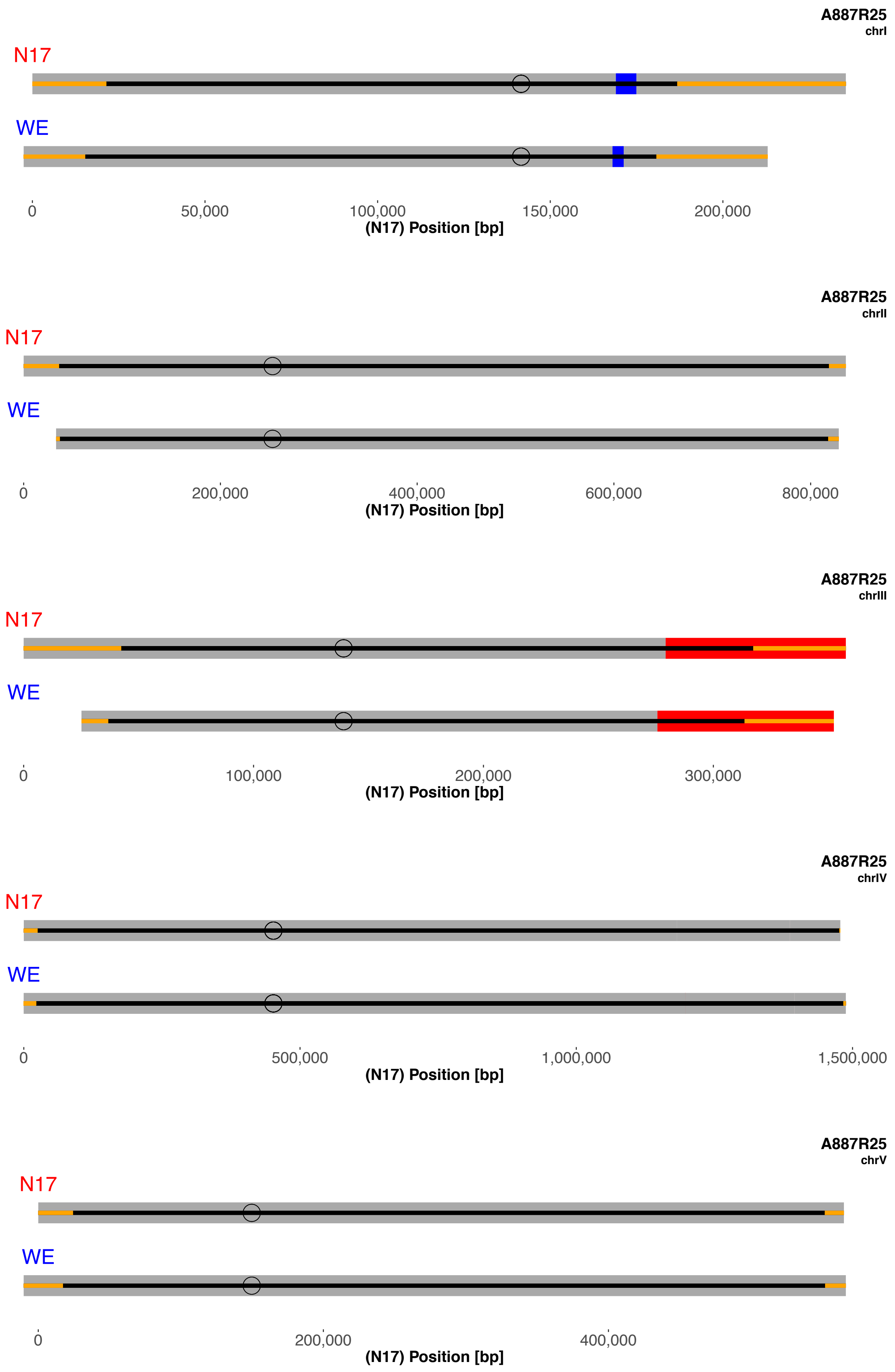
A887R25
chrI
N17
WE
0
50,000
100,000
150,000
200,000
(N17) Position [bp]
A887R25
chrII
N17
WE
0
200,000
400,000
600,000
800,000
(N17) Position [bp]
A887R25
chrIII
N17
WE
0
100,000
200,000
300,000
(N17) Position [bp]
A887R25
chrIV
N17
WE
0
500,000
1,000,000
1,500,000
(N17) Position [bp]
A887R25
chrV
N17
WE
0
200,000
400,000
(N17) Position [bp]

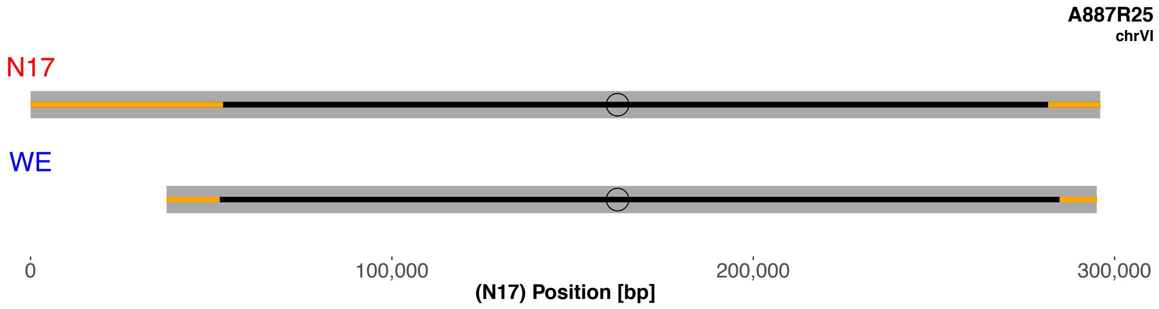
A887R25
chrVI
N17
WE
0
100,000
200,000
300,000
(N17) Position [bp]

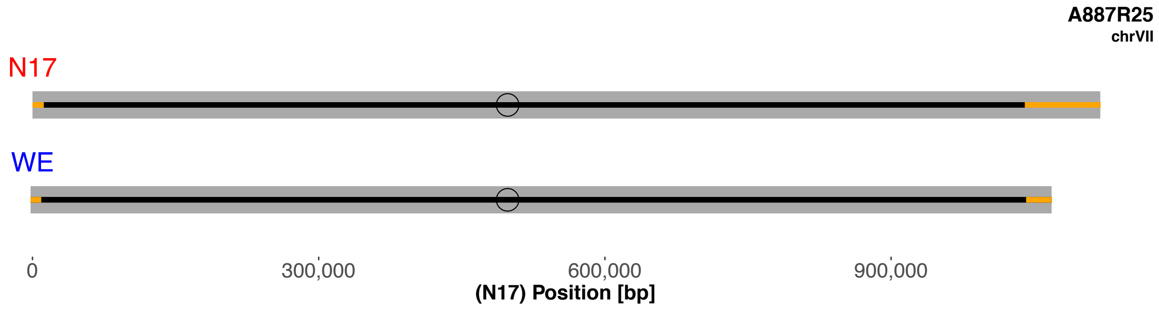
A887R25
chrVII
N17
WE
0
300,000
600,000
900,000
(N17) Position [bp]

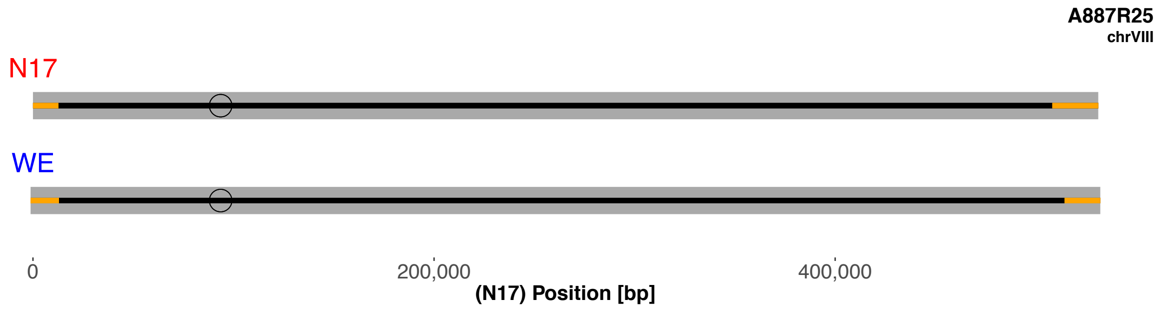
A887R25
chrVIII
N17
WE
0
200,000
400,000
(N17) Position [bp]

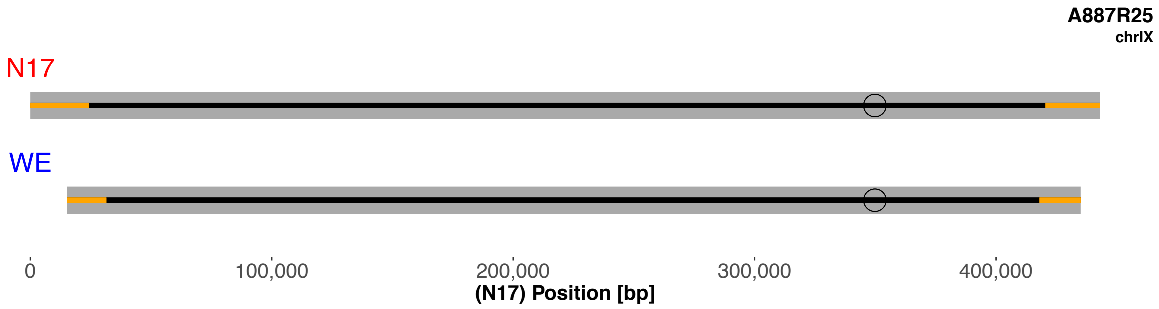
A887R25
chrIX
N17
WE
0
100,000
200,000
300,000
400,000
(N17) Position [bp]

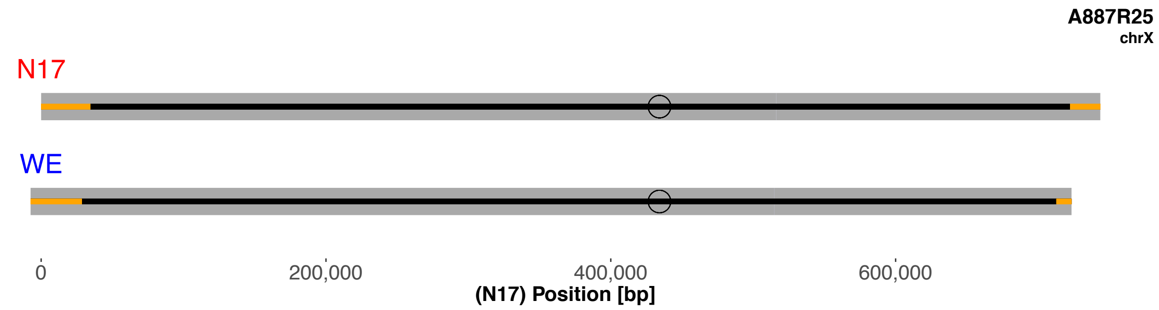
A887R25
chrX
N17
WE
0
200,000
400,000
600,000
(N17) Position [bp]

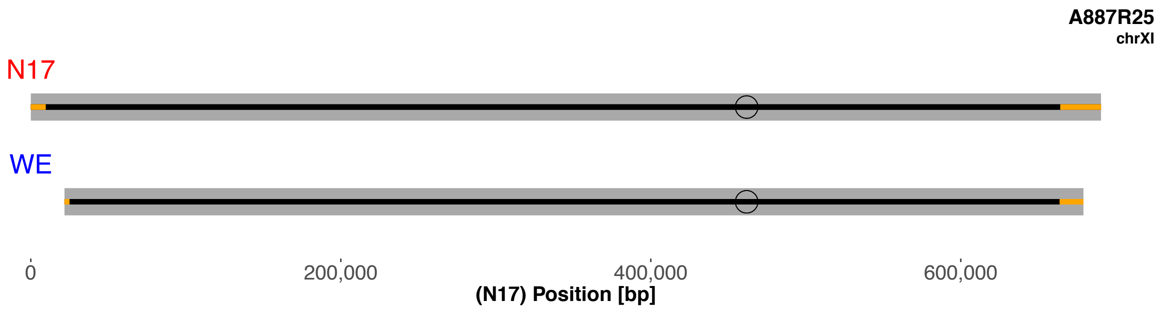
A887R25
chrXI
N17
WE
0
200,000
400,000
600,000
(N17) Position [bp]

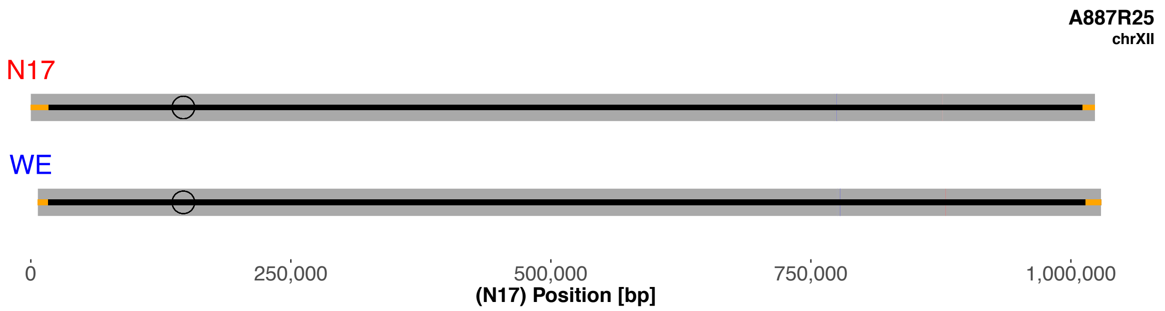
A887R25
chrXII
N17
WE
0
250,000
500,000
750,000
1,000,000
(N17) Position [bp]

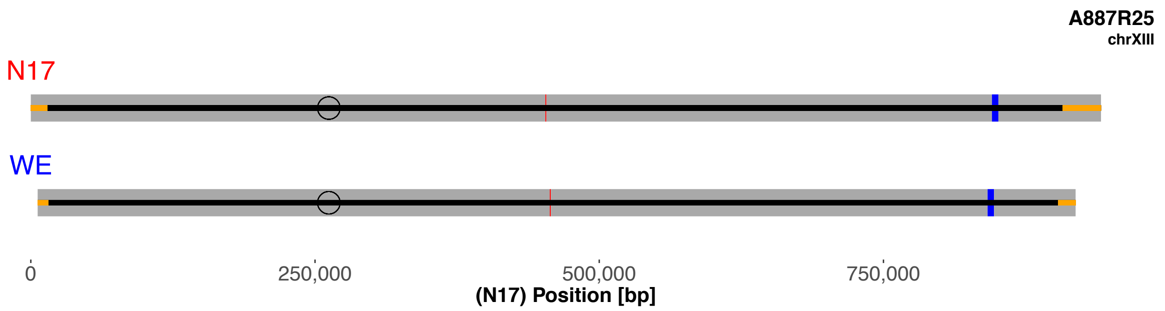
A887R25
chrXIII
N17
WE
0
250,000
500,000
750,000
(N17) Position [bp]

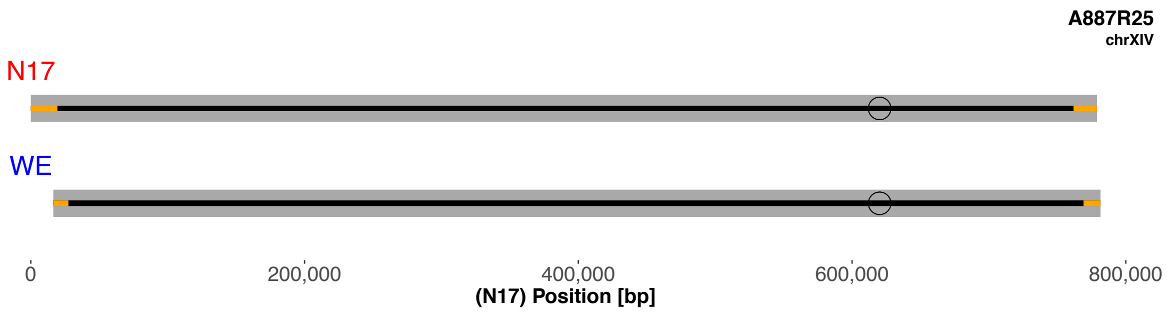
A887R25
chrXIV
N17
WE
0
200,000
400,000
600,000
800,000
(N17) Position [bp]

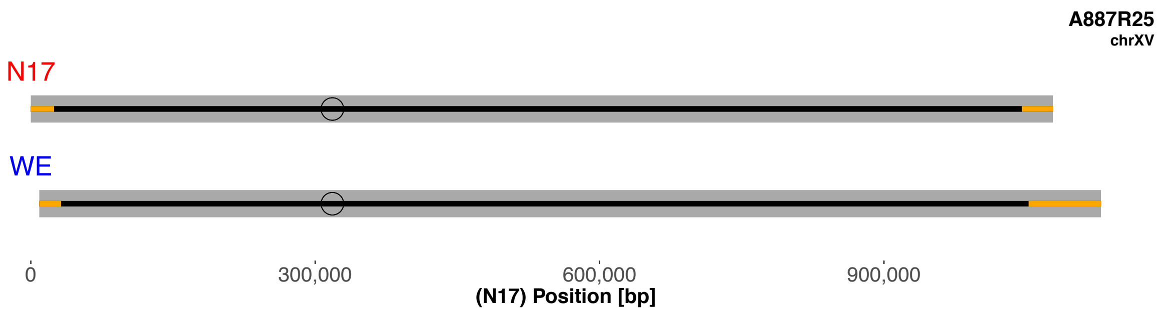
A887R25
chrXV
N17
WE
0
300,000
600,000
900,000
(N17) Position [bp]

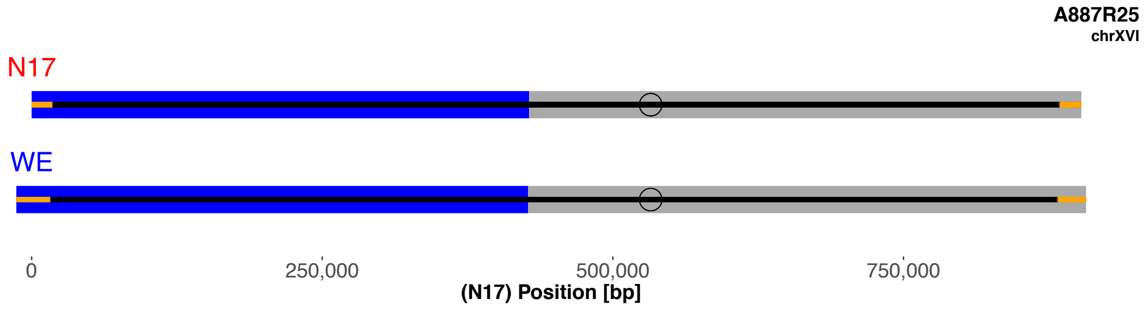
A887R25
chrXVI
N17
WE
0
250,000
500,000
750,000
(N17) Position [bp]

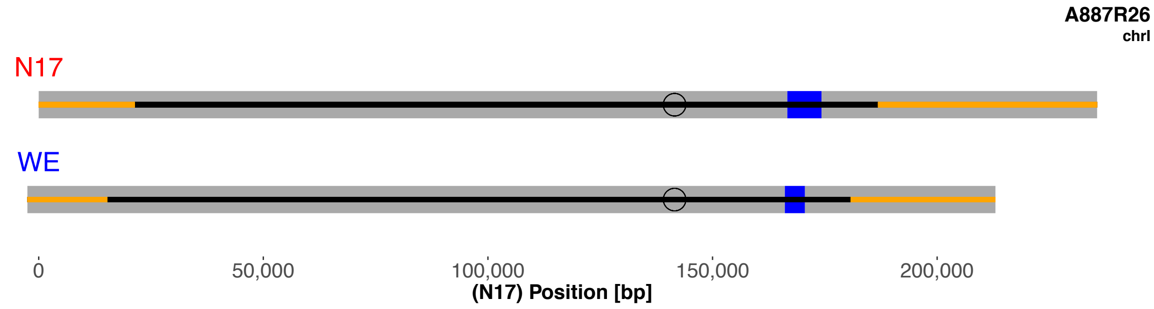
A887R26
chrI
N17
WE
0
50,000
100,000
150,000
200,000
(N17) Position [bp]

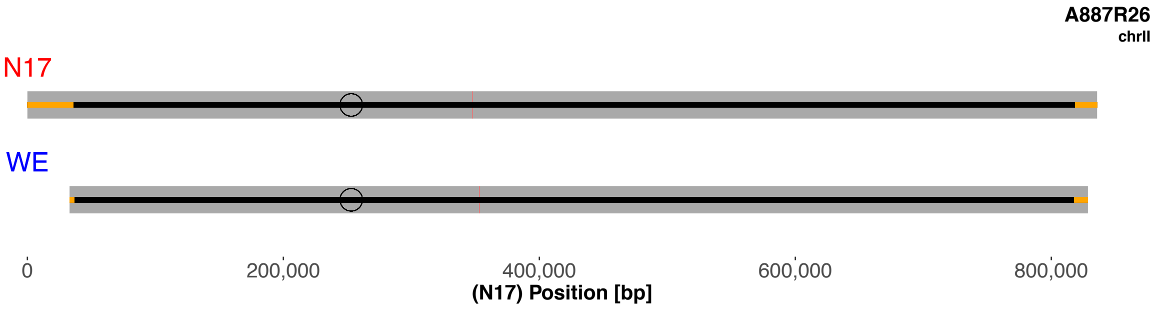
A887R26
chrII
N17
WE
0
200,000
400,000
600,000
800,000
(N17) Position [bp]

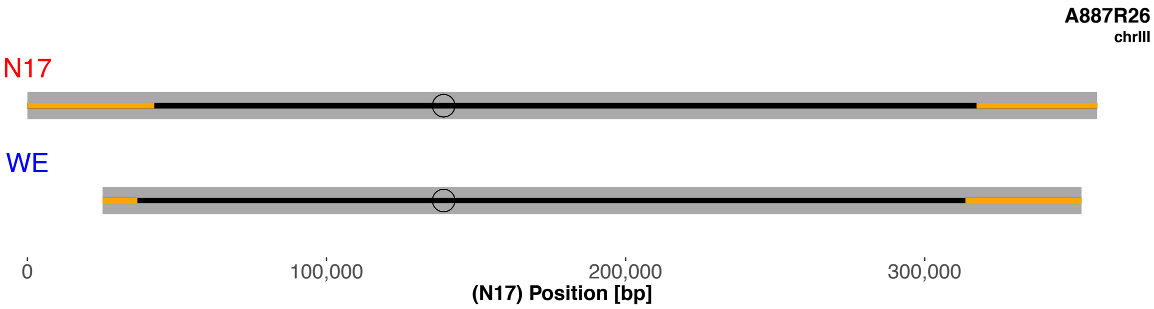
A887R26
chrIII
N17
WE
0
100,000
200,000
300,000
(N17) Position [bp]

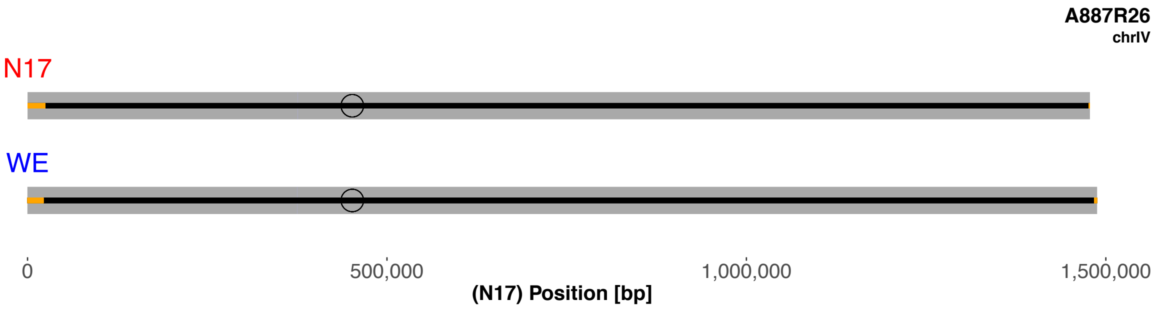
A887R26
chrIV
N17
WE
0
500,000
1,000,000
1,500,000
(N17) Position [bp]

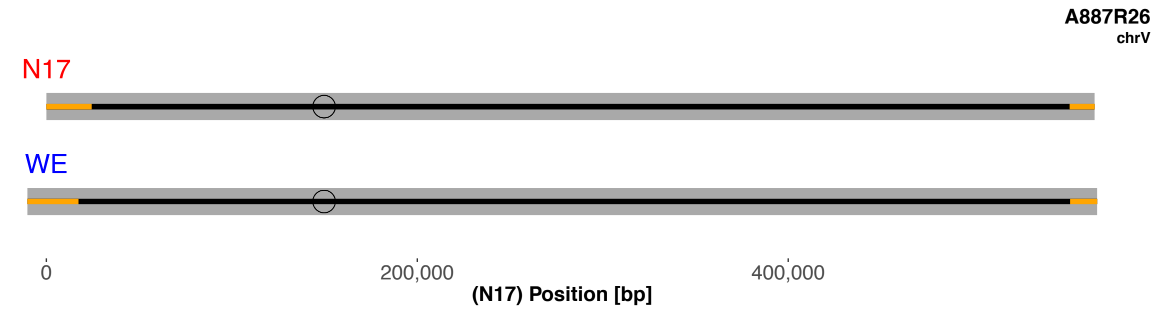
A887R26
chrV
N17
WE
0
200,000
400,000
(N17) Position [bp]

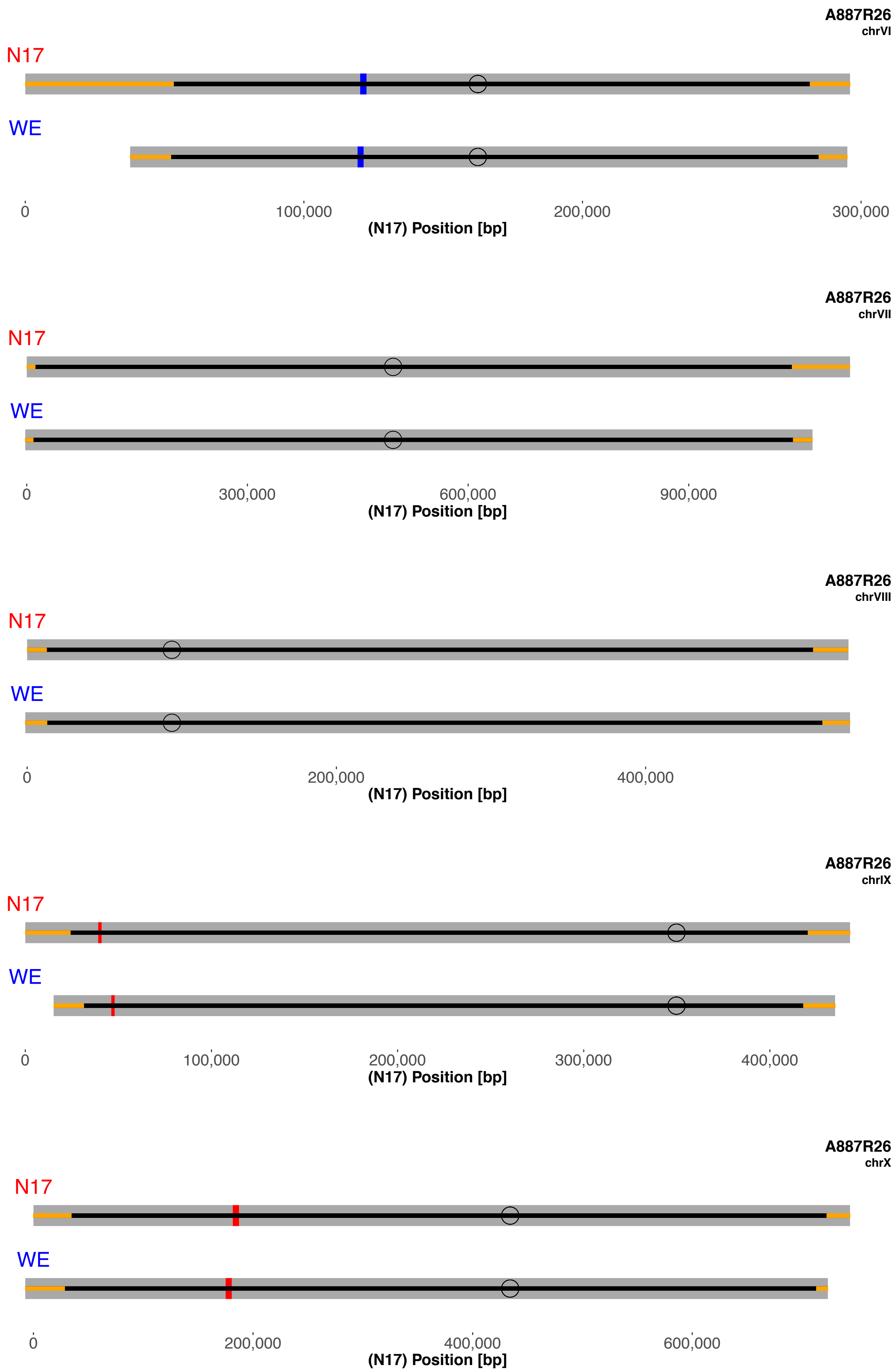
A887R26
chrVI
N17
WE
0
100,000
200,000
300,000
(N17) Position [bp]
A887R26
chrVII
N17
WE
0
300,000
600,000
900,000
(N17) Position [bp]
A887R26
chrVIII
N17
WE
0
200,000
400,000
(N17) Position [bp]
A887R26
chrIX
N17
WE
0
100,000
200,000
300,000
400,000
(N17) Position [bp]
A887R26
chrX
N17
WE
0
200,000
400,000
600,000
(N17) Position [bp]

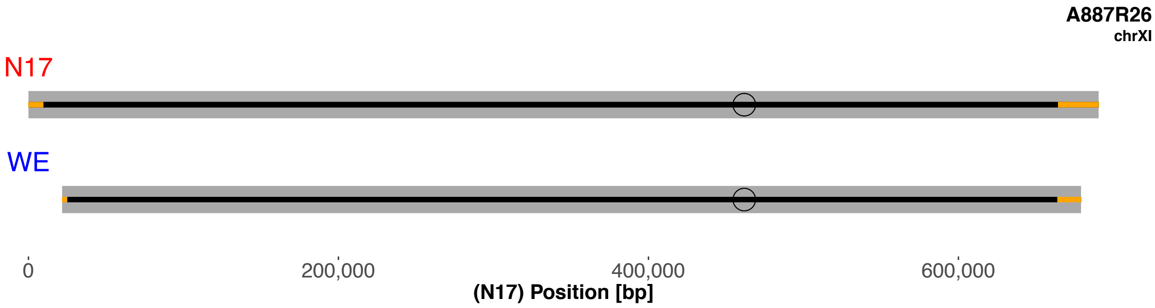
A887R26
chrXI
N17
WE
0
200,000
400,000
600,000
(N17) Position [bp]

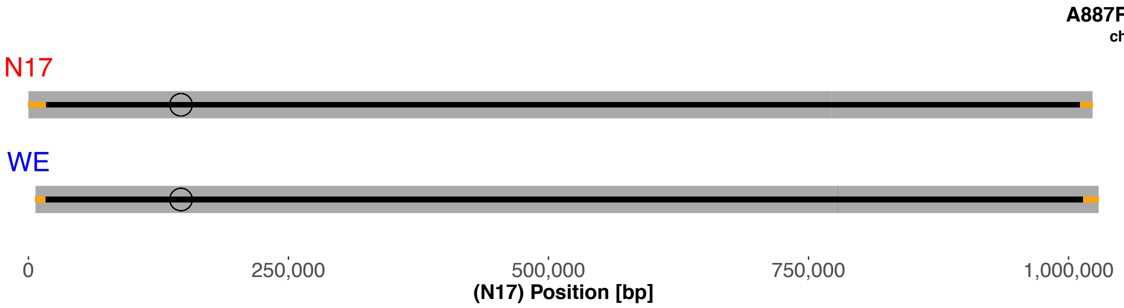
A887R26
chrXII
N17
WE
0
250,000
500,000
750,000
1,000,000
(N17) Position [bp]

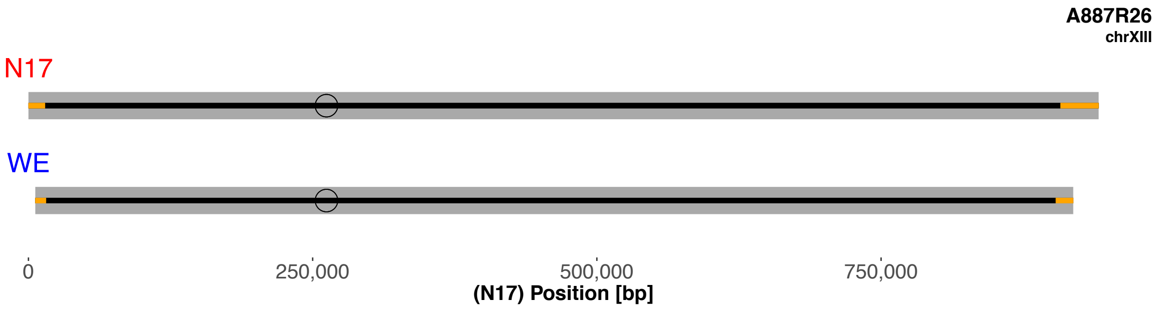
A887R26
chrXIII
N17
WE
0
250,000
500,000
750,000
(N17) Position [bp]

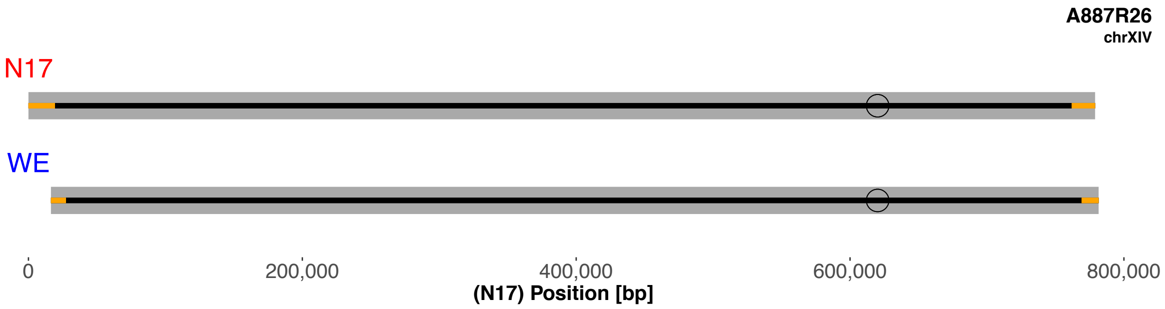
A887R26
chrXIV
N17
WE
0
200,000
400,000
600,000
800,000
(N17) Position [bp]

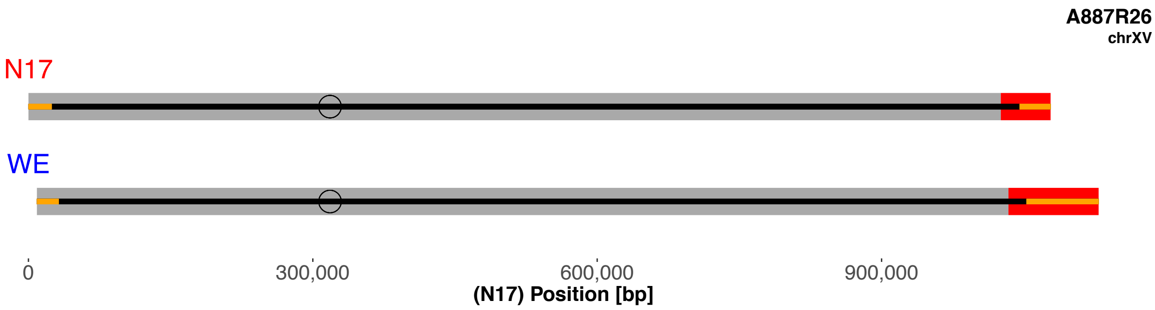
A887R26
chrXV
N17
WE
0
300,000
600,000
900,000
(N17) Position [bp]

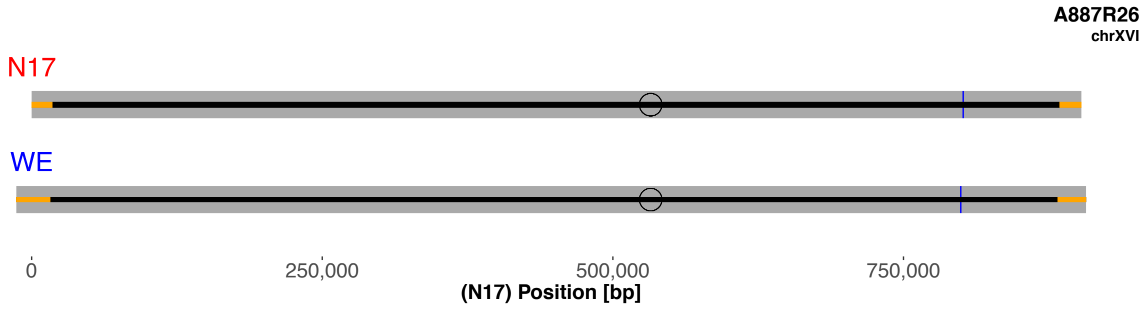
A887R26
chrXVI
N17
WE
0
250,000
500,000
750,000
(N17) Position [bp]

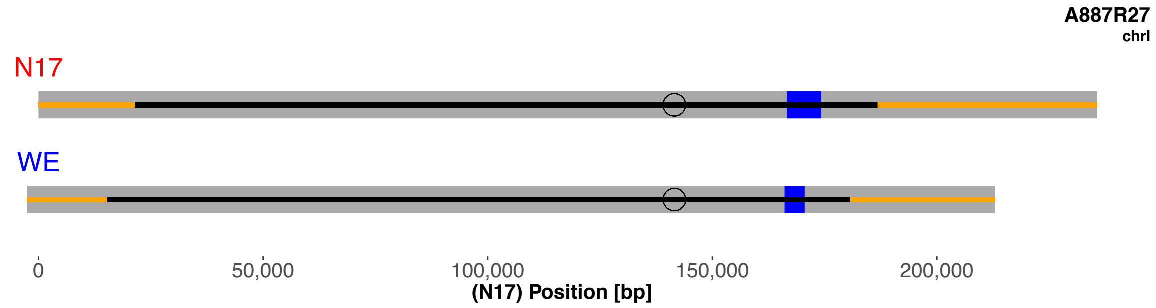
A887R27
chrI
N17
WE
0
50,000
100,000
150,000
200,000
(N17) Position [bp]

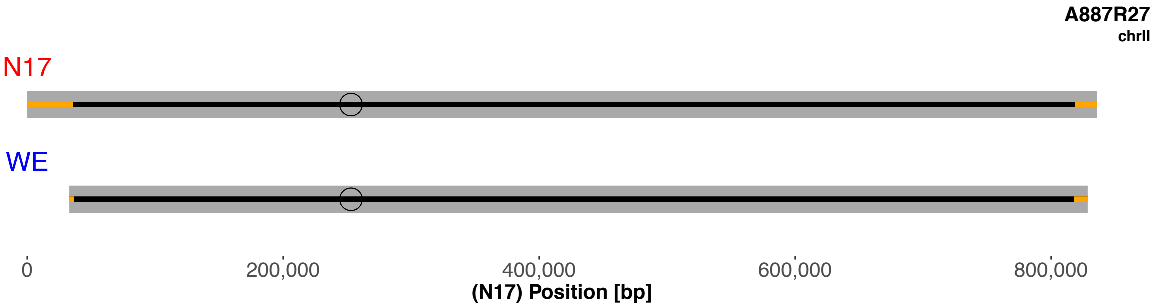
A887R27
chrII
N17
WE
0
200,000
400,000
600,000
800,000
(N17) Position [bp]

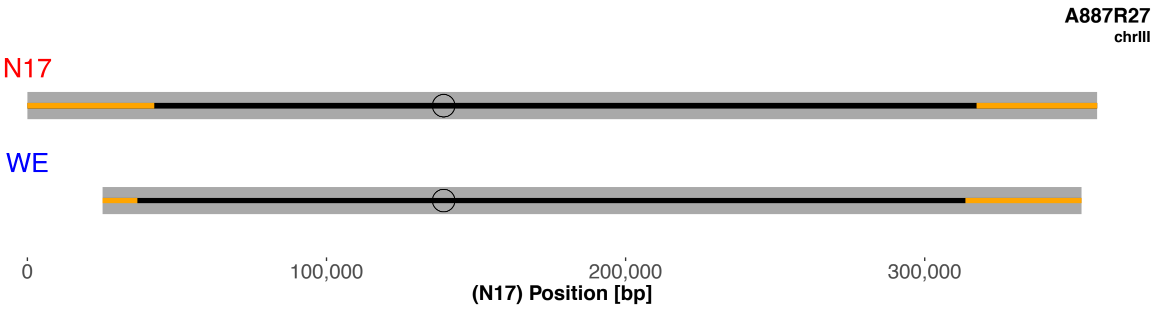
A887R27
chrIII
N17
WE
0
100,000
200,000
300,000
(N17) Position [bp]

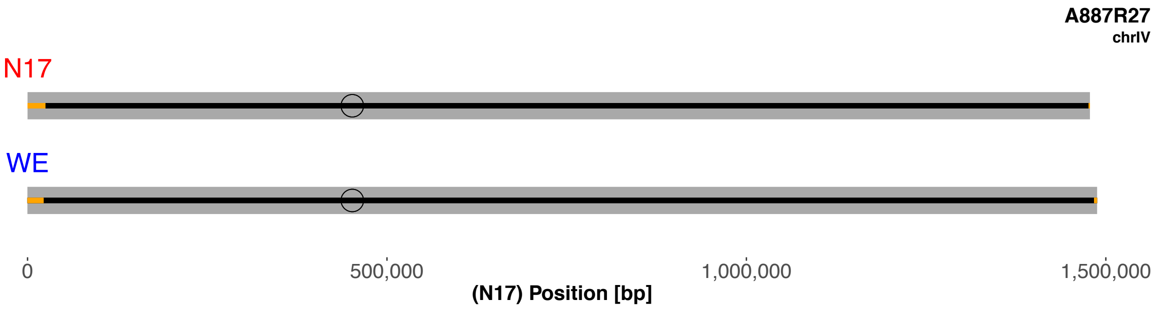
A887R27
chrIV
N17
WE
0
500,000
1,000,000
1,500,000
(N17) Position [bp]

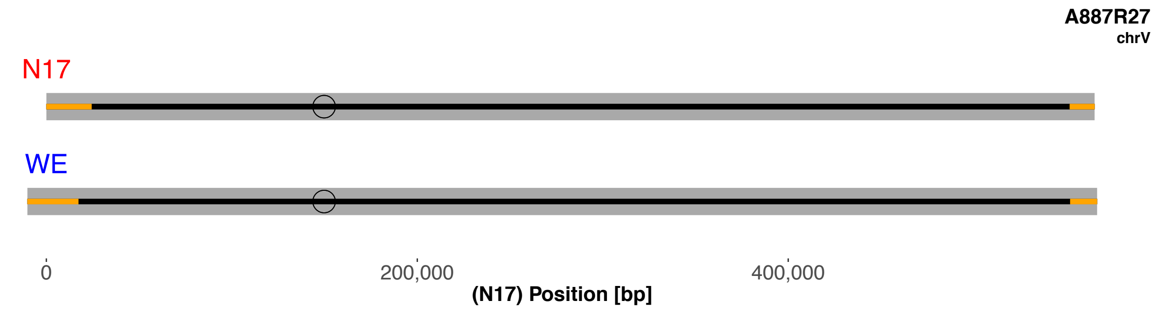
A887R27
chrV
N17
WE
0
200,000
400,000
(N17) Position [bp]

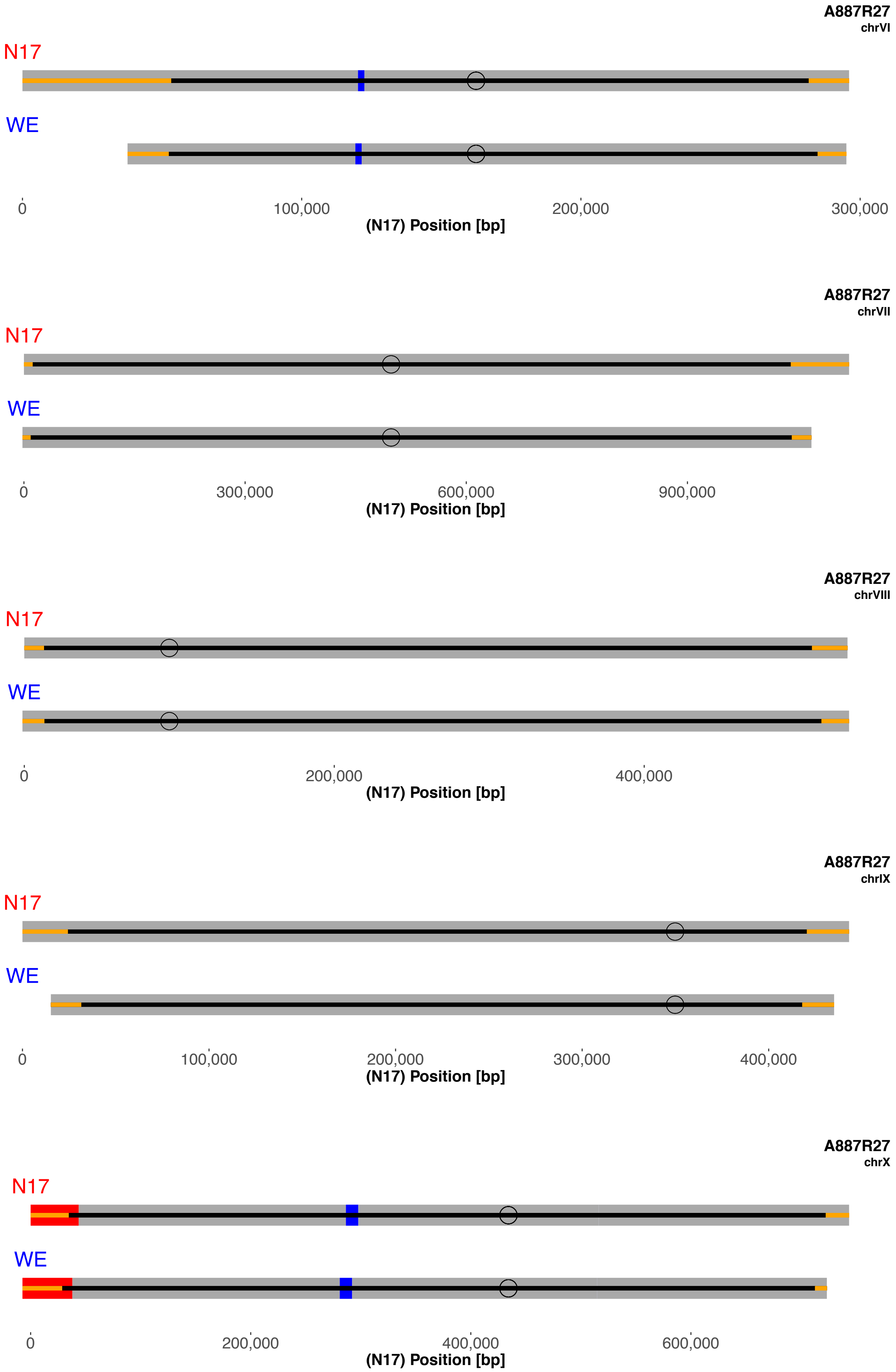
A887R27
chrVI
N17
WE
0
100,000
200,000
300,000
(N17) Position [bp]
A887R27
chrVII
N17
WE
0
300,000
600,000
900,000
(N17) Position [bp]
A887R27
chrVIII
N17
WE
0
200,000
400,000
(N17) Position [bp]
A887R27
chrIX
N17
WE
0
100,000
200,000
300,000
400,000
(N17) Position [bp]
A887R27
chrX
N17
WE
0
200,000
400,000
600,000
(N17) Position [bp]

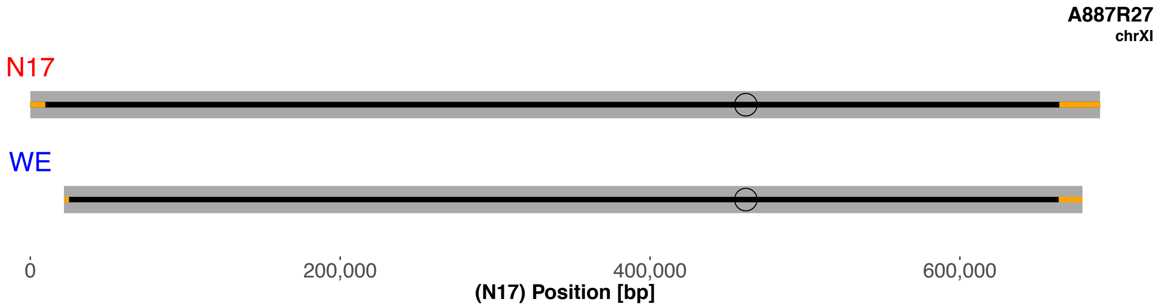
A887R27
chrXI
N17
WE
0
200,000
400,000
600,000
(N17) Position [bp]

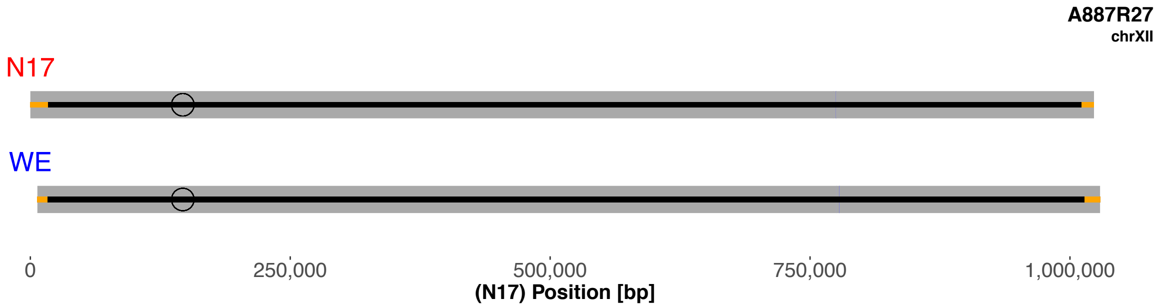
A887R27
chrXII
N17
WE
0
250,000
500,000
750,000
1,000,000
(N17) Position [bp]

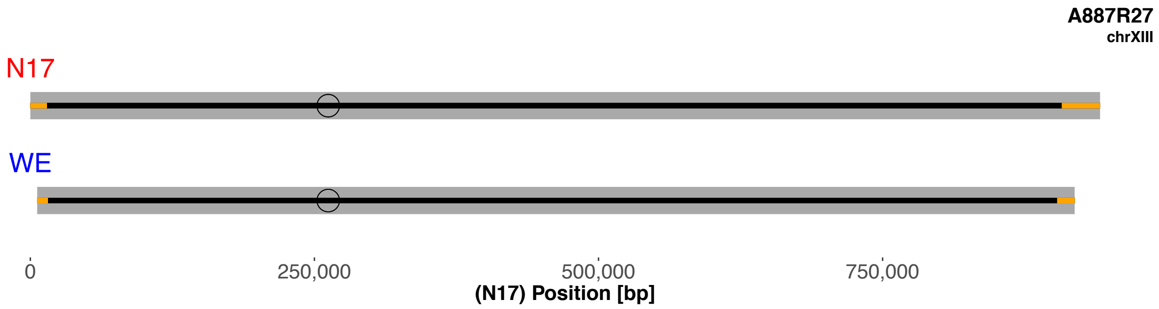
A887R27
chrXIII
N17
WE
0
250,000
500,000
750,000
(N17) Position [bp]

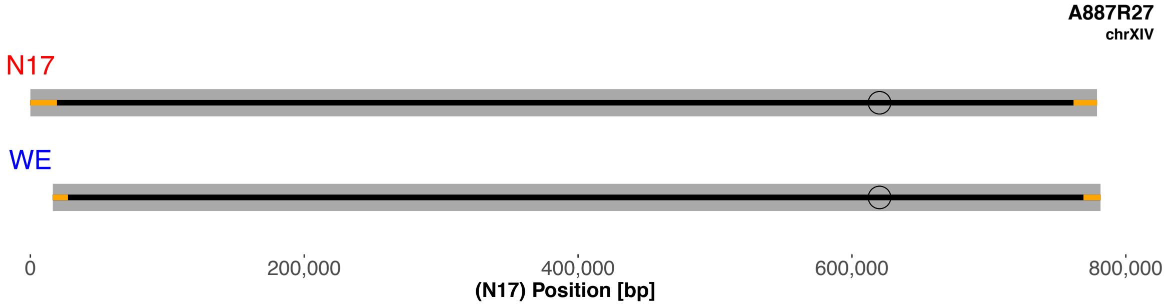
A887R27
chrXIV
N17
WE
0
200,000
400,000
600,000
800,000
(N17) Position [bp]

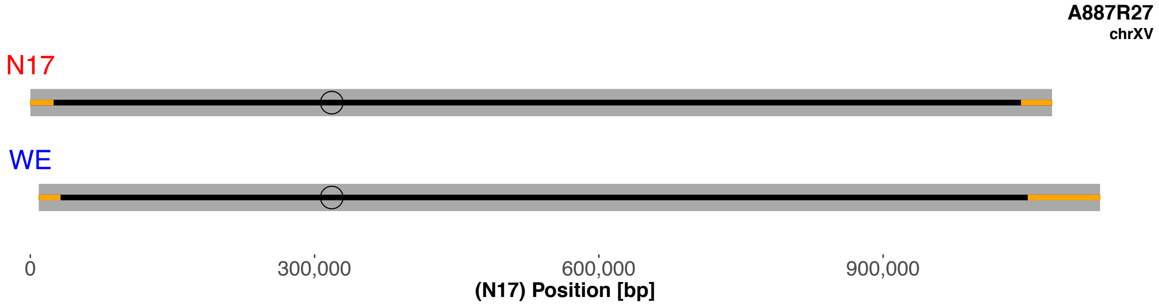
A887R27
chrXV
N17
WE
0
300,000
600,000
900,000
(N17) Position [bp]

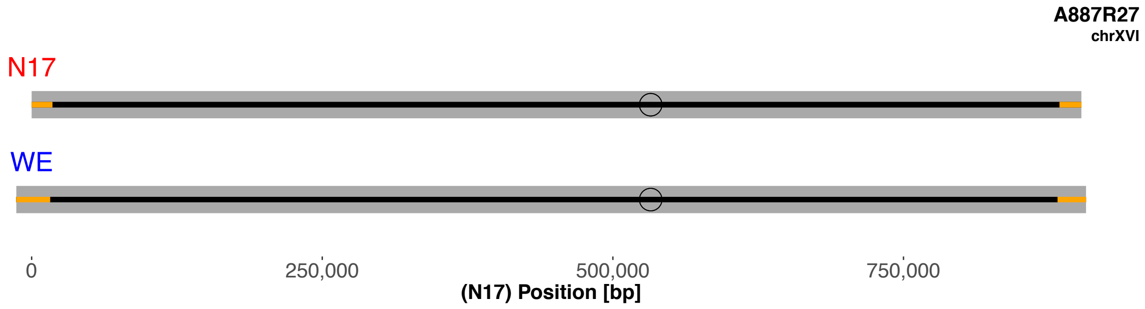

A887R27
chrXVI
N17
WE
0
250,000
500,000
750,000
(N17) Position [bp]
